# A configuration-resolved benchmark of differential abundance analysis methods for human gut 16S rRNA microbiome data

**DOI:** 10.64898/2026.09.01.748571

**Authors:** Nader Abdelmalek, Carola Di Meo, Giovanni Fiorito, Sergio Uzzau, Alessandro Tanca

**Affiliations:** Unit of Microbiology and Virology, University Hospital of Sassari, Sassari, Italy; Clinical Bioinformatics Unit, IRCCS Istituto Giannina Gaslini, Genoa, Italy; Department of Biomedical Sciences, University of Sassari, Sassari, Italy

**Keywords:** 16S rRNA, differential abundance, benchmarking, normalisation, compositional data, false discovery rate

## Abstract

Differential abundance (DA) analysis decides which taxa a 16S rRNA study reports. Each DA tool runs under choices of normalisation, transformation, zero handling and reference, yet benchmarks usually compare tools at their defaults. We benchmarked 11 tools in 43 configurations on 1,064 datasets, with signals implanted into healthy stool and inflammatory bowel disease (IBD) mucosal profiles or data simulated from the stool profiles, and in two real cohorts with permuted labels. Designs varied sample size, effect size, signal structure, confounding and read depth. The study design explained most of the variation in the Matthews correlation coefficient (MCC), followed by the tool and, much less, by its configuration. A few settings changed error control as much as the choice of tool, including the sensitivity analysis of ANCOM-BC2, median comparison in MaAsLin 3 and zero handling in LinDA. Linear models, mostly on log scales with a small pseudo-count, gave the highest MCC at an empirical false discovery rate (FDR) near the nominal level. Empirical FDR exceeded the nominal level for negative binomial count models, and for methods testing proportions in larger studies and under shifts in one direction. Fixed pseudo-counts on log-ratio scales failed when read depth differed about 1.35-fold between groups, and references assuming most taxa unchanged failed when most taxa shifted. The ranking held across taxonomic levels, communities and simulations, and agreed with how liberal each configuration was on permuted sex and IBD labels. We derive recommendations for gut 16S studies and a permutation check applicable to any dataset.

**Importance:** Microbiome studies usually identify disease-associated bacteria with a single tool run at its default settings. Yet the same data analysed with different tools, or with different settings of the same tool, can point to different bacteria. A false association can then enter follow-up studies, biomarker panels and clinical decisions, and results of different studies cannot be compared. Benchmarks that test tools only at their defaults cannot tell researchers which of these choices matter. By testing the settings themselves, this work shows which choices decide the reliability of a result and when common choices produce false findings. These results support condition-specific recommendations for choosing, checking and reporting a differential abundance analysis that can prevent false conclusions in clinical microbiome research.

## Introduction

16S rRNA amplicon sequencing remains the most widely used census of microbial communities in clinical and population studies (1, 2). 16S rRNA datasets can differ in cohort-related features, such as sparsity, inter-individual variability and the proportion of taxa that change between groups (3), as well as in study design characteristics, such as sample size, sequencing depth and confounders. Almost every such study ends with a differential abundance (DA) test, which ultimately determines which taxa are reported.

DA methods have multiplied. Early analyses relied on rarefaction and proportion tests (4). Count models from RNA sequencing followed, such as edgeR (5) with trimmed mean of M-values (TMM) normalisation (6), DESeq2 (7) and the weighted linear model limma-voom (8). Methods that address compositionality (9, 10) include ALDEx2 (11), ANCOM (12), its bias-corrected extension ANCOM-BC2 (13) and LinDA (14). ZicoSeq (15) and the linear decomposition model LDM (16) test by permutation, while MaAsLin 2 (17) and MaAsLin 3 (18) fit covariate-adjusted models. Each tool makes its own assumptions about normalisation, transformation and zeros (19) and exposes them as settings, which turns each tool into a family of analyses.

In practice, a study usually runs one tool at its default and reports the result. Benchmarks have ranked tools and shown that they return different sets of significant taxa (15, 20–23). Several tools fail to control the false discovery rate (FDR) in sparse data (21, 22, 24), and signals implanted into real profiles give more realistic data than parametric simulation (25). These benchmarks primarily compared tools using their default configurations. The extent to which choosing different configurations within the same tool affects the conclusions has not been quantified systematically. It is also unclear how performance scales with sample size and effect size, when error control fails and whether conclusions hold across data-generating models and in real cohorts. Without this knowledge, a study cannot tell whether its reported taxa reflect the biology or the settings. Answering these questions requires varying the design factors systematically (26, 27).

Here we evaluate 43 configurations of 11 DA tools. Signals were implanted into Human Microbiome Project (HMP) stool profiles of healthy adults at genus and operational taxonomic unit (OTU) level and into mucosal profiles of patients with inflammatory bowel disease (IBD). The same stool profiles served to simulate data with SparseDOSSA 2 (28). In addition, two real cohorts, compared by sex and by IBD status, tested the configurations on permuted labels. We quantify how much each tool and its configuration contribute to performance, identify the conditions under which error control fails and derive a guide for choosing and reporting a configuration.

## Results and Discussion

### Benchmark design

The benchmark tests 43 configurations of 11 tools in two-group comparisons (Figure 1, Table S1). Signals were implanted into real HMP profiles, thereby preserving the original structure of the surrounding microbial community (25). Stool of healthy adults was analysed at genus and OTU level (Figure 1A), after removing genera and OTUs present in fewer than 5% and 10% of the samples, respectively. Mucosal biopsies of patients with IBD were analysed at genus level only. Genus-level data were also simulated with SparseDOSSA 2 fitted to the stool genus table (28). A joint signal affected both the abundance of a taxon in the samples that carry it and its prevalence, whereas a prevalence-only signal affected prevalence alone. Signals were placed at random on features present in at least 10% of the samples of a dataset. The main design crossed 50, 100 and 200 samples, effect sizes of 1.0 to 2.5 on the natural-log scale and both signal types, and each other design changed one of its factors (Figure 1B). A scenario is one combination of design factors and data source, and each was generated four times, giving 1,064 datasets over 266 scenarios. Performance was measured by the Matthews correlation coefficient (MCC) and the empirical FDR at a nominal FDR level of 0.05. Figure 1C lists the configurations and explains their names, in which TSS stands for total sum scaling, CLR for centred log-ratio, PLOG for pseudo-log and Median for median comparison.

**Figure 1.**
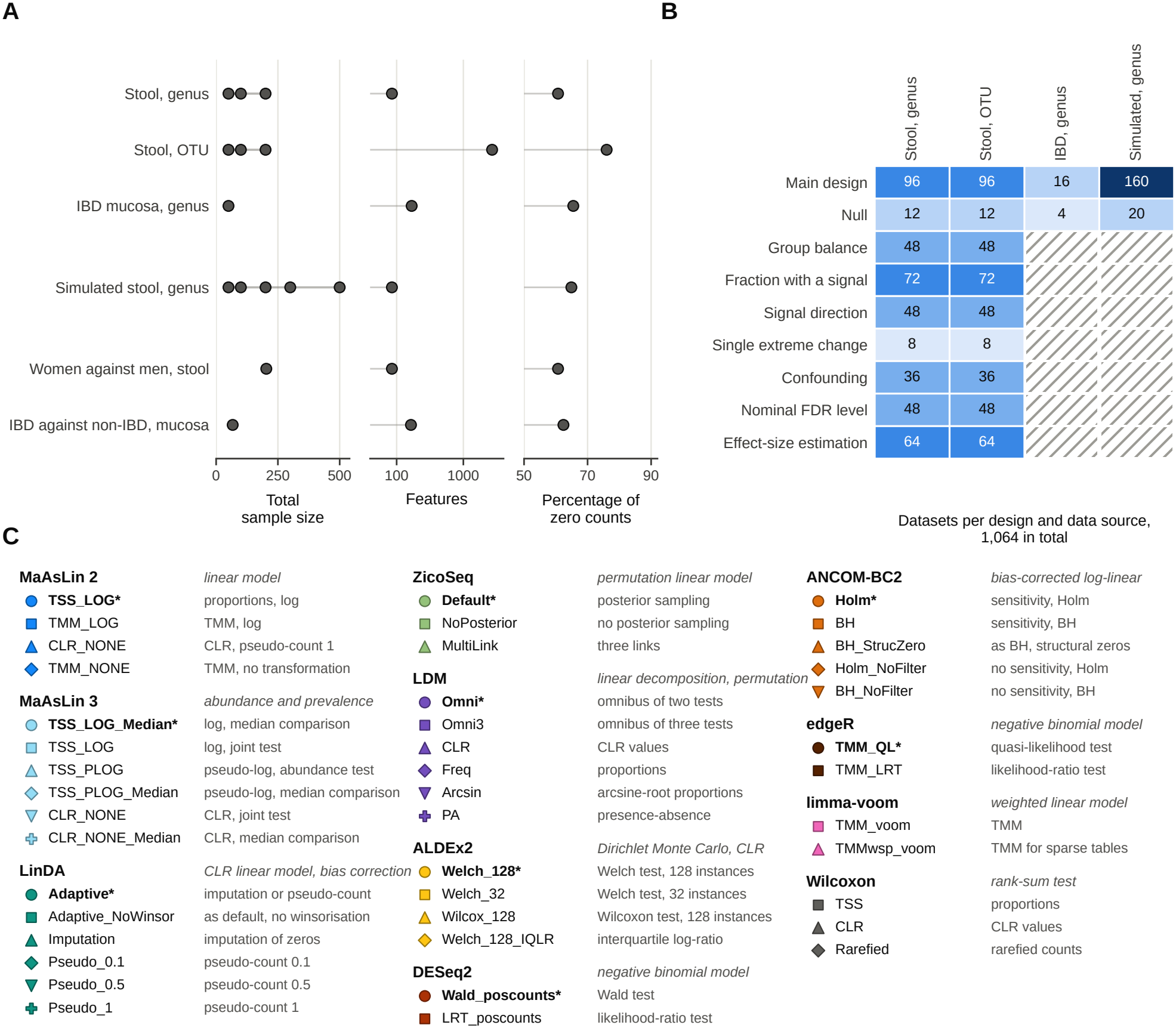
Study design. (A) Datasets of the benchmark. Signals were implanted into the stool and IBD tables, and simulated data were generated with SparseDOSSA 2 fitted to the stool genera. The real cohorts compare women with men in stool and patients with IBD with non-IBD controls in rectal biopsies. In the left section, each filled circle is one total sample size analysed, from 50 to 500. In the middle and right sections, each circle gives the number of features and the percentage of zero counts of the table. (B) Number of datasets in each design and data source, counting the four replicates of every scenario. The main design crosses sample size, effect size and signal type, and the null design implants no signal. The other designs vary one factor, namely the proportion of cases (group balance), the percentage of features with a signal, increases and decreases against increases only (signal direction), one feature increased strongly (single extreme change), a confounder correlated with the group (confounding), the nominal FDR level, and abundance-only against joint signals (effect-size estimation). Hatched cells mark combinations that were not run. (C) The 43 configurations, grouped by tool with its model family. Each configuration keeps its marker in all figures. Package defaults are shown as circles and in bold with an asterisk. Table S2 gives the full settings.

### MCC and error control of the configurations

Linear models, mostly on log scales with a small pseudo-count, gave the highest MCC (Figure 2A and B, Table S3, Text S1, section 1.1). At genus level, LinDA with a pseudo-count of 0.1 and MaAsLin 2 TMM_LOG ranked first. Nine further configurations could not be separated from the best by a paired test (grey band in Figure 2A). At OTU level, ZicoSeq without posterior sampling ranked first, followed by LinDA with a pseudo-count of 0.1 or 0.5, the MaAsLin 3 pseudo-log configurations and MaAsLin 2 TMM_LOG. Twelve further configurations could not be separated from the best. Linear models and the Wilcoxon test were shown to control false discoveries on implanted data in an earlier benchmark (25) and gave the most replicable results across real datasets (29).

**Figure 2.**
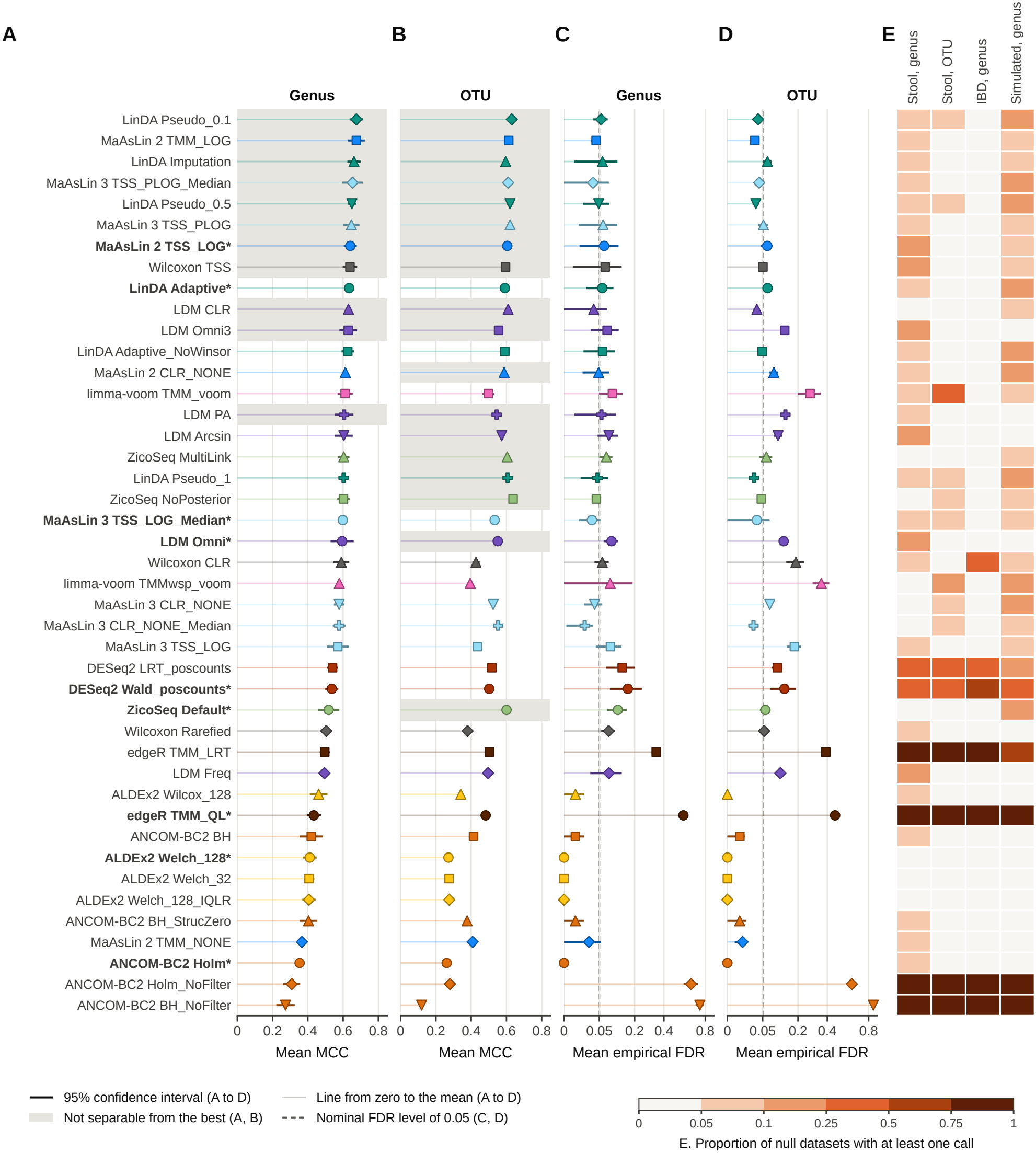
MCC and empirical FDR of the 43 configurations. Mean MCC (A, B) and mean empirical FDR on a square-root scale (C, D) in the main design at genus (A, C) and OTU level (B, D) in semi-synthetic stool data. Thick dark segments are 95% confidence intervals across four replicates, and thin pale lines lead from zero to the mean. Grey bands in A and B mark the configurations that could not be separated from the configuration with the best mean rank by a paired t test with Holm adjustment. The test depends on how consistently a configuration trails the best one across scenarios, not only on its mean MCC, which is why the bands are not contiguous. All calls were made at a nominal FDR level of 0.05, marked by the dashed line in C and D. (E) Proportion of null datasets with at least one call. Each stool column rests on 12 datasets, the IBD column on four and the simulated column on 20. In data without signal, a configuration that controls the FDR at 0.05 makes a call in at most 5% of the datasets. Configurations are ordered by mean MCC at genus level, and circles, bold labels and asterisks mark package defaults.

Error control distinguished the configurations more clearly than the MCC (Figure 2C and D, Text S1, section 1.2). At genus level, the empirical FDR of 8 configurations was below the nominal level of 0.05, that of 10 above it, and 25 could not be distinguished from it with four replicates. With 20 replicates, a check run at genus level, 13 were below, 16 above and 14 undetermined, and none moved from below to above the nominal level or back (Table S4). With four replicates, the four count models of edgeR and DESeq2 and the two ANCOM-BC2 configurations without sensitivity analysis had the highest empirical FDR. The ZicoSeq and LDM omnibus defaults, ZicoSeq MultiLink and the Wilcoxon test after rarefaction exceeded the nominal level by a small margin. Neither filtering with filterByExpr and robust fits in edgeR nor the sparse-count settings of DESeq2 without outlier replacement lowered their empirical FDR (Table S5), in line with earlier reports (21, 22, 24). In data without signal, DESeq2, edgeR and ANCOM-BC2 without its sensitivity analysis made calls in more datasets than the nominal level allows (Figure 2E, Table S6).

ALDEx2 and ANCOM-BC2 with its sensitivity analysis showed the opposite pattern. Their empirical FDR was at or near zero, but they recovered a quarter to a third of the implanted differences at genus level. Their defaults made no call in more than a third of the genus datasets (Table S3).

Calibration against the nominal level also differed between configurations (Figure S1, Table S7). LinDA with a pseudo-count of 0.5 tracked it, whereas MaAsLin 2 and MaAsLin 3 on proportions without median comparison stayed above it at every level. At the largest nominal level that kept the empirical FDR at 0.05, MaAsLin 2 TMM_LOG, LinDA with a pseudo-count of 0.5, MaAsLin 3 TSS_PLOG_Median and LDM CLR reached the highest recall (Figure S2). The nominal level should therefore be matched to the configuration.

### Contribution of design, tool and configuration

Over all scenarios, configurations and replicates of the main design, the design explained about two thirds of the variation in MCC, the tool 7% at genus and 13% at OTU level, and the configuration within tool 2% and 4% (Figure 3A, Table S8, Text S1, section 1.3). For the empirical FDR, the configuration explained as much of the variation as the tool (30% against 25% at genus level). Most of the variation explained by the configuration came from three extreme settings, MaAsLin 2 TMM_NONE and the two ANCOM-BC2 configurations without sensitivity analysis. Without these three settings, the configuration explained 3% of the variation in empirical FDR. Without the four most liberal configurations, the two of edgeR and the two of ANCOM-BC2 without sensitivity analysis, the tool still explained more of the variation in MCC than its configuration (5% against 2% at genus level). At genus level, the tool and the configuration then each explained 6% or less of the variation in empirical FDR, and most of the rest lay between replicates (Table S8). At OTU level, the gain from configuring a tool exceeded the gap between tools in about a third of the scenarios, mostly at 200 samples (Table S9).

**Figure 3.**
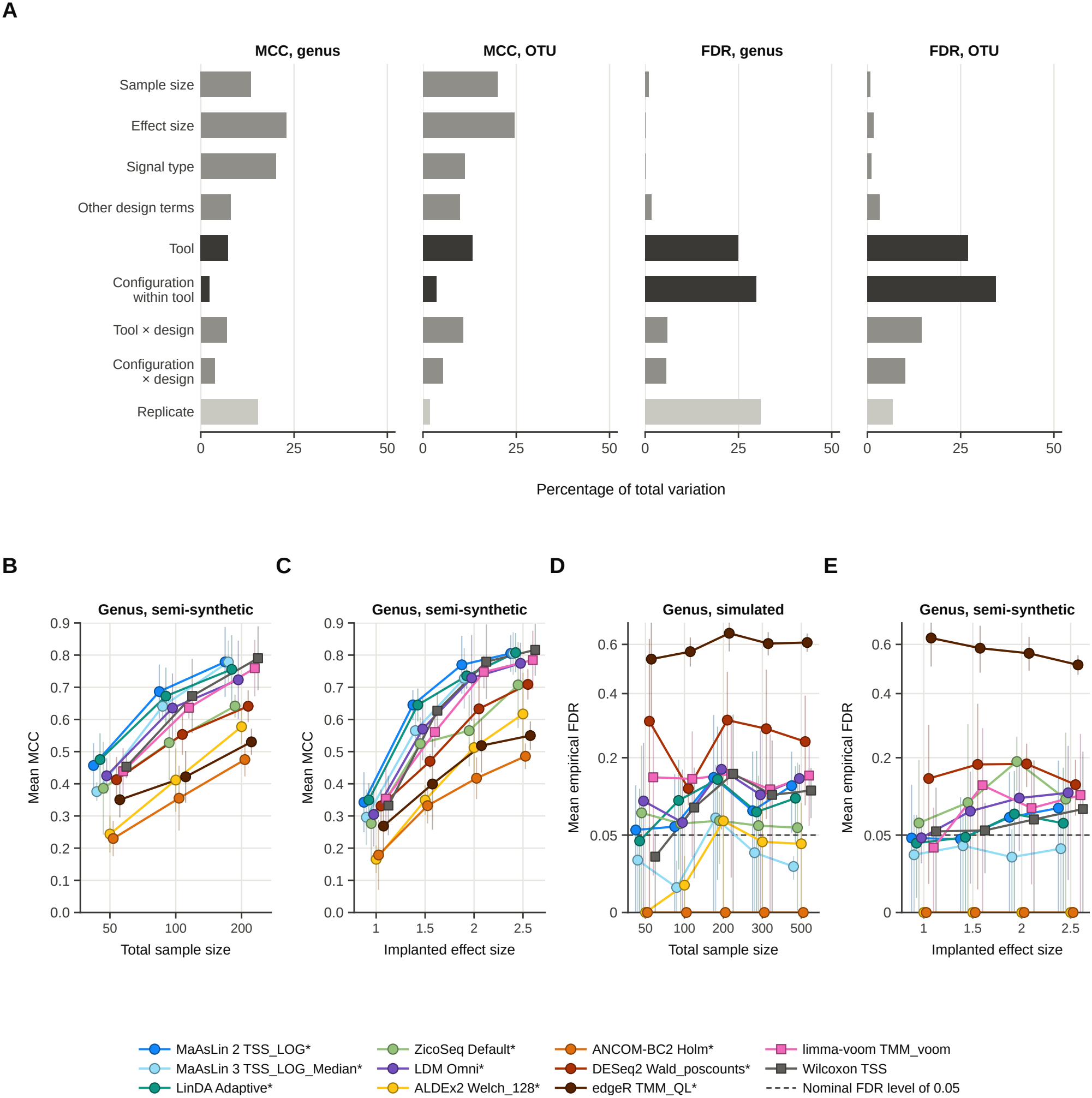
Contribution of design, tool and configuration. (A) Percentage of the variation in MCC and in empirical FDR explained by each term, computed over all scenarios, configurations and replicates of the main design at genus and OTU level. Other design terms are the interactions of sample size, effect size and signal type, and tool × design groups the interactions of the tool with these factors. Dark bars mark the tool and the configuration within tool. (B, C) Mean MCC at 50, 100 and 200 samples (B) and at implanted effect sizes of 1.0 to 2.5 (C), at genus level in semi-synthetic data. (D, E) Mean empirical FDR on a square-root scale at 50 to 500 samples at genus level in simulated data (D) and at the effect sizes of C at genus level in semi-synthetic data (E). B to E show the package default of each tool, or limma-voom with TMM normalisation and the Wilcoxon test on proportions for the two tools without one, as means over the other factors of the main design with 95% confidence intervals across four replicates.

Five settings had the largest effect on the empirical FDR (Figure S3, Text S1, section 1.3). First, switching off the sensitivity analysis of ANCOM-BC2 moved it from the lowest empirical FDR in the benchmark to the highest. Second, median comparison in MaAsLin 3, which tests each abundance coefficient against the median coefficient (18), protected the log and CLR models against an inflated FDR. Third, posterior sampling in ZicoSeq lowered the MCC and raised the empirical FDR. Fourth, LDM on CLR values stayed near the nominal level, whereas the LDM omnibus default exceeded it. Fifth, in LinDA, imputation raised the empirical FDR wherever read depth followed the signal, whereas the value of a fixed pseudo-count mainly changed recall (Table S10).

Of the package defaults, only that of MaAsLin 2 at genus level (grey band in Figure 2A) and those of MaAsLin 2, LDM and ZicoSeq at OTU level (Figure 2B) were in the leading group. Choosing the configuration on three replicates and scoring it on the fourth improved most on the default for ZicoSeq, ANCOM-BC2 and edgeR at genus level and for ANCOM-BC2 and MaAsLin 3 at OTU level (Table S11). For ANCOM-BC2, Benjamini–Hochberg adjustment with the sensitivity analysis gave most of this gain at an empirical FDR near zero. The configuration chosen by MCC, however, often lacked the sensitivity analysis, and its empirical FDR was then far above the nominal level at both taxonomic levels (Text S1, section 1.3). Slower configurations had no higher MCC (Table S2, Text S1, section 1.1).

The MCC of every configuration increased with sample size and effect size (Figure 3B and C, Figures S4 and S5, Text S1, section 1.4). With 50 samples, the leading configurations found about a third of the implanted differences at genus level. Larger studies, however, also raised the empirical FDR of methods that test proportions. In simulated data, the empirical FDR of the Wilcoxon test on proportions and MaAsLin 2 TSS_LOG increased with sample size and stayed above the nominal level at 500 samples. That of edgeR exceeded it at every sample size (Figure 3D, Figure S6). Each implanted signal slightly changes the proportions of every other taxon, and larger studies make these relative changes significant (9, 30). Most of this increase came from a higher proportion of falsely called taxa in each dataset (Figure S7). Larger effects raised the empirical FDR in the same way (Figure 3E, Figure S8).

### Conditions under which error control fails

Error control failed under four conditions, each linked to a property of the method. These were signals in one direction, many changed taxa, unequal read depth and an omitted confounder (Figure 4, Text S1, section 1.5). Signals in one direction lower the proportions of the unchanged taxa, because proportions sum to one (9, 30). Methods that test proportions, or log ratios against a reference that includes the changed taxa, then call unchanged taxa decreased. At the read depth produced by implantation, the empirical FDR greatly exceeded the nominal level for the configurations on proportions, the CLR configurations of MaAsLin 2 and the Wilcoxon test, limma-voom, the LDM omnibus test and the count models (Table S12). The bias correction of ANCOM-BC2, median comparison in MaAsLin 3 on log or CLR values and the references of LinDA with a fixed pseudo-count and LDM CLR largely contained the inflation (14, 18, 31, 32). The reference of ZicoSeq did so at genus level only. Implantation, however, increased the read depth of the cases about twofold, which spared the unchanged taxa from dilution. At fixed depth, LinDA, LDM CLR and most other configurations exceeded the nominal level. ANCOM-BC2 with sensitivity analysis and MaAsLin 3 with median comparison on log or CLR values had the lowest empirical FDR (Table S5). Most extra calls of the configurations on proportions were real changes in relative abundance, and counting these as true left almost no false calls at fixed depth (Text S1, section 1.6).

**Figure 4.**
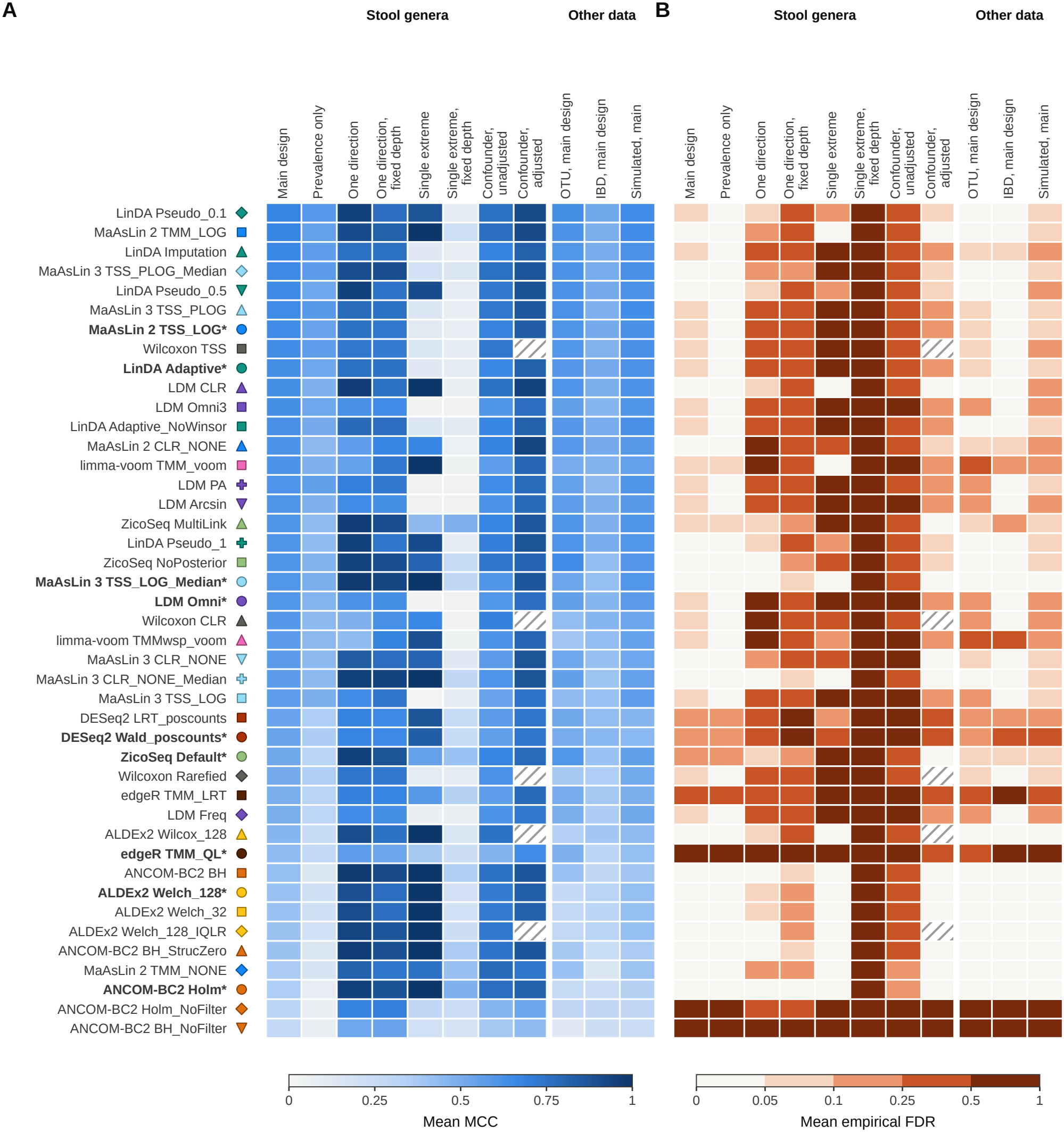
MCC and empirical FDR across conditions. Mean MCC (A) and mean empirical FDR (B) of every configuration in eleven conditions. The first eight are stool genera in semi-synthetic data, namely the main design, prevalence-only signals, signals in one direction and a single feature changed by a factor of e10, each of the last two at the read depth produced by implantation and at fixed read depth, and a confounder correlated 0.6 with the group without and with adjustment. The last three are the main design at OTU level, in the IBD community and in simulated data. Each condition is a design or data source, and its values are means over all its scenarios and replicates. The FDR colour classes break at 0.05, 0.10, 0.25 and 0.50. Hatched cells mark configurations without a covariate-adjusted form. Configurations are ordered by mean MCC in the main design at genus level.

A single bloom, in which one taxon increased by a natural-log fold change of 10, is the limiting case of a signal in one direction. At implantation depth, references insensitive to a single taxon, such as TMM trimming in linear models (6), median comparison on log or CLR values and bias correction, kept the empirical FDR near the nominal level at genus level. At fixed depth no configuration did, because the bloom truly lowers the proportion of every other taxon (Table S5).

Methods that compare each taxon with a reference estimated from the observed community, such as a geometric mean, rely on most taxa remaining unchanged (32). When many taxa shift in one direction, the reference shifts with them. With 20% of the features changed, the empirical FDR of the three ZicoSeq configurations exceeded the nominal level severalfold at OTU level, as did that of MaAsLin 3 TSS_PLOG_Median. MaAsLin 3 with median comparison on log or CLR values, LinDA with a fixed pseudo-count and LDM CLR stayed close to it (Figure S9). With 50% or more of the genera shifted, no configuration kept the empirical FDR at 0.05. At 70%, the reference moved with the shifted taxa, and the MCC turned negative in at least one direction of shift. This held for every configuration of ZicoSeq, ANCOM-BC2, LinDA and ALDEx2 and for several others, including the CLR configurations of MaAsLin 2, LDM and the Wilcoxon test (Figure S10C and D, Table S13). Such cases require a measure of total microbial load (33).

Unequal read depth between the groups produced false calls with fixed pseudo-counts on log-ratio scales (Figure S10A, Table S14). A pseudo-count of fixed size, such as 0.5, weighs more on the smaller counts of the shallower group, and taxa then appear to differ between groups that differ only in depth. When the reads of the cases were thinned, LinDA with a fixed pseudo-count, limma-voom and the Wilcoxon test on CLR values began to call taxa in null data once mean read depth differed about 1.35-fold. At 2-fold, these configurations, MaAsLin 2 CLR_NONE and LDM CLR made calls in nearly every null dataset. Imputation in LinDA, MaAsLin 3 TSS_PLOG_Median, the sensitivity analysis of ANCOM-BC2 and the Wilcoxon test on rarefied counts kept the rate near the nominal level up to 2-fold (34). Halving the reads of the cases in data with a signal gave about a 1.8-fold difference in depth. ZicoSeq without posterior sampling and MaAsLin 3 TSS_PLOG_Median then combined the highest MCC with error control near the nominal level (Figure S10B, Text S1, section 1.6).

An omitted confounder, a covariate with a correlation of 0.6 with the group, raised the empirical FDR of every configuration far above the nominal level at genus level (Figure 4, Figure S11). With adjustment, the empirical FDR was 0.085 or less at both taxonomic levels for LinDA with a fixed pseudo-count, MaAsLin 2 TMM_LOG, MaAsLin 3 with median comparison, LDM CLR, ZicoSeq without posterior sampling, ALDEx2 and ANCOM-BC2 with sensitivity analysis, and at genus level for the ZicoSeq default and MultiLink (Table S15). Unbalanced groups lowered the MCC slightly without changing which configurations led (Figure S9).

### Prevalence signals and effect-size estimation

Prevalence-only signals separated the configurations by whether their model can detect a change in presence (Figure 5A and B, Text S1, section 1.7). LinDA with a pseudo-count of 0.1, the MaAsLin 3 pseudo-log configurations, the Wilcoxon test on proportions and the LDM presence-absence test kept most of their MCC. ANCOM-BC2 and ALDEx2 lost most of it, and DESeq2 and edgeR about half. The abundance model of MaAsLin 3 on log values, unlike that on pseudo-log values, is fitted on non-zero values only and misses these signals, whereas its prevalence model and joint test detect them (Table S16).

**Figure 5.**
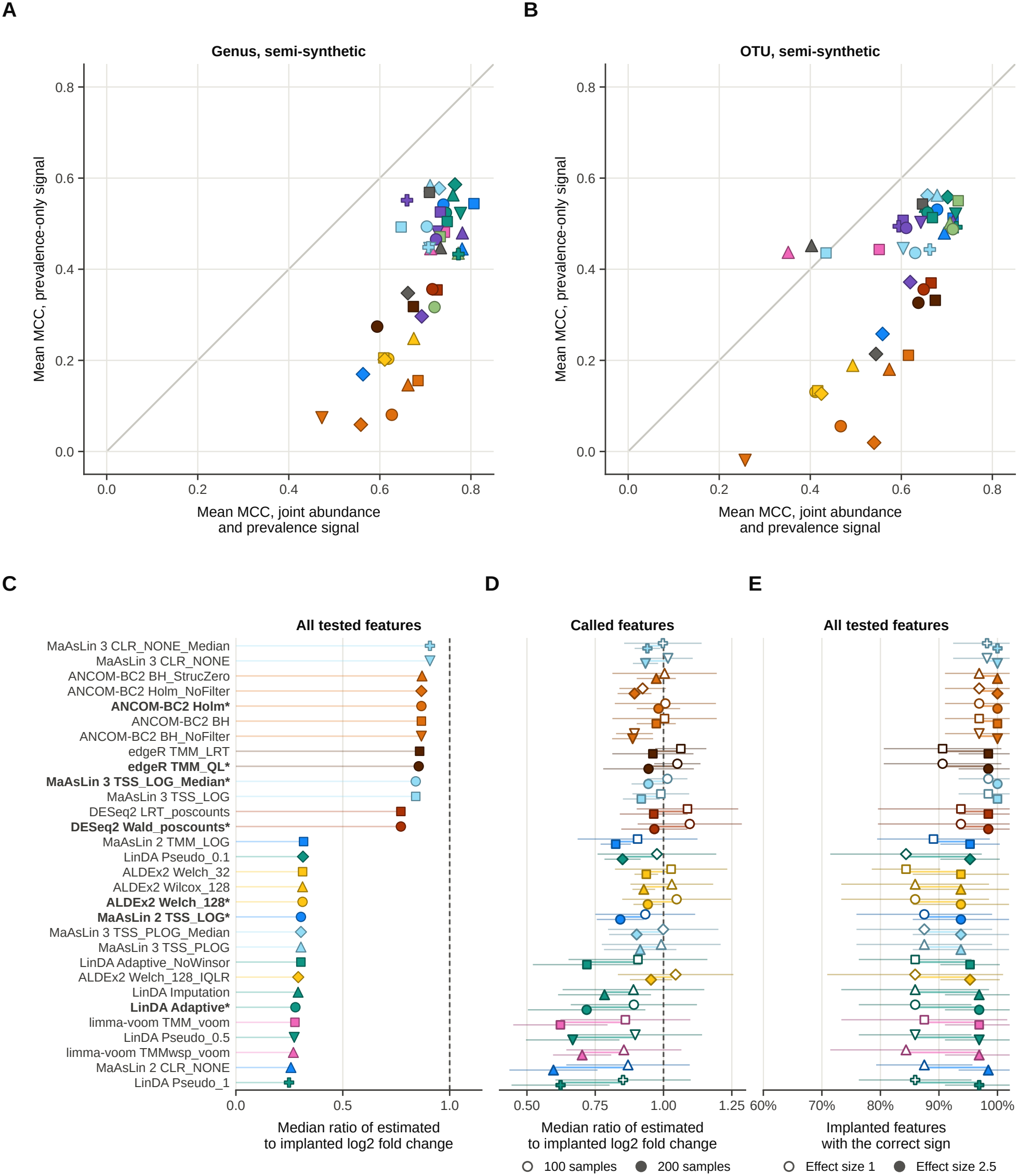
Signal type and effect-size estimation. (A, B) Mean MCC on joint signals against prevalence-only signals at genus (A) and OTU level (B) in semi-synthetic data. Both axes have the same scale, and the grey line marks equal values. (C) Median ratio of the estimated to the implanted log2 fold change over all tested implanted features under abundance-only signals at genus level, over all effect sizes at 100 and 200 samples. (D) The same ratio among the implanted features that were called, at 100 samples (open markers) and 200 samples (filled markers), with 95% confidence intervals across replicates. (E) Proportion of all tested implanted features whose estimate has the correct sign, at effect sizes of 1.0 (open markers) and 2.5 (filled markers) over 100 and 200 samples, with 95% confidence intervals. Dashed lines mark a ratio of one. ZicoSeq, LDM and the Wilcoxon test return no coefficient and are not shown in C to E.

Effect-size estimates depended on whether a model includes the zeros of a taxon (Figure 5C to E, Table S17). Under abundance-only signals, MaAsLin 3 on log or CLR values, ANCOM-BC2 and edgeR recovered most of the implanted log fold change among carriers, and DESeq2 somewhat less. An abundance-only signal multiplies the counts of the samples that carry a taxon and leaves its zeros unchanged. LinDA, MaAsLin 2 and limma-voom add a pseudo-count and estimate the change in mean log abundance over all samples, which therefore changes less, and these tools recovered less than a third of the implanted change. This reflects a different estimand, not an error, and only coefficients with the same estimand should be combined across studies. Among called features under abundance-only signals, estimates were larger, especially in small studies, because a feature passes the threshold more easily when noise enlarges its estimate (35). Under joint signals, over 99% of the called implanted features had the correct sign. Under prevalence-only signals, ANCOM-BC2 and MaAsLin 3 on log or CLR values, whose abundance models leave zeros out, gave the correct sign for only 54% to 84% (Text S1, section 1.7).

### Transfer across taxonomic levels, communities and data-generating models

The ranking of the configurations held between genus and OTU level (Figure 6A to C, Table S3, Text S1, section 1.8). MCC was lower at OTU level, mainly through recall, because OTU tables contain about 30 times more features and more zeros (36). At OTU level, the empirical FDR of the two limma-voom configurations was three to four times that at genus level in the main design. That of the three ZicoSeq configurations also increased under signals in one direction (Figure 6B and C). The IBD community lowered the MCC without reordering the configurations (Figure 6D). A configuration chosen on stool genera lost little MCC in the IBD community, at OTU level or in simulated data (Table S18).

**Figure 6.**
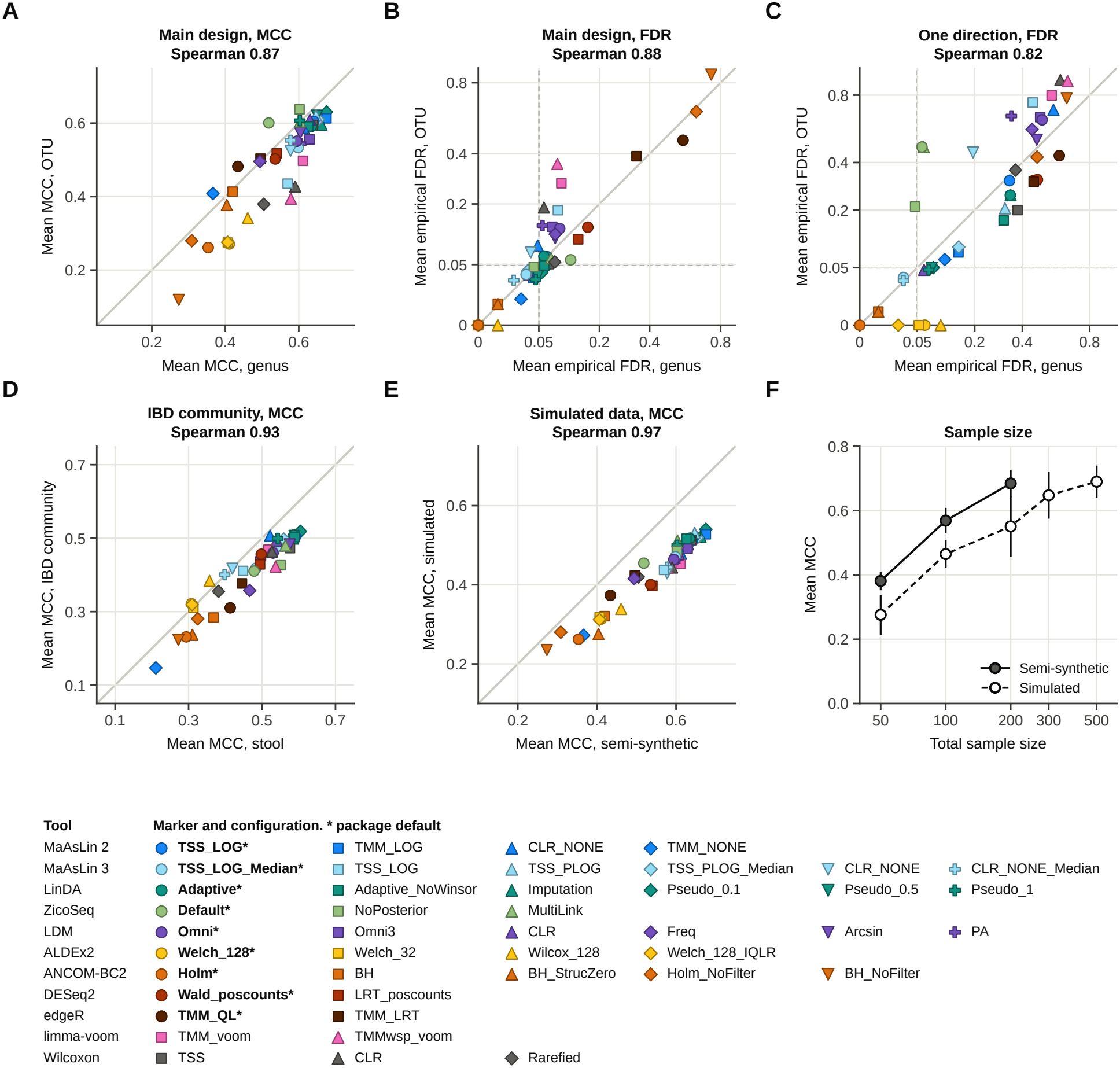
Agreement between taxonomic levels, communities and data-generating models. Each point is one configuration. (A to C) Genus against OTU level in semi-synthetic data, for mean MCC (A) and mean empirical FDR (B) in the main design and mean empirical FDR with signals in one direction (C). (D) Stool against the IBD community, for mean MCC on the four scenarios that they have in common at 50 samples. (E) Semi-synthetic against simulated data, for mean MCC on the 24 genus-level scenarios common to both approaches. The grey line marks equal values, and the Spearman correlation across the 43 configurations is given above each panel. (F) Mean MCC over the 43 configurations at each total sample size in semi-synthetic and simulated data, with 95% confidence intervals across replicates.

Simulated data ranked the configurations like the semi-synthetic data (Spearman 0.97 for MCC). LinDA with a pseudo-count of 0.1 and MaAsLin 2 TMM_LOG were among the three best in both (Figure 6E, Figure S12, Text S1, section 1.9). Simulated data gave a lower MCC and needed 500 samples to reach the MCC that semi-synthetic data reached with 200 (Figure 6F, Tables S19 and S20). SparseDOSSA 2 reproduced the prevalence, abundance and variance of each genus but not their correlations, and it lost part of each signal when drawing the reads (Figure S13, Table S21).

### Error control in real cohorts

In the two real cohorts, HMP stool profiles grouped by sex and HMP2 biopsy profiles grouped by IBD status, the labels were permuted 200 times (Figure 7, Table S22, Text S1, section 1.10). With the true sex labels, 18 configurations called no genus and 19 called a single genus, Akkermansia, in line with the few differences expected between the sexes (3). With the IBD labels, differences are expected, but with only 19 controls 34 configurations called at most one genus. Permuted labels carry no signal, and a configuration that controls the FDR at 0.05 should therefore make a call in about 5% of the permutations. edgeR, DESeq2 and the two ANCOM-BC2 configurations without sensitivity analysis made calls in about a quarter or more of the permutations in both cohorts. Both limma-voom configurations did so with the IBD labels only, and all others made calls in 13% or fewer. The proportion of permutations with a call correlated with the benchmark FDR (Spearman 0.72 for sex and 0.74 for IBD). Label permutation can therefore reveal excess calls in a dataset (37), but not compositional false discoveries, which permuting the labels removes.

**Figure 7.**
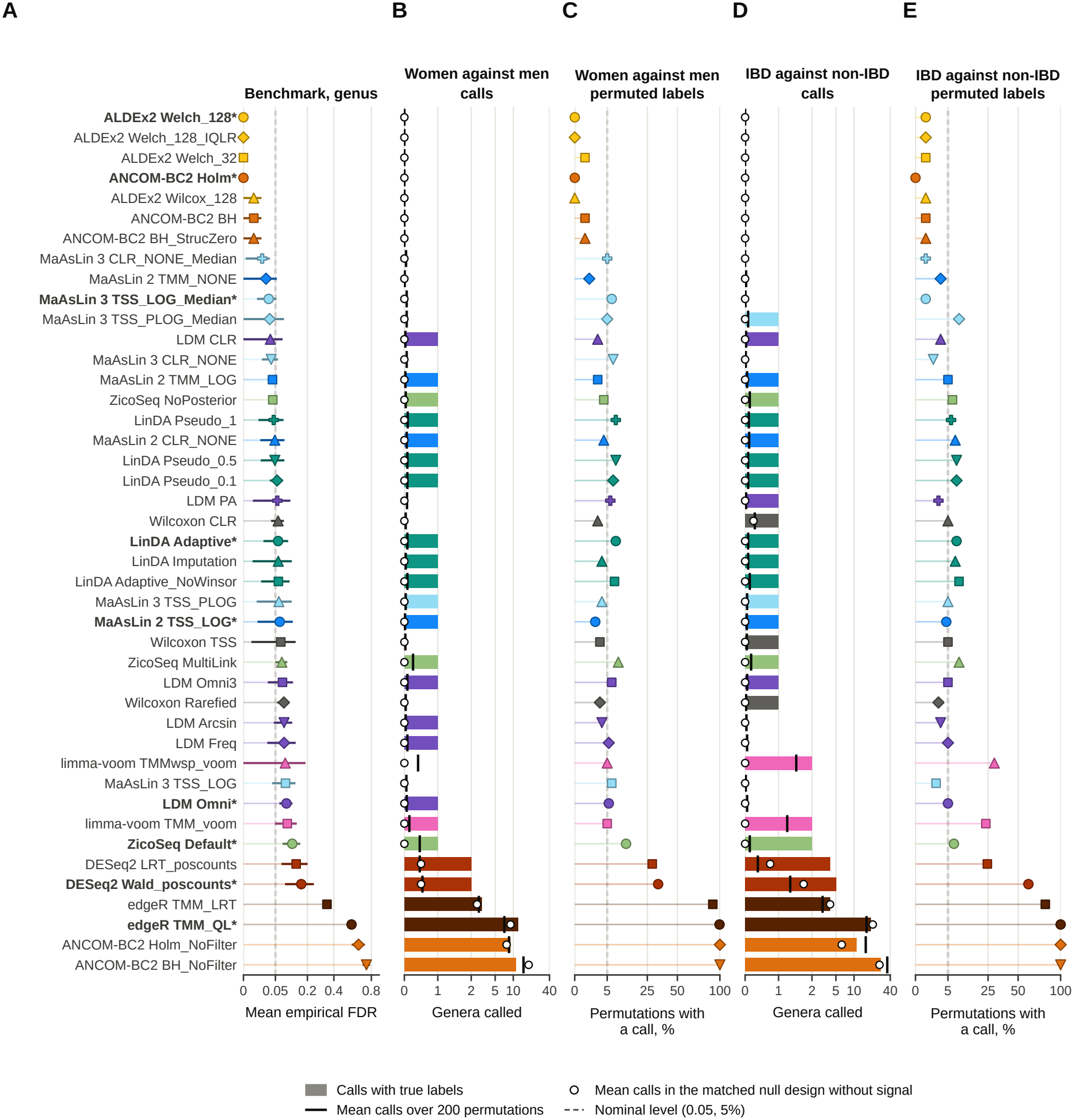
Error control in two real cohorts with permuted labels. (A) Mean empirical FDR of each configuration at genus level in semi-synthetic data, with 95% confidence intervals. (B) Genera with an adjusted p-value of 0.05 or less in the stool of 95 women and 108 men from HMP, on a symmetric logarithmic axis. Bars give the calls with the true labels, black ticks the mean over 200 permutations of the labels and open circles the mean calls in the matched null design without signal. (C) Percentage of the 200 permutations of the sex labels with at least one call, where every call is false, on a square-root axis. The dashed line marks the 5% expected for a calibrated test. (D) Genera called as in B in the rectal biopsies of 48 patients with IBD and 19 non-IBD controls from HMP2, with the IBD community at 50 samples as the matched null design. (E) Percentage of the 200 permutations of the IBD labels with at least one call, as in C.

### Limitations

The implanted signals follow a defined structure, whereas real differences may be correlated across taxa, follow phylogeny or arise from changes in total microbial load (33). Effect sizes were fixed within each scenario rather than drawn from the effects observed in a particular disease. The semi-synthetic, simulated and sex analyses derive from the same HMP stool subjects, and replicates at 200 samples reuse most of them. Their agreement therefore supports robustness but is not an external validation, which the IBD cohort provides only for error control. The confidence intervals reflect the assignment of groups and signals more than the sampling of subjects. The stool data are V3–V5 OTU profiles from 454 pyrosequencing at about 4,500 reads per sample. Deeper Illumina tables of amplicon sequence variants may change the role of zeros and the depth ratio at which a fixed pseudo-count fails. Other prevalence filters may change the results. The confounder entered the data and the adjusted model in the same linear form, the most favourable case for adjustment. Other body sites, with different community structures, need a dedicated evaluation. Multi-group, longitudinal and repeated-measures designs remain to be tested (13, 18), as do other methods, including metagenomeSeq, corncob, reference-frame approaches, phylogenetic transforms and the scale models of ALDEx2 (38–42).

## Conclusions

A DA result is underspecified when it names only the tool. A minimal report should state the tool and its version, the taxonomic level, any prevalence filter and the ratio of mean read depth between the groups. It should also give the normalisation, transformation, zero handling, reference, internal filters and sensitivity options, the test, the multiplicity procedure and nominal level, the covariates and whether a permutation check was run.

Table 1 turns the results of this benchmark into a guide for two-group comparisons of human gut 16S rRNA datasets. In the main design, the recommended configurations kept the empirical FDR close to the nominal level at both taxonomic levels. The guide rests on the order of the configurations, which held across taxonomic levels, data-generating models and the IBD community, and not on their absolute MCC.

**Table 1.** Configuration guide for two-group differential abundance analysis of human gut 16S rRNA data.

| **Situation** | **Best-performing options** | **Avoid** |
|---|---|---|
| Genus level, balanced signal, discovery | LinDA with pseudo-count 0.1 or 0.5, MaAsLin 2 TMM_LOG, MaAsLin 3 TSS_PLOG_Median, LDM CLR | edgeR, DESeq2*, ANCOM-BC2 without sensitivity analysis, MaAsLin 2 TMM_NONE, ZicoSeq*, LDM omnibus* |
| OTU level | ZicoSeq without posterior sampling, LinDA with pseudo-count 0.1 or 0.5, MaAsLin 2 TMM_LOG, MaAsLin 3 TSS_PLOG_Median, LDM CLR. With about 20% of taxa changed, prefer LinDA with a fixed pseudo-count or LDM CLR, because ZicoSeq and MaAsLin 3 TSS_PLOG_Median exceed the nominal FDR | MaAsLin 3 TSS_LOG without median comparison, limma-voom, Wilcoxon test on CLR values, LDM omnibus*, LinDA with imputation or the LinDA default*, edgeR, DESeq2* |
| Small cohort, about 50 samples | LinDA with pseudo-count 0.1, MaAsLin 2 TMM_LOG, MaAsLin 3 TSS_PLOG_Median. Treat calls as exploratory | ALDEx2* and ANCOM-BC2*, which often make no call |
| Large cohort, 300 samples or more | MaAsLin 3 with median comparison (TSS_LOG_Median*, TSS_PLOG_Median), ZicoSeq without posterior sampling, ANCOM-BC2 with sensitivity analysis | Configurations on proportions (Wilcoxon test, MaAsLin 2 TSS_LOG*, LDM omnibus*), edgeR, DESeq2*, LinDA with a fixed pseudo-count and LDM CLR (empirical FDR about 0.1) |
| Read depth differs between groups by about 1.35- to 2-fold | ZicoSeq without posterior sampling, ANCOM-BC2 with sensitivity analysis, MaAsLin 3 TSS_PLOG_Median (Table S23). Beyond 2-fold, the Wilcoxon test on rarefied counts was the most robust to depth alone (Table S14), but its empirical FDR with a signal exceeded the nominal level | Fixed pseudo-counts on log-ratio scales (LinDA with a pseudo-count, MaAsLin 2 CLR_NONE, LDM CLR, Wilcoxon test on CLR values), MaAsLin 3 TSS_LOG_Median* and CLR_NONE_Median from 2-fold, limma-voom, DESeq2*, edgeR |
| Shift in one direction of up to about a third of the taxa | MaAsLin 3 with median comparison on log or CLR values (TSS_LOG_Median*, CLR_NONE_Median), ANCOM-BC2 with sensitivity analysis, with an empirical FDR of 0.13 or less at fixed read depth | MaAsLin 2 TSS_LOG* and CLR_NONE, MaAsLin 3 without median comparison, LinDA, LDM CLR and ALDEx2*, which exceeded the nominal FDR at fixed read depth, ZicoSeq (empirical FDR about 0.11), Wilcoxon test, limma-voom, DESeq2*, edgeR, LDM omnibus* |
| Half or more of the taxa shift in one direction | No configuration kept the empirical FDR at 0.05. Measure total microbial load, or report changes in relative abundance with MaAsLin 3 TSS_PLOG, MaAsLin 2 TSS_LOG* or TMM_LOG or the Wilcoxon test on proportions, which kept a positive MCC | References that assume most taxa unchanged (ZicoSeq, MaAsLin 3 TSS_PLOG_Median, ANCOM-BC2*, LinDA, LDM CLR, ALDEx2*, CLR values in MaAsLin 2 and the Wilcoxon test), which inverted the result at 70% |
| A single taxon with a very large change (bloom) | No configuration controlled the empirical FDR at fixed read depth when a bloom took about 99.5% of the reads. Report the bloom itself and treat calls on other taxa as changes in relative abundance, or measure total microbial load | Reading the decrease of other taxa as a biological change |
| Presence or absence expected to change | LinDA with pseudo-count 0.1, MaAsLin 3 TSS_PLOG_Median, Wilcoxon test on proportions below 200 samples | ANCOM-BC2*, ALDEx2*, DESeq2*, edgeR |
| Known confounder | Adjusted models with LinDA with a fixed pseudo-count, MaAsLin 2 TMM_LOG, MaAsLin 3 with median comparison*, LDM CLR, ZicoSeq without posterior sampling, ANCOM-BC2* or ALDEx2* | Unadjusted analyses, the Wilcoxon test (no adjusted form), the LinDA default*, DESeq2*, edgeR, limma-voom |
| Confirmation with minimal false discoveries | ANCOM-BC2 with sensitivity analysis and Benjamini–Hochberg adjustment, ALDEx2* (Welch or Wilcoxon). Expect low recall below 200 samples | ANCOM-BC2 without its sensitivity analysis |
| Effect sizes for comparison across studies | Coefficients with the same estimand. MaAsLin 3 TSS_LOG_Median* or CLR_NONE_Median or ANCOM-BC2 for changes among carriers, except when only presence changes, where their sign is often wrong. Report the study size with them | MaAsLin 2 TMM_NONE, whose coefficients are not on a log scale. Coefficients of called taxa from small cohorts taken as true effects |

Each row is a study situation that draws on one or more designs. Best-performing options had the highest MCC in that situation among configurations that did not exceed the nominal FDR of 0.05 there, and they may fail in other situations. Where they exceeded the nominal FDR, the cell gives their empirical FDR. When half or more of the taxa shift or one taxon blooms, no configuration controlled the FDR, and the cell says how to report the result. Configurations to avoid exceeded the nominal FDR or lost most of their MCC, and the list gives the clearest cases. Unless stated, rows assume mean read depths differing less than about 1.35-fold between groups. The large-cohort row rests on simulated data. Asterisks mark package defaults.

Three rules apply to every study. First, the read depth of the groups should be compared before a configuration is chosen, because a fixed pseudo-count on a log-ratio scale makes false calls from about a 1.35-fold difference. Second, median comparison in MaAsLin 3 on log or CLR values and the sensitivity analysis of ANCOM-BC2 should stay active, because switching them off inflated the empirical FDR. Third, a known confounder should be included in the model. When the characteristics of a study are uncertain and read depth is similar in the two groups, MaAsLin 3 with median comparison on log or CLR values is the safest choice. It kept the empirical FDR at 0.13 or less in every design except unadjusted confounding, read depth differing 2-fold or more between groups, shifts of half the taxa or more and a single bloom at fixed depth, at a cost in recall. No configuration protects against shifts of half the taxa or a bloom, which require a measure of total microbial load.

## Materials and Methods

### Data sources

Stool profiles of healthy adults from the HMP V3–V5 16S rRNA data (43) were obtained with the HMP16SData package (44), and mucosal biopsy profiles of patients with Crohn’s disease or ulcerative colitis from the HMP2 IBD cohort (45) with the HMP2Data package (46). Samples with fewer than 1,000 reads were removed, and one sample per subject was kept. Genera present in at least 5% of all samples and OTUs present in at least 10% were retained. The stool data comprised 203 subjects, 85 genera and 2,697 OTUs, and the IBD data 56 subjects and 168 genera (Table S24, Text S1, section 2.1). The real cohorts were the 95 women and 108 men of the healthy HMP stool cohort, with the 85 genera of the stool table, and the rectal biopsies of 48 patients with IBD and 19 non-IBD controls of HMP2, with 164 genera after the same filters. Only public, de-identified data were used.

### Signal implantation and simulation

For each dataset, subjects were drawn without replacement and assigned at random to two groups. A feature could carry a signal only if it was detected in at least 10% of the drawn samples and in at least three samples. By default, 10% of the features were chosen among these, half increased and half decreased in the cases unless the design specified one direction. An abundance effect multiplied the non-zero counts of a feature in the cases by exp(±e), and a prevalence effect shifted the log-odds of its presence by ±e. A joint signal applied both. Only implanted features were true positives, and calls on any other feature, including compositional changes, counted as false. SparseDOSSA 2 (28), which generated the simulated datasets, used the same effect definitions (Text S1, sections 2.2 and 2.3).

### Designs

Secondary designs on stool used 100 and 200 samples and effect sizes of 1.5, 2.0 and 2.5. They varied group balance (30/70), the fraction of features with a signal (5%, 10% and 20%), signal direction (both directions or increases only) and the nominal FDR level (0.05, 0.10 and 0.25). A single-feature design increased one feature with an effect size of 10. A confounding design, run at 200 samples only, added a confounder, a covariate correlated 0, 0.3 or 0.6 with the group. A null design implanted no signal and measured how often a configuration makes any call. The IBD community ran four main scenarios and the null design at 50 samples. The simulated data ran the main and null designs at 50 to 500 samples, the largest two beyond the 203 stool subjects.

### Tools and configurations

The 43 configurations vary the settings that each tool exposes for normalisation, transformation, zero handling, reference, test and filtering (Table S2, Text S1, section 2.6). Each tool contributed two to six configurations (Figure 1C). The quasi-likelihood test recommended in the edgeR user guide was treated as the edgeR default. Prevalence filters inside the tools were switched off, except the 10% filter of ANCOM-BC2, and DESeq2 kept its independent filtering. The adaptive switch of LinDA, which does not take effect in MicrobiomeStat 1.4, was applied as documented (14). A feature was called when its adjusted p-value was at or below the nominal level.

### Metrics and statistical analysis

The MCC, computed per dataset from the confusion matrix of implanted against called features, was the primary metric (47, 48). It ranges from −1 to 1, with 0 for a random call set and 1 for a perfect one, and datasets without calls were scored 0 (Text S1, section 2.7). The false discovery proportion of a dataset is FP/(TP + FP), set to 0 without calls, where FP and TP are the false and true positives. Its mean over datasets is the empirical FDR, which was also divided into the proportion of datasets with a call and the FDR of the lists with calls (49). A called implanted feature counted as a true positive whatever its sign, which was scored separately. Metrics are means of the four replicate values with a 95% t interval. The empirical FDR was classed as below or above the nominal level when its interval lay at or below it or entirely above it, and as not distinguishable otherwise. The variation of MCC and FDR was decomposed into sums of squares for scenario, tool, configuration within tool, their interactions and replicates. Configurations were compared with the one of best mean rank by paired t tests with Holm adjustment (50). They were also ranked within each scenario with the Friedman test and the Nemenyi critical difference (Table S3, Text S1, section 2.8).

### Robustness analyses and real cohorts

At genus level, the main and null designs were extended to 20 replicates per scenario. The main, signal-direction, single-feature and null designs were reanalysed after every sample was resampled to its depth before implantation, and signals in one direction were also scored against a relative-abundance truth. The reads of the cases were thinned to 90% to 10% of their depth, and real stool samples were split by depth. In a further design, 10% to 70% of the genera were shifted in one direction (Text S1, section 2.9). In both real cohorts, the labels were permuted 200 times (Text S1, section 2.10).

### Implementation

The benchmark was implemented in R 4.5.2 with the package versions listed in Text S1 (section 2.11). Seeds were fixed for every scenario and replicate, and every dataset can be regenerated exactly.

## Data availability

The R script of the benchmark and all its result tables are deposited in Zenodo at https://zenodo.org/records/24505993 (version 2, DOI 10.5281/zenodo.24505993). The HMP data are available through the HMP16SData package (44) and the HMP2 IBD data through the HMP2Data package (46).

## References

1. Hamady M, Knight R. Microbial community profiling for human microbiome projects: tools, techniques, and challenges. Genome Res. 2009;19:1141–1152.

2. Gilbert JA, Blaser MJ, Caporaso JG, et al. Current understanding of the human microbiome. Nat Med. 2018;24:392–400.

3. Falony G, Joossens M, Vieira-Silva S, et al. Population-level analysis of gut microbiome variation. Science. 2016;352:560–564.

4. McMurdie PJ, Holmes S. Waste not, want not: why rarefying microbiome data is inadmissible. PLoS Comput Biol. 2014;10:e1003531.

5. Robinson MD, McCarthy DJ, Smyth GK. edgeR: a Bioconductor package for differential expression analysis of digital gene expression data. Bioinformatics. 2010;26:139–140.

6. Robinson MD, Oshlack A. A scaling normalization method for differential expression analysis of RNA-seq data. Genome Biol. 2010;11:R25.

7. Love MI, Huber W, Anders S. Moderated estimation of fold change and dispersion for RNA-seq data with DESeq2. Genome Biol. 2014;15:550.

8. Law CW, Chen Y, Shi W, Smyth GK. voom: precision weights unlock linear model analysis tools for RNA-seq read counts. Genome Biol. 2014;15:R29.

9. Gloor GB, Macklaim JM, Pawlowsky-Glahn V, Egozcue JJ. Microbiome datasets are compositional and this is not optional. Front Microbiol. 2017;8:2224.

10. Quinn TP, Erb I, Richardson MF, Crowley TM. Understanding sequencing data as compositions: an outlook and review. Bioinformatics. 2018;34:2870–2878.

11. Fernandes AD, Reid JN, Macklaim JM, et al. Unifying the analysis of high-throughput sequencing datasets: characterizing RNA-seq, 16S rRNA gene sequencing and selective growth experiments by compositional data analysis. Microbiome. 2014;2:15.

12. Mandal S, Van Treuren W, White RA, et al. Analysis of composition of microbiomes: a novel method for studying microbial composition. Microb Ecol Health Dis. 2015;26:27663.

13. Lin H, Peddada SD. Multigroup analysis of compositions of microbiomes with covariate adjustments and repeated measures. Nat Methods. 2024;21:83–91.

14. Zhou H, He K, Chen J, Zhang X. LinDA: linear models for differential abundance analysis of microbiome compositional data. Genome Biol. 2022;23:95.

15. Yang L, Chen J. A comprehensive evaluation of microbial differential abundance analysis methods. Microbiome. 2022;10:130.

16. Hu YJ, Satten GA. Testing hypotheses about the microbiome using the linear decomposition model (LDM). Bioinformatics. 2020;36:4106–4115.

17. Mallick H, Rahnavard A, McIver LJ, et al. Multivariable association discovery in population-scale meta-omics studies. PLoS Comput Biol. 2021;17:e1009442.

18. Nickols WA, Kuntz T, Shen J, et al. MaAsLin 3: refining and extending generalized multivariable linear models for meta-omic association discovery. Nat Methods. 2026;23:554–564. doi:10.1038/s41592-025-02923-9.

19. 1. Lin H, Peddada SD. Analysis of microbial compositions: a review of normalization and differential abundance analysis. npj Biofilms Microbiomes. 2020;6:60.

20. Nearing JT, Douglas GM, Hayes MG, et al. Microbiome differential abundance methods produce different results across 38 datasets. Nat Commun. 2022;13:342.

21. Calgaro M, Romualdi C, Waldron L, et al. Assessment of statistical methods from single cell, bulk RNA-seq, and metagenomics applied to microbiome data. Genome Biol. 2020;21:191.

22. Hawinkel S, Mattiello F, Bijnens L, Thas O. A broken promise: microbiome differential abundance methods do not control the false discovery rate. Brief Bioinform. 2019;20:210–221.

23. Calgaro M, Romualdi C, Risso D, Vitulo N. benchdamic: benchmarking of differential abundance methods for microbiome data. Bioinformatics. 2023;39:btac778.

24. Thorsen J, Brejnrod A, Mortensen M, et al. Large-scale benchmarking reveals false discoveries and count transformation sensitivity in 16S rRNA gene amplicon data analysis methods used in microbiome studies. Microbiome. 2016;4:62.

25. Wirbel J, Essex M, Forslund SK, Zeller G. A realistic benchmark for differential abundance testing and confounder adjustment in human microbiome studies. Genome Biol. 2024;25:247.

26. Weber LM, Saelens W, Cannoodt R, et al. Essential guidelines for computational method benchmarking. Genome Biol. 2019;20:125.

27. Costea PI, Zeller G, Sunagawa S, Bork P. A fair comparison. Nat Methods. 2014;11:359–360.

28. Ma S, Ren B, Mallick H, et al. A statistical model for describing and simulating microbial community profiles. PLoS Comput Biol. 2021;17:e1008913.

29. Pelto J, Auranen K, Kujala JV, Lahti L. Elementary methods provide more replicable results in microbial differential abundance analysis. Brief Bioinform. 2025;26:bbaf130.

30. Kumar MS, Slud EV, Okrah K, et al. Analysis and correction of compositional bias in sparse sequencing count data. BMC Genomics. 2018;19:799.

31. Lin H, Peddada SD. Analysis of compositions of microbiomes with bias correction. Nat Commun. 2020;11:3514.

32. Brill B, Amir A, Heller R. Testing for differential abundance in compositional counts data, with application to microbiome studies. Ann Appl Stat. 2022;16:2648–2671.

33. Vandeputte D, Kathagen G, D’hoe K, et al. Quantitative microbiome profiling links gut community variation to microbial load. Nature. 2017;551:507–511.

34. Schloss PD. Rarefaction is currently the best approach to control for uneven sequencing effort in amplicon sequence analyses. mSphere. 2024;9:e00354–23.

35. Ioannidis JPA. Why most discovered true associations are inflated. Epidemiology. 2008;19:640–648.

36. Weiss S, Xu ZZ, Peddada S, et al. Normalization and microbial differential abundance strategies depend upon data characteristics. Microbiome. 2017;5:27.

37. Zeng K, Fodor A. A permutation-based framework for evaluating bias in microbiome differential abundance analysis. bioRxiv. 2026. doi:10.64898/2026.03.14.711836 (preprint).

38. Paulson JN, Stine OC, Bravo HC, Pop M. Differential abundance analysis for microbial marker-gene surveys. Nat Methods. 2013;10:1200–1202.

39. 1. Martin BD, Witten D, Willis AD. Modeling microbial abundances and dysbiosis with beta-binomial regression. Ann Appl Stat. 2020;14:94–115.

40. Morton JT, Marotz C, Washburne A, et al. Establishing microbial composition measurement standards with reference frames. Nat Commun. 2019;10:2719.

41. Silverman JD, Washburne AD, Mukherjee S, David LA. A phylogenetic transform enhances analysis of compositional microbiota data. eLife. 2017;6:e21887.

42. Nixon MP, Gloor GB, Silverman JD. Incorporating scale uncertainty in microbiome and gene expression analysis as an extension of normalization. Genome Biol. 2025;26:139.

43. Human Microbiome Project Consortium. Structure, function and diversity of the healthy human microbiome. Nature. 2012;486:207–214.

44. Schiffer L, Azhar R, Shepherd L, et al. HMP16SData: efficient access to the Human Microbiome Project through Bioconductor. Am J Epidemiol. 2019;188:1023–1026.

45. Lloyd-Price J, Arze C, Ananthakrishnan AN, et al. Multi-omics of the gut microbial ecosystem in inflammatory bowel diseases. Nature. 2019;569:655–662.

46. Stansfield J, Smirnova E, Zhao N, et al. HMP2Data: 16s rRNA sequencing data from the Human Microbiome Project 2. R package version 0.99.4. https://github.com/dozmorovlab/HMP2Data. doi:10.18129/B9.bioc.HMP2Data.

47. Matthews BW. Comparison of the predicted and observed secondary structure of T4 phage lysozyme. Biochim Biophys Acta. 1975;405:442–451.

48. Chicco D, Jurman G. The advantages of the Matthews correlation coefficient (MCC) over F1 score and accuracy in binary classification evaluation. BMC Genomics. 2020;21:6.

49. Storey JD. A direct approach to false discovery rates. J R Stat Soc B. 2002;64:479–498.

50. Holm S. A simple sequentially rejective multiple test procedure. Scand J Stat. 1979;6:65–70.

