## Supplementary Figures for "A configuration-resolved benchmark of differential abundance analysis methods for human gut 16S rRNA microbiome data"

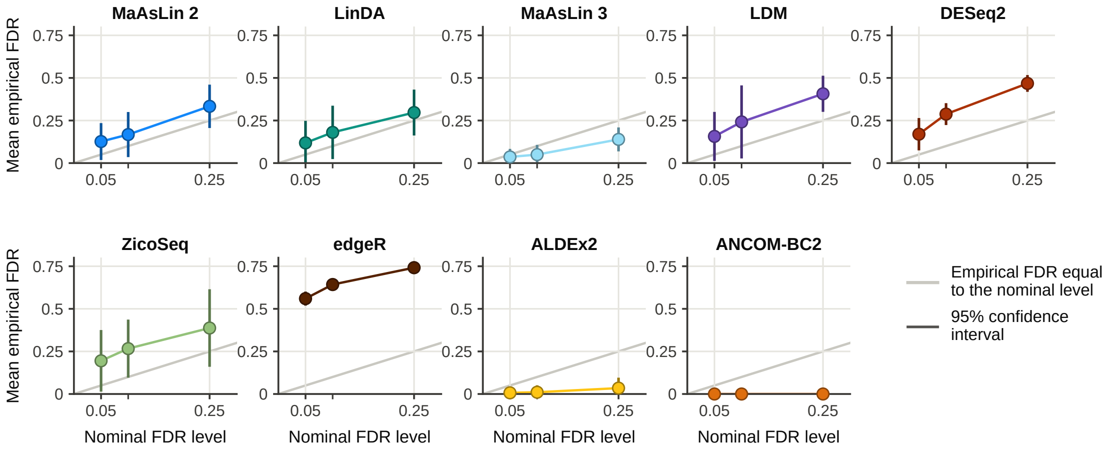


**Figure S1. Calibration of the package defaults.** Mean empirical FDR of the package default of each tool at nominal FDR levels of 0.05, 0.10 and 0.25 in the threshold design (genus level, 100 and 200 samples, effect sizes of 1.5 to 2.5, both signal types), with every tool refitted at each level. Bars are 95% confidence intervals across replicates, and the grey line marks an empirical FDR equal to the nominal level. Because this design excludes 50 samples and an effect size of 1.0, its FDR at 0.05 is higher than in the main design.

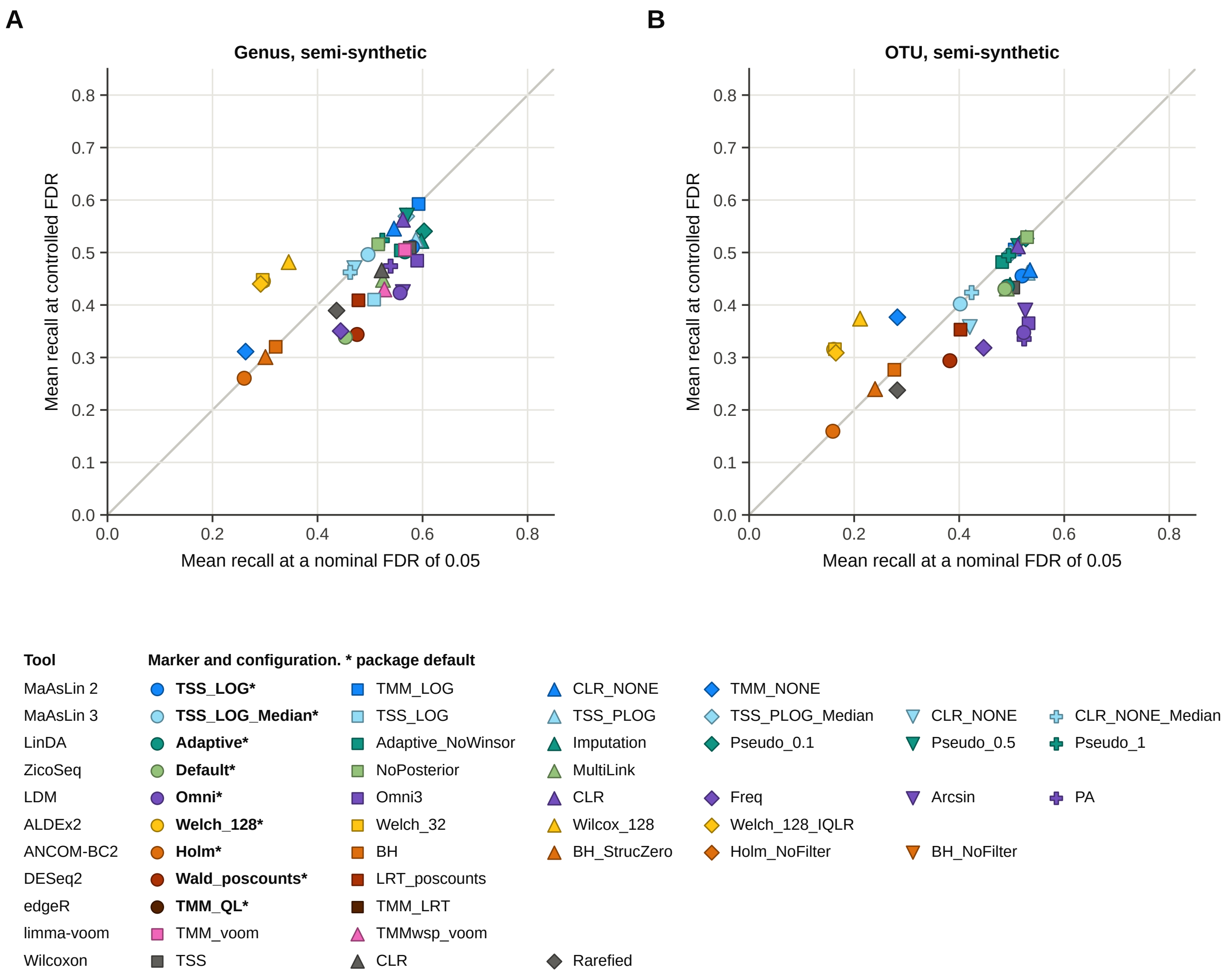


**Figure S2. Recall at controlled FDR.** Mean recall at a nominal FDR of 0.05 against the mean recall at the largest nominal level at which the empirical FDR stayed at or below 0.05, at genus (A) and OTU level (B) in semi-synthetic data. Configurations above the grey line gain recall when the threshold is relaxed to this level, and those below it need a stricter threshold to keep the FDR at 0.05.

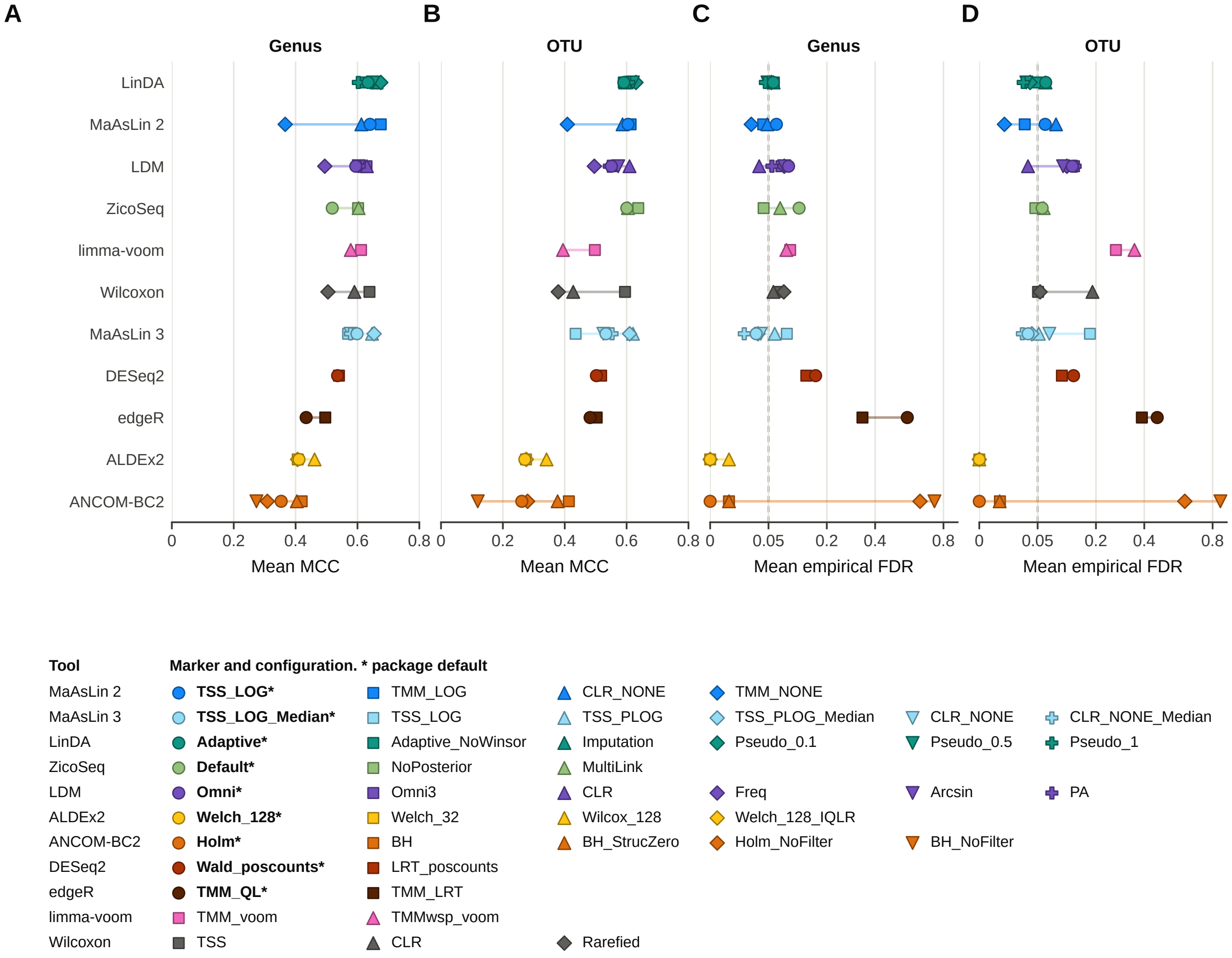


**Figure S3. Every configuration of each tool.** Mean MCC at genus (A) and OTU level (B) and mean empirical FDR on a square-root scale at genus (C) and OTU level (D) of every configuration in the main design in semi-synthetic data. Each row is one tool, and the line spans its configurations. The dashed line marks the nominal FDR level of 0.05.

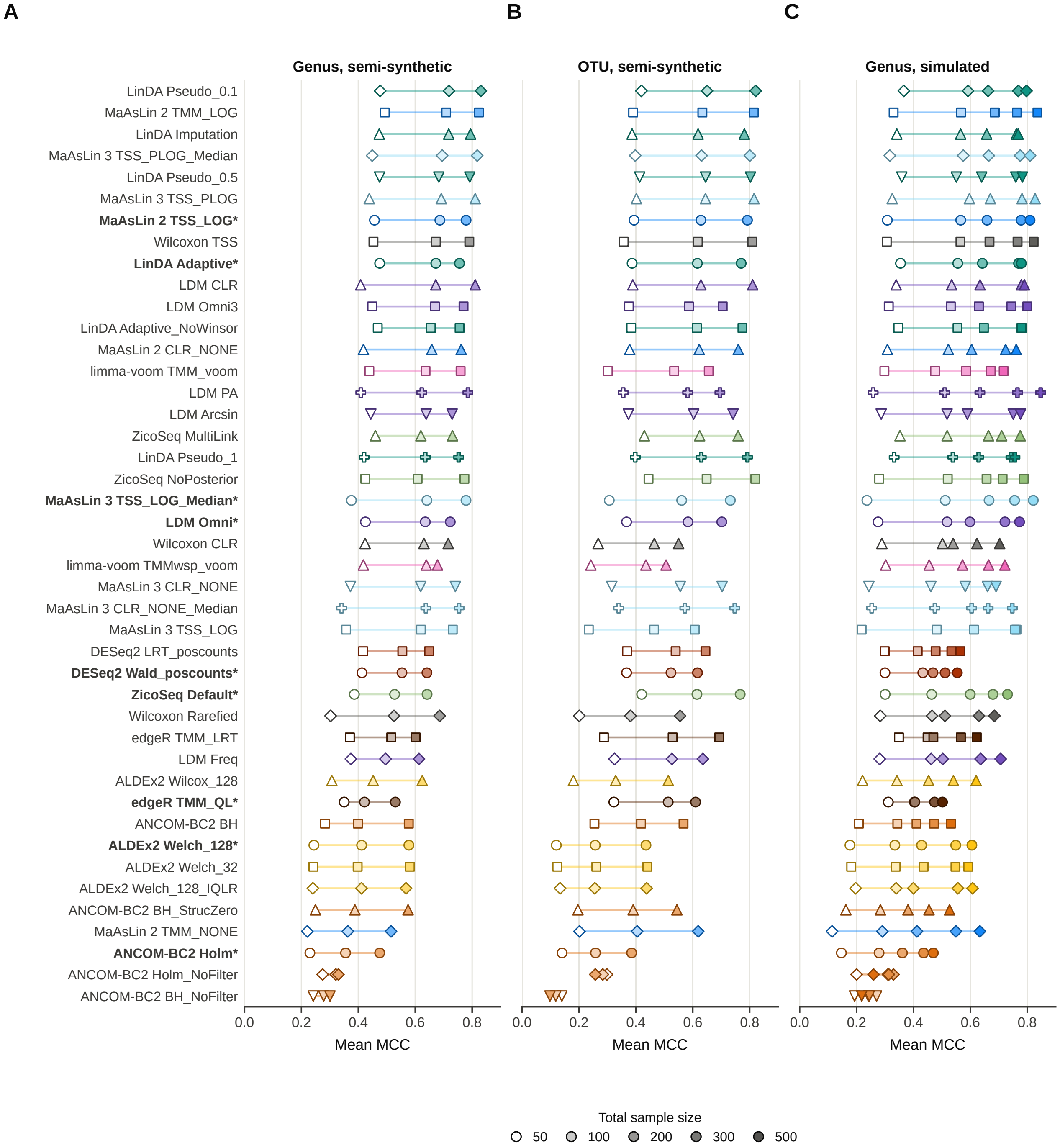


**Figure S4. MCC by sample size.** Mean MCC of every configuration at 50, 100 and 200 samples at genus (A) and OTU level (B) in semi-synthetic data, and at 50 to 500 samples at genus level in simulated data (C). Values are means over the other factors of the main detection design and the four replicates.

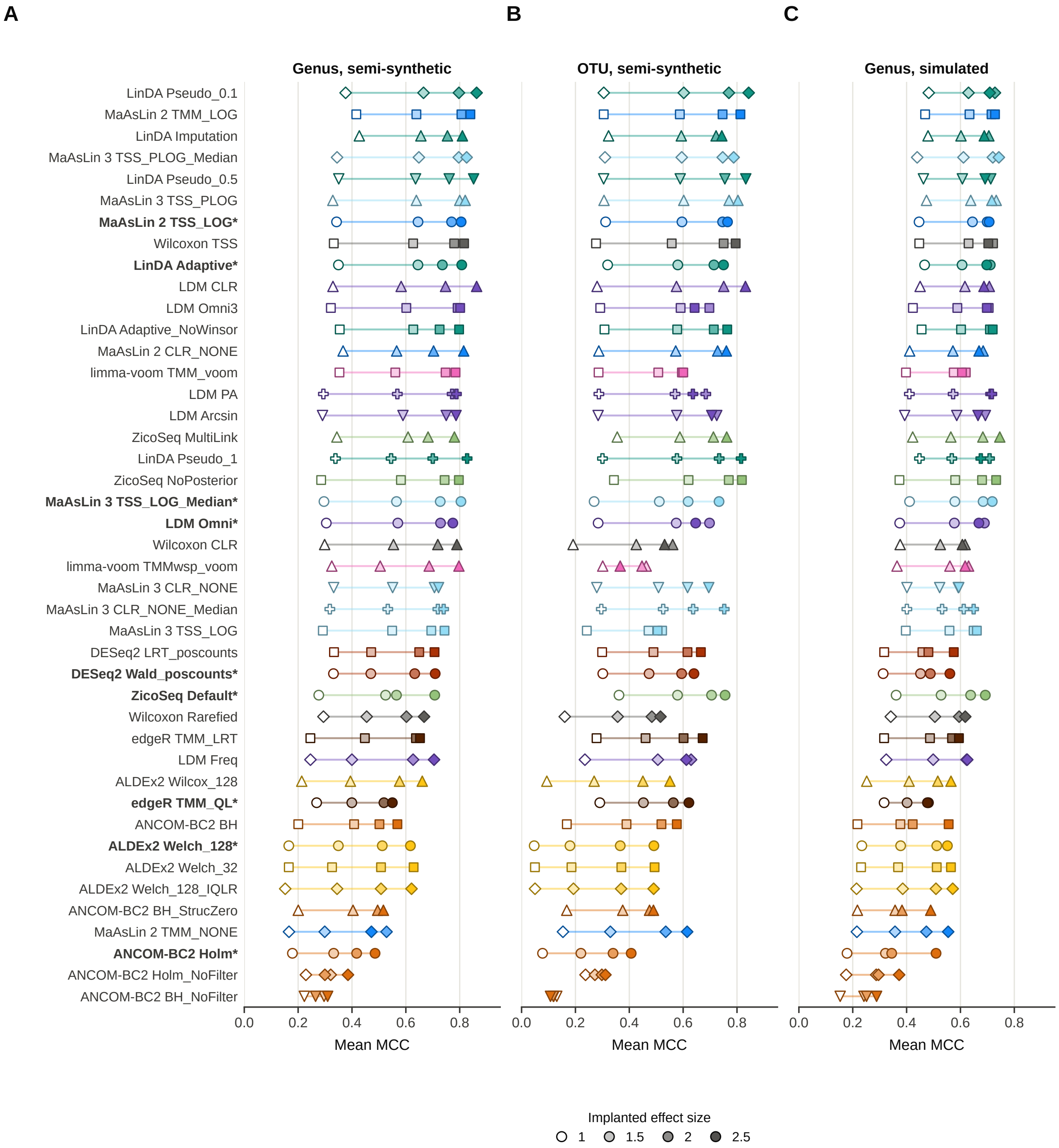


**Figure S5. MCC by effect size.** Mean MCC of every configuration at implanted effect sizes of 1.0, 1.5, 2.0 and 2.5 at genus (A) and OTU level (B) in semi-synthetic data and at genus level in simulated data (C). Values are means over the other factors of the main detection design and the four replicates.

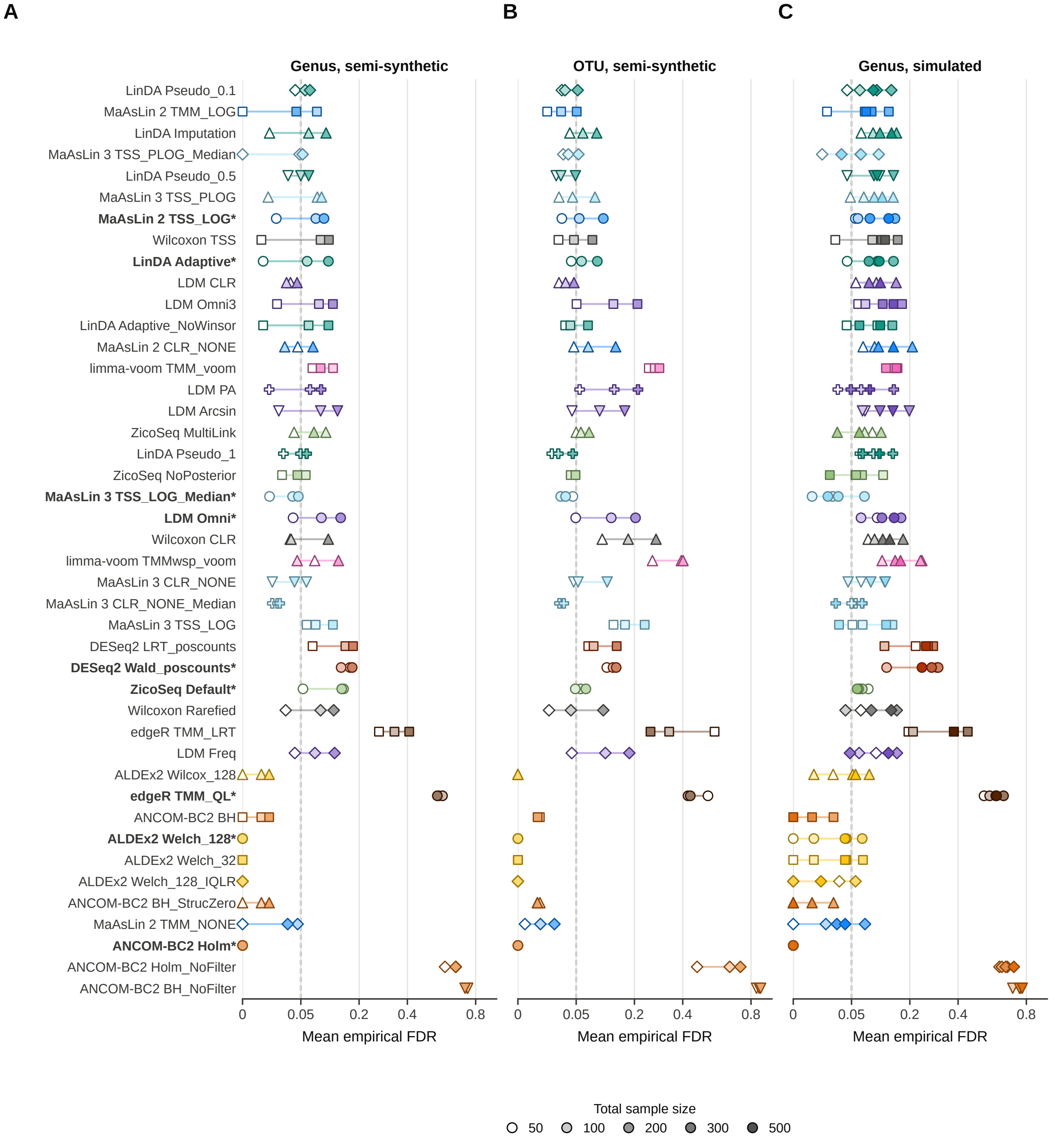


**Figure S6. Empirical FDR by sample size.** Mean empirical FDR of every configuration at 50, 100 and 200 samples at genus (A) and OTU level (B) in semi-synthetic data, and at 50 to 500 samples at genus level in simulated data (C). Values are means over the other factors of the main detection design and the four replicates.

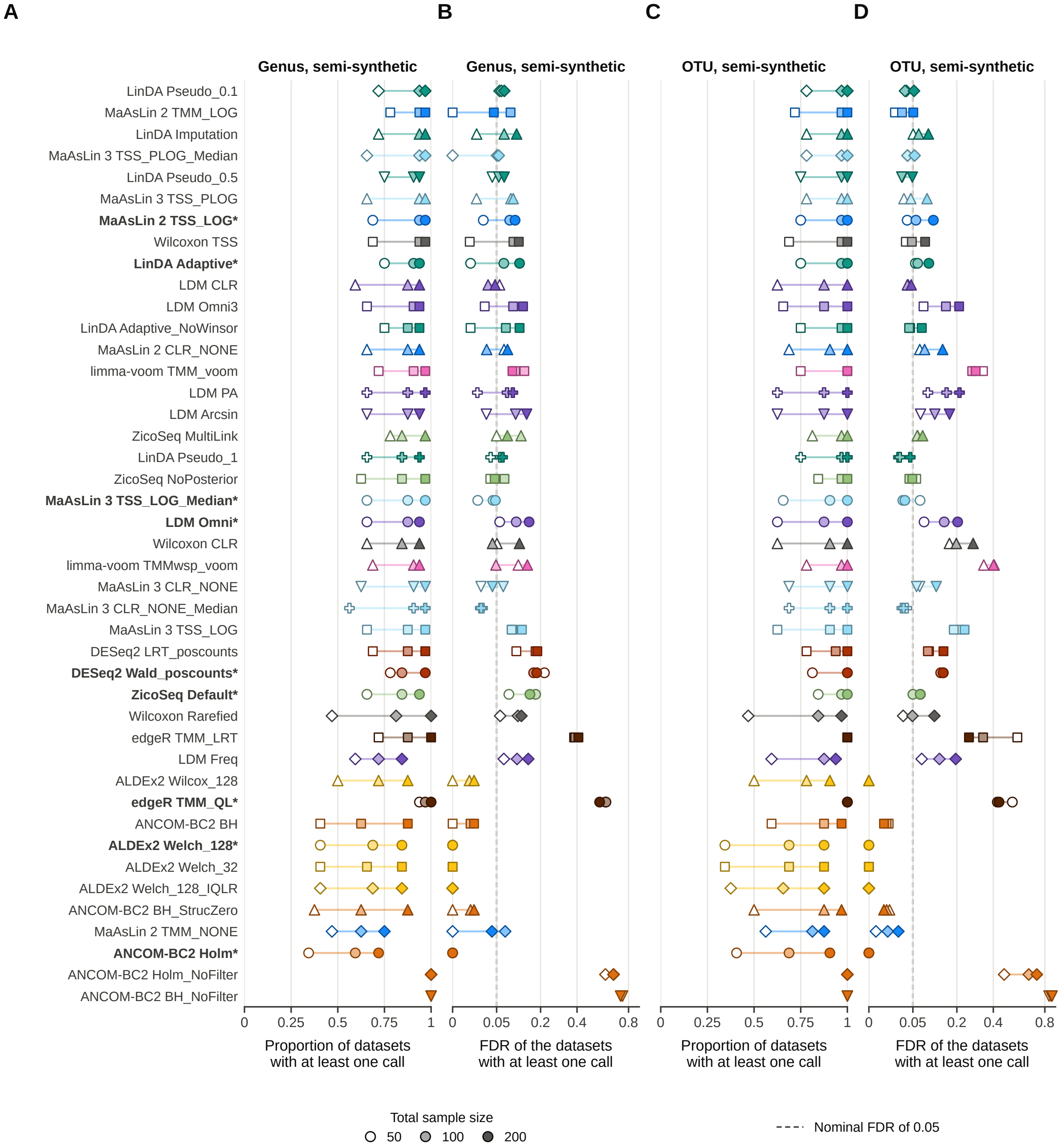


**Figure S7. Decomposition of the empirical FDR.** Proportion of datasets with at least one call (A, C) and FDR of the datasets with at least one call (B, D), for every configuration at 50, 100 and 200 samples (open to filled markers) at genus (A, B) and OTU level (C, D) in semi-synthetic data.

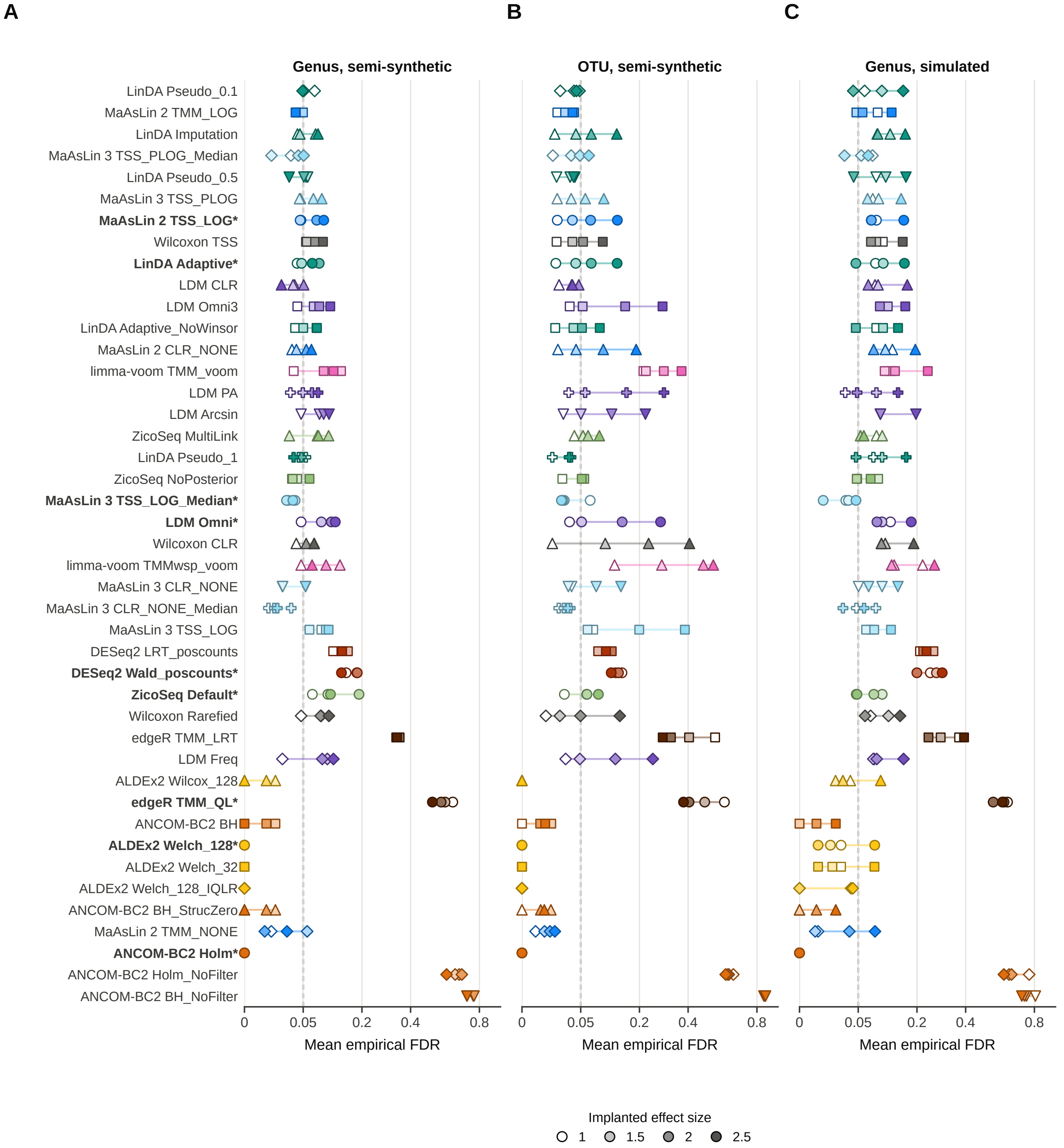


**Figure S8. Empirical FDR by effect size.** Mean empirical FDR of every configuration at implanted effect sizes of 1.0, 1.5, 2.0 and 2.5 at genus (A) and OTU level (B) in semi-synthetic data and at genus level in simulated data (C). Values are means over the other factors of the main detection design and the four replicates.

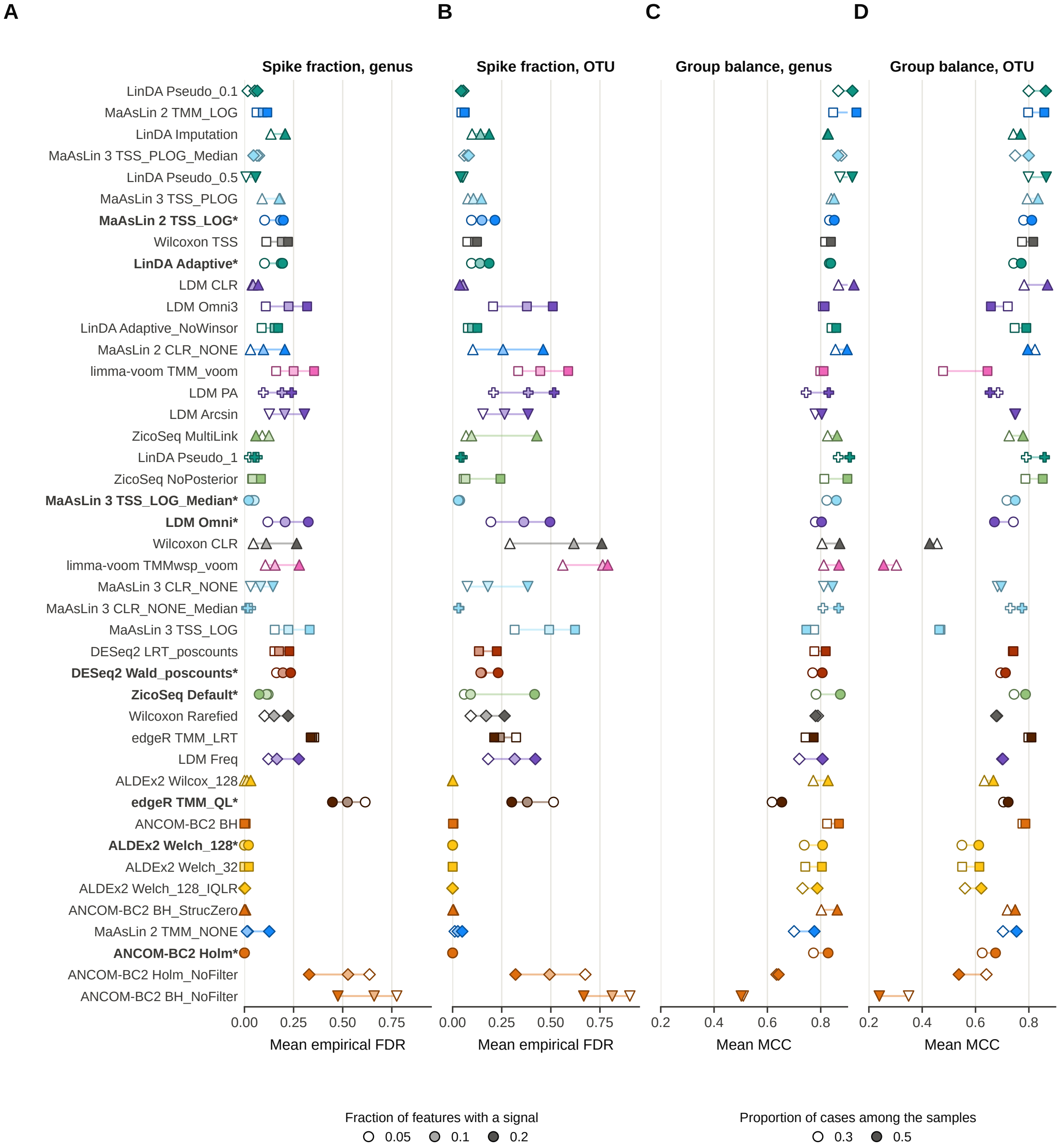


**Figure S9. Fraction of features with a signal and group balance.** Mean empirical FDR with signals implanted into 5%, 10% or 20% of the features at genus (A) and OTU level (B), and mean MCC with 30% or 50% of the samples in the case group at genus (C) and OTU level (D), in semi-synthetic data.

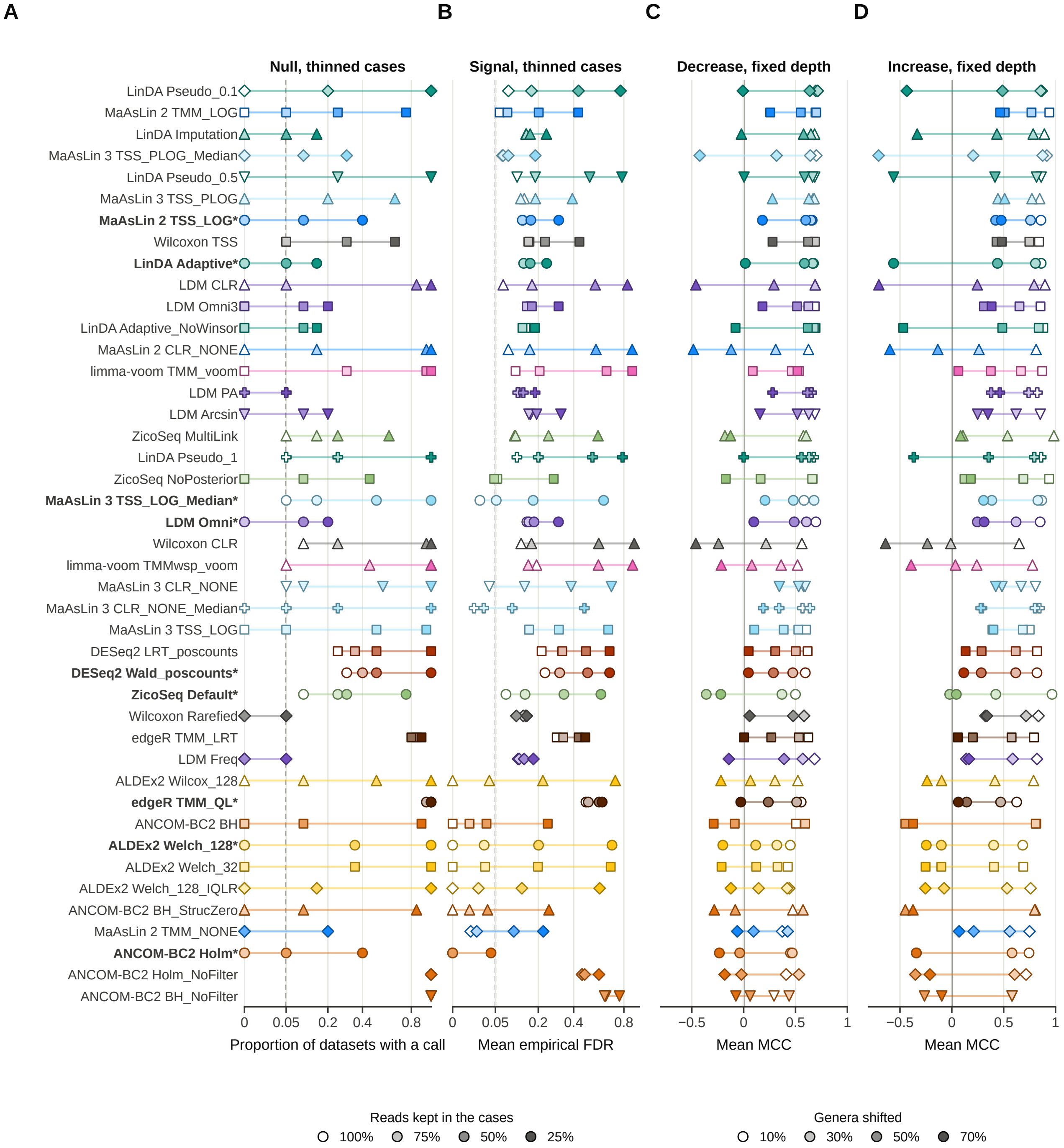


**Figure S10. Read depth that differs between groups and shifts of most taxa.** (A) Proportion of null datasets with at least one call after the reads of the case samples were thinned to 100%, 75%, 50% or 25%, at 100 and 200 samples on stool genera (20 datasets per level). The mean depth of the controls was then 1.0, 1.35, 2.0 and 4.0 times that of the cases. (B) Mean empirical FDR with joint signals implanted into 10% of the genera, thinned in the same way (16 datasets per level). (C, D) Mean MCC at fixed read depth when 10%, 30%, 50% or 70% of the genera were decreased (C) or increased (D) together, with an effect size of 1.5.

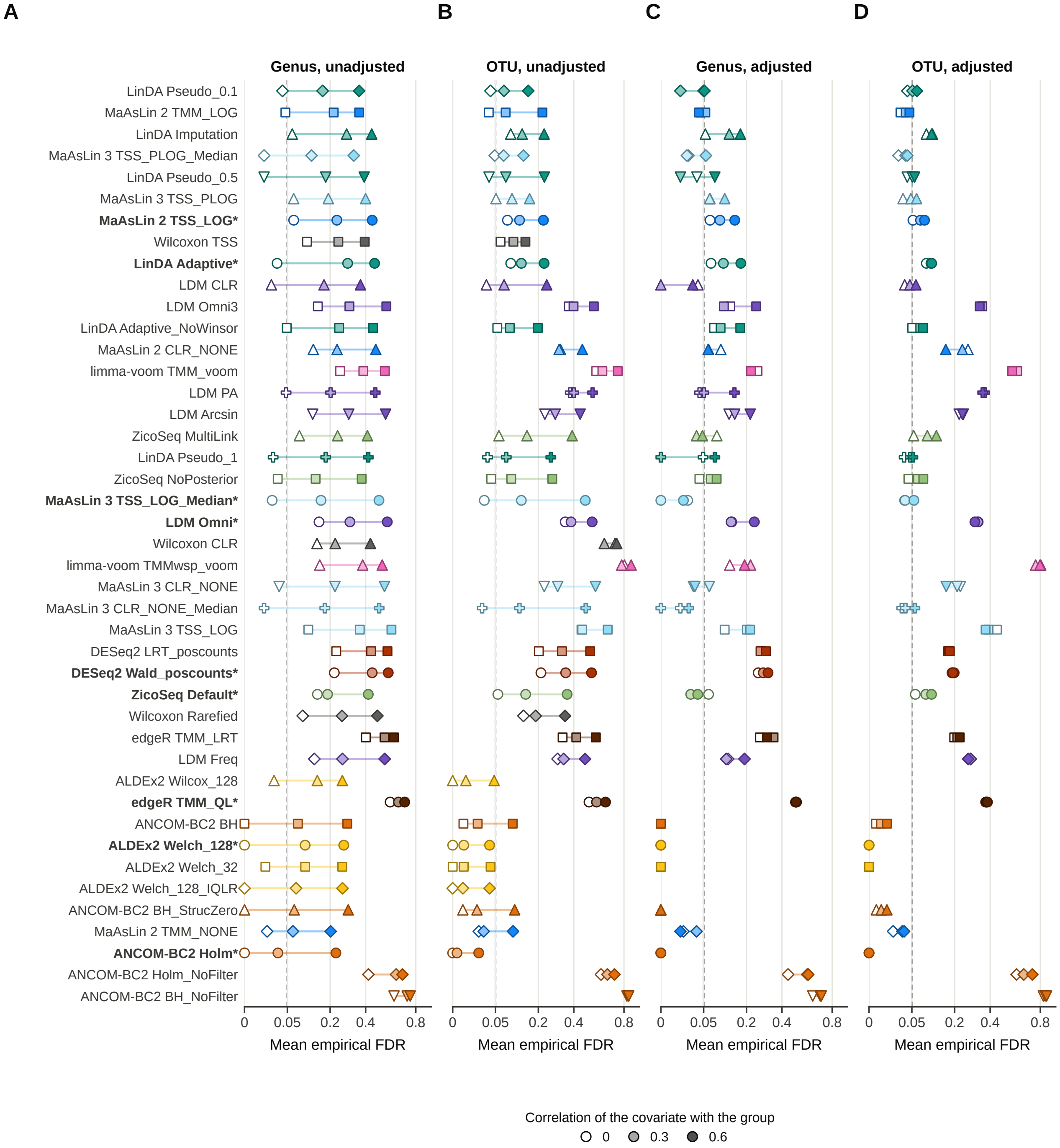


**Figure S11. Confounding.** Mean empirical FDR of every configuration with a confounder, a covariate correlated 0, 0.3 or 0.6 with the group, analysed without (A, B) and with (C, D) the covariate in the model, at genus (A, C) and OTU level (B, D).

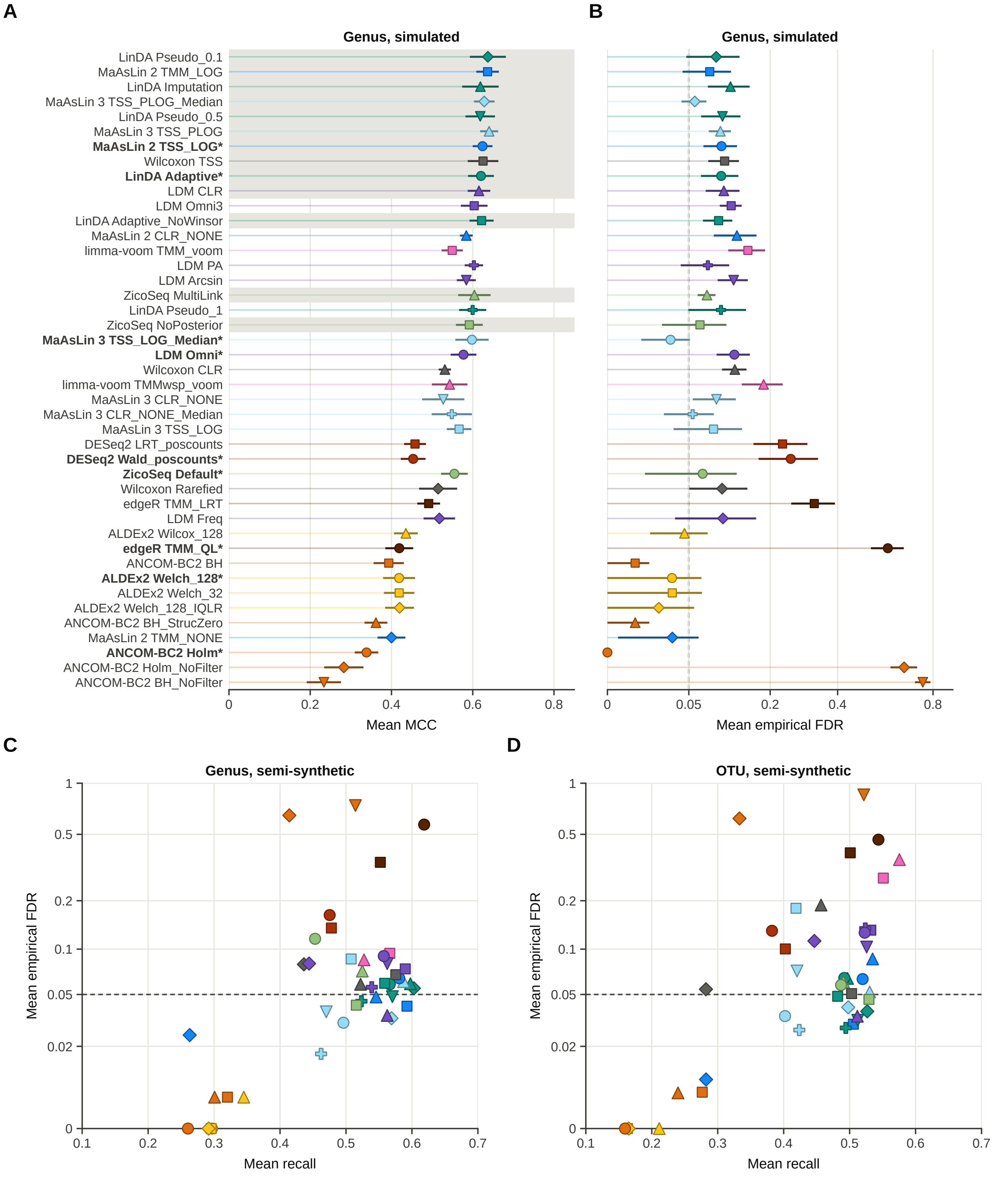


**Figure S12. Simulated data and recall against empirical FDR.** Mean MCC (A) and mean empirical FDR on a square-root scale (B) of every configuration in the main design at genus level in simulated data, with 95% confidence intervals across four replicates. Grey bands in A mark the configurations that could not be separated from the best one by a paired t test with Holm adjustment, and the dashed line in B the nominal FDR level of 0.05. Values are means over the 40 scenarios of the simulated main design. (C, D) Mean recall against mean empirical FDR at genus (C) and OTU level (D) in semi-synthetic data, over the 24 scenarios of the main design. The FDR axis is logarithmic after adding 0.01, which keeps configurations without false discoveries visible, and the dashed line marks the nominal FDR level of 0.05.

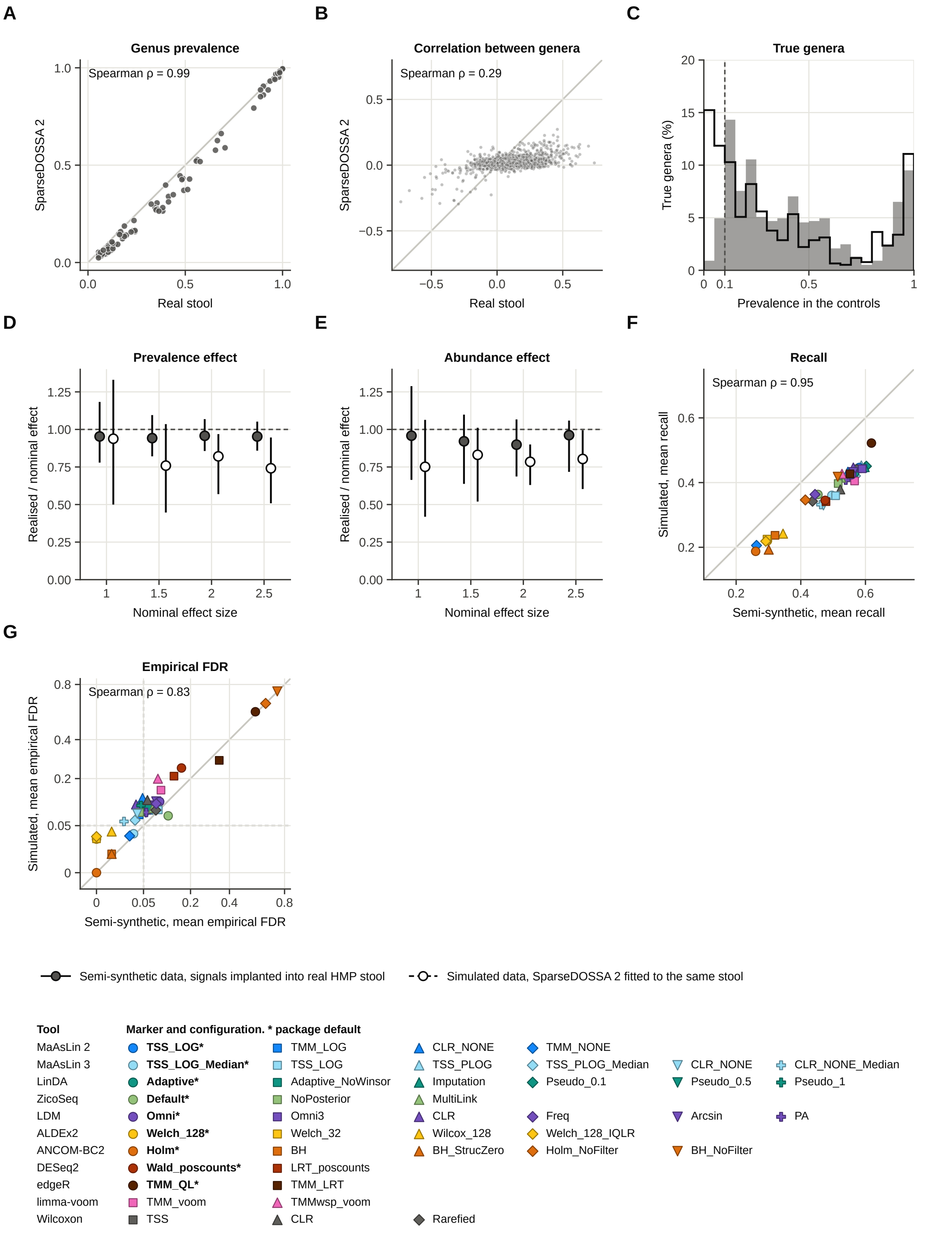


**Figure S13. Real stool and SparseDOSSA 2 tables and realised signals.** (A) Prevalence of each genus and (B) Spearman correlation of each pair of genera present in at least 10% of the samples, in the real stool genus table and in 20 tables simulated without signal from the fitted SparseDOSSA 2 model for the same genera and number of samples (mean over the 20 tables). (C) Prevalence in the control group of the true genera of the main design, as a percentage of all true genera, grey for semi-synthetic and outlined for simulated data. The dashed line marks the minimum prevalence of 10% that implantation requires in all samples of a dataset, which is why a few genera are below it in the controls. (D, E) Effect measured in the generated counts divided by the nominal effect for the true genera, in the log-odds of presence under prevalence-only signals (D) and in the mean log abundance of carriers under joint signals (E). A value of one means that the full effect reached the data. Points are medians and bars interquartile ranges over all true genera at 50 to 200 samples. (F, G) Mean recall (F) and mean empirical FDR (G) of every configuration on the 24 genus-level scenarios common to both approaches.

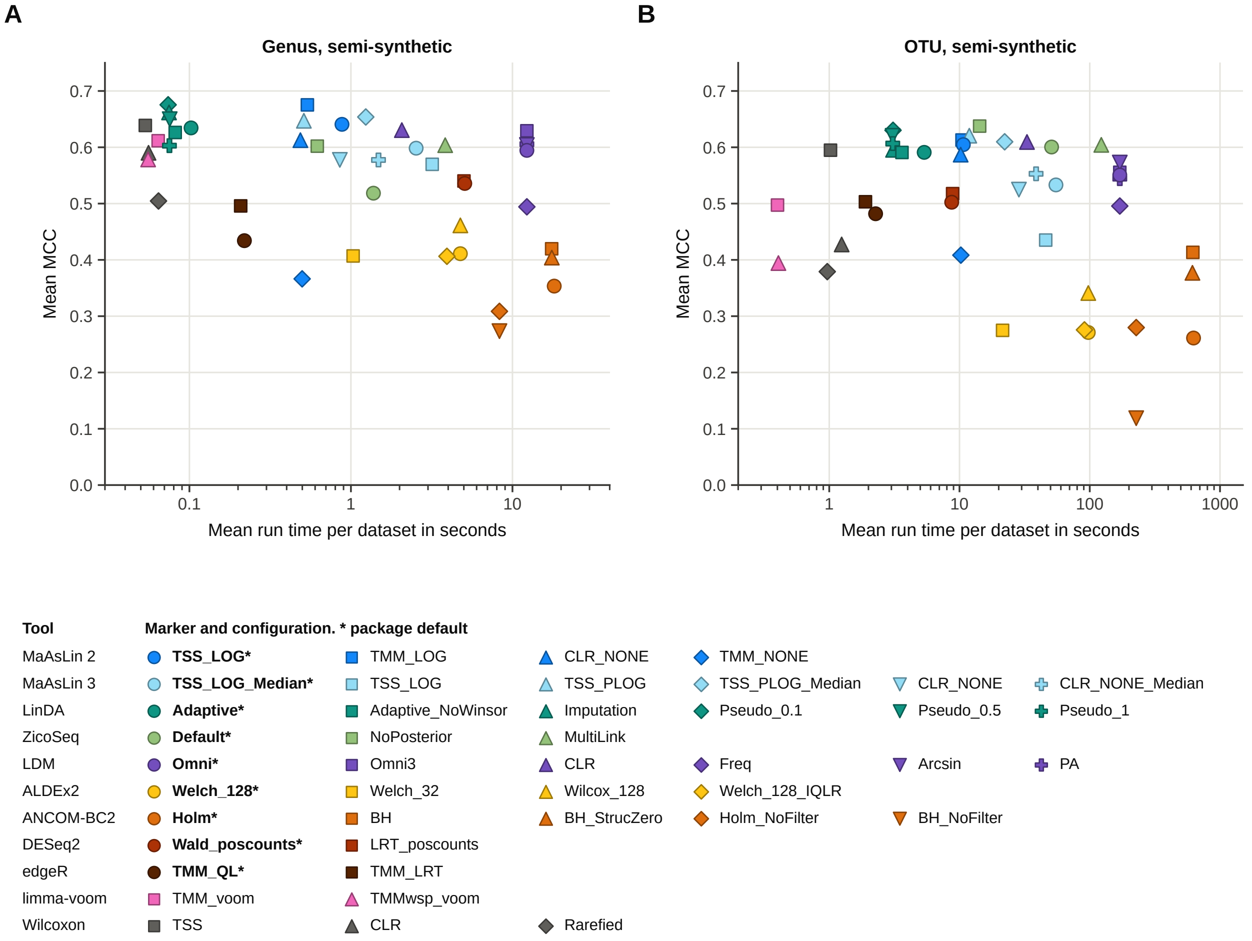


**Figure S14. Run time and MCC.** Mean run time per dataset against mean MCC at genus (A) and OTU level (B) in semi-synthetic data.

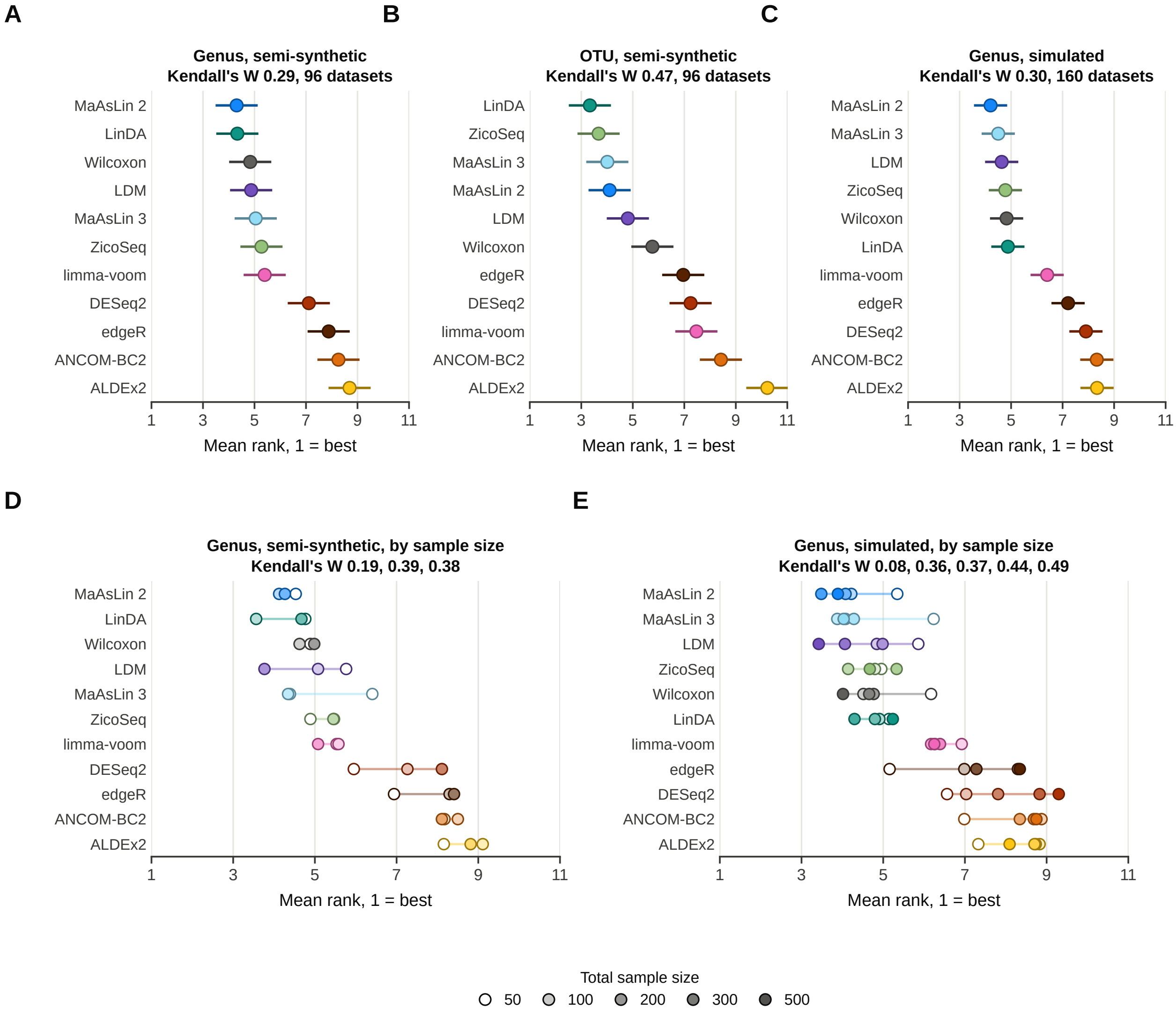


**Figure S15. Tool rankings.** Mean rank of the 11 tools over the datasets of the main detection design at genus (A) and OTU level (B) in semi-synthetic data and at genus level in simulated data (C), and at each total sample size at genus level in semi-synthetic (D) and simulated data (E). Each tool is represented by the configuration chosen on the other three replicates. Bars span the Nemenyi critical difference. Kendall's W and the number of datasets are given above each panel.

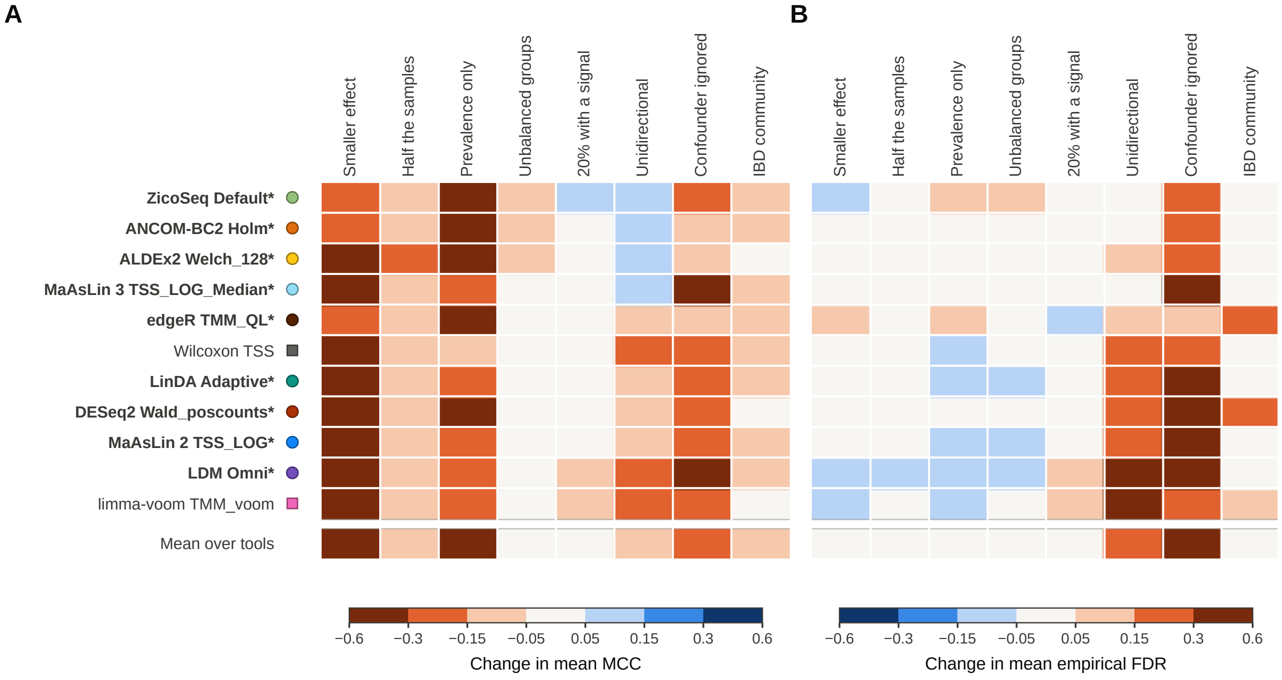


**Figure S16. Cost of each design factor.** Change in mean MCC (A) and mean empirical FDR (B) in each condition against its matched reference, at genus level in semi-synthetic data. Each row is the package default of a tool, or limma-voom with TMM normalisation and the Wilcoxon test on proportions for the two tools without one. References are an effect size of 2.0, 200 samples, joint signals, balanced groups, 10% of features with a signal, bidirectional signals, no confounding and stool.

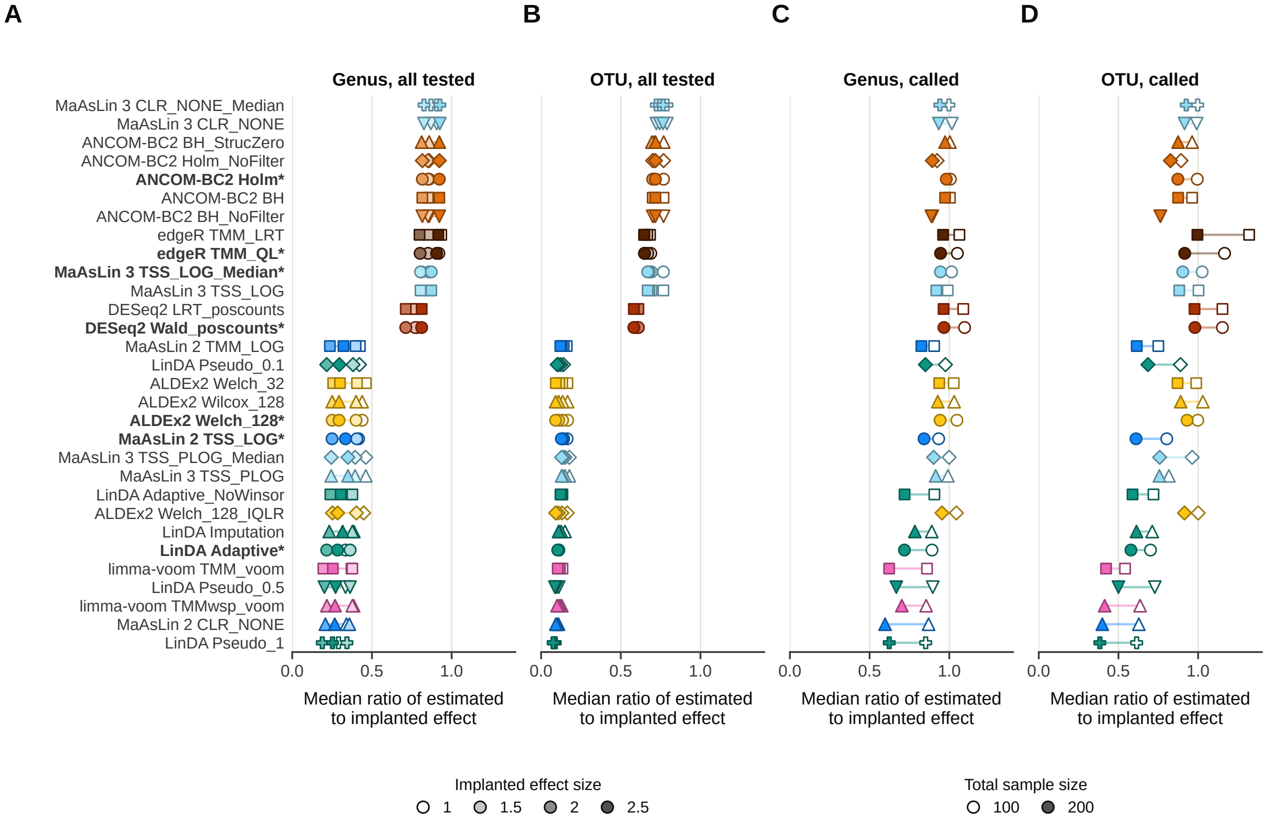


**Figure S17. Effect-size estimation by effect size and sample size.** Median ratio of the estimated to the implanted log2 fold change under abundance-only signals, for all tested implanted features at each effect size at genus (A) and OTU level (B), and for the implanted features that were called at 100 and 200 samples at genus (C) and OTU level (D).

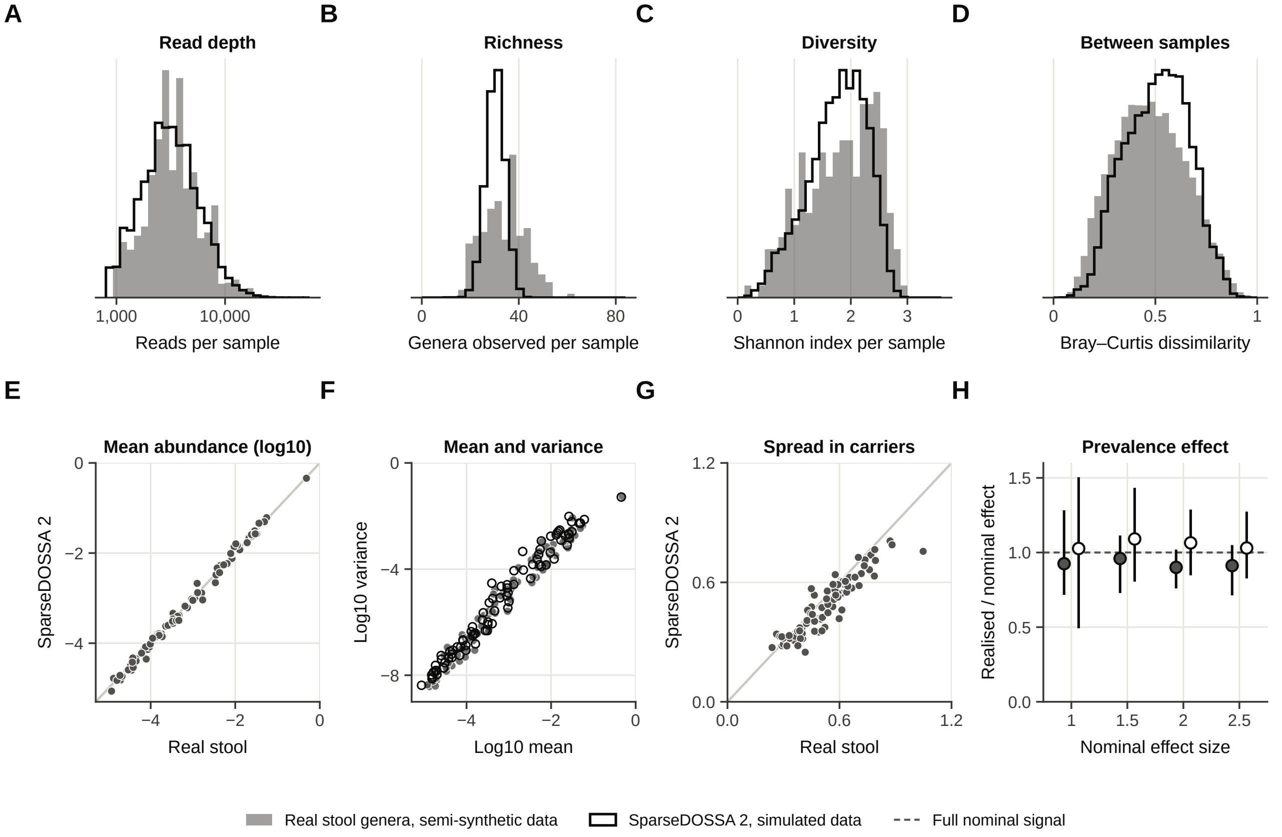


**Figure S18. Further properties of real and simulated stool tables.** The real stool genus table against 20 tables simulated without signal from the fitted SparseDOSSA 2 model for the same genera and number of samples. (A) Reads per sample, on a logarithmic axis. (B) Genera observed per sample. (C) Shannon index per sample. (D) Bray–Curtis dissimilarity between pairs of samples, for the real table and one simulated table. Histograms are scaled to equal area. (E) Mean relative abundance of each genus, real against simulated, on log10 scales. (F) Mean against variance of the relative abundance of each genus, on log10 scales, with filled markers for real and open markers for simulated genera. (G) Standard deviation of the log10 relative abundance among the samples that carry each genus, real against simulated. The grey line in E and G marks equal values. (H) Effect measured in the generated counts divided by the nominal effect, in the log-odds of presence of the true genera under joint signals, at 50 to 200 samples.

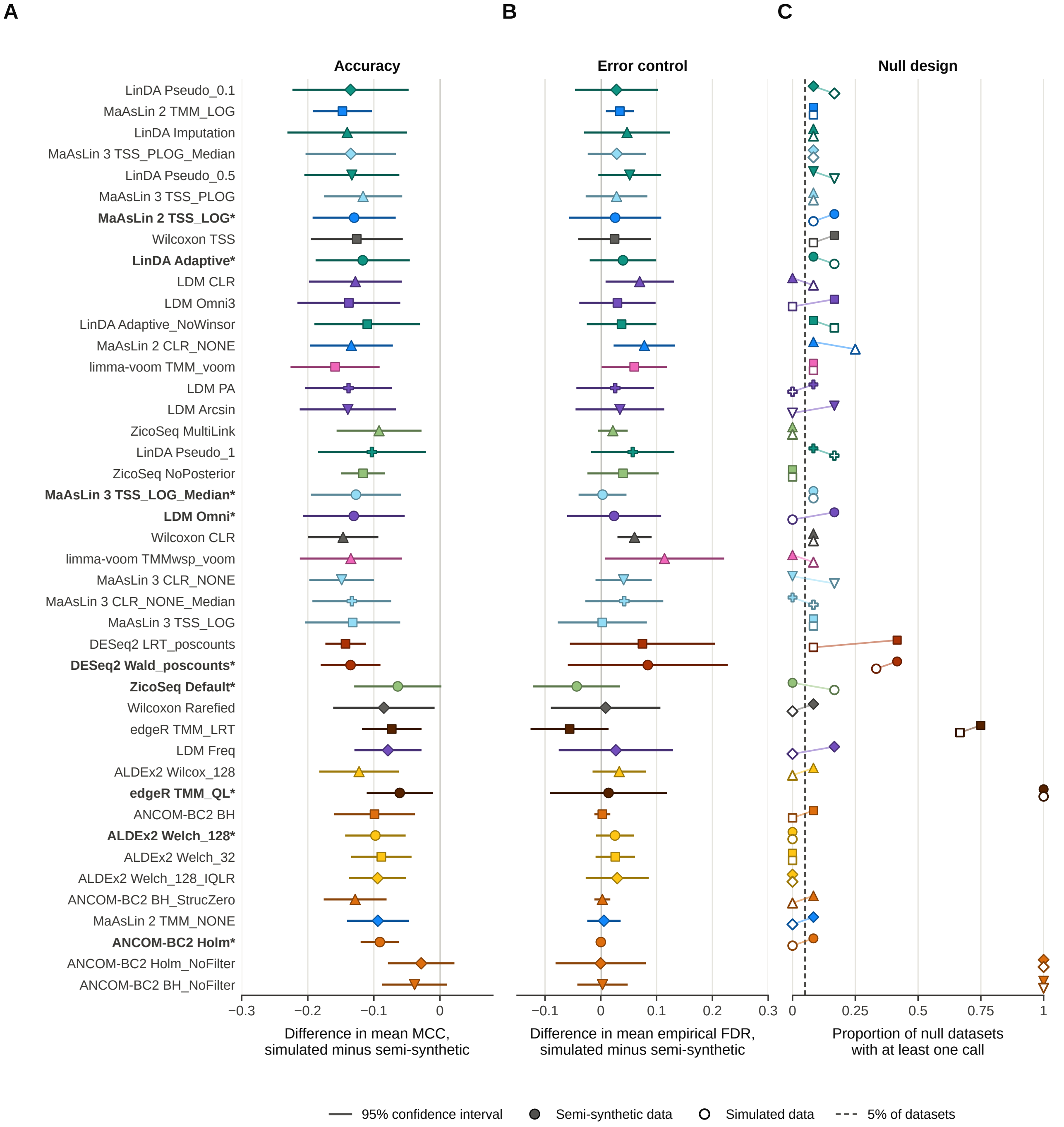


**Figure S19. Differences between simulated and semi-synthetic data for every configuration.** (A, B) Difference in mean MCC (A) and in mean empirical FDR (B) between simulated and semi-synthetic data on the 24 scenarios common to both approaches, with Welch 95% confidence intervals across replicates. Negative values in A mean a lower MCC in simulated data. (C) Proportion of the 12 null datasets of each approach at 50, 100 and 200 samples with at least one call, with filled markers for the semi-synthetic and open markers for simulated data.
