## Supplementary Tables for "A configuration-resolved benchmark of differential abundance analysis methods for human gut 16S rRNA microbiome data"

**Table S1. Designs and scenario grids.** Each design compares two groups, and each scenario was generated four times with independent seeds. Sample sizes are totals, split equally between the groups unless stated otherwise. Every configuration was run on every dataset, except in the effect-size design, which includes only configurations with a coefficient on an interpretable scale, and in the adjusted confounding analysis. Effect sizes are on the natural-log scale, a fold change of exp(e) in abundance and a shift of e in the log-odds of presence.

| Design | Data | Community | Resolution | Signal | Total samples | Effect sizes | Other factor | Scenarios | Datasets |
|---|---|---|---|---|---|---|---|---|---|
| Main detection | Semi-synthetic | Stool | Genus, OTU | Joint, prevalence only | 50, 100, 200 | 1.0, 1.5, 2.0, 2.5 | 10% of features, balanced | 48 | 192 |
| IBD community | Semi-synthetic | IBD | Genus | Joint, prevalence only | 50 | 1.5, 2.5 | 10% of features, balanced | 4 | 16 |
| Null | Semi-synthetic | Stool (both levels), IBD (genus) | Genus, OTU | None | 50, 100, 200 (stool), 50 (IBD) | None | No signal | 7 | 28 |
| Group balance | Semi-synthetic | Stool | Genus, OTU | Joint | 100, 200 | 1.5, 2.0, 2.5 | Cases 50% or 30% of samples | 24 | 96 |
| Fraction with a signal | Semi-synthetic | Stool | Genus, OTU | Joint | 100, 200 | 1.5, 2.0, 2.5 | 5%, 10% or 20% of features | 36 | 144 |
| Single extreme change | Semi-synthetic | Stool | Genus, OTU | Joint, one feature | 100, 200 | 10 | Single feature increased | 4 | 16 |
| Signal direction | Semi-synthetic | Stool | Genus, OTU | Joint | 100, 200 | 1.5, 2.0, 2.5 | Balanced or all increases | 24 | 96 |
| Effect-size estimation | Semi-synthetic | Stool | Genus, OTU | Abundance only, joint | 100, 200 | 1.0, 1.5, 2.0, 2.5 | Configurations with interpretable coefficients | 32 | 128 |
| Confounding | Semi-synthetic | Stool | Genus, OTU | Joint | 200 | 1.5, 2.0, 2.5 | Covariate correlation 0, 0.3, 0.6, analysed unadjusted and adjusted | 18 | 72 |
| Nominal level | Semi-synthetic | Stool | Genus, OTU | Joint, prevalence only | 100, 200 | 1.5, 2.0, 2.5 | Adjusted p-value thresholds 0.05, 0.10, 0.25 (refitted) | 24 | 96 |
| Main detection, simulated | Simulated | SparseDOSSA 2 fitted to stool | Genus | Joint, prevalence only | 50, 100, 200, 300, 500 | 1.0, 1.5, 2.0, 2.5 | 10% of features, balanced | 40 | 160 |
| Null, simulated | Simulated | SparseDOSSA 2 fitted to stool | Genus | None | 50, 100, 200, 300, 500 | None | No signal | 5 | 20 |
| Total | | | | | | | | 266 | 1064 |

**Table S2. Tools, configurations and run time.** Defaults are the package defaults, and for edgeR the quasi-likelihood test recommended in its user guide. In the other tables, package defaults are marked with an asterisk. Run time is the mean wall-clock time per dataset of one fit on one core of a 12-core workstation in the main detection design. BH, Benjamini–Hochberg.

| Tool | Configuration | Model and settings | Zero handling | Adjustment | Default | Extreme | Refitted at each level | Scale | Run time, genus (s) | Run time, OTU (s) | Run time, simulated (s) |
|---|---|---|---|---|---|---|---|---|---|---|---|
| MaAsLin 2 | TSS_LOG | TSS normalisation, log2 transformation, linear model | zeros replaced by half the smallest non-zero value per feature | BH | yes | | | log2 | 0.9 | 10.7 | 0.5 |
| MaAsLin 2 | CLR_NONE | CLR normalisation, no transformation, linear model | pseudo-count of 1 | BH | | | | ln | 0.5 | 10.2 | 0.4 |
| MaAsLin 2 | TMM_LOG | TMM normalisation, log2 transformation, linear model | half the smallest non-zero value | BH | | | | log2 | 0.5 | 10.5 | 0.5 |
| MaAsLin 2 | TMM_NONE | TMM normalisation, no transformation, linear model | none | BH | | yes | | none | 0.5 | 10.2 | 0.4 |
| MaAsLin 3 | TSS_LOG | TSS, log2, abundance model on non-zero values and prevalence model, joint test | zeros modelled by the prevalence model | BH | | | | log2 | 3.2 | 45.9 | 2.0 |
| MaAsLin 3 | TSS_LOG_Median | as TSS_LOG, with median comparison of abundance coefficients | prevalence model | BH | yes | | | log2 | 2.5 | 55.0 | 2.0 |
| MaAsLin 3 | TSS_PLOG | TSS, pseudo-log transformation, abundance test | pseudo-log keeps zeros on the log scale | BH | | | | log2 | 0.5 | 11.9 | 0.4 |
| MaAsLin 3 | TSS_PLOG_Median | as TSS_PLOG, with median comparison | pseudo-log | BH | | | | log2 | 1.2 | 22.2 | 1.0 |
| MaAsLin 3 | CLR_NONE | CLR normalisation, joint test | prevalence model | BH | | | | ln | 0.9 | 28.6 | 0.7 |
| MaAsLin 3 | CLR_NONE_Median | as CLR_NONE, with median comparison | prevalence model | BH | | | | ln | 1.5 | 38.7 | 1.2 |
| edgeR | TMM_LRT | TMM normalisation, negative binomial GLM, likelihood-ratio test | count model | BH | | | | log2 | 0.2 | 1.9 | 0.2 |
| edgeR | TMM_QL | TMM normalisation, quasi-likelihood F test | count model | BH | yes | | | log2 | 0.2 | 2.3 | 0.3 |
| ALDEx2 | Welch_128 | 128 Monte Carlo instances, denominator all features, Welch t test | Dirichlet prior 0.5 | expected BH | yes | | | log2 | 4.8 | 97.5 | 6.6 |
| ALDEx2 | Wilcox_128 | same fit as Welch_128, Wilcoxon test | Dirichlet prior 0.5 | expected BH | | | | log2 | 4.8 | 97.5 | 6.6 |
| ALDEx2 | Welch_32 | 32 Monte Carlo instances, Welch t test | Dirichlet prior 0.5 | expected BH | | | | log2 | 1.0 | 21.4 | 1.6 |
| ALDEx2 | Welch_128_IQLR | 128 instances, interquartile log-ratio denominator, Welch t test | Dirichlet prior 0.5 | expected BH | | | | log2 | 3.9 | 91.0 | 6.3 |
| ANCOM-BC2 | Holm | bias-corrected log-linear model, sensitivity analysis for pseudo-counts, 10% prevalence filter | zeros left out, sensitivity analysis over pseudo-counts of 0.01 to 0.5 | Holm | yes | | yes | ln | 18.1 | 626.2 | 13.5 |
| ANCOM-BC2 | BH | as Holm, with Benjamini-Hochberg adjustment | as Holm | BH | | | yes | ln | 17.5 | 618.8 | 13.3 |
| ANCOM-BC2 | BH_StrucZero | as BH, with structural zeros detected by their asymptotic lower bound | as Holm | BH | | | yes | ln | 17.5 | 614.2 | 13.3 |
| ANCOM-BC2 | BH_NoFilter | sensitivity analysis switched off | zeros left out | BH | | yes | | ln | 8.3 | 227.3 | 5.6 |

| Tool | Configuration | Model and settings | Zero handling | Adjustment | Default | Extreme | Refitted at each level | Scale | Run time, genus (s) | Run time, OTU (s) | Run time, simulated (s) |
|---|---|---|---|---|---|---|---|---|---|---|---|
| ANCOM-BC2 | Holm_NoFilter | sensitivity analysis switched off | zeros left out | Holm | | yes | | ln | 8.3 | 227.3 | 5.6 |
| LinDA | Adaptive | linear model on CLR values, bias correction by the mode of coefficients, winsorisation | adaptive, imputation when library size is associated with the model terms (P <= 0.1), otherwise pseudo-count 0.5, switch applied as documented | BH | yes | | | log2 | 0.1 | 5.4 | 0.5 |
| LinDA | Pseudo_0.1 | as Adaptive, fixed zero handling | pseudo-count 0.1 | BH | | | | log2 | 0.1 | 3.1 | 0.1 |
| LinDA | Pseudo_0.5 | as Adaptive, fixed zero handling | pseudo-count 0.5 | BH | | | | log2 | 0.1 | 3.1 | 0.1 |
| LinDA | Pseudo_1 | as Adaptive, fixed zero handling | pseudo-count 1 | BH | | | | log2 | 0.1 | 3.1 | 0.1 |
| LinDA | Imputation | as Adaptive, fixed zero handling | imputation of zeros | BH | | | | log2 | 0.1 | 3.1 | 0.1 |
| LinDA | Adaptive_NoWinsor | as Adaptive, without winsorisation | adaptive, switch applied as documented | BH | | | | log2 | 0.1 | 3.6 | 0.1 |
| ZicoSeq | Default | permutation linear model, reference-based correction, square-root link, posterior sampling, 99 permutations | posterior sampling of proportions | permutation FDR | yes | | | none | 1.4 | 50.9 | 1.7 |
| ZicoSeq | MultiLink | three links (powers 0.25, 0.5, 0.75) combined by their maximum | posterior sampling | permutation FDR | | | | none | 3.8 | 122.6 | 4.6 |
| ZicoSeq | NoPosterior | square-root link, no posterior sampling | none | permutation FDR | | | | none | 0.6 | 14.3 | 0.6 |
| LDM | Omni | omnibus of the frequency and arcsine-root tests, at most 5,000 permutations | none | permutation FDR | yes | | yes | none | 12.3 | 170.0 | 27.2 |
| LDM | Freq | test on relative abundances | none | permutation FDR | | | yes | none | 12.3 | 170.0 | 27.2 |
| LDM | Arcsin | test on arcsine-root relative abundances | none | permutation FDR | | | yes | none | 12.3 | 170.0 | 27.2 |
| LDM | PA | presence-absence test over rarefactions | presence-absence | permutation FDR | | | yes | none | 12.3 | 170.0 | 27.2 |
| LDM | Omni3 | omnibus of frequency, arcsine-root and presence-absence tests | none | permutation FDR | | | yes | none | 12.3 | 170.0 | 27.2 |
| LDM | CLR | LDM on CLR values (compositional analysis) | pseudo-count 0.5 | permutation FDR | | | yes | none | 2.1 | 33.0 | 3.0 |
| DESeq2 | Wald_poscounts | negative binomial GLM, Wald test, poscounts size factors, default filters | count model | BH with independent filtering | yes | | yes | log2 | 5.1 | 8.7 | 3.8 |
| DESeq2 | LRT_poscounts | negative binomial GLM, likelihood-ratio test, poscounts size factors | count model | BH with independent filtering | | | yes | log2 | 5.0 | 8.9 | 3.9 |
| limma-voom | TMM_voom | TMM normalisation, voom weights, moderated t test | log-CPM with offset | BH | | | | log2 | 0.1 | 0.4 | 0.1 |
| limma-voom | TMMwsp_voom | TMM with singleton pairing, voom weights, moderated t test | log-CPM with offset | BH | | | | log2 | 0.1 | 0.4 | 0.1 |
| Wilcoxon | TSS | rank-sum test on relative abundances | none | BH | | | | none | 0.1 | 1.0 | 0.1 |
| Wilcoxon | CLR | rank-sum test on CLR values | pseudo-count of 1 | BH | | | | none | 0.1 | 1.2 | 0.1 |
| Wilcoxon | Rarefied | rank-sum test on counts rarefied to the smallest library | rarefaction | BH | | | | none | 0.1 | 1.0 | 0.1 |

**Table S3. Overall performance in the main detection design.** Means over scenarios with 95% confidence intervals across replicates. No call is the proportion of datasets without a significant feature, scored as an MCC of 0. FDR against 0.05 gives below or above when the 95% interval of the empirical FDR lies entirely below or above the nominal level, and is empty when the interval includes it. Configurations were ranked within each scenario by MCC, rank 1 being the best. Holm P compares each configuration with the best in a paired t test across scenarios.

| Tool | Configuration | MCC | Recall | FDR | F1 | No call (%) | Calls per dataset | FDR against 0.05 | Mean rank | Holm P vs best | Within critical difference of best |
|---|---|---|---|---|---|---|---|---|---|---|---|
| ***Semi-synthetic stool, genus level (24 scenarios, 96 datasets). Friedman chi-squared 457.1, P 3.7e-71, Kendall's W 0.45, Nemenyi critical difference 14.2*** | | | | | | | | | | | |
| LinDA | Pseudo_0.1 | 0.675 [0.637, 0.714] | 0.603 [0.561, 0.644] | 0.055 [0.034, 0.076] | 0.668 [0.630, 0.706] | 12.5 | 5.2 | | 9.4 | 0.99 | yes |
| MaAsLin 2 | TMM_LOG | 0.675 [0.628, 0.723] | 0.592 [0.564, 0.621] | 0.041 [0.029, 0.054] | 0.660 [0.615, 0.704] | 10.4 | 5.1 | | 7.5 | | yes |
| LinDA | Imputation | 0.662 [0.624, 0.699] | 0.598 [0.565, 0.630] | 0.059 [0.004, 0.114] | 0.652 [0.611, 0.692] | 12.5 | 5.8 | | 12.1 | 0.92 | yes |
| MaAsLin 3 | TSS_PLOG_Median | 0.654 [0.596, 0.712] | 0.569 [0.512, 0.626] | 0.034 [0.000, 0.080] | 0.642 [0.580, 0.704] | 14.6 | 5.0 | | 12.6 | 0.92 | yes |
| LinDA | Pseudo_0.5 | 0.650 [0.625, 0.675] | 0.570 [0.556, 0.585] | 0.048 [0.015, 0.082] | 0.638 [0.616, 0.661] | 13.5 | 4.9 | | 11.6 | 0.29 | yes |
| MaAsLin 3 | TSS_PLOG | 0.647 [0.601, 0.693] | 0.587 [0.559, 0.615] | 0.061 [0.009, 0.114] | 0.637 [0.593, 0.680] | 14.6 | 5.5 | | 14.0 | 0.82 | yes |
| MaAsLin 2 | TSS_LOG * | 0.641 [0.604, 0.677] | 0.581 [0.561, 0.600] | 0.064 [0.010, 0.119] | 0.627 [0.597, 0.658] | 13.5 | 5.6 | | 12.8 | 0.43 | yes |
| Wilcoxon | TSS | 0.639 [0.598, 0.680] | 0.576 [0.560, 0.591] | 0.068 [0.003, 0.133] | 0.627 [0.584, 0.670] | 13.5 | 5.5 | | 15.6 | 0.45 | yes |
| LinDA | Adaptive * | 0.634 [0.611, 0.657] | 0.566 [0.552, 0.581] | 0.059 [0.020, 0.098] | 0.621 [0.590, 0.651] | 13.5 | 5.5 | | 15.1 | 0.032 | yes |
| LDM | CLR | 0.631 [0.623, 0.638] | 0.562 [0.545, 0.580] | 0.035 [0.000, 0.075] | 0.621 [0.608, 0.635] | 19.8 | 4.8 | | 13.5 | 0.17 | yes |
| LDM | Omni3 | 0.629 [0.579, 0.679] | 0.590 [0.555, 0.625] | 0.074 [0.029, 0.120] | 0.621 [0.569, 0.673] | 16.7 | 6.4 | | 13.9 | 0.29 | yes |
| LinDA | Adaptive_NoWinsor | 0.626 [0.591, 0.661] | 0.559 [0.540, 0.577] | 0.060 [0.015, 0.104] | 0.613 [0.577, 0.649] | 14.6 | 5.3 | | 15.9 | 0.0094 | yes |
| MaAsLin 2 | CLR_NONE | 0.613 [0.605, 0.620] | 0.546 [0.524, 0.567] | 0.048 [0.014, 0.082] | 0.598 [0.587, 0.609] | 17.7 | 5.0 | | 15.2 | 0.026 | yes |
| limma-voom | TMM_voom | 0.612 [0.569, 0.655] | 0.566 [0.539, 0.594] | 0.094 [0.049, 0.139] | 0.600 [0.567, 0.632] | 13.5 | 6.2 | | 16.7 | 0.0014 | yes |
| LDM | PA | 0.606 [0.552, 0.659] | 0.539 [0.491, 0.587] | 0.056 [0.004, 0.108] | 0.593 [0.542, 0.644] | 16.7 | 5.6 | | 19.2 | 0.12 | yes |
| LDM | Arcsin | 0.604 [0.554, 0.655] | 0.562 [0.545, 0.580] | 0.081 [0.045, 0.116] | 0.594 [0.540, 0.647] | 17.7 | 6.1 | | 17.3 | 0.038 | yes |
| ZicoSeq | MultiLink | 0.604 [0.571, 0.636] | 0.525 [0.493, 0.557] | 0.072 [0.050, 0.094] | 0.590 [0.566, 0.614] | 13.5 | 5.0 | above | 16.8 | 0.035 | yes |
| LinDA | Pseudo_1 | 0.603 [0.586, 0.620] | 0.523 [0.508, 0.539] | 0.045 [0.011, 0.079] | 0.591 [0.571, 0.611] | 18.8 | 4.5 | | 16.7 | 0.019 | yes |
| ZicoSeq | NoPosterior | 0.602 [0.567, 0.637] | 0.516 [0.486, 0.545] | 0.042 [0.032, 0.052] | 0.590 [0.555, 0.625] | 18.8 | 4.4 | | 17.7 | 0.024 | yes |
| MaAsLin 3 | TSS_LOG_Median * | 0.598 [0.580, 0.617] | 0.496 [0.469, 0.523] | 0.031 [0.009, 0.053] | 0.582 [0.557, 0.606] | 16.7 | 4.2 | | 20.2 | 0.0084 | yes |
| LDM | Omni * | 0.595 [0.529, 0.660] | 0.557 [0.528, 0.587] | 0.090 [0.063, 0.118] | 0.584 [0.518, 0.651] | 17.7 | 6.2 | above | 19.0 | 0.0046 | yes |
| Wilcoxon | CLR | 0.590 [0.545, 0.635] | 0.522 [0.501, 0.543] | 0.059 [0.037, 0.080] | 0.578 [0.534, 0.622] | 18.8 | 5.1 | | 20.2 | 0.00031 | yes |
| limma-voom | TMMwsp_voom | 0.578 [0.564, 0.593] | 0.527 [0.479, 0.575] | 0.085 [0.000, 0.189] | 0.563 [0.556, 0.570] | 15.6 | 6.3 | | 21.0 | 0.00034 | yes |
| MaAsLin 3 | CLR_NONE | 0.578 [0.548, 0.607] | 0.470 [0.439, 0.501] | 0.038 [0.017, 0.059] | 0.559 [0.522, 0.595] | 16.7 | 4.0 | | 21.1 | 0.0035 | yes |
| MaAsLin 3 | CLR_NONE_Median | 0.577 [0.541, 0.614] | 0.462 [0.435, 0.490] | 0.017 [0.000, 0.034] | 0.559 [0.520, 0.597] | 18.8 | 3.8 | below | 20.1 | 0.015 | yes |
| MaAsLin 3 | TSS_LOG | 0.570 [0.508, 0.632] | 0.508 [0.476, 0.540] | 0.086 [0.040, 0.132] | 0.559 [0.497, 0.621] | 16.7 | 5.8 | | 24.8 | 0.0017 | |
| DESeq2 | LRT_poscounts | 0.540 [0.509, 0.571] | 0.478 [0.474, 0.482] | 0.136 [0.071, 0.201] | 0.533 [0.505, 0.560] | 15.6 | 4.7 | above | 24.3 | 3.1e-05 | |
| DESeq2 | Wald_poscounts * | 0.536 [0.499, 0.573] | 0.475 [0.467, 0.483] | 0.163 [0.084, 0.242] | 0.530 [0.499, 0.562] | 13.5 | 4.7 | above | 25.6 | 3.1e-05 | |
| ZicoSeq | Default * | 0.518 [0.458, 0.579] | 0.453 [0.405, 0.501] | 0.116 [0.075, 0.158] | 0.512 [0.458, 0.566] | 18.8 | 5.0 | above | 23.8 | 0.002 | |
| Wilcoxon | Rarefied | 0.505 [0.483, 0.526] | 0.436 [0.404, 0.468] | 0.080 [0.056, 0.104] | 0.492 [0.468, 0.515] | 24.0 | 4.5 | above | 27.8 | 0.00014 | |
| edgeR | TMM_LRT | 0.496 [0.468, 0.524] | 0.552 [0.517, 0.587] | 0.341 [0.308, 0.374] | 0.516 [0.486, 0.546] | 13.5 | 7.2 | above | 30.5 | 1.1e-06 | |
| LDM | Freq | 0.494 [0.470, 0.518] | 0.444 [0.419, 0.469] | 0.081 [0.028, 0.134] | 0.483 [0.461, 0.506] | 28.1 | 4.9 | | 27.9 | 0.0001 | |
| ALDEx2 | Wilcox_128 | 0.461 [0.411, 0.511] | 0.345 [0.292, 0.398] | 0.005 [0.000, 0.016] | 0.436 [0.387, 0.486] | 30.2 | 2.8 | below | 29.6 | 5.2e-05 | |
| edgeR | TMM_QL * | 0.434 [0.394, 0.474] | 0.618 [0.574, 0.663] | 0.571 [0.527, 0.615] | 0.470 [0.436, 0.503] | 3.1 | 11.6 | above | 34.0 | 2.1e-08 | |
| ANCOM-BC2 | BH | 0.420 [0.355, 0.485] | 0.320 [0.272, 0.368] | 0.005 [0.000, 0.016] | 0.395 [0.336, 0.454] | 36.5 | 2.6 | below | 29.1 | 0.00035 | |
| ALDEx2 | Welch_128 * | 0.411 [0.372, 0.450] | 0.297 [0.256, 0.337] | 0.000 [0.000, 0.000] | 0.383 [0.338, 0.428] | 35.4 | 2.4 | below | 33.2 | 9.6e-06 | |
| ALDEx2 | Welch_32 | 0.407 [0.377, 0.437] | 0.296 [0.268, 0.323] | 0.000 [0.000, 0.000] | 0.380 [0.349, 0.411] | 36.5 | 2.4 | below | 33.3 | 1.4e-05 | |
| ALDEx2 | Welch_128_IQLR | 0.406 [0.368, 0.444] | 0.292 [0.241, 0.342] | 0.000 [0.000, 0.000] | 0.377 [0.332, 0.421] | 35.4 | 2.3 | below | 33.7 | 9.6e-06 | |
| ANCOM-BC2 | BH_StrucZero | 0.404 [0.355, 0.453] | 0.301 [0.266, 0.336] | 0.005 [0.000, 0.016] | 0.379 [0.334, 0.423] | 37.5 | 2.4 | below | 31.4 | 0.00014 | |
| MaAsLin 2 | TMM_NONE | 0.366 [0.360, 0.372] | 0.263 [0.247, 0.279] | 0.025 [0.000, 0.055] | 0.340 [0.326, 0.353] | 38.5 | 2.2 | | 35.3 | 1.4e-06 | |
| ANCOM-BC2 | Holm * | 0.353 [0.328, 0.379] | 0.260 [0.234, 0.287] | 0.000 [0.000, 0.000] | 0.329 [0.303, 0.355] | 44.8 | 2.1 | below | 33.2 | 0.00014 | |
| ANCOM-BC2 | Holm_NoFilter | 0.309 [0.261, 0.357] | 0.414 [0.377, 0.451] | 0.647 [0.573, 0.722] | 0.357 [0.316, 0.397] | 0.0 | 8.7 | above | 36.2 | 1.1e-05 | |

| Tool | Configuration | MCC | Recall | FDR | F1 | No call (%) | Calls per dataset | FDR against 0.05 | Mean rank | Holm P vs best | Within critical difference of best |
|---|---|---|---|---|---|---|---|---|---|---|---|
| ANCOM-BC2 | BH_NoFilter | 0.273 [0.221, 0.326] | 0.514 [0.463, 0.566] | 0.739 [0.694, 0.785] | 0.336 [0.288, 0.383] | 0.0 | 15.4 | above | 36.9 | 7.3e-07 | |
| ***Semi-synthetic stool, OTU level (24 scenarios, 96 datasets). Friedman chi-squared 567.0, P 3.8e-93, Kendall's W 0.56, Nemenyi critical difference 14.2*** | | | | | | | | | | | |
| LinDA | Pseudo_0.1 | 0.630 [0.617, 0.644] | 0.527 [0.517, 0.536] | 0.038 [0.037, 0.039] | 0.613 [0.605, 0.621] | 8.3 | 149.3 | below | 8.7 | 0.42 | yes |
| MaAsLin 2 | TMM_LOG | 0.613 [0.599, 0.627] | 0.506 [0.497, 0.514] | 0.030 [0.021, 0.039] | 0.593 [0.582, 0.604] | 10.4 | 142.6 | below | 10.6 | 0.082 | yes |
| LinDA | Imputation | 0.596 [0.584, 0.607] | 0.497 [0.489, 0.505] | 0.064 [0.050, 0.078] | 0.581 [0.571, 0.591] | 8.3 | 149.2 | above | 14.8 | 0.052 | yes |
| MaAsLin 3 | TSS_PLOG_Median | 0.610 [0.599, 0.621] | 0.498 [0.496, 0.499] | 0.041 [0.036, 0.045] | 0.593 [0.589, 0.598] | 8.3 | 142.1 | below | 12.2 | 0.11 | yes |
| LinDA | Pseudo_0.5 | 0.620 [0.601, 0.640] | 0.512 [0.503, 0.521] | 0.033 [0.025, 0.040] | 0.601 [0.588, 0.614] | 9.4 | 144.3 | below | 9.1 | 0.11 | yes |
| MaAsLin 3 | TSS_PLOG | 0.621 [0.608, 0.633] | 0.530 [0.522, 0.539] | 0.052 [0.039, 0.066] | 0.604 [0.595, 0.613] | 8.3 | 157.1 | | 9.5 | 0.33 | yes |
| MaAsLin 2 | TSS_LOG * | 0.605 [0.589, 0.620] | 0.520 [0.507, 0.532] | 0.064 [0.045, 0.082] | 0.588 [0.578, 0.599] | 9.4 | 159.8 | | 12.5 | 0.082 | yes |
| Wilcoxon | TSS | 0.595 [0.589, 0.601] | 0.503 [0.497, 0.509] | 0.051 [0.041, 0.061] | 0.577 [0.567, 0.586] | 11.5 | 148.1 | | 13.3 | 0.082 | yes |
| LinDA | Adaptive * | 0.591 [0.574, 0.608] | 0.492 [0.486, 0.498] | 0.065 [0.054, 0.076] | 0.575 [0.564, 0.586] | 9.4 | 147.8 | above | 14.6 | 0.0094 | yes |
| LDM | CLR | 0.609 [0.605, 0.614] | 0.512 [0.506, 0.518] | 0.035 [0.027, 0.042] | 0.596 [0.592, 0.600] | 16.7 | 144.5 | below | 12.2 | 0.11 | yes |
| LDM | Omni3 | 0.556 [0.543, 0.568] | 0.532 [0.520, 0.544] | 0.132 [0.113, 0.151] | 0.546 [0.538, 0.553] | 15.6 | 237.0 | above | 16.8 | 0.082 | yes |
| LinDA | Adaptive_NoWinsor | 0.591 [0.581, 0.601] | 0.482 [0.477, 0.487] | 0.049 [0.043, 0.054] | 0.574 [0.566, 0.581] | 9.4 | 139.9 | | 15.1 | 0.00091 | yes |
| MaAsLin 2 | CLR_NONE | 0.587 [0.574, 0.600] | 0.535 [0.530, 0.540] | 0.086 [0.069, 0.104] | 0.570 [0.558, 0.583] | 13.5 | 175.5 | above | 14.8 | 0.07 | yes |
| limma-voom | TMM_voom | 0.497 [0.463, 0.532] | 0.551 [0.499, 0.603] | 0.274 [0.199, 0.350] | 0.493 [0.452, 0.534] | 8.3 | 321.4 | above | 25.8 | 0.00013 | |
| LDM | PA | 0.544 [0.538, 0.550] | 0.524 [0.509, 0.539] | 0.135 [0.112, 0.158] | 0.535 [0.528, 0.542] | 16.7 | 238.7 | above | 20.8 | 0.065 | yes |
| LDM | Arcsin | 0.573 [0.564, 0.582] | 0.526 [0.513, 0.538] | 0.103 [0.086, 0.120] | 0.562 [0.552, 0.571] | 16.7 | 192.1 | above | 16.7 | 0.068 | yes |
| ZicoSeq | MultiLink | 0.604 [0.590, 0.619] | 0.491 [0.475, 0.506] | 0.061 [0.041, 0.081] | 0.587 [0.567, 0.607] | 7.3 | 143.0 | | 14.1 | 0.068 | yes |
| LinDA | Pseudo_1 | 0.607 [0.590, 0.623] | 0.494 [0.489, 0.500] | 0.028 [0.020, 0.036] | 0.585 [0.576, 0.594] | 9.4 | 138.6 | below | 12.3 | 0.062 | yes |
| ZicoSeq | NoPosterior | 0.638 [0.619, 0.656] | 0.529 [0.517, 0.542] | 0.046 [0.036, 0.056] | 0.619 [0.603, 0.635] | 6.2 | 150.9 | | 6.9 | | yes |
| MaAsLin 3 | TSS_LOG_Median * | 0.533 [0.519, 0.547] | 0.402 [0.393, 0.411] | 0.035 [0.000, 0.072] | 0.508 [0.495, 0.521] | 14.6 | 111.8 | | 25.5 | 0.00057 | |
| LDM | Omni * | 0.551 [0.545, 0.556] | 0.523 [0.511, 0.535] | 0.127 [0.107, 0.147] | 0.540 [0.536, 0.545] | 16.7 | 225.6 | above | 19.2 | 0.067 | yes |
| Wilcoxon | CLR | 0.427 [0.407, 0.448] | 0.457 [0.431, 0.482] | 0.188 [0.139, 0.237] | 0.406 [0.390, 0.421] | 15.6 | 348.3 | above | 30.5 | 0.00035 | |
| limma-voom | TMMwsp_voom | 0.394 [0.378, 0.411] | 0.575 [0.552, 0.599] | 0.354 [0.291, 0.416] | 0.401 [0.390, 0.412] | 8.3 | 664.1 | above | 27.4 | 0.0024 | |
| MaAsLin 3 | CLR_NONE | 0.525 [0.516, 0.534] | 0.420 [0.415, 0.425] | 0.072 [0.065, 0.079] | 0.504 [0.501, 0.507] | 13.5 | 130.8 | above | 25.4 | 0.0003 | |
| MaAsLin 3 | CLR_NONE_Median | 0.553 [0.539, 0.567] | 0.424 [0.414, 0.434] | 0.027 [0.020, 0.034] | 0.529 [0.518, 0.541] | 13.5 | 118.2 | below | 21.8 | 0.0029 | |
| MaAsLin 3 | TSS_LOG | 0.435 [0.421, 0.449] | 0.419 [0.412, 0.426] | 0.180 [0.142, 0.217] | 0.422 [0.413, 0.432] | 15.6 | 253.4 | above | 30.7 | 0.00084 | |
| DESeq2 | LRT_poscounts | 0.518 [0.507, 0.528] | 0.403 [0.400, 0.405] | 0.100 [0.080, 0.121] | 0.496 [0.487, 0.505] | 9.4 | 123.5 | above | 22.3 | 0.00015 | |
| DESeq2 | Wald_poscounts * | 0.502 [0.494, 0.511] | 0.382 [0.379, 0.385] | 0.131 [0.072, 0.189] | 0.479 [0.473, 0.485] | 6.2 | 118.8 | above | 24.4 | 0.00014 | |
| ZicoSeq | Default * | 0.600 [0.583, 0.618] | 0.487 [0.472, 0.502] | 0.058 [0.042, 0.074] | 0.582 [0.563, 0.602] | 6.2 | 141.6 | | 14.5 | 0.082 | yes |
| Wilcoxon | Rarefied | 0.379 [0.365, 0.393] | 0.282 [0.270, 0.294] | 0.054 [0.046, 0.063] | 0.349 [0.334, 0.365] | 24.0 | 92.8 | | 33.1 | 3.6e-06 | |
| edgeR | TMM_LRT | 0.503 [0.485, 0.521] | 0.501 [0.488, 0.514] | 0.388 [0.362, 0.415] | 0.516 [0.499, 0.532] | 0.0 | 190.3 | above | 21.1 | 0.00085 | yes |
| LDM | Freq | 0.496 [0.482, 0.509] | 0.447 [0.437, 0.456] | 0.113 [0.094, 0.131] | 0.479 [0.468, 0.489] | 19.8 | 178.7 | above | 24.8 | 0.00035 | |
| ALDEx2 | Wilcox_128 | 0.341 [0.331, 0.351] | 0.211 [0.204, 0.219] | 0.000 [0.000, 0.000] | 0.296 [0.286, 0.307] | 27.1 | 57.1 | below | 34.4 | 5.1e-07 | |
| edgeR | TMM_QL * | 0.482 [0.471, 0.493] | 0.544 [0.534, 0.553] | 0.465 [0.453, 0.477] | 0.504 [0.490, 0.519] | 0.0 | 248.4 | above | 24.1 | 5.6e-05 | |
| ANCOM-BC2 | BH | 0.413 [0.407, 0.419] | 0.277 [0.270, 0.283] | 0.006 [0.000, 0.013] | 0.367 [0.359, 0.375] | 18.8 | 75.0 | below | 28.3 | 4.9e-05 | |
| ALDEx2 | Welch_128 * | 0.271 [0.266, 0.276] | 0.161 [0.159, 0.163] | 0.000 [0.000, 0.000] | 0.229 [0.226, 0.233] | 36.5 | 43.5 | below | 37.2 | 3e-07 | |
| ALDEx2 | Welch_32 | 0.275 [0.265, 0.285] | 0.163 [0.159, 0.168] | 0.000 [0.000, 0.000] | 0.233 [0.226, 0.240] | 36.5 | 44.1 | below | 36.7 | 3e-07 | |
| ALDEx2 | Welch_128_IQLR | 0.276 [0.267, 0.285] | 0.165 [0.161, 0.169] | 0.000 [0.000, 0.000] | 0.235 [0.228, 0.242] | 36.5 | 44.6 | below | 36.7 | 4.4e-07 | |
| ANCOM-BC2 | BH_StrucZero | 0.377 [0.372, 0.382] | 0.240 [0.234, 0.245] | 0.006 [0.000, 0.015] | 0.328 [0.321, 0.335] | 21.9 | 65.0 | below | 32.0 | 7.7e-06 | |
| MaAsLin 2 | TMM_NONE | 0.408 [0.387, 0.430] | 0.282 [0.273, 0.292] | 0.009 [0.002, 0.016] | 0.372 [0.362, 0.383] | 25.0 | 77.8 | below | 30.3 | 4e-06 | |
| ANCOM-BC2 | Holm * | 0.261 [0.253, 0.270] | 0.159 [0.154, 0.165] | 0.000 [0.000, 0.000] | 0.219 [0.211, 0.227] | 33.3 | 43.0 | below | 35.5 | 4e-06 | |
| ANCOM-BC2 | Holm_NoFilter | 0.280 [0.274, 0.286] | 0.333 [0.329, 0.337] | 0.621 [0.605, 0.637] | 0.320 [0.316, 0.324] | 0.0 | 223.7 | above | 32.1 | 0.00027 | |
| ANCOM-BC2 | BH_NoFilter | 0.119 [0.112, 0.126] | 0.521 [0.514, 0.529] | 0.854 [0.850, 0.858] | 0.226 [0.222, 0.231] | 0.0 | 942.6 | above | 37.3 | 2.5e-07 | |
| ***Simulated stool, genus level (40 scenarios, 160 datasets). Friedman chi-squared 838.6, P 9.8e-149, Kendall's W 0.50, Nemenyi critical difference 11.0*** | | | | | | | | | | | |
| LinDA | Pseudo_0.1 | 0.637 [0.593, 0.682] | 0.563 [0.518, 0.608] | 0.089 [0.047, 0.132] | 0.623 [0.587, 0.659] | 8.1 | 5.6 | | 10.5 | 1 | yes |
| MaAsLin 2 | TMM_LOG | 0.637 [0.609, 0.664] | 0.569 [0.534, 0.604] | 0.079 [0.043, 0.115] | 0.623 [0.599, 0.648] | 11.9 | 5.3 | | 10.1 | 1 | yes |
| LinDA | Imputation | 0.619 [0.574, 0.664] | 0.562 [0.530, 0.595] | 0.114 [0.076, 0.153] | 0.609 [0.572, 0.647] | 10.0 | 5.9 | above | 12.8 | 0.095 | yes |

| Tool | Configuration | MCC | Recall | FDR | F1 | No call (%) | Calls per dataset | FDR against 0.05 | Mean rank | Holm P vs best | Within critical difference of best |
|---|---|---|---|---|---|---|---|---|---|---|---|
| MaAsLin 3 | TSS_PLOG_Median | 0.628 [0.603, 0.654] | 0.534 [0.511, 0.558] | 0.058 [0.041, 0.074] | 0.613 [0.593, 0.633] | 11.2 | 4.9 | | 11.6 | 0.37 | yes |
| LinDA | Pseudo_0.5 | 0.618 [0.582, 0.655] | 0.543 [0.506, 0.580] | 0.100 [0.066, 0.133] | 0.605 [0.574, 0.637] | 8.8 | 5.5 | above | 13.6 | 0.13 | yes |
| MaAsLin 3 | TSS_PLOG | 0.640 [0.618, 0.663] | 0.585 [0.548, 0.622] | 0.096 [0.078, 0.115] | 0.627 [0.611, 0.644] | 10.0 | 6.0 | above | 8.2 | | yes |
| MaAsLin 2 | TSS_LOG * | 0.624 [0.600, 0.648] | 0.577 [0.542, 0.612] | 0.098 [0.070, 0.127] | 0.613 [0.594, 0.632] | 12.5 | 6.1 | above | 11.2 | 0.13 | yes |
| Wilcoxon | TSS | 0.625 [0.588, 0.663] | 0.577 [0.537, 0.617] | 0.104 [0.077, 0.131] | 0.615 [0.582, 0.647] | 11.9 | 6.0 | above | 11.9 | 0.15 | yes |
| LinDA | Adaptive * | 0.620 [0.589, 0.652] | 0.545 [0.514, 0.577] | 0.098 [0.066, 0.129] | 0.607 [0.580, 0.633] | 8.8 | 5.5 | above | 13.2 | 0.15 | yes |
| LDM | CLR | 0.615 [0.588, 0.643] | 0.564 [0.529, 0.600] | 0.102 [0.073, 0.132] | 0.606 [0.582, 0.631] | 12.5 | 6.1 | above | 13.1 | 0.095 | yes |
| LDM | Omni3 | 0.604 [0.571, 0.637] | 0.573 [0.548, 0.599] | 0.116 [0.095, 0.136] | 0.594 [0.567, 0.621] | 14.4 | 6.5 | above | 13.8 | 0.0047 | yes |
| LinDA | Adaptive_NoWinsor | 0.622 [0.592, 0.651] | 0.545 [0.513, 0.576] | 0.093 [0.069, 0.118] | 0.609 [0.586, 0.632] | 9.4 | 5.3 | above | 12.3 | 0.15 | yes |
| MaAsLin 2 | CLR_NONE | 0.584 [0.568, 0.600] | 0.546 [0.510, 0.582] | 0.127 [0.085, 0.168] | 0.578 [0.563, 0.593] | 14.4 | 6.3 | above | 18.0 | 0.00012 | yes |
| limma-voom | TMM_voom | 0.550 [0.523, 0.576] | 0.511 [0.495, 0.526] | 0.149 [0.110, 0.188] | 0.541 [0.521, 0.560] | 11.9 | 7.2 | above | 23.2 | 3e-07 | |
| LDM | PA | 0.603 [0.580, 0.626] | 0.555 [0.532, 0.577] | 0.076 [0.041, 0.112] | 0.593 [0.577, 0.609] | 17.5 | 5.8 | | 13.8 | 0.026 | yes |
| LDM | Arcsin | 0.584 [0.561, 0.608] | 0.560 [0.516, 0.605] | 0.120 [0.092, 0.149] | 0.576 [0.559, 0.594] | 16.9 | 6.5 | above | 17.1 | 0.00047 | yes |
| ZicoSeq | MultiLink | 0.604 [0.564, 0.644] | 0.512 [0.465, 0.558] | 0.075 [0.061, 0.088] | 0.590 [0.552, 0.628] | 12.5 | 4.5 | above | 15.9 | 0.17 | yes |
| LinDA | Pseudo_1 | 0.600 [0.566, 0.633] | 0.525 [0.489, 0.561] | 0.097 [0.050, 0.145] | 0.588 [0.562, 0.613] | 11.2 | 5.4 | | 15.9 | 0.0021 | yes |
| ZicoSeq | NoPosterior | 0.592 [0.559, 0.625] | 0.502 [0.469, 0.536] | 0.065 [0.022, 0.107] | 0.573 [0.539, 0.608] | 14.4 | 4.3 | | 17.6 | 0.15 | yes |
| MaAsLin 3 | TSS_LOG_Median * | 0.598 [0.557, 0.639] | 0.494 [0.449, 0.539] | 0.030 [0.009, 0.051] | 0.580 [0.539, 0.621] | 16.2 | 4.2 | | 17.7 | 0.017 | yes |
| LDM | Omni * | 0.577 [0.546, 0.609] | 0.549 [0.520, 0.579] | 0.122 [0.090, 0.153] | 0.569 [0.545, 0.593] | 16.9 | 6.4 | above | 18.6 | 0.00011 | yes |
| Wilcoxon | CLR | 0.531 [0.516, 0.546] | 0.471 [0.415, 0.527] | 0.123 [0.099, 0.146] | 0.518 [0.494, 0.543] | 13.8 | 6.3 | above | 25.3 | 3.9e-09 | |
| limma-voom | TMMwsp_voom | 0.543 [0.500, 0.587] | 0.526 [0.491, 0.560] | 0.184 [0.136, 0.232] | 0.541 [0.507, 0.576] | 11.9 | 7.7 | above | 23.7 | 1.6e-08 | |
| MaAsLin 3 | CLR_NONE | 0.527 [0.475, 0.579] | 0.430 [0.383, 0.478] | 0.090 [0.055, 0.124] | 0.508 [0.450, 0.566] | 15.0 | 4.3 | above | 26.4 | 8.2e-11 | |
| MaAsLin 3 | CLR_NONE_Median | 0.549 [0.499, 0.598] | 0.431 [0.378, 0.484] | 0.055 [0.024, 0.085] | 0.527 [0.472, 0.582] | 15.6 | 3.8 | | 23.2 | 2.6e-06 | |
| MaAsLin 3 | TSS_LOG | 0.566 [0.536, 0.597] | 0.494 [0.459, 0.528] | 0.085 [0.033, 0.137] | 0.554 [0.526, 0.581] | 16.2 | 5.2 | | 21.8 | 3e-07 | |
| DESeq2 | LRT_poscounts | 0.458 [0.431, 0.485] | 0.411 [0.372, 0.450] | 0.231 [0.161, 0.302] | 0.455 [0.425, 0.485] | 15.0 | 4.7 | above | 29.8 | 6.5e-07 | |
| DESeq2 | Wald_poscounts * | 0.453 [0.423, 0.484] | 0.407 [0.373, 0.441] | 0.254 [0.172, 0.335] | 0.451 [0.420, 0.481] | 12.5 | 4.7 | above | 29.7 | 2.6e-07 | |
| ZicoSeq | Default * | 0.555 [0.522, 0.588] | 0.463 [0.434, 0.491] | 0.068 [0.011, 0.126] | 0.540 [0.509, 0.571] | 18.1 | 4.2 | | 21.0 | 0.0036 | |
| Wilcoxon | Rarefied | 0.515 [0.468, 0.562] | 0.443 [0.401, 0.485] | 0.099 [0.051, 0.148] | 0.498 [0.460, 0.536] | 18.8 | 4.8 | above | 25.6 | 7.4e-08 | |
| edgeR | TMM_LRT | 0.492 [0.464, 0.520] | 0.530 [0.497, 0.564] | 0.323 [0.255, 0.391] | 0.503 [0.483, 0.523] | 12.5 | 6.8 | above | 26.4 | 8.3e-06 | |
| LDM | Freq | 0.518 [0.479, 0.557] | 0.468 [0.438, 0.498] | 0.101 [0.035, 0.167] | 0.501 [0.470, 0.532] | 21.2 | 5.2 | | 22.4 | 2e-05 | |
| ALDEx2 | Wilcox_128 | 0.435 [0.406, 0.465] | 0.329 [0.294, 0.363] | 0.045 [0.014, 0.076] | 0.407 [0.380, 0.435] | 27.5 | 3.1 | | 31.9 | 2.1e-10 | |
| edgeR | TMM_QL * | 0.419 [0.385, 0.453] | 0.627 [0.602, 0.653] | 0.593 [0.524, 0.663] | 0.452 [0.426, 0.478] | 2.5 | 12.9 | above | 32.3 | 8.7e-10 | |
| ANCOM-BC2 | BH | 0.393 [0.356, 0.431] | 0.299 [0.260, 0.338] | 0.006 [0.000, 0.013] | 0.371 [0.336, 0.406] | 40.6 | 2.5 | below | 30.5 | 2.6e-06 | |
| ALDEx2 | Welch_128 * | 0.419 [0.379, 0.458] | 0.308 [0.263, 0.352] | 0.031 [0.000, 0.067] | 0.388 [0.349, 0.426] | 29.4 | 2.8 | | 33.2 | 2.6e-11 | |
| ALDEx2 | Welch_32 | 0.419 [0.381, 0.456] | 0.309 [0.272, 0.346] | 0.032 [0.000, 0.068] | 0.389 [0.353, 0.425] | 30.0 | 2.8 | | 32.9 | 7.6e-11 | |
| ALDEx2 | Welch_128_IQLR | 0.420 [0.384, 0.455] | 0.305 [0.259, 0.351] | 0.020 [0.000, 0.057] | 0.389 [0.354, 0.423] | 30.6 | 2.7 | | 32.5 | 1.3e-09 | |
| ANCOM-BC2 | BH_StrucZero | 0.362 [0.334, 0.390] | 0.267 [0.235, 0.300] | 0.006 [0.000, 0.013] | 0.338 [0.312, 0.363] | 43.1 | 2.2 | below | 33.5 | 1.5e-07 | |
| MaAsLin 2 | TMM_NONE | 0.400 [0.366, 0.434] | 0.318 [0.301, 0.335] | 0.032 [0.001, 0.063] | 0.379 [0.348, 0.410] | 38.1 | 2.8 | | 32.9 | 4.7e-08 | |
| ANCOM-BC2 | Holm * | 0.339 [0.310, 0.368] | 0.253 [0.220, 0.286] | 0.000 [0.000, 0.000] | 0.318 [0.288, 0.348] | 48.8 | 2.0 | below | 34.6 | 4e-07 | |
| ANCOM-BC2 | Holm_NoFilter | 0.283 [0.234, 0.331] | 0.375 [0.322, 0.428] | 0.664 [0.604, 0.724] | 0.337 [0.291, 0.383] | 0.6 | 8.2 | above | 37.9 | 7.2e-10 | |
| ANCOM-BC2 | BH_NoFilter | 0.234 [0.191, 0.276] | 0.430 [0.383, 0.478] | 0.751 [0.714, 0.788] | 0.307 [0.270, 0.344] | 0.0 | 13.6 | above | 38.5 | 1.8e-11 | |

**Table S4. Twenty replicates per scenario.** Main detection design on stool genera and null design on stool and IBD genera, extended from four to 20 replicates per scenario (480 main and 80 null datasets). Replicates 1 to 4 are the benchmark datasets, regenerated from their seeds with identical results. Means with 95% confidence intervals across replicates. Null datasets with a call are given with exact binomial 95% intervals, out of 60 stool datasets at 50, 100 and 200 samples and 20 IBD datasets at 50 samples. The FDR is controlled when the upper limit is 0.05 or less, and exceeded when the lower limit is above 0.05.

| Tool | Configuration | MCC, 4 replicates | MCC, 20 replicates | FDR, 4 replicates | FDR, 20 replicates | FDR verdict, 20 replicates | Null datasets with a call, stool | Null datasets with a call, IBD | False calls per stool null dataset |
|---|---|---|---|---|---|---|---|---|---|
| LinDA | Pseudo_0.1 | 0.675 [0.637, 0.714] | 0.670 [0.658, 0.682] | 0.055 [0.034, 0.076] | 0.053 [0.041, 0.064] | not distinguishable | 5 of 60 [0.03, 0.18] | 0 of 20 [0.00, 0.17] | 0.20 |
| MaAsLin 2 | TMM_LOG | 0.675 [0.628, 0.723] | 0.664 [0.651, 0.676] | 0.041 [0.029, 0.054] | 0.038 [0.028, 0.047] | controlled | 5 of 60 [0.03, 0.18] | 0 of 20 [0.00, 0.17] | 0.18 |
| LinDA | Imputation | 0.662 [0.624, 0.699] | 0.652 [0.637, 0.666] | 0.059 [0.004, 0.114] | 0.063 [0.051, 0.075] | exceeded | 6 of 60 [0.04, 0.21] | 1 of 20 [0.00, 0.25] | 0.22 |
| MaAsLin 3 | TSS_PLOG_Median | 0.654 [0.596, 0.712] | 0.647 [0.630, 0.663] | 0.034 [0.000, 0.080] | 0.025 [0.018, 0.033] | controlled | 5 of 60 [0.03, 0.18] | 0 of 20 [0.00, 0.17] | 0.20 |
| LinDA | Pseudo_0.5 | 0.650 [0.625, 0.675] | 0.639 [0.627, 0.652] | 0.048 [0.015, 0.082] | 0.047 [0.038, 0.056] | not distinguishable | 6 of 60 [0.04, 0.21] | 0 of 20 [0.00, 0.17] | 0.28 |
| MaAsLin 3 | TSS_PLOG | 0.647 [0.601, 0.693] | 0.655 [0.642, 0.667] | 0.061 [0.009, 0.114] | 0.051 [0.040, 0.063] | not distinguishable | 5 of 60 [0.03, 0.18] | 0 of 20 [0.00, 0.17] | 0.17 |
| MaAsLin 2 | TSS_LOG * | 0.641 [0.604, 0.677] | 0.644 [0.631, 0.656] | 0.064 [0.010, 0.119] | 0.051 [0.040, 0.062] | not distinguishable | 5 of 60 [0.03, 0.18] | 0 of 20 [0.00, 0.17] | 0.17 |
| Wilcoxon | TSS | 0.639 [0.598, 0.680] | 0.641 [0.625, 0.658] | 0.068 [0.003, 0.133] | 0.058 [0.044, 0.071] | not distinguishable | 6 of 60 [0.04, 0.21] | 0 of 20 [0.00, 0.17] | 0.15 |
| LinDA | Adaptive * | 0.634 [0.611, 0.657] | 0.630 [0.616, 0.644] | 0.059 [0.020, 0.098] | 0.060 [0.047, 0.072] | not distinguishable | 6 of 60 [0.04, 0.21] | 0 of 20 [0.00, 0.17] | 0.17 |
| LDM | CLR | 0.631 [0.623, 0.638] | 0.635 [0.618, 0.652] | 0.035 [0.000, 0.075] | 0.038 [0.031, 0.044] | controlled | 2 of 60 [0.00, 0.12] | 0 of 20 [0.00, 0.17] | 0.22 |
| LDM | Omni3 | 0.629 [0.579, 0.679] | 0.631 [0.614, 0.647] | 0.074 [0.029, 0.120] | 0.068 [0.057, 0.080] | exceeded | 7 of 60 [0.05, 0.23] | 1 of 20 [0.00, 0.25] | 0.25 |
| LinDA | Adaptive_NoWinsor | 0.626 [0.591, 0.661] | 0.627 [0.611, 0.642] | 0.060 [0.015, 0.104] | 0.060 [0.048, 0.072] | not distinguishable | 6 of 60 [0.04, 0.21] | 0 of 20 [0.00, 0.17] | 0.18 |
| MaAsLin 2 | CLR_NONE | 0.613 [0.605, 0.620] | 0.607 [0.596, 0.618] | 0.048 [0.014, 0.082] | 0.054 [0.045, 0.062] | not distinguishable | 4 of 60 [0.02, 0.16] | 0 of 20 [0.00, 0.17] | 0.45 |
| limma-voom | TMM_voom | 0.612 [0.569, 0.655] | 0.586 [0.569, 0.603] | 0.094 [0.049, 0.139] | 0.095 [0.077, 0.113] | exceeded | 7 of 60 [0.05, 0.23] | 3 of 20 [0.03, 0.38] | 0.28 |
| LDM | PA | 0.606 [0.552, 0.659] | 0.617 [0.598, 0.637] | 0.056 [0.004, 0.108] | 0.056 [0.044, 0.068] | not distinguishable | 5 of 60 [0.03, 0.18] | 1 of 20 [0.00, 0.25] | 0.18 |
| LDM | Arcsin | 0.604 [0.554, 0.655] | 0.604 [0.584, 0.624] | 0.081 [0.045, 0.116] | 0.068 [0.056, 0.080] | exceeded | 7 of 60 [0.05, 0.23] | 0 of 20 [0.00, 0.17] | 0.18 |
| ZicoSeq | MultiLink | 0.604 [0.571, 0.636] | 0.587 [0.572, 0.601] | 0.072 [0.050, 0.094] | 0.079 [0.064, 0.094] | exceeded | 5 of 60 [0.03, 0.18] | 1 of 20 [0.00, 0.25] | 0.20 |
| LinDA | Pseudo_1 | 0.603 [0.586, 0.620] | 0.610 [0.598, 0.623] | 0.045 [0.011, 0.079] | 0.044 [0.034, 0.053] | not distinguishable | 7 of 60 [0.05, 0.23] | 0 of 20 [0.00, 0.17] | 0.30 |
| ZicoSeq | NoPosterior | 0.602 [0.567, 0.637] | 0.591 [0.576, 0.605] | 0.042 [0.032, 0.052] | 0.043 [0.034, 0.051] | not distinguishable | 4 of 60 [0.02, 0.16] | 0 of 20 [0.00, 0.17] | 0.12 |
| MaAsLin 3 | TSS_LOG_Median * | 0.598 [0.580, 0.617] | 0.583 [0.570, 0.596] | 0.031 [0.009, 0.053] | 0.024 [0.018, 0.031] | controlled | 3 of 60 [0.01, 0.14] | 1 of 20 [0.00, 0.25] | 0.10 |
| LDM | Omni * | 0.595 [0.529, 0.660] | 0.594 [0.575, 0.613] | 0.090 [0.063, 0.118] | 0.070 [0.057, 0.083] | exceeded | 7 of 60 [0.05, 0.23] | 1 of 20 [0.00, 0.25] | 0.17 |
| Wilcoxon | CLR | 0.590 [0.545, 0.635] | 0.578 [0.563, 0.593] | 0.059 [0.037, 0.080] | 0.063 [0.054, 0.073] | exceeded | 4 of 60 [0.02, 0.16] | 1 of 20 [0.00, 0.25] | 0.82 |
| limma-voom | TMMwsp_voom | 0.578 [0.564, 0.593] | 0.573 [0.557, 0.589] | 0.085 [0.000, 0.189] | 0.097 [0.078, 0.117] | exceeded | 7 of 60 [0.05, 0.23] | 0 of 20 [0.00, 0.17] | 1.48 |
| MaAsLin 3 | CLR_NONE | 0.578 [0.548, 0.607] | 0.560 [0.547, 0.574] | 0.038 [0.017, 0.059] | 0.042 [0.031, 0.053] | not distinguishable | 4 of 60 [0.02, 0.16] | 0 of 20 [0.00, 0.17] | 0.12 |
| MaAsLin 3 | CLR_NONE_Median | 0.577 [0.541, 0.614] | 0.559 [0.546, 0.573] | 0.017 [0.000, 0.034] | 0.024 [0.014, 0.034] | controlled | 2 of 60 [0.00, 0.12] | 0 of 20 [0.00, 0.17] | 0.07 |
| MaAsLin 3 | TSS_LOG | 0.570 [0.508, 0.632] | 0.562 [0.548, 0.575] | 0.086 [0.040, 0.132] | 0.067 [0.057, 0.078] | exceeded | 3 of 60 [0.01, 0.14] | 0 of 20 [0.00, 0.17] | 0.10 |
| DESeq2 | LRT_poscounts | 0.540 [0.509, 0.571] | 0.530 [0.514, 0.546] | 0.136 [0.071, 0.201] | 0.137 [0.115, 0.159] | exceeded | 22 of 60 [0.25, 0.50] | 4 of 20 [0.06, 0.44] | 0.57 |
| DESeq2 | Wald_poscounts * | 0.536 [0.499, 0.573] | 0.526 [0.509, 0.542] | 0.163 [0.084, 0.242] | 0.163 [0.139, 0.188] | exceeded | 23 of 60 [0.26, 0.52] | 12 of 20 [0.36, 0.81] | 0.63 |
| ZicoSeq | Default * | 0.518 [0.458, 0.579] | 0.521 [0.506, 0.537] | 0.116 [0.075, 0.158] | 0.100 [0.083, 0.118] | exceeded | 3 of 60 [0.01, 0.14] | 0 of 20 [0.00, 0.17] | 0.12 |
| Wilcoxon | Rarefied | 0.505 [0.483, 0.526] | 0.509 [0.498, 0.520] | 0.080 [0.056, 0.104] | 0.054 [0.042, 0.065] | not distinguishable | 2 of 60 [0.00, 0.12] | 0 of 20 [0.00, 0.17] | 0.07 |
| edgeR | TMM_LRT | 0.496 [0.468, 0.524] | 0.496 [0.476, 0.517] | 0.341 [0.308, 0.374] | 0.353 [0.322, 0.385] | exceeded | 40 of 60 [0.53, 0.78] | 15 of 20 [0.51, 0.91] | 1.80 |
| LDM | Freq | 0.494 [0.470, 0.518] | 0.505 [0.487, 0.522] | 0.081 [0.028, 0.134] | 0.059 [0.046, 0.073] | not distinguishable | 5 of 60 [0.03, 0.18] | 1 of 20 [0.00, 0.25] | 0.12 |
| ALDEx2 | Wilcox_128 | 0.461 [0.411, 0.511] | 0.454 [0.439, 0.468] | 0.005 [0.000, 0.016] | 0.004 [0.001, 0.007] | controlled | 1 of 60 [0.00, 0.09] | 0 of 20 [0.00, 0.17] | 0.02 |
| edgeR | TMM_QL * | 0.434 [0.394, 0.474] | 0.421 [0.406, 0.437] | 0.571 [0.527, 0.615] | 0.583 [0.564, 0.602] | exceeded | 56 of 60 [0.84, 0.98] | 20 of 20 [0.83, 1.00] | 5.70 |
| ANCOM-BC2 | BH | 0.420 [0.355, 0.485] | 0.403 [0.387, 0.418] | 0.005 [0.000, 0.016] | 0.003 [0.001, 0.006] | controlled | 1 of 60 [0.00, 0.09] | 0 of 20 [0.00, 0.17] | 0.02 |
| ALDEx2 | Welch_128 * | 0.411 [0.372, 0.450] | 0.410 [0.397, 0.423] | 0.000 [0.000, 0.000] | 0.002 [0.000, 0.004] | controlled | 1 of 60 [0.00, 0.09] | 0 of 20 [0.00, 0.17] | 0.02 |
| ALDEx2 | Welch_32 | 0.407 [0.377, 0.437] | 0.410 [0.399, 0.421] | 0.000 [0.000, 0.000] | 0.003 [0.000, 0.006] | controlled | 1 of 60 [0.00, 0.09] | 0 of 20 [0.00, 0.17] | 0.02 |
| ALDEx2 | Welch_128_IQLR | 0.406 [0.368, 0.444] | 0.401 [0.386, 0.416] | 0.000 [0.000, 0.000] | 0.002 [0.000, 0.004] | controlled | 1 of 60 [0.00, 0.09] | 0 of 20 [0.00, 0.17] | 0.02 |
| ANCOM-BC2 | BH_StrucZero | 0.404 [0.355, 0.453] | 0.387 [0.372, 0.401] | 0.005 [0.000, 0.016] | 0.003 [0.001, 0.006] | controlled | 1 of 60 [0.00, 0.09] | 0 of 20 [0.00, 0.17] | 0.02 |
| MaAsLin 2 | TMM_NONE | 0.366 [0.360, 0.372] | 0.347 [0.331, 0.362] | 0.025 [0.000, 0.055] | 0.012 [0.004, 0.020] | controlled | 1 of 60 [0.00, 0.09] | 0 of 20 [0.00, 0.17] | 0.02 |
| ANCOM-BC2 | Holm * | 0.353 [0.328, 0.379] | 0.344 [0.332, 0.357] | 0.000 [0.000, 0.000] | 0.000 [0.000, 0.000] | controlled | 1 of 60 [0.00, 0.09] | 0 of 20 [0.00, 0.17] | 0.02 |
| ANCOM-BC2 | Holm_NoFilter | 0.309 [0.261, 0.357] | 0.308 [0.296, 0.320] | 0.647 [0.573, 0.722] | 0.645 [0.626, 0.663] | exceeded | 58 of 60 [0.88, 1.00] | 20 of 20 [0.83, 1.00] | 6.67 |

| Tool | Configuration | MCC, 4 replicates | MCC, 20 replicates | FDR, 4 replicates | FDR, 20 replicates | FDR verdict, 20 replicates | Null datasets with a call, stool | Null datasets with a call, IBD | False calls per stool null dataset |
|---|---|---|---|---|---|---|---|---|---|
| ANCOM-BC2 | BH_NoFilter | 0.273 [0.221, 0.326] | 0.260 [0.247, 0.273] | 0.739 [0.694, 0.785] | 0.747 [0.735, 0.759] | exceeded | 60 of 60 [0.94, 1.00] | 20 of 20 [0.83, 1.00] | 14.20 |

**Table S5. Fixed read depth and relative truth.** Stool genera, four replicates. Observed refers to the benchmark datasets, in which implantation changes the read depth of the cases. Fixed depth refers to the same datasets after every sample was resampled from a multinomial distribution to its read depth before implantation. Without resampling, the read depth of the cases increased 1.9-fold on average with signals in one direction and 208-fold with a single extreme feature. The relative truth adds every genus whose mean relative abundance in the cases changed by more than 0.2 on the natural-log scale. In the null design every call is false, and the FDR equals the proportion of datasets with at least one call, with exact binomial 95% intervals. Across configurations, the Spearman correlation between observed and fixed depth in the main detection design was 0.99 for MCC and 0.87 for FDR. Configurations marked † are not among the 43, namely the edgeR user-guide pipeline with filterByExpr and robust fits, the edgeR 3 quasi-likelihood fit and DESeq2 with its settings for sparse counts.

| Tool | Configuration | MCC observed | MCC fixed depth | FDR observed | FDR fixed depth | FDR relative truth, observed | FDR relative truth, fixed | Calls per dataset, observed | Calls per dataset, fixed |
|---|---|---|---|---|---|---|---|---|---|
| ***Main detection design (24 scenarios)*** | | | | | | | | | |
| LinDA | Pseudo_0.1 | 0.675 | 0.629 | 0.055 [0.034, 0.076] | 0.065 [0.037, 0.092] | 0.043 | 0.032 | 5.2 | 5.5 |
| MaAsLin 2 | TMM_LOG | 0.675 | 0.622 | 0.041 [0.029, 0.054] | 0.047 [0.009, 0.085] | 0.028 | 0.024 | 5.1 | 5.1 |
| LinDA | Imputation | 0.662 | 0.627 | 0.059 [0.004, 0.114] | 0.050 [0.009, 0.091] | 0.027 | 0.019 | 5.8 | 5.5 |
| MaAsLin 3 | TSS_PLOG_Median | 0.654 | 0.609 | 0.034 [0.000, 0.080] | 0.027 [0.000, 0.077] | 0.013 | 0.007 | 5.0 | 4.7 |
| LinDA | Pseudo_0.5 | 0.650 | 0.603 | 0.048 [0.015, 0.082] | 0.066 [0.025, 0.107] | 0.036 | 0.029 | 4.9 | 5.3 |
| MaAsLin 3 | TSS_PLOG | 0.647 | 0.612 | 0.061 [0.009, 0.114] | 0.059 [0.007, 0.111] | 0.028 | 0.024 | 5.5 | 5.6 |
| MaAsLin 2 | TSS_LOG * | 0.641 | 0.597 | 0.064 [0.010, 0.119] | 0.050 [0.000, 0.111] | 0.026 | 0.016 | 5.6 | 5.4 |
| Wilcoxon | TSS | 0.639 | 0.587 | 0.068 [0.003, 0.133] | 0.055 [0.010, 0.101] | 0.033 | 0.020 | 5.5 | 5.3 |
| LinDA | Adaptive * | 0.634 | 0.607 | 0.059 [0.020, 0.098] | 0.058 [0.019, 0.097] | 0.026 | 0.021 | 5.5 | 5.3 |
| LDM | CLR | 0.631 | 0.585 | 0.035 [0.000, 0.075] | 0.056 [0.022, 0.090] | 0.027 | 0.025 | 4.8 | 5.3 |
| LDM | Omni3 | 0.629 | 0.594 | 0.074 [0.029, 0.120] | 0.069 [0.026, 0.112] | 0.032 | 0.029 | 6.4 | 5.8 |
| LinDA | Adaptive_NoWinsor | 0.626 | 0.611 | 0.060 [0.015, 0.104] | 0.060 [0.015, 0.105] | 0.027 | 0.026 | 5.3 | 5.3 |
| MaAsLin 2 | CLR_NONE | 0.613 | 0.584 | 0.048 [0.014, 0.082] | 0.053 [0.016, 0.090] | 0.029 | 0.020 | 5.0 | 5.5 |
| limma-voom | TMM_voom | 0.612 | 0.550 | 0.094 [0.049, 0.139] | 0.115 [0.097, 0.133] | 0.063 | 0.058 | 6.2 | 7.2 |
| LDM | PA | 0.606 | 0.579 | 0.056 [0.004, 0.108] | 0.040 [0.000, 0.097] | 0.022 | 0.013 | 5.6 | 5.1 |
| LDM | Arcsin | 0.604 | 0.565 | 0.081 [0.045, 0.116] | 0.078 [0.041, 0.115] | 0.036 | 0.037 | 6.1 | 5.7 |
| ZicoSeq | MultiLink | 0.604 | 0.583 | 0.072 [0.050, 0.094] | 0.039 [0.035, 0.044] | 0.054 | 0.019 | 5.0 | 4.2 |
| LinDA | Pseudo_1 | 0.603 | 0.575 | 0.045 [0.011, 0.079] | 0.067 [0.027, 0.107] | 0.033 | 0.028 | 4.5 | 5.1 |
| ZicoSeq | NoPosterior | 0.602 | 0.549 | 0.042 [0.032, 0.052] | 0.035 [0.009, 0.060] | 0.025 | 0.012 | 4.4 | 4.0 |
| MaAsLin 3 | TSS_LOG_Median * | 0.598 | 0.521 | 0.031 [0.009, 0.053] | 0.032 [0.007, 0.058] | 0.027 | 0.017 | 4.2 | 3.8 |
| LDM | Omni * | 0.595 | 0.549 | 0.090 [0.063, 0.118] | 0.085 [0.050, 0.121] | 0.042 | 0.040 | 6.2 | 5.6 |
| Wilcoxon | CLR | 0.590 | 0.533 | 0.059 [0.037, 0.080] | 0.054 [0.000, 0.125] | 0.041 | 0.022 | 5.1 | 5.4 |
| limma-voom | TMMwsp_voom | 0.578 | 0.545 | 0.085 [0.000, 0.189] | 0.105 [0.015, 0.195] | 0.053 | 0.067 | 6.3 | 6.6 |
| MaAsLin 3 | CLR_NONE | 0.578 | 0.494 | 0.038 [0.017, 0.059] | 0.052 [0.017, 0.088] | 0.028 | 0.024 | 4.0 | 3.9 |
| MaAsLin 3 | CLR_NONE_Median | 0.577 | 0.506 | 0.017 [0.000, 0.034] | 0.031 [0.000, 0.074] | 0.015 | 0.017 | 3.8 | 3.5 |
| MaAsLin 3 | TSS_LOG | 0.570 | 0.512 | 0.086 [0.040, 0.132] | 0.061 [0.022, 0.099] | 0.037 | 0.020 | 5.8 | 4.6 |
| DESeq2 | LRT_poscounts | 0.540 | 0.496 | 0.136 [0.071, 0.201] | 0.136 [0.051, 0.220] | 0.113 | 0.101 | 4.7 | 4.4 |
| DESeq2 | Wald_poscounts * | 0.536 | 0.490 | 0.163 [0.084, 0.242] | 0.171 [0.087, 0.255] | 0.139 | 0.134 | 4.7 | 4.5 |
| ZicoSeq | Default * | 0.518 | 0.504 | 0.116 [0.075, 0.158] | 0.051 [0.001, 0.101] | 0.098 | 0.030 | 5.0 | 3.8 |
| Wilcoxon | Rarefied | 0.505 | 0.490 | 0.080 [0.056, 0.104] | 0.051 [0.000, 0.104] | 0.034 | 0.014 | 4.5 | 4.3 |
| edgeR | TMM_LRT | 0.496 | 0.484 | 0.341 [0.308, 0.374] | 0.322 [0.278, 0.365] | 0.293 | 0.262 | 7.2 | 6.7 |
| LDM | Freq | 0.494 | 0.486 | 0.081 [0.028, 0.134] | 0.075 [0.027, 0.123] | 0.037 | 0.033 | 4.9 | 4.7 |
| ALDEx2 | Wilcox_128 | 0.461 | 0.429 | 0.005 [0.000, 0.016] | 0.025 [0.000, 0.072] | 0.003 | 0.006 | 2.8 | 2.9 |
| edgeR | TMM_QL * | 0.434 | 0.425 | 0.571 [0.527, 0.615] | 0.579 [0.532, 0.626] | 0.492 | 0.489 | 11.6 | 11.9 |
| ANCOM-BC2 | BH | 0.420 | 0.380 | 0.005 [0.000, 0.016] | 0.006 [0.000, 0.019] | 0.003 | 0.000 | 2.6 | 2.3 |
| ALDEx2 | Welch_128 * | 0.411 | 0.378 | 0.000 [0.000, 0.000] | 0.014 [0.000, 0.050] | 0.000 | 0.000 | 2.4 | 2.4 |
| ALDEx2 | Welch_32 | 0.407 | 0.380 | 0.000 [0.000, 0.000] | 0.014 [0.000, 0.050] | 0.000 | 0.000 | 2.4 | 2.4 |
| ALDEx2 | Welch_128_IQLR | 0.406 | 0.366 | 0.000 [0.000, 0.000] | 0.015 [0.000, 0.054] | 0.000 | 0.000 | 2.3 | 2.3 |
| ANCOM-BC2 | BH_StrucZero | 0.404 | 0.367 | 0.005 [0.000, 0.016] | 0.006 [0.000, 0.019] | 0.003 | 0.000 | 2.4 | 2.2 |

| Tool | Configuration | MCC observed | MCC fixed depth | FDR observed | FDR fixed depth | FDR relative truth, observed | FDR relative truth, fixed | Calls per dataset, observed | Calls per dataset, fixed |
|---|---|---|---|---|---|---|---|---|---|
| MaAsLin 2 | TMM_NONE | 0.366 | 0.318 | 0.025 [0.000, 0.055] | 0.038 [0.000, 0.098] | 0.022 | 0.022 | 2.2 | 2.1 |
| ANCOM-BC2 | Holm * | 0.353 | 0.322 | 0.000 [0.000, 0.000] | 0.000 [0.000, 0.000] | 0.000 | 0.000 | 2.1 | 1.9 |
| ANCOM-BC2 | Holm_NoFilter | 0.309 | 0.290 | 0.647 [0.573, 0.722] | 0.628 [0.547, 0.709] | 0.563 | 0.548 | 8.7 | 7.7 |
| ANCOM-BC2 | BH_NoFilter | 0.273 | 0.269 | 0.739 [0.694, 0.785] | 0.727 [0.667, 0.787] | 0.628 | 0.622 | 15.4 | 14.3 |
| DESeq2 | LRT_sparse † | 0.511 | 0.473 | 0.164 [0.104, 0.223] | 0.173 [0.130, 0.217] | 0.134 | 0.131 | 4.7 | 4.5 |
| edgeR | TMM_QL_Legacy † | 0.376 | 0.358 | 0.690 [0.667, 0.713] | 0.704 [0.681, 0.728] | 0.584 | 0.594 | 16.5 | 18.0 |
| edgeR | TMM_QL_Robust † | 0.196 | 0.183 | 0.392 [0.229, 0.556] | 0.462 [0.400, 0.523] | 0.344 | 0.396 | 2.7 | 2.7 |
| ***Signals in one direction (6 scenarios)*** | | | | | | | | | |
| LinDA | Pseudo_0.1 | 0.951 | 0.764 | 0.082 [0.000, 0.184] | 0.332 [0.194, 0.470] | 0.023 | 0.031 | 8.9 | 14.5 |
| MaAsLin 2 | TMM_LOG | 0.910 | 0.818 | 0.147 [0.001, 0.293] | 0.268 [0.241, 0.296] | 0.017 | 0.035 | 9.8 | 12.2 |
| LinDA | Imputation | 0.757 | 0.754 | 0.345 [0.219, 0.471] | 0.344 [0.197, 0.491] | 0.025 | 0.024 | 14.4 | 14.9 |
| MaAsLin 3 | TSS_PLOG_Median | 0.902 | 0.901 | 0.148 [0.080, 0.216] | 0.143 [0.019, 0.268] | 0.005 | 0.005 | 9.8 | 10.2 |
| LinDA | Pseudo_0.5 | 0.952 | 0.745 | 0.078 [0.000, 0.183] | 0.350 [0.231, 0.469] | 0.023 | 0.026 | 8.9 | 15.3 |
| MaAsLin 3 | TSS_PLOG | 0.774 | 0.752 | 0.319 [0.195, 0.442] | 0.341 [0.187, 0.495] | 0.023 | 0.024 | 13.3 | 15.0 |
| MaAsLin 2 | TSS_LOG * | 0.753 | 0.724 | 0.339 [0.213, 0.465] | 0.372 [0.254, 0.491] | 0.021 | 0.025 | 14.6 | 16.0 |
| Wilcoxon | TSS | 0.730 | 0.719 | 0.377 [0.268, 0.486] | 0.379 [0.247, 0.511] | 0.024 | 0.024 | 15.3 | 16.3 |
| LinDA | Adaptive * | 0.759 | 0.752 | 0.343 [0.227, 0.459] | 0.338 [0.195, 0.481] | 0.023 | 0.015 | 14.2 | 15.1 |
| LDM | CLR | 0.963 | 0.760 | 0.063 [0.000, 0.151] | 0.325 [0.169, 0.481] | 0.012 | 0.027 | 8.7 | 15.0 |
| LDM | Omni3 | 0.625 | 0.643 | 0.493 [0.364, 0.621] | 0.467 [0.324, 0.610] | 0.027 | 0.023 | 23.0 | 20.9 |
| LinDA | Adaptive_NoWinsor | 0.785 | 0.766 | 0.313 [0.194, 0.433] | 0.326 [0.199, 0.452] | 0.021 | 0.020 | 13.8 | 14.6 |
| MaAsLin 2 | CLR_NONE | 0.563 | 0.671 | 0.568 [0.507, 0.628] | 0.451 [0.323, 0.580] | 0.070 | 0.042 | 26.5 | 19.2 |
| limma-voom | TMM_voom | 0.544 | 0.722 | 0.557 [0.440, 0.675] | 0.366 [0.190, 0.543] | 0.056 | 0.039 | 31.2 | 18.9 |
| LDM | PA | 0.698 | 0.720 | 0.348 [0.237, 0.459] | 0.319 [0.209, 0.430] | 0.027 | 0.027 | 15.1 | 13.0 |
| LDM | Arcsin | 0.644 | 0.635 | 0.474 [0.344, 0.604] | 0.475 [0.345, 0.604] | 0.031 | 0.026 | 20.6 | 21.1 |
| ZicoSeq | MultiLink | 0.961 | 0.903 | 0.062 [0.004, 0.120] | 0.113 [0.000, 0.270] | 0.008 | 0.009 | 8.6 | 10.0 |
| LinDA | Pseudo_1 | 0.957 | 0.735 | 0.072 [0.000, 0.169] | 0.362 [0.250, 0.475] | 0.019 | 0.026 | 8.8 | 15.6 |
| ZicoSeq | NoPosterior | 0.943 | 0.902 | 0.046 [0.004, 0.089] | 0.105 [0.000, 0.259] | 0.005 | 0.008 | 8.1 | 9.3 |
| MaAsLin 3 | TSS_LOG_Median * | 0.974 | 0.936 | 0.029 [0.000, 0.103] | 0.085 [0.000, 0.226] | 0.004 | 0.004 | 8.2 | 8.7 |
| LDM | Omni * | 0.617 | 0.632 | 0.503 [0.415, 0.591] | 0.479 [0.367, 0.590] | 0.031 | 0.021 | 22.4 | 20.8 |
| Wilcoxon | CLR | 0.479 | 0.628 | 0.609 [0.537, 0.682] | 0.483 [0.384, 0.582] | 0.078 | 0.053 | 33.5 | 20.5 |
| limma-voom | TMMwsp_voom | 0.430 | 0.677 | 0.654 [0.568, 0.740] | 0.425 [0.150, 0.699] | 0.091 | 0.039 | 39.5 | 21.5 |
| MaAsLin 3 | CLR_NONE | 0.846 | 0.766 | 0.194 [0.080, 0.308] | 0.313 [0.229, 0.396] | 0.017 | 0.017 | 10.7 | 12.8 |
| MaAsLin 3 | CLR_NONE_Median | 0.955 | 0.940 | 0.029 [0.000, 0.103] | 0.052 [0.000, 0.113] | 0.004 | 0.004 | 7.9 | 8.1 |
| MaAsLin 3 | TSS_LOG | 0.639 | 0.731 | 0.453 [0.368, 0.537] | 0.358 [0.278, 0.439] | 0.015 | 0.010 | 21.8 | 15.0 |
| DESeq2 | LRT_poscounts | 0.679 | 0.651 | 0.470 [0.375, 0.564] | 0.503 [0.464, 0.542] | 0.058 | 0.053 | 16.5 | 17.7 |
| DESeq2 | Wald_poscounts * | 0.673 | 0.653 | 0.478 [0.382, 0.573] | 0.501 [0.461, 0.541] | 0.056 | 0.054 | 16.7 | 17.5 |
| ZicoSeq | Default * | 0.955 | 0.872 | 0.059 [0.000, 0.132] | 0.122 [0.000, 0.307] | 0.005 | 0.009 | 8.4 | 10.5 |
| Wilcoxon | Rarefied | 0.734 | 0.721 | 0.367 [0.295, 0.439] | 0.367 [0.258, 0.476] | 0.013 | 0.012 | 15.5 | 15.5 |
| edgeR | TMM_LRT | 0.691 | 0.674 | 0.458 [0.383, 0.533] | 0.465 [0.398, 0.533] | 0.068 | 0.051 | 15.5 | 16.8 |
| LDM | Freq | 0.640 | 0.635 | 0.449 [0.334, 0.565] | 0.456 [0.349, 0.563] | 0.015 | 0.017 | 18.3 | 17.7 |
| ALDEx2 | Wilcox_128 | 0.904 | 0.754 | 0.099 [0.004, 0.195] | 0.305 [0.186, 0.424] | 0.017 | 0.020 | 8.5 | 12.6 |
| edgeR | TMM_QL * | 0.571 | 0.533 | 0.603 [0.563, 0.642] | 0.634 [0.613, 0.655] | 0.100 | 0.108 | 20.9 | 23.8 |
| ANCOM-BC2 | BH | 0.973 | 0.912 | 0.005 [0.000, 0.022] | 0.075 [0.019, 0.131] | 0.000 | 0.005 | 7.8 | 8.1 |
| ALDEx2 | Welch_128 * | 0.897 | 0.771 | 0.064 [0.000, 0.154] | 0.237 [0.124, 0.351] | 0.010 | 0.015 | 7.6 | 10.1 |
| ALDEx2 | Welch_32 | 0.906 | 0.765 | 0.053 [0.000, 0.143] | 0.246 [0.121, 0.371] | 0.014 | 0.015 | 7.7 | 10.3 |
| ALDEx2 | Welch_128_IQLR | 0.934 | 0.870 | 0.022 [0.000, 0.076] | 0.113 [0.002, 0.225] | 0.000 | 0.000 | 7.5 | 8.4 |
| ANCOM-BC2 | BH_StrucZero | 0.967 | 0.889 | 0.005 [0.000, 0.022] | 0.078 [0.012, 0.145] | 0.000 | 0.005 | 7.7 | 7.8 |
| MaAsLin 2 | TMM_NONE | 0.813 | 0.732 | 0.109 [0.000, 0.231] | 0.215 [0.142, 0.288] | 0.010 | 0.000 | 7.3 | 8.8 |
| ANCOM-BC2 | Holm * | 0.957 | 0.882 | 0.000 [0.000, 0.000] | 0.034 [0.000, 0.118] | 0.000 | 0.000 | 7.5 | 6.8 |
| ANCOM-BC2 | Holm_NoFilter | 0.677 | 0.697 | 0.476 [0.416, 0.535] | 0.436 [0.400, 0.472] | 0.088 | 0.072 | 16.1 | 14.5 |

| Tool | Configuration | MCC observed | MCC fixed depth | FDR observed | FDR fixed depth | FDR relative truth, observed | FDR relative truth, fixed | Calls per dataset, observed | Calls per dataset, fixed |
|---|---|---|---|---|---|---|---|---|---|
| ANCOM-BC2 | BH_NoFilter | 0.525 | 0.542 | 0.648 [0.615, 0.680] | 0.627 [0.576, 0.678] | 0.112 | 0.087 | 23.5 | 22.4 |
| DESeq2 | LRT_sparse † | 0.635 | 0.587 | 0.476 [0.395, 0.558] | 0.531 [0.495, 0.567] | 0.077 | 0.066 | 15.0 | 16.6 |
| edgeR | TMM_QL_Legacy † | 0.501 | 0.447 | 0.671 [0.645, 0.698] | 0.711 [0.699, 0.722] | 0.124 | 0.119 | 25.4 | 29.7 |
| edgeR | TMM_QL_Robust † | 0.429 | 0.342 | 0.509 [0.334, 0.685] | 0.613 [0.558, 0.668] | 0.033 | 0.070 | 10.5 | 10.8 |
| ***Single extreme feature (2 scenarios)*** | | | | | | | | | |
| LinDA | Pseudo_0.1 | 0.872 | 0.098 | 0.208 [0.000, 0.438] | 0.978 [0.972, 0.983] | 0.104 | 0.000 | 1.5 | 48.8 |
| MaAsLin 2 | TMM_LOG | 1.000 | 0.215 | 0.000 [0.000, 0.000] | 0.902 [0.729, 1.075] | 0.000 | 0.003 | 1.0 | 35.0 |
| LinDA | Imputation | 0.124 | 0.082 | 0.972 [0.966, 0.979] | 0.981 [0.980, 0.983] | 0.000 | 0.000 | 38.9 | 54.6 |
| MaAsLin 3 | TSS_PLOG_Median | 0.189 | 0.146 | 0.951 [0.941, 0.961] | 0.966 [0.959, 0.973] | 0.000 | 0.000 | 22.8 | 32.2 |
| LinDA | Pseudo_0.5 | 0.909 | 0.092 | 0.146 [0.000, 0.419] | 0.979 [0.973, 0.985] | 0.062 | 0.000 | 1.4 | 51.4 |
| MaAsLin 3 | TSS_PLOG | 0.148 | 0.082 | 0.965 [0.955, 0.975] | 0.981 [0.977, 0.984] | 0.003 | 0.002 | 31.9 | 55.4 |
| MaAsLin 2 | TSS_LOG * | 0.115 | 0.080 | 0.974 [0.968, 0.980] | 0.981 [0.978, 0.984] | 0.003 | 0.002 | 42.0 | 56.4 |
| Wilcoxon | TSS | 0.160 | 0.087 | 0.963 [0.961, 0.964] | 0.980 [0.976, 0.984] | 0.010 | 0.005 | 27.4 | 53.4 |
| LinDA | Adaptive * | 0.124 | 0.092 | 0.972 [0.966, 0.979] | 0.979 [0.973, 0.984] | 0.000 | 0.000 | 38.9 | 51.1 |
| LDM | CLR | 1.000 | 0.076 | 0.000 [0.000, 0.000] | 0.982 [0.980, 0.984] | 0.000 | 0.000 | 1.0 | 57.5 |
| LDM | Omni3 | 0.030 | 0.042 | 0.986 [0.985, 0.988] | 0.985 [0.983, 0.987] | 0.004 | 0.004 | 75.5 | 70.8 |
| LinDA | Adaptive_NoWinsor | 0.152 | 0.107 | 0.965 [0.960, 0.969] | 0.976 [0.968, 0.983] | 0.000 | 0.000 | 30.0 | 45.5 |
| MaAsLin 2 | CLR_NONE | 0.666 | 0.057 | 0.428 [0.000, 1.021] | 0.985 [0.983, 0.986] | 0.057 | 0.004 | 6.8 | 66.9 |
| limma-voom | TMM_voom | 1.000 | 0.049 | 0.000 [0.000, 0.000] | 0.986 [0.985, 0.986] | 0.000 | 0.003 | 1.0 | 70.5 |
| LDM | PA | 0.037 | 0.042 | 0.986 [0.984, 0.988] | 0.985 [0.982, 0.988] | 0.004 | 0.004 | 72.9 | 70.2 |
| LDM | Arcsin | 0.038 | 0.033 | 0.985 [0.982, 0.988] | 0.986 [0.984, 0.988] | 0.004 | 0.004 | 72.0 | 74.2 |
| ZicoSeq | MultiLink | 0.445 | 0.481 | 0.674 [0.278, 1.070] | 0.593 [0.245, 0.941] | 0.006 | 0.000 | 14.8 | 22.0 |
| LinDA | Pseudo_1 | 0.909 | 0.093 | 0.146 [0.000, 0.419] | 0.979 [0.973, 0.985] | 0.062 | 0.000 | 1.4 | 51.0 |
| ZicoSeq | NoPosterior | 0.802 | 0.261 | 0.308 [0.000, 0.735] | 0.838 [0.467, 1.210] | 0.000 | 0.000 | 2.0 | 31.1 |
| MaAsLin 3 | TSS_LOG_Median * | 1.000 | 0.288 | 0.000 [0.000, 0.000] | 0.882 [0.794, 0.970] | 0.000 | 0.000 | 1.0 | 17.2 |
| LDM | Omni * | 0.029 | 0.037 | 0.987 [0.985, 0.988] | 0.986 [0.985, 0.987] | 0.004 | 0.004 | 76.0 | 72.6 |
| Wilcoxon | CLR | 0.663 | 0.041 | 0.417 [0.000, 0.895] | 0.986 [0.986, 0.987] | 0.008 | 0.003 | 9.2 | 74.0 |
| limma-voom | TMMwsp_voom | 0.892 | 0.049 | 0.163 [0.000, 0.477] | 0.986 [0.985, 0.986] | 0.025 | 0.004 | 1.6 | 70.6 |
| MaAsLin 3 | CLR_NONE | 0.808 | 0.135 | 0.302 [0.000, 0.712] | 0.969 [0.959, 0.978] | 0.000 | 0.000 | 1.9 | 35.4 |
| MaAsLin 3 | CLR_NONE_Median | 1.000 | 0.287 | 0.000 [0.000, 0.000] | 0.825 [0.466, 1.185] | 0.000 | 0.000 | 1.0 | 26.8 |
| MaAsLin 3 | TSS_LOG | 0.027 | 0.101 | 0.987 [0.986, 0.988] | 0.978 [0.976, 0.980] | 0.000 | 0.000 | 77.8 | 46.5 |
| DESeq2 | LRT_poscounts | 0.872 | 0.253 | 0.208 [0.000, 0.438] | 0.895 [0.721, 1.070] | 0.000 | 0.000 | 1.5 | 19.6 |
| DESeq2 | Wald_poscounts * | 0.835 | 0.252 | 0.271 [0.000, 0.652] | 0.896 [0.721, 1.071] | 0.000 | 0.000 | 1.6 | 19.6 |
| ZicoSeq | Default * | 0.547 | 0.409 | 0.566 [0.205, 0.927] | 0.720 [0.417, 1.022] | 0.000 | 0.000 | 9.9 | 20.0 |
| Wilcoxon | Rarefied | 0.097 | 0.105 | 0.978 [0.976, 0.981] | 0.977 [0.974, 0.980] | 0.002 | 0.003 | 48.6 | 45.5 |
| edgeR | TMM_LRT | 0.583 | 0.337 | 0.613 [0.345, 0.882] | 0.778 [0.414, 1.142] | 0.056 | 0.004 | 3.8 | 25.8 |
| LDM | Freq | 0.063 | 0.072 | 0.983 [0.981, 0.986] | 0.982 [0.980, 0.985] | 0.004 | 0.002 | 63.9 | 59.6 |
| ALDEx2 | Wilcox_128 | 1.000 | 0.158 | 0.000 [0.000, 0.000] | 0.960 [0.956, 0.964] | 0.000 | 0.000 | 1.0 | 32.0 |
| edgeR | TMM_QL * | 0.377 | 0.223 | 0.832 [0.668, 0.995] | 0.912 [0.796, 1.027] | 0.013 | 0.004 | 7.8 | 31.0 |
| ANCOM-BC2 | BH | 1.000 | 0.351 | 0.000 [0.000, 0.000] | 0.849 [0.760, 0.938] | 0.000 | 0.000 | 1.0 | 9.9 |
| ALDEx2 | Welch_128 * | 1.000 | 0.195 | 0.000 [0.000, 0.000] | 0.947 [0.938, 0.956] | 0.000 | 0.000 | 1.0 | 23.8 |
| ALDEx2 | Welch_32 | 1.000 | 0.196 | 0.000 [0.000, 0.000] | 0.947 [0.940, 0.954] | 0.000 | 0.000 | 1.0 | 23.4 |
| ALDEx2 | Welch_128_IQLR | 1.000 | 0.220 | 0.000 [0.000, 0.000] | 0.939 [0.929, 0.949] | 0.000 | 0.000 | 1.0 | 17.6 |
| ANCOM-BC2 | BH_StrucZero | 1.000 | 0.366 | 0.000 [0.000, 0.000] | 0.839 [0.741, 0.938] | 0.000 | 0.000 | 1.0 | 9.0 |
| MaAsLin 2 | TMM_NONE | 0.750 | 0.417 | 0.000 [0.000, 0.000] | 0.681 [0.287, 1.075] | 0.000 | 0.000 | 0.8 | 13.1 |
| ANCOM-BC2 | Holm * | 1.000 | 0.481 | 0.000 [0.000, 0.000] | 0.705 [0.406, 1.004] | 0.000 | 0.000 | 1.0 | 6.5 |
| ANCOM-BC2 | Holm_NoFilter | 0.288 | 0.232 | 0.904 [0.877, 0.930] | 0.933 [0.899, 0.966] | 0.000 | 0.000 | 11.4 | 16.2 |
| ANCOM-BC2 | BH_NoFilter | 0.201 | 0.176 | 0.947 [0.927, 0.966] | 0.957 [0.942, 0.972] | 0.000 | 0.000 | 20.5 | 24.4 |
| DESeq2 | LRT_sparse † | 0.851 | 0.239 | 0.250 [0.000, 0.575] | 0.900 [0.722, 1.078] | 0.000 | 0.000 | 1.5 | 23.1 |
| edgeR | TMM_QL_Legacy † | 0.311 | 0.165 | 0.882 [0.771, 0.993] | 0.950 [0.908, 0.991] | 0.019 | 0.003 | 10.9 | 36.8 |

| Tool | Configuration | MCC observed | MCC fixed depth | FDR observed | FDR fixed depth | FDR relative truth, observed | FDR relative truth, fixed | Calls per dataset, observed | Calls per dataset, fixed |
|---|---|---|---|---|---|---|---|---|---|
| edgeR | TMM_QL_Robust † | 0.595 | 0.395 | 0.622 [0.485, 0.758] | 0.809 [0.655, 0.962] | 0.000 | 0.000 | 3.4 | 8.4 |
| ***Null design (3 scenarios)*** | | | | | | | | | |
| LinDA | Pseudo_0.1 | | | 0.08 [0.00, 0.38] | 0.08 [0.00, 0.38] | | | 0.1 | 0.1 |
| MaAsLin 2 | TMM_LOG | | | 0.08 [0.00, 0.38] | 0.17 [0.02, 0.48] | | | 0.1 | 0.2 |
| LinDA | Imputation | | | 0.08 [0.00, 0.38] | 0.08 [0.00, 0.38] | | | 0.1 | 0.1 |
| MaAsLin 3 | TSS_PLOG_Median | | | 0.08 [0.00, 0.38] | 0.08 [0.00, 0.38] | | | 0.1 | 0.1 |
| LinDA | Pseudo_0.5 | | | 0.08 [0.00, 0.38] | 0.08 [0.00, 0.38] | | | 0.1 | 0.1 |
| MaAsLin 3 | TSS_PLOG | | | 0.08 [0.00, 0.38] | 0.08 [0.00, 0.38] | | | 0.1 | 0.1 |
| MaAsLin 2 | TSS_LOG * | | | 0.17 [0.02, 0.48] | 0.17 [0.02, 0.48] | | | 0.2 | 0.2 |
| Wilcoxon | TSS | | | 0.17 [0.02, 0.48] | 0.17 [0.02, 0.48] | | | 0.2 | 0.2 |
| LinDA | Adaptive * | | | 0.08 [0.00, 0.38] | 0.08 [0.00, 0.38] | | | 0.1 | 0.1 |
| LDM | CLR | | | 0.00 [0.00, 0.26] | 0.00 [0.00, 0.26] | | | 0.0 | 0.0 |
| LDM | Omni3 | | | 0.17 [0.02, 0.48] | 0.17 [0.02, 0.48] | | | 0.2 | 0.2 |
| LinDA | Adaptive_NoWinsor | | | 0.08 [0.00, 0.38] | 0.08 [0.00, 0.38] | | | 0.1 | 0.1 |
| MaAsLin 2 | CLR_NONE | | | 0.08 [0.00, 0.38] | 0.08 [0.00, 0.38] | | | 0.1 | 0.1 |
| limma-voom | TMM_voom | | | 0.08 [0.00, 0.38] | 0.08 [0.00, 0.38] | | | 0.4 | 0.1 |
| LDM | PA | | | 0.08 [0.00, 0.38] | 0.08 [0.00, 0.38] | | | 0.1 | 0.1 |
| LDM | Arcsin | | | 0.17 [0.02, 0.48] | 0.17 [0.02, 0.48] | | | 0.2 | 0.2 |
| ZicoSeq | MultiLink | | | 0.00 [0.00, 0.26] | 0.08 [0.00, 0.38] | | | 0.0 | 0.1 |
| LinDA | Pseudo_1 | | | 0.08 [0.00, 0.38] | 0.08 [0.00, 0.38] | | | 0.1 | 0.1 |
| ZicoSeq | NoPosterior | | | 0.00 [0.00, 0.26] | 0.00 [0.00, 0.26] | | | 0.0 | 0.0 |
| MaAsLin 3 | TSS_LOG_Median * | | | 0.08 [0.00, 0.38] | 0.00 [0.00, 0.26] | | | 0.1 | 0.0 |
| LDM | Omni * | | | 0.17 [0.02, 0.48] | 0.17 [0.02, 0.48] | | | 0.2 | 0.2 |
| Wilcoxon | CLR | | | 0.08 [0.00, 0.38] | 0.08 [0.00, 0.38] | | | 0.1 | 0.1 |
| limma-voom | TMMwsp_voom | | | 0.00 [0.00, 0.26] | 0.00 [0.00, 0.26] | | | 0.0 | 0.0 |
| MaAsLin 3 | CLR_NONE | | | 0.00 [0.00, 0.26] | 0.00 [0.00, 0.26] | | | 0.0 | 0.0 |
| MaAsLin 3 | CLR_NONE_Median | | | 0.00 [0.00, 0.26] | 0.00 [0.00, 0.26] | | | 0.0 | 0.0 |
| MaAsLin 3 | TSS_LOG | | | 0.08 [0.00, 0.38] | 0.00 [0.00, 0.26] | | | 0.1 | 0.0 |
| DESeq2 | LRT_poscounts | | | 0.42 [0.15, 0.72] | 0.17 [0.02, 0.48] | | | 0.4 | 0.2 |
| DESeq2 | Wald_poscounts * | | | 0.42 [0.15, 0.72] | 0.33 [0.10, 0.65] | | | 0.6 | 0.3 |
| ZicoSeq | Default * | | | 0.00 [0.00, 0.26] | 0.00 [0.00, 0.26] | | | 0.0 | 0.0 |
| Wilcoxon | Rarefied | | | 0.08 [0.00, 0.38] | 0.08 [0.00, 0.38] | | | 0.1 | 0.1 |
| edgeR | TMM_LRT | | | 0.75 [0.43, 0.95] | 0.83 [0.52, 0.98] | | | 2.2 | 1.9 |
| LDM | Freq | | | 0.17 [0.02, 0.48] | 0.17 [0.02, 0.48] | | | 0.2 | 0.2 |
| ALDEx2 | Wilcox_128 | | | 0.08 [0.00, 0.38] | 0.08 [0.00, 0.38] | | | 0.1 | 0.1 |
| edgeR | TMM_QL * | | | 1.00 [0.74, 1.00] | 1.00 [0.74, 1.00] | | | 7.2 | 7.0 |
| ANCOM-BC2 | BH | | | 0.08 [0.00, 0.38] | 0.00 [0.00, 0.26] | | | 0.1 | 0.0 |
| ALDEx2 | Welch_128 * | | | 0.00 [0.00, 0.26] | 0.00 [0.00, 0.26] | | | 0.0 | 0.0 |
| ALDEx2 | Welch_32 | | | 0.00 [0.00, 0.26] | 0.00 [0.00, 0.26] | | | 0.0 | 0.0 |
| ALDEx2 | Welch_128_IQLR | | | 0.00 [0.00, 0.26] | 0.00 [0.00, 0.26] | | | 0.0 | 0.0 |
| ANCOM-BC2 | BH_StrucZero | | | 0.08 [0.00, 0.38] | 0.00 [0.00, 0.26] | | | 0.1 | 0.0 |
| MaAsLin 2 | TMM_NONE | | | 0.08 [0.00, 0.38] | 0.08 [0.00, 0.38] | | | 0.1 | 0.1 |
| ANCOM-BC2 | Holm * | | | 0.08 [0.00, 0.38] | 0.00 [0.00, 0.26] | | | 0.1 | 0.0 |
| ANCOM-BC2 | Holm_NoFilter | | | 1.00 [0.74, 1.00] | 1.00 [0.74, 1.00] | | | 5.8 | 6.1 |
| ANCOM-BC2 | BH_NoFilter | | | 1.00 [0.74, 1.00] | 1.00 [0.74, 1.00] | | | 15.3 | 14.6 |
| DESeq2 | LRT_sparse † | | | 0.33 [0.10, 0.65] | 0.33 [0.10, 0.65] | | | 0.4 | 0.7 |
| edgeR | TMM_QL_Legacy † | | | 1.00 [0.74, 1.00] | 1.00 [0.74, 1.00] | | | 12.0 | 12.3 |
| edgeR | TMM_QL_Robust † | | | 0.67 [0.35, 0.90] | 0.67 [0.35, 0.90] | | | 0.9 | 1.2 |

**Table S6. Null designs.** No signal was implanted, and every call is false. With Benjamini–Hochberg adjustment at 0.05, at most 5% of the datasets should contain a call. Stool at 50, 100 and 200 samples, the IBD community at 50 samples and simulated data at 50 to 500 samples, four replicates each. False positives per dataset with 95% confidence intervals across replicates, and datasets with a call with exact binomial 95% intervals.

| Tool | Configuration | False positives per dataset | Datasets with at least one call |
|---|---|---|---|
| ***Semi-synthetic, stool, genus level*** | | | |
| LinDA | Pseudo_0.1 | 0.08 [0.00, 0.35] | 1 of 12 [0.00, 0.38] |
| MaAsLin 2 | TMM_LOG | 0.08 [0.00, 0.35] | 1 of 12 [0.00, 0.38] |
| LinDA | Imputation | 0.08 [0.00, 0.35] | 1 of 12 [0.00, 0.38] |
| MaAsLin 3 | TSS_PLOG_Median | 0.08 [0.00, 0.35] | 1 of 12 [0.00, 0.38] |
| LinDA | Pseudo_0.5 | 0.08 [0.00, 0.35] | 1 of 12 [0.00, 0.38] |
| MaAsLin 3 | TSS_PLOG | 0.08 [0.00, 0.35] | 1 of 12 [0.00, 0.38] |
| MaAsLin 2 | TSS_LOG * | 0.17 [0.00, 0.47] | 2 of 12 [0.02, 0.48] |
| Wilcoxon | TSS | 0.17 [0.00, 0.47] | 2 of 12 [0.02, 0.48] |
| LinDA | Adaptive * | 0.08 [0.00, 0.35] | 1 of 12 [0.00, 0.38] |
| LDM | CLR | 0.00 [0.00, 0.00] | 0 of 12 [0.00, 0.26] |
| LDM | Omni3 | 0.17 [0.00, 0.47] | 2 of 12 [0.02, 0.48] |
| LinDA | Adaptive_NoWinsor | 0.08 [0.00, 0.35] | 1 of 12 [0.00, 0.38] |
| MaAsLin 2 | CLR_NONE | 0.08 [0.00, 0.35] | 1 of 12 [0.00, 0.38] |
| limma-voom | TMM_voom | 0.42 [0.00, 1.74] | 1 of 12 [0.00, 0.38] |
| LDM | PA | 0.08 [0.00, 0.35] | 1 of 12 [0.00, 0.38] |
| LDM | Arcsin | 0.17 [0.00, 0.47] | 2 of 12 [0.02, 0.48] |
| ZicoSeq | MultiLink | 0.00 [0.00, 0.00] | 0 of 12 [0.00, 0.26] |
| LinDA | Pseudo_1 | 0.08 [0.00, 0.35] | 1 of 12 [0.00, 0.38] |
| ZicoSeq | NoPosterior | 0.00 [0.00, 0.00] | 0 of 12 [0.00, 0.26] |
| MaAsLin 3 | TSS_LOG_Median * | 0.08 [0.00, 0.35] | 1 of 12 [0.00, 0.38] |
| LDM | Omni * | 0.17 [0.00, 0.47] | 2 of 12 [0.02, 0.48] |
| Wilcoxon | CLR | 0.08 [0.00, 0.35] | 1 of 12 [0.00, 0.38] |
| limma-voom | TMMwsp_voom | 0.00 [0.00, 0.00] | 0 of 12 [0.00, 0.26] |
| MaAsLin 3 | CLR_NONE | 0.00 [0.00, 0.00] | 0 of 12 [0.00, 0.26] |
| MaAsLin 3 | CLR_NONE_Median | 0.00 [0.00, 0.00] | 0 of 12 [0.00, 0.26] |
| MaAsLin 3 | TSS_LOG | 0.08 [0.00, 0.35] | 1 of 12 [0.00, 0.38] |
| DESeq2 | LRT_poscounts | 0.42 [0.15, 0.68] | 5 of 12 [0.15, 0.72] |
| DESeq2 | Wald_poscounts * | 0.58 [0.32, 0.85] | 5 of 12 [0.15, 0.72] |
| ZicoSeq | Default * | 0.00 [0.00, 0.00] | 0 of 12 [0.00, 0.26] |
| Wilcoxon | Rarefied | 0.08 [0.00, 0.35] | 1 of 12 [0.00, 0.38] |
| edgeR | TMM_LRT | 2.25 [0.92, 3.58] | 9 of 12 [0.43, 0.95] |
| LDM | Freq | 0.17 [0.00, 0.47] | 2 of 12 [0.02, 0.48] |
| ALDEx2 | Wilcox_128 | 0.08 [0.00, 0.35] | 1 of 12 [0.00, 0.38] |
| edgeR | TMM_QL * | 7.25 [4.82, 9.68] | 12 of 12 [0.74, 1.00] |
| ANCOM-BC2 | BH | 0.08 [0.00, 0.35] | 1 of 12 [0.00, 0.38] |
| ALDEx2 | Welch_128 * | 0.00 [0.00, 0.00] | 0 of 12 [0.00, 0.26] |
| ALDEx2 | Welch_32 | 0.00 [0.00, 0.00] | 0 of 12 [0.00, 0.26] |
| ALDEx2 | Welch_128_IQLR | 0.00 [0.00, 0.00] | 0 of 12 [0.00, 0.26] |
| ANCOM-BC2 | BH_StrucZero | 0.08 [0.00, 0.35] | 1 of 12 [0.00, 0.38] |
| MaAsLin 2 | TMM_NONE | 0.08 [0.00, 0.35] | 1 of 12 [0.00, 0.38] |
| ANCOM-BC2 | Holm * | 0.08 [0.00, 0.35] | 1 of 12 [0.00, 0.38] |
| ANCOM-BC2 | Holm_NoFilter | 5.75 [4.57, 6.93] | 12 of 12 [0.74, 1.00] |
| ANCOM-BC2 | BH_NoFilter | 15.33 [13.45, 17.22] | 12 of 12 [0.74, 1.00] |
| ***Semi-synthetic, stool, OTU level*** | | | |
| LinDA | Pseudo_0.1 | 0.08 [0.00, 0.35] | 1 of 12 [0.00, 0.38] |

| Tool | Configuration | False positives per dataset | Datasets with at least one call |
|---|---|---|---|
| MaAsLin 2 | TMM_LOG | 0.00 [0.00, 0.00] | 0 of 12 [0.00, 0.26] |
| LinDA | Imputation | 0.00 [0.00, 0.00] | 0 of 12 [0.00, 0.26] |
| MaAsLin 3 | TSS_PLOG_Median | 0.00 [0.00, 0.00] | 0 of 12 [0.00, 0.26] |
| LinDA | Pseudo_0.5 | 0.08 [0.00, 0.35] | 1 of 12 [0.00, 0.38] |
| MaAsLin 3 | TSS_PLOG | 0.00 [0.00, 0.00] | 0 of 12 [0.00, 0.26] |
| MaAsLin 2 | TSS_LOG * | 0.00 [0.00, 0.00] | 0 of 12 [0.00, 0.26] |
| Wilcoxon | TSS | 0.00 [0.00, 0.00] | 0 of 12 [0.00, 0.26] |
| LinDA | Adaptive * | 0.00 [0.00, 0.00] | 0 of 12 [0.00, 0.26] |
| LDM | CLR | 0.00 [0.00, 0.00] | 0 of 12 [0.00, 0.26] |
| LDM | Omni3 | 0.00 [0.00, 0.00] | 0 of 12 [0.00, 0.26] |
| LinDA | Adaptive_NoWinsor | 0.00 [0.00, 0.00] | 0 of 12 [0.00, 0.26] |
| MaAsLin 2 | CLR_NONE | 0.00 [0.00, 0.00] | 0 of 12 [0.00, 0.26] |
| limma-voom | TMM_voom | 5.50 [0.00, 20.23] | 5 of 12 [0.15, 0.72] |
| LDM | PA | 0.00 [0.00, 0.00] | 0 of 12 [0.00, 0.26] |
| LDM | Arcsin | 0.00 [0.00, 0.00] | 0 of 12 [0.00, 0.26] |
| ZicoSeq | MultiLink | 0.00 [0.00, 0.00] | 0 of 12 [0.00, 0.26] |
| LinDA | Pseudo_1 | 0.08 [0.00, 0.35] | 1 of 12 [0.00, 0.38] |
| ZicoSeq | NoPosterior | 0.17 [0.00, 0.70] | 1 of 12 [0.00, 0.38] |
| MaAsLin 3 | TSS_LOG_Median * | 0.08 [0.00, 0.35] | 1 of 12 [0.00, 0.38] |
| LDM | Omni * | 0.00 [0.00, 0.00] | 0 of 12 [0.00, 0.26] |
| Wilcoxon | CLR | 0.00 [0.00, 0.00] | 0 of 12 [0.00, 0.26] |
| limma-voom | TMMwsp_voom | 0.58 [0.00, 2.44] | 2 of 12 [0.02, 0.48] |
| MaAsLin 3 | CLR_NONE | 0.17 [0.00, 0.70] | 1 of 12 [0.00, 0.38] |
| MaAsLin 3 | CLR_NONE_Median | 0.17 [0.00, 0.70] | 1 of 12 [0.00, 0.38] |
| MaAsLin 3 | TSS_LOG | 0.00 [0.00, 0.00] | 0 of 12 [0.00, 0.26] |
| DESeq2 | LRT_poscounts | 0.25 [0.00, 0.76] | 3 of 12 [0.05, 0.57] |
| DESeq2 | Wald_poscounts * | 0.58 [0.00, 1.84] | 4 of 12 [0.10, 0.65] |
| ZicoSeq | Default * | 0.00 [0.00, 0.00] | 0 of 12 [0.00, 0.26] |
| Wilcoxon | Rarefied | 0.00 [0.00, 0.00] | 0 of 12 [0.00, 0.26] |
| edgeR | TMM_LRT | 22.08 [7.01, 37.16] | 12 of 12 [0.74, 1.00] |
| LDM | Freq | 0.00 [0.00, 0.00] | 0 of 12 [0.00, 0.26] |
| ALDEx2 | Wilcox_128 | 0.00 [0.00, 0.00] | 0 of 12 [0.00, 0.26] |
| edgeR | TMM_QL * | 53.92 [19.40, 88.44] | 12 of 12 [0.74, 1.00] |
| ANCOM-BC2 | BH | 0.00 [0.00, 0.00] | 0 of 12 [0.00, 0.26] |
| ALDEx2 | Welch_128 * | 0.00 [0.00, 0.00] | 0 of 12 [0.00, 0.26] |
| ALDEx2 | Welch_32 | 0.00 [0.00, 0.00] | 0 of 12 [0.00, 0.26] |
| ALDEx2 | Welch_128_IQLR | 0.00 [0.00, 0.00] | 0 of 12 [0.00, 0.26] |
| ANCOM-BC2 | BH_StrucZero | 0.00 [0.00, 0.00] | 0 of 12 [0.00, 0.26] |
| MaAsLin 2 | TMM_NONE | 0.00 [0.00, 0.00] | 0 of 12 [0.00, 0.26] |
| ANCOM-BC2 | Holm * | 0.00 [0.00, 0.00] | 0 of 12 [0.00, 0.26] |
| ANCOM-BC2 | Holm_NoFilter | 139.25 [123.99, 154.51] | 12 of 12 [0.74, 1.00] |
| ANCOM-BC2 | BH_NoFilter | 858.08 [802.81, 913.35] | 12 of 12 [0.74, 1.00] |
| ***Semi-synthetic, IBD, genus level*** | | | |
| LinDA | Pseudo_0.1 | 0.00 [0.00, 0.00] | 0 of 4 [0.00, 0.60] |
| MaAsLin 2 | TMM_LOG | 0.00 [0.00, 0.00] | 0 of 4 [0.00, 0.60] |
| LinDA | Imputation | 0.00 [0.00, 0.00] | 0 of 4 [0.00, 0.60] |
| MaAsLin 3 | TSS_PLOG_Median | 0.00 [0.00, 0.00] | 0 of 4 [0.00, 0.60] |
| LinDA | Pseudo_0.5 | 0.00 [0.00, 0.00] | 0 of 4 [0.00, 0.60] |
| MaAsLin 3 | TSS_PLOG | 0.00 [0.00, 0.00] | 0 of 4 [0.00, 0.60] |
| MaAsLin 2 | TSS_LOG * | 0.00 [0.00, 0.00] | 0 of 4 [0.00, 0.60] |
| Wilcoxon | TSS | 0.00 [0.00, 0.00] | 0 of 4 [0.00, 0.60] |

| Tool | Configuration | False positives per dataset | Datasets with at least one call |
|---|---|---|---|
| LinDA | Adaptive * | 0.00 [0.00, 0.00] | 0 of 4 [0.00, 0.60] |
| LDM | CLR | 0.00 [0.00, 0.00] | 0 of 4 [0.00, 0.60] |
| LDM | Omni3 | 0.00 [0.00, 0.00] | 0 of 4 [0.00, 0.60] |
| LinDA | Adaptive_NoWinsor | 0.00 [0.00, 0.00] | 0 of 4 [0.00, 0.60] |
| MaAsLin 2 | CLR_NONE | 0.00 [0.00, 0.00] | 0 of 4 [0.00, 0.60] |
| limma-voom | TMM_voom | 0.00 [0.00, 0.00] | 0 of 4 [0.00, 0.60] |
| LDM | PA | 0.00 [0.00, 0.00] | 0 of 4 [0.00, 0.60] |
| LDM | Arcsin | 0.00 [0.00, 0.00] | 0 of 4 [0.00, 0.60] |
| ZicoSeq | MultiLink | 0.00 [0.00, 0.00] | 0 of 4 [0.00, 0.60] |
| LinDA | Pseudo_1 | 0.00 [0.00, 0.00] | 0 of 4 [0.00, 0.60] |
| ZicoSeq | NoPosterior | 0.00 [0.00, 0.00] | 0 of 4 [0.00, 0.60] |
| MaAsLin 3 | TSS_LOG_Median * | 0.00 [0.00, 0.00] | 0 of 4 [0.00, 0.60] |
| LDM | Omni * | 0.00 [0.00, 0.00] | 0 of 4 [0.00, 0.60] |
| Wilcoxon | CLR | 0.25 [0.00, 1.05] | 1 of 4 [0.01, 0.81] |
| limma-voom | TMMwsp_voom | 0.00 [0.00, 0.00] | 0 of 4 [0.00, 0.60] |
| MaAsLin 3 | CLR_NONE | 0.00 [0.00, 0.00] | 0 of 4 [0.00, 0.60] |
| MaAsLin 3 | CLR_NONE_Median | 0.00 [0.00, 0.00] | 0 of 4 [0.00, 0.60] |
| MaAsLin 3 | TSS_LOG | 0.00 [0.00, 0.00] | 0 of 4 [0.00, 0.60] |
| DESeq2 | LRT_poscounts | 0.75 [0.00, 3.14] | 1 of 4 [0.01, 0.81] |
| DESeq2 | Wald_poscounts * | 1.75 [0.00, 5.03] | 2 of 4 [0.07, 0.93] |
| ZicoSeq | Default * | 0.00 [0.00, 0.00] | 0 of 4 [0.00, 0.60] |
| Wilcoxon | Rarefied | 0.00 [0.00, 0.00] | 0 of 4 [0.00, 0.60] |
| edgeR | TMM_LRT | 4.00 [0.56, 7.44] | 4 of 4 [0.40, 1.00] |
| LDM | Freq | 0.00 [0.00, 0.00] | 0 of 4 [0.00, 0.60] |
| ALDEx2 | Wilcox_128 | 0.00 [0.00, 0.00] | 0 of 4 [0.00, 0.60] |
| edgeR | TMM_QL * | 20.50 [8.00, 33.00] | 4 of 4 [0.40, 1.00] |
| ANCOM-BC2 | BH | 0.00 [0.00, 0.00] | 0 of 4 [0.00, 0.60] |
| ALDEx2 | Welch_128 * | 0.00 [0.00, 0.00] | 0 of 4 [0.00, 0.60] |
| ALDEx2 | Welch_32 | 0.00 [0.00, 0.00] | 0 of 4 [0.00, 0.60] |
| ALDEx2 | Welch_128_IQLR | 0.00 [0.00, 0.00] | 0 of 4 [0.00, 0.60] |
| ANCOM-BC2 | BH_StrucZero | 0.00 [0.00, 0.00] | 0 of 4 [0.00, 0.60] |
| MaAsLin 2 | TMM_NONE | 0.00 [0.00, 0.00] | 0 of 4 [0.00, 0.60] |
| ANCOM-BC2 | Holm * | 0.00 [0.00, 0.00] | 0 of 4 [0.00, 0.60] |
| ANCOM-BC2 | Holm_NoFilter | 6.25 [1.16, 11.34] | 4 of 4 [0.40, 1.00] |
| ANCOM-BC2 | BH_NoFilter | 26.50 [0.00, 53.89] | 4 of 4 [0.40, 1.00] |
| ***Simulated, stool, genus level*** | | | |
| LinDA | Pseudo_0.1 | 0.15 [0.00, 0.45] | 3 of 20 [0.03, 0.38] |
| MaAsLin 2 | TMM_LOG | 0.05 [0.00, 0.21] | 1 of 20 [0.00, 0.25] |
| LinDA | Imputation | 0.05 [0.00, 0.21] | 1 of 20 [0.00, 0.25] |
| MaAsLin 3 | TSS_PLOG_Median | 0.10 [0.00, 0.42] | 2 of 20 [0.01, 0.32] |
| LinDA | Pseudo_0.5 | 0.10 [0.00, 0.28] | 2 of 20 [0.01, 0.32] |
| MaAsLin 3 | TSS_PLOG | 0.05 [0.00, 0.21] | 1 of 20 [0.00, 0.25] |
| MaAsLin 2 | TSS_LOG * | 0.05 [0.00, 0.21] | 1 of 20 [0.00, 0.25] |
| Wilcoxon | TSS | 0.05 [0.00, 0.21] | 1 of 20 [0.00, 0.25] |
| LinDA | Adaptive * | 0.10 [0.00, 0.28] | 2 of 20 [0.01, 0.32] |
| LDM | CLR | 0.05 [0.00, 0.21] | 1 of 20 [0.00, 0.25] |
| LDM | Omni3 | 0.00 [0.00, 0.00] | 0 of 20 [0.00, 0.17] |
| LinDA | Adaptive_NoWinsor | 0.10 [0.00, 0.28] | 2 of 20 [0.01, 0.32] |
| MaAsLin 2 | CLR_NONE | 0.20 [0.00, 0.57] | 4 of 20 [0.06, 0.44] |
| limma-voom | TMM_voom | 0.05 [0.00, 0.21] | 1 of 20 [0.00, 0.25] |
| LDM | PA | 0.00 [0.00, 0.00] | 0 of 20 [0.00, 0.17] |

| Tool | Configuration | False positives per dataset | Datasets with at least one call |
|---|---|---|---|
| LDM | Arcsin | 0.00 [0.00, 0.00] | 0 of 20 [0.00, 0.17] |
| ZicoSeq | MultiLink | 0.05 [0.00, 0.21] | 1 of 20 [0.00, 0.25] |
| LinDA | Pseudo_1 | 0.10 [0.00, 0.28] | 2 of 20 [0.01, 0.32] |
| ZicoSeq | NoPosterior | 0.05 [0.00, 0.21] | 1 of 20 [0.00, 0.25] |
| MaAsLin 3 | TSS_LOG_Median * | 0.05 [0.00, 0.21] | 1 of 20 [0.00, 0.25] |
| LDM | Omni * | 0.00 [0.00, 0.00] | 0 of 20 [0.00, 0.17] |
| Wilcoxon | CLR | 0.05 [0.00, 0.21] | 1 of 20 [0.00, 0.25] |
| limma-voom | TMMwsp_voom | 0.10 [0.00, 0.28] | 2 of 20 [0.01, 0.32] |
| MaAsLin 3 | CLR_NONE | 0.15 [0.00, 0.63] | 2 of 20 [0.01, 0.32] |
| MaAsLin 3 | CLR_NONE_Median | 0.05 [0.00, 0.21] | 1 of 20 [0.00, 0.25] |
| MaAsLin 3 | TSS_LOG | 0.05 [0.00, 0.21] | 1 of 20 [0.00, 0.25] |
| DESeq2 | LRT_poscounts | 0.20 [0.20, 0.20] | 4 of 20 [0.06, 0.44] |
| DESeq2 | Wald_poscounts * | 0.35 [0.05, 0.65] | 7 of 20 [0.15, 0.59] |
| ZicoSeq | Default * | 0.15 [0.00, 0.45] | 3 of 20 [0.03, 0.38] |
| Wilcoxon | Rarefied | 0.00 [0.00, 0.00] | 0 of 20 [0.00, 0.17] |
| edgeR | TMM_LRT | 1.25 [0.71, 1.79] | 14 of 20 [0.46, 0.88] |
| LDM | Freq | 0.00 [0.00, 0.00] | 0 of 20 [0.00, 0.17] |
| ALDEx2 | Wilcox_128 | 0.00 [0.00, 0.00] | 0 of 20 [0.00, 0.17] |
| edgeR | TMM_QL * | 6.65 [3.85, 9.45] | 20 of 20 [0.83, 1.00] |
| ANCOM-BC2 | BH | 0.00 [0.00, 0.00] | 0 of 20 [0.00, 0.17] |
| ALDEx2 | Welch_128 * | 0.00 [0.00, 0.00] | 0 of 20 [0.00, 0.17] |
| ALDEx2 | Welch_32 | 0.00 [0.00, 0.00] | 0 of 20 [0.00, 0.17] |
| ALDEx2 | Welch_128_IQLR | 0.00 [0.00, 0.00] | 0 of 20 [0.00, 0.17] |
| ANCOM-BC2 | BH_StrucZero | 0.00 [0.00, 0.00] | 0 of 20 [0.00, 0.17] |
| MaAsLin 2 | TMM_NONE | 0.00 [0.00, 0.00] | 0 of 20 [0.00, 0.17] |
| ANCOM-BC2 | Holm * | 0.00 [0.00, 0.00] | 0 of 20 [0.00, 0.17] |
| ANCOM-BC2 | Holm_NoFilter | 6.65 [6.11, 7.19] | 20 of 20 [0.83, 1.00] |
| ANCOM-BC2 | BH_NoFilter | 11.65 [9.53, 13.77] | 20 of 20 [0.83, 1.00] |

**Table S7. Calibration against the nominal level.** Threshold design, at 100 and 200 samples, effect sizes 1.5 to 2.5 and both signal types, with every tool refitted at each adjusted p-value threshold. Empirical FDR with 95% confidence intervals across replicates, and mean recall.

| Tool | Configuration | FDR at 0.05 | Recall at 0.05 | FDR at 0.10 | Recall at 0.10 | FDR at 0.25 | Recall at 0.25 |
|---|---|---|---|---|---|---|---|
| ***Genus level*** | | | | | | | |
| LinDA | Pseudo_0.1 | 0.069 [0.000, 0.138] | 0.80 | 0.097 [0.019, 0.175] | 0.86 | 0.243 [0.207, 0.279] | 0.90 |
| MaAsLin 2 | TMM_LOG | 0.069 [0.014, 0.125] | 0.80 | 0.114 [0.018, 0.211] | 0.84 | 0.238 [0.178, 0.298] | 0.89 |
| LinDA | Imputation | 0.125 [0.005, 0.246] | 0.78 | 0.184 [0.058, 0.309] | 0.84 | 0.338 [0.251, 0.426] | 0.88 |
| MaAsLin 3 | TSS_PLOG_Median | 0.051 [0.000, 0.108] | 0.75 | 0.077 [0.000, 0.164] | 0.80 | 0.189 [0.055, 0.323] | 0.85 |
| LinDA | Pseudo_0.5 | 0.047 [0.006, 0.089] | 0.76 | 0.107 [0.054, 0.160] | 0.79 | 0.202 [0.135, 0.268] | 0.87 |
| MaAsLin 3 | TSS_PLOG | 0.119 [0.013, 0.224] | 0.82 | 0.157 [0.028, 0.285] | 0.86 | 0.307 [0.207, 0.407] | 0.91 |
| MaAsLin 2 | TSS_LOG * | 0.127 [0.026, 0.227] | 0.79 | 0.168 [0.043, 0.292] | 0.83 | 0.333 [0.214, 0.453] | 0.89 |
| Wilcoxon | TSS | 0.137 [0.039, 0.235] | 0.80 | 0.181 [0.079, 0.283] | 0.84 | 0.345 [0.230, 0.461] | 0.90 |
| LinDA | Adaptive * | 0.119 [0.000, 0.241] | 0.77 | 0.180 [0.031, 0.329] | 0.79 | 0.297 [0.170, 0.424] | 0.88 |
| LDM | CLR | 0.051 [0.011, 0.091] | 0.76 | 0.093 [0.044, 0.141] | 0.80 | 0.253 [0.205, 0.301] | 0.88 |
| LDM | Omni3 | 0.177 [0.017, 0.337] | 0.78 | 0.227 [0.066, 0.388] | 0.84 | 0.410 [0.294, 0.527] | 0.90 |
| LinDA | Adaptive_NoWinsor | 0.100 [0.000, 0.243] | 0.74 | 0.156 [0.014, 0.298] | 0.78 | 0.315 [0.180, 0.450] | 0.88 |
| MaAsLin 2 | CLR_NONE | 0.075 [0.033, 0.118] | 0.73 | 0.136 [0.045, 0.227] | 0.77 | 0.282 [0.185, 0.378] | 0.84 |
| limma-voom | TMM_voom | 0.119 [0.000, 0.242] | 0.72 | 0.171 [0.080, 0.263] | 0.76 | 0.313 [0.212, 0.414] | 0.83 |
| LDM | PA | 0.116 [0.004, 0.227] | 0.76 | 0.187 [0.024, 0.349] | 0.82 | 0.383 [0.277, 0.488] | 0.89 |
| LDM | Arcsin | 0.148 [0.008, 0.287] | 0.73 | 0.232 [0.085, 0.379] | 0.79 | 0.399 [0.289, 0.509] | 0.84 |
| ZicoSeq | MultiLink | 0.126 [0.024, 0.229] | 0.69 | 0.205 [0.075, 0.334] | 0.74 | 0.343 [0.215, 0.472] | 0.82 |
| LinDA | Pseudo_1 | 0.045 [0.014, 0.077] | 0.71 | 0.097 [0.044, 0.149] | 0.76 | 0.195 [0.154, 0.236] | 0.82 |
| ZicoSeq | NoPosterior | 0.058 [0.018, 0.098] | 0.69 | 0.112 [0.000, 0.231] | 0.73 | 0.243 [0.168, 0.318] | 0.81 |
| MaAsLin 3 | TSS_LOG_Median * | 0.036 [0.000, 0.076] | 0.73 | 0.050 [0.000, 0.099] | 0.77 | 0.139 [0.076, 0.202] | 0.83 |
| LDM | Omni * | 0.157 [0.021, 0.293] | 0.72 | 0.242 [0.035, 0.448] | 0.77 | 0.407 [0.308, 0.505] | 0.84 |
| Wilcoxon | CLR | 0.073 [0.042, 0.105] | 0.71 | 0.145 [0.084, 0.206] | 0.75 | 0.263 [0.210, 0.317] | 0.81 |
| limma-voom | TMMwsp_voom | 0.181 [0.096, 0.266] | 0.72 | 0.243 [0.145, 0.340] | 0.77 | 0.362 [0.221, 0.504] | 0.82 |
| MaAsLin 3 | CLR_NONE | 0.045 [0.018, 0.072] | 0.68 | 0.076 [0.009, 0.143] | 0.71 | 0.266 [0.205, 0.327] | 0.76 |
| MaAsLin 3 | CLR_NONE_Median | 0.024 [0.000, 0.058] | 0.68 | 0.044 [0.000, 0.103] | 0.71 | 0.146 [0.088, 0.205] | 0.75 |
| MaAsLin 3 | TSS_LOG | 0.143 [0.012, 0.273] | 0.75 | 0.175 [0.046, 0.304] | 0.78 | 0.325 [0.248, 0.402] | 0.85 |
| DESeq2 | LRT_poscounts | 0.162 [0.072, 0.252] | 0.62 | 0.280 [0.240, 0.320] | 0.64 | 0.491 [0.444, 0.538] | 0.71 |
| DESeq2 | Wald_poscounts * | 0.170 [0.082, 0.257] | 0.61 | 0.288 [0.231, 0.344] | 0.63 | 0.468 [0.426, 0.509] | 0.69 |
| ZicoSeq | Default * | 0.195 [0.022, 0.368] | 0.62 | 0.266 [0.104, 0.429] | 0.70 | 0.387 [0.167, 0.607] | 0.77 |
| Wilcoxon | Rarefied | 0.126 [0.025, 0.227] | 0.64 | 0.190 [0.060, 0.319] | 0.68 | 0.338 [0.229, 0.447] | 0.75 |
| edgeR | TMM_LRT | 0.355 [0.278, 0.432] | 0.67 | 0.449 [0.324, 0.574] | 0.71 | 0.605 [0.509, 0.702] | 0.77 |
| LDM | Freq | 0.134 [0.024, 0.244] | 0.60 | 0.199 [0.000, 0.412] | 0.62 | 0.374 [0.234, 0.514] | 0.70 |
| ALDEx2 | Wilcox_128 | 0.000 [0.000, 0.000] | 0.52 | 0.005 [0.000, 0.022] | 0.55 | 0.046 [0.000, 0.113] | 0.64 |
| edgeR | TMM_QL * | 0.560 [0.527, 0.593] | 0.73 | 0.643 [0.618, 0.668] | 0.75 | 0.742 [0.731, 0.752] | 0.84 |
| ANCOM-BC2 | BH | 0.000 [0.000, 0.000] | 0.44 | 0.008 [0.000, 0.035] | 0.48 | 0.047 [0.016, 0.078] | 0.55 |
| ALDEx2 | Welch_128 * | 0.007 [0.000, 0.029] | 0.47 | 0.010 [0.000, 0.044] | 0.52 | 0.035 [0.000, 0.088] | 0.61 |
| ALDEx2 | Welch_32 | 0.007 [0.000, 0.029] | 0.47 | 0.012 [0.000, 0.038] | 0.52 | 0.043 [0.000, 0.112] | 0.61 |
| ALDEx2 | Welch_128_IQLR | 0.012 [0.000, 0.051] | 0.45 | 0.009 [0.000, 0.036] | 0.52 | 0.021 [0.000, 0.061] | 0.59 |
| ANCOM-BC2 | BH_StrucZero | 0.000 [0.000, 0.000] | 0.43 | 0.008 [0.000, 0.035] | 0.46 | 0.048 [0.019, 0.078] | 0.51 |
| MaAsLin 2 | TMM_NONE | 0.012 [0.000, 0.036] | 0.38 | 0.042 [0.000, 0.122] | 0.45 | 0.114 [0.000, 0.253] | 0.54 |
| ANCOM-BC2 | Holm * | 0.000 [0.000, 0.000] | 0.36 | 0.000 [0.000, 0.000] | 0.38 | 0.000 [0.000, 0.000] | 0.42 |
| ANCOM-BC2 | Holm_NoFilter | 0.676 [0.589, 0.763] | 0.48 | 0.691 [0.623, 0.759] | 0.50 | 0.712 [0.657, 0.766] | 0.53 |
| ANCOM-BC2 | BH_NoFilter | 0.743 [0.697, 0.789] | 0.55 | 0.772 [0.741, 0.803] | 0.58 | 0.813 [0.801, 0.826] | 0.64 |
| ***OTU level*** | | | | | | | |
| LinDA | Pseudo_0.1 | 0.047 [0.030, 0.065] | 0.76 | 0.096 [0.070, 0.123] | 0.81 | 0.236 [0.170, 0.302] | 0.88 |
| MaAsLin 2 | TMM_LOG | 0.040 [0.030, 0.049] | 0.74 | 0.090 [0.055, 0.125] | 0.80 | 0.227 [0.181, 0.274] | 0.88 |
| LinDA | Imputation | 0.092 [0.061, 0.123] | 0.71 | 0.154 [0.111, 0.197] | 0.76 | 0.312 [0.248, 0.375] | 0.84 |

| Tool | Configuration | FDR at 0.05 | Recall at 0.05 | FDR at 0.10 | Recall at 0.10 | FDR at 0.25 | Recall at 0.25 |
|---|---|---|---|---|---|---|---|
| MaAsLin 3 | TSS_PLOG_Median | 0.053 [0.036, 0.069] | 0.71 | 0.101 [0.072, 0.129] | 0.76 | 0.242 [0.174, 0.310] | 0.84 |
| LinDA | Pseudo_0.5 | 0.044 [0.030, 0.057] | 0.74 | 0.090 [0.063, 0.118] | 0.79 | 0.227 [0.166, 0.287] | 0.86 |
| MaAsLin 3 | TSS_PLOG | 0.070 [0.044, 0.096] | 0.78 | 0.134 [0.086, 0.182] | 0.83 | 0.303 [0.240, 0.365] | 0.90 |
| MaAsLin 2 | TSS_LOG * | 0.099 [0.065, 0.132] | 0.76 | 0.166 [0.115, 0.216] | 0.81 | 0.342 [0.278, 0.405] | 0.90 |
| Wilcoxon | TSS | 0.073 [0.047, 0.099] | 0.76 | 0.125 [0.080, 0.169] | 0.82 | 0.280 [0.215, 0.345] | 0.89 |
| LinDA | Adaptive * | 0.092 [0.067, 0.116] | 0.70 | 0.153 [0.120, 0.187] | 0.75 | 0.311 [0.248, 0.373] | 0.83 |
| LDM | CLR | 0.043 [0.034, 0.053] | 0.74 | 0.092 [0.069, 0.115] | 0.80 | 0.225 [0.160, 0.289] | 0.87 |
| LDM | Omni3 | 0.216 [0.187, 0.246] | 0.78 | 0.297 [0.242, 0.353] | 0.83 | 0.461 [0.383, 0.539] | 0.90 |
| LinDA | Adaptive_NoWinsor | 0.065 [0.042, 0.087] | 0.69 | 0.122 [0.088, 0.155] | 0.74 | 0.272 [0.201, 0.343] | 0.83 |
| MaAsLin 2 | CLR_NONE | 0.134 [0.121, 0.148] | 0.79 | 0.215 [0.187, 0.242] | 0.84 | 0.394 [0.332, 0.457] | 0.91 |
| limma-voom | TMM_voom | 0.333 [0.185, 0.481] | 0.79 | 0.412 [0.283, 0.541] | 0.83 | 0.570 [0.480, 0.661] | 0.89 |
| LDM | PA | 0.221 [0.192, 0.249] | 0.77 | 0.299 [0.240, 0.358] | 0.82 | 0.463 [0.387, 0.538] | 0.90 |
| LDM | Arcsin | 0.166 [0.144, 0.188] | 0.77 | 0.249 [0.201, 0.296] | 0.82 | 0.423 [0.347, 0.500] | 0.90 |
| ZicoSeq | MultiLink | 0.075 [0.052, 0.099] | 0.68 | 0.139 [0.100, 0.179] | 0.73 | 0.317 [0.254, 0.379] | 0.82 |
| LinDA | Pseudo_1 | 0.041 [0.030, 0.052] | 0.72 | 0.084 [0.058, 0.109] | 0.77 | 0.219 [0.163, 0.275] | 0.84 |
| ZicoSeq | NoPosterior | 0.053 [0.031, 0.075] | 0.75 | 0.102 [0.065, 0.139] | 0.80 | 0.242 [0.175, 0.309] | 0.88 |
| MaAsLin 3 | TSS_LOG_Median * | 0.030 [0.020, 0.040] | 0.60 | 0.059 [0.044, 0.074] | 0.66 | 0.170 [0.144, 0.197] | 0.76 |
| LDM | Omni * | 0.210 [0.182, 0.237] | 0.76 | 0.290 [0.238, 0.343] | 0.82 | 0.456 [0.386, 0.526] | 0.89 |
| Wilcoxon | CLR | 0.336 [0.308, 0.364] | 0.73 | 0.404 [0.323, 0.484] | 0.78 | 0.519 [0.387, 0.652] | 0.85 |
| limma-voom | TMMwsp_voom | 0.476 [0.361, 0.591] | 0.81 | 0.551 [0.418, 0.683] | 0.86 | 0.670 [0.532, 0.807] | 0.91 |
| MaAsLin 3 | CLR_NONE | 0.108 [0.098, 0.118] | 0.63 | 0.174 [0.166, 0.181] | 0.69 | 0.331 [0.313, 0.349] | 0.78 |
| MaAsLin 3 | CLR_NONE_Median | 0.032 [0.024, 0.040] | 0.63 | 0.066 [0.053, 0.079] | 0.68 | 0.174 [0.155, 0.193] | 0.76 |
| MaAsLin 3 | TSS_LOG | 0.265 [0.225, 0.304] | 0.66 | 0.310 [0.266, 0.355] | 0.72 | 0.437 [0.368, 0.506] | 0.82 |
| DESeq2 | LRT_poscounts | 0.118 [0.091, 0.144] | 0.55 | 0.199 [0.150, 0.248] | 0.59 | 0.395 [0.341, 0.450] | 0.65 |
| DESeq2 | Wald_poscounts * | 0.127 [0.100, 0.153] | 0.52 | 0.204 [0.163, 0.245] | 0.55 | 0.391 [0.339, 0.442] | 0.61 |
| ZicoSeq | Default * | 0.074 [0.044, 0.103] | 0.67 | 0.136 [0.091, 0.180] | 0.73 | 0.298 [0.237, 0.359] | 0.82 |
| Wilcoxon | Rarefied | 0.101 [0.089, 0.114] | 0.44 | 0.161 [0.136, 0.185] | 0.51 | 0.325 [0.284, 0.366] | 0.62 |
| edgeR | TMM_LRT | 0.265 [0.235, 0.295] | 0.71 | 0.353 [0.328, 0.379] | 0.77 | 0.524 [0.504, 0.545] | 0.86 |
| LDM | Freq | 0.187 [0.160, 0.215] | 0.65 | 0.263 [0.222, 0.304] | 0.71 | 0.420 [0.355, 0.486] | 0.80 |
| ALDEx2 | Wilcox_128 | 0.000 [0.000, 0.001] | 0.34 | 0.001 [0.000, 0.002] | 0.38 | 0.004 [0.000, 0.009] | 0.46 |
| edgeR | TMM_QL * | 0.409 [0.382, 0.435] | 0.75 | 0.492 [0.468, 0.515] | 0.80 | 0.635 [0.621, 0.649] | 0.88 |
| ANCOM-BC2 | BH | 0.007 [0.004, 0.011] | 0.41 | 0.015 [0.008, 0.023] | 0.46 | 0.052 [0.030, 0.074] | 0.54 |
| ALDEx2 | Welch_128 * | 0.000 [0.000, 0.000] | 0.27 | 0.000 [0.000, 0.001] | 0.31 | 0.001 [0.000, 0.002] | 0.39 |
| ALDEx2 | Welch_32 | 0.000 [0.000, 0.000] | 0.27 | 0.000 [0.000, 0.001] | 0.31 | 0.001 [0.000, 0.002] | 0.40 |
| ALDEx2 | Welch_128_IQLR | 0.000 [0.000, 0.000] | 0.27 | 0.000 [0.000, 0.000] | 0.31 | 0.001 [0.000, 0.002] | 0.38 |
| ANCOM-BC2 | BH_StrucZero | 0.008 [0.003, 0.012] | 0.37 | 0.014 [0.008, 0.019] | 0.41 | 0.048 [0.031, 0.065] | 0.46 |
| MaAsLin 2 | TMM_NONE | 0.018 [0.011, 0.024] | 0.45 | 0.041 [0.034, 0.047] | 0.53 | 0.140 [0.135, 0.144] | 0.65 |
| ANCOM-BC2 | Holm * | 0.000 [0.000, 0.000] | 0.26 | 0.000 [0.000, 0.000] | 0.27 | 0.000 [0.000, 0.000] | 0.29 |
| ANCOM-BC2 | Holm_NoFilter | 0.685 [0.677, 0.694] | 0.40 | 0.710 [0.704, 0.716] | 0.41 | 0.742 [0.737, 0.746] | 0.43 |
| ANCOM-BC2 | BH_NoFilter | 0.862 [0.858, 0.867] | 0.55 | 0.876 [0.872, 0.879] | 0.58 | 0.891 [0.888, 0.894] | 0.63 |

**Table S8. Variance decomposition.** Sums of squares of each term as a percentage of the total in the main detection design. The extreme configurations are MaAsLin 2 TMM_NONE and the two ANCOM-BC2 configurations without sensitivity analysis. The third configuration set leaves out edgeR and the two ANCOM-BC2 configurations without sensitivity analysis, the most liberal configurations (39 configurations). The configuration fraction of method is the configuration term divided by the sum of the tool and configuration terms. The replicate term measures how single datasets vary around their scenario, which more replicates estimate more precisely but do not reduce.

| Data | Replicates | Resolution | Metric | Configurations | Scenario (%) | Tool (%) | Configuration within tool (%) | Scenario x tool (%) | Scenario x configuration (%) | Replicate (%) | Configuration fraction of method (%) |
|---|---|---|---|---|---|---|---|---|---|---|---|
| Semi-synthetic | 4 | Genus | MCC | all | 64.37 | 7.33 | 2.33 | 7.01 | 3.68 | 15.28 | 24.1 |
| Semi-synthetic | 4 | Genus | MCC | without extreme | 68.42 | 5.75 | 1.01 | 6.56 | 2.72 | 15.54 | 14.9 |
| Semi-synthetic | 4 | Genus | MCC | without edgeR and ANCOM-BC2 NoFilter | 67.52 | 4.96 | 2.24 | 6.31 | 3.21 | 15.77 | 31.1 |
| Semi-synthetic | 4 | Genus | FDR | all | 2.87 | 24.89 | 29.75 | 5.94 | 5.65 | 30.91 | 54.4 |
| Semi-synthetic | 4 | Genus | FDR | without extreme | 5.58 | 28.76 | 2.94 | 7.47 | 6.04 | 49.21 | 9.3 |
| Semi-synthetic | 4 | Genus | FDR | without edgeR and ANCOM-BC2 NoFilter | 8.68 | 5.55 | 1.21 | 7.98 | 7.78 | 68.79 | 17.9 |
| Semi-synthetic | 4 | Genus | Recall | all | 70.83 | 5.82 | 3.16 | 6.10 | 2.89 | 11.19 | 35.2 |
| Semi-synthetic | 4 | Genus | Recall | without extreme | 72.96 | 7.35 | 1.07 | 5.37 | 2.11 | 11.13 | 12.7 |
| Semi-synthetic | 4 | Genus | Recall | without edgeR and ANCOM-BC2 NoFilter | 71.60 | 6.75 | 2.59 | 5.14 | 2.70 | 11.22 | 27.8 |
| Semi-synthetic | 4 | OTU | MCC | all | 65.44 | 13.26 | 3.55 | 10.68 | 5.27 | 1.79 | 21.1 |
| Semi-synthetic | 4 | OTU | MCC | without extreme | 71.45 | 11.03 | 2.11 | 9.68 | 3.84 | 1.90 | 16.0 |
| Semi-synthetic | 4 | OTU | MCC | without edgeR and ANCOM-BC2 NoFilter | 70.90 | 10.68 | 2.88 | 9.36 | 4.27 | 1.91 | 21.2 |
| Semi-synthetic | 4 | OTU | FDR | all | 6.87 | 27.01 | 34.47 | 14.58 | 10.20 | 6.87 | 56.1 |
| Semi-synthetic | 4 | OTU | FDR | without extreme | 13.83 | 34.37 | 3.83 | 22.57 | 13.39 | 12.01 | 10.0 |
| Semi-synthetic | 4 | OTU | FDR | without edgeR and ANCOM-BC2 NoFilter | 21.70 | 19.89 | 5.10 | 19.80 | 18.02 | 15.48 | 20.4 |
| Semi-synthetic | 4 | OTU | Recall | all | 72.68 | 10.87 | 3.79 | 8.99 | 2.66 | 1.02 | 25.8 |
| Semi-synthetic | 4 | OTU | Recall | without extreme | 75.11 | 13.17 | 1.41 | 7.49 | 1.77 | 1.04 | 9.7 |
| Semi-synthetic | 4 | OTU | Recall | without edgeR and ANCOM-BC2 NoFilter | 74.40 | 12.78 | 2.50 | 7.11 | 2.15 | 1.07 | 16.3 |
| Semi-synthetic | 20 | Genus | MCC | all | 62.15 | 7.35 | 2.31 | 5.65 | 2.48 | 20.06 | 23.9 |
| Semi-synthetic | 20 | Genus | MCC | without extreme | 65.88 | 5.80 | 0.92 | 5.22 | 1.48 | 20.70 | 13.7 |
| Semi-synthetic | 20 | Genus | FDR | all | 1.51 | 26.72 | 30.29 | 3.52 | 2.92 | 35.03 | 53.1 |
| Semi-synthetic | 20 | Genus | FDR | without extreme | 2.89 | 31.69 | 2.66 | 4.12 | 2.03 | 56.62 | 7.7 |
| Semi-synthetic | 20 | Genus | Recall | all | 67.86 | 5.34 | 3.16 | 4.99 | 1.95 | 16.70 | 37.2 |
| Semi-synthetic | 20 | Genus | Recall | without extreme | 69.89 | 6.83 | 0.97 | 4.21 | 1.23 | 16.87 | 12.5 |
| Simulated | 4 | Genus | MCC | all | 62.99 | 8.22 | 1.74 | 6.90 | 2.77 | 17.38 | 17.4 |
| Simulated | 4 | Genus | MCC | without extreme | 66.61 | 6.17 | 0.74 | 6.75 | 1.68 | 18.06 | 10.7 |
| Simulated | 4 | Genus | MCC | without edgeR and ANCOM-BC2 NoFilter | 66.85 | 5.49 | 1.51 | 6.17 | 2.22 | 17.76 | 21.5 |
| Simulated | 4 | Genus | FDR | all | 10.23 | 17.03 | 24.22 | 8.17 | 4.91 | 35.44 | 58.7 |
| Simulated | 4 | Genus | FDR | without extreme | 15.62 | 19.17 | 2.38 | 9.56 | 4.77 | 48.51 | 11.0 |
| Simulated | 4 | Genus | FDR | without edgeR and ANCOM-BC2 NoFilter | 19.71 | 6.84 | 0.77 | 9.60 | 5.68 | 57.39 | 10.2 |
| Simulated | 4 | Genus | Recall | all | 70.98 | 6.52 | 2.27 | 4.97 | 2.11 | 13.15 | 25.8 |
| Simulated | 4 | Genus | Recall | without extreme | 72.23 | 7.63 | 0.97 | 4.43 | 1.53 | 13.20 | 11.3 |
| Simulated | 4 | Genus | Recall | without edgeR and ANCOM-BC2 NoFilter | 71.97 | 6.94 | 1.92 | 4.13 | 1.92 | 13.12 | 21.7 |
| IBD, semi-synthetic | 4 | Genus | MCC | all | 70.52 | 5.73 | 3.31 | 4.13 | 2.59 | 13.71 | 36.6 |
| IBD, semi-synthetic | 4 | Genus | MCC | without extreme | 74.07 | 5.26 | 1.06 | 4.15 | 1.45 | 14.02 | 16.7 |
| IBD, semi-synthetic | 4 | Genus | FDR | all | 0.45 | 44.28 | 29.21 | 5.10 | 3.73 | 17.23 | 39.7 |
| IBD, semi-synthetic | 4 | Genus | FDR | without extreme | 0.33 | 59.98 | 3.27 | 7.01 | 2.71 | 26.70 | 5.2 |
| IBD, semi-synthetic | 4 | Genus | Recall | all | 71.96 | 4.61 | 5.45 | 3.68 | 2.75 | 11.55 | 54.2 |
| IBD, semi-synthetic | 4 | Genus | Recall | without extreme | 75.41 | 6.58 | 1.49 | 3.65 | 1.33 | 11.54 | 18.5 |

**Table S9. Configuration against tool in each scenario.** Main detection design, semi-synthetic data. Within tools is the mean over tools of the difference in MCC between the configuration chosen on three replicates and the median configuration of the tool. Between tools is the difference in MCC between the best and the median tool, each with its chosen configuration. With 20 replicates at genus level, the gap between tools exceeded the gain within tools in 24 of 24 scenarios.

| Samples | Effect size | Signal | Within tools | Between tools | Best tool |
|---|---|---|---|---|---|
| ***Genus level*** | | | | | |
| 50 | 1.0 | joint | 0.051 | 0.085 | MaAsLin 2 |
| 50 | 1.0 | prevalence only | 0.007 | 0.076 | DESeq2 |
| 50 | 1.5 | joint | 0.018 | 0.090 | MaAsLin 2 |
| 50 | 1.5 | prevalence only | 0.013 | 0.177 | ZicoSeq |
| 50 | 2.0 | joint | 0.045 | 0.049 | limma-voom |
| 50 | 2.0 | prevalence only | 0.053 | 0.186 | Wilcoxon |
| 50 | 2.5 | joint | 0.011 | 0.038 | DESeq2 |
| 50 | 2.5 | prevalence only | 0.055 | 0.186 | LinDA |
| 100 | 1.0 | joint | 0.031 | 0.181 | LinDA |
| 100 | 1.0 | prevalence only | 0.011 | 0.130 | LinDA |
| 100 | 1.5 | joint | 0.011 | 0.094 | LinDA |
| 100 | 1.5 | prevalence only | 0.038 | 0.324 | MaAsLin 3 |
| 100 | 2.0 | joint | 0.020 | 0.065 | ZicoSeq |
| 100 | 2.0 | prevalence only | 0.037 | 0.122 | MaAsLin 3 |
| 100 | 2.5 | joint | 0.060 | 0.091 | LDM |
| 100 | 2.5 | prevalence only | 0.042 | 0.055 | MaAsLin 3 |
| 200 | 1.0 | joint | 0.017 | 0.059 | LDM |
| 200 | 1.0 | prevalence only | 0.075 | 0.185 | MaAsLin 3 |
| 200 | 1.5 | joint | 0.021 | 0.102 | LDM |
| 200 | 1.5 | prevalence only | 0.060 | 0.208 | MaAsLin 3 |
| 200 | 2.0 | joint | 0.023 | 0.085 | LDM |
| 200 | 2.0 | prevalence only | 0.092 | 0.073 | LDM |
| 200 | 2.5 | joint | −0.027 | 0.176 | limma-voom |
| 200 | 2.5 | prevalence only | 0.008 | 0.026 | MaAsLin 3 |
| ***OTU level*** | | | | | |
| 50 | 1.0 | joint | 0.056 | 0.310 | ANCOM-BC2 |
| 50 | 1.0 | prevalence only | 0.003 | 0.020 | ZicoSeq |
| 50 | 1.5 | joint | 0.055 | 0.114 | ZicoSeq |
| 50 | 1.5 | prevalence only | 0.026 | 0.120 | ZicoSeq |
| 50 | 2.0 | joint | 0.051 | 0.049 | MaAsLin 2 |
| 50 | 2.0 | prevalence only | 0.088 | 0.138 | ZicoSeq |
| 50 | 2.5 | joint | 0.060 | 0.047 | MaAsLin 2 |
| 50 | 2.5 | prevalence only | 0.061 | 0.063 | LinDA |
| 100 | 1.0 | joint | 0.054 | 0.081 | ZicoSeq |
| 100 | 1.0 | prevalence only | 0.013 | 0.037 | ZicoSeq |
| 100 | 1.5 | joint | 0.056 | 0.076 | MaAsLin 2 |
| 100 | 1.5 | prevalence only | 0.063 | 0.132 | MaAsLin 3 |
| 100 | 2.0 | joint | 0.043 | 0.076 | MaAsLin 2 |
| 100 | 2.0 | prevalence only | 0.056 | 0.046 | LinDA |
| 100 | 2.5 | joint | 0.099 | 0.051 | LDM |
| 100 | 2.5 | prevalence only | 0.050 | 0.051 | MaAsLin 3 |
| 200 | 1.0 | joint | 0.055 | 0.060 | MaAsLin 2 |
| 200 | 1.0 | prevalence only | 0.065 | 0.104 | LinDA |
| 200 | 1.5 | joint | 0.056 | 0.048 | MaAsLin 2 |
| 200 | 1.5 | prevalence only | 0.074 | 0.078 | Wilcoxon |

| Samples | Effect size | Signal | Within tools | Between tools | Best tool |
|---|---|---|---|---|---|
| 200 | 2.0 | joint | 0.093 | 0.027 | LDM |
| 200 | 2.0 | prevalence only | 0.059 | 0.048 | Wilcoxon |
| 200 | 2.5 | joint | 0.149 | 0.109 | LinDA |
| 200 | 2.5 | prevalence only | 0.045 | 0.023 | Wilcoxon |

**Table S10. Zero handling in LinDA.** Paired differences from LinDA with a pseudo-count of 0.5 on the same datasets in the main detection design, with 95% confidence intervals across replicates. The adaptive configurations use imputation when library size is associated with the group and a pseudo-count of 0.5 otherwise, with the switch applied.

| Configuration | Difference in MCC | Difference in FDR | Difference in recall |
|---|---|---|---|
| ***Semi-synthetic, genus level*** | | | |
| Adaptive | −0.015 [−0.059, 0.029] | 0.010 [−0.034, 0.055] | −0.004 [−0.018, 0.010] |
| Adaptive_NoWinsor | −0.024 [−0.081, 0.034] | 0.011 [−0.042, 0.065] | −0.012 [−0.036, 0.013] |
| Imputation | 0.012 [−0.044, 0.068] | 0.011 [−0.054, 0.076] | 0.027 [−0.013, 0.068] |
| Pseudo_0.1 | 0.026 [−0.014, 0.066] | 0.007 [−0.009, 0.023] | 0.033 [−0.019, 0.084] |
| Pseudo_1 | −0.047 [−0.069, −0.025] | −0.003 [−0.015, 0.008] | −0.047 [−0.076, −0.017] |
| ***Semi-synthetic, OTU level*** | | | |
| Adaptive | −0.029 [−0.035, −0.024] | 0.032 [0.021, 0.043] | −0.020 [−0.029, −0.012] |
| Adaptive_NoWinsor | −0.030 [−0.039, −0.020] | 0.016 [0.010, 0.022] | −0.030 [−0.035, −0.026] |
| Imputation | −0.025 [−0.033, −0.016] | 0.032 [0.016, 0.048] | −0.015 [−0.026, −0.005] |
| Pseudo_0.1 | 0.010 [0.002, 0.018] | 0.005 [−0.002, 0.012] | 0.014 [0.009, 0.019] |
| Pseudo_1 | −0.014 [−0.017, −0.011] | −0.004 [−0.006, −0.002] | −0.018 [−0.022, −0.014] |
| ***Simulated, genus level*** | | | |
| Adaptive | 0.002 [−0.005, 0.009] | −0.002 [−0.006, 0.002] | 0.002 [−0.006, 0.011] |
| Adaptive_NoWinsor | 0.003 [−0.006, 0.013] | −0.007 [−0.018, 0.005] | 0.002 [−0.007, 0.010] |
| Imputation | 0.000 [−0.009, 0.009] | 0.014 [0.000, 0.028] | 0.020 [0.015, 0.024] |
| Pseudo_0.1 | 0.019 [0.007, 0.031] | −0.011 [−0.027, 0.005] | 0.020 [0.011, 0.030] |
| Pseudo_1 | −0.019 [−0.022, −0.016] | −0.003 [−0.019, 0.014] | −0.018 [−0.026, −0.009] |
| ***IBD community, semi-synthetic, genus level*** | | | |
| Adaptive | 0.002 [−0.005, 0.009] | −0.004 [−0.017, 0.009] | 0.000 [0.000, 0.000] |
| Adaptive_NoWinsor | −0.003 [−0.016, 0.010] | −0.001 [−0.022, 0.020] | −0.007 [−0.021, 0.006] |
| Imputation | −0.007 [−0.033, 0.018] | 0.028 [−0.079, 0.136] | −0.007 [−0.052, 0.037] |
| Pseudo_0.1 | 0.012 [−0.034, 0.059] | 0.003 [−0.018, 0.024] | 0.015 [−0.028, 0.057] |
| Pseudo_1 | −0.007 [−0.032, 0.017] | 0.005 [−0.033, 0.043] | −0.007 [−0.038, 0.023] |

**Table S11. Configuration chosen out of sample.** For each scenario and tool, the configuration with the highest mean MCC on three replicates was scored on the fourth, in rotation, with ties resolved in favour of the default. Means with 95% confidence intervals across replicates. Most often chosen gives the proportion of scenario and replicate combinations. The last two columns repeat the choice on 19 replicates, scored on the twentieth. limma-voom and the Wilcoxon test have no single default.

| Tool | MCC default | MCC chosen | Gain over default | FDR default | FDR chosen | Most often chosen | Gain over default, 20 replicates | FDR change, 20 replicates |
|---|---|---|---|---|---|---|---|---|
| ***Semi-synthetic, genus level*** | | | | | | | | |
| ALDEx2 | 0.411 [0.372, 0.450] | 0.442 [0.388, 0.496] | 0.031 [−0.014, 0.076] | 0.000 | 0.005 | Wilcox_128 56%; Welch_128 38%; Welch_32 4% | 0.041 [0.033, 0.050] | 0.002 |
| ANCOM-BC2 | 0.353 [0.328, 0.379] | 0.425 [0.391, 0.460] | 0.072 [0.040, 0.104] | 0.000 | 0.235 | BH 57%; BH_NoFilter 15%; Holm_NoFilter 15% | 0.086 [0.073, 0.099] | 0.271 |
| DESeq2 | 0.536 [0.499, 0.573] | 0.539 [0.498, 0.579] | 0.003 [−0.006, 0.012] | 0.163 | 0.159 | Wald_poscounts 60%; LRT_poscounts 40% | 0.008 [0.003, 0.013] | −0.008 |
| LDM | 0.595 [0.529, 0.660] | 0.624 [0.575, 0.672] | 0.029 [−0.036, 0.094] | 0.090 | 0.052 | CLR 32%; Omni3 21%; PA 17% | 0.061 [0.047, 0.076] | −0.027 |
| LinDA | 0.634 [0.611, 0.657] | 0.669 [0.638, 0.700] | 0.035 [0.011, 0.059] | 0.059 | 0.055 | Pseudo_0.1 36%; Imputation 33%; Adaptive 19% | 0.044 [0.034, 0.055] | −0.013 |
| MaAsLin 2 | 0.641 [0.604, 0.677] | 0.657 [0.607, 0.707] | 0.016 [−0.003, 0.036] | 0.064 | 0.058 | TMM_LOG 43%; TSS_LOG 40%; CLR_NONE 18% | 0.025 [0.016, 0.034] | −0.016 |
| MaAsLin 3 | 0.598 [0.580, 0.617] | 0.638 [0.597, 0.678] | 0.039 [0.016, 0.063] | 0.031 | 0.047 | TSS_PLOG 41%; TSS_PLOG_Median 23%; CLR_NONE 15% | 0.070 [0.059, 0.081] | 0.014 |
| Wilcoxon | | 0.645 [0.614, 0.676] | | | 0.067 | TSS 60%; CLR 34%; Rarefied 5% | | |
| ZicoSeq | 0.518 [0.458, 0.579] | 0.622 [0.585, 0.659] | 0.104 [0.028, 0.180] | 0.116 | 0.058 | NoPosterior 42%; MultiLink 38%; Default 21% | 0.072 [0.056, 0.088] | −0.039 |
| edgeR | 0.434 [0.394, 0.474] | 0.497 [0.457, 0.538] | 0.063 [0.058, 0.068] | 0.571 | 0.406 | TMM_LRT 75%; TMM_QL 25% | 0.075 [0.064, 0.085] | −0.193 |
| limma-voom | | 0.605 [0.588, 0.622] | | | 0.079 | TMM_voom 78%; TMMwsp_voom 22% | | |
| ***Semi-synthetic, OTU level*** | | | | | | | | |
| ALDEx2 | 0.271 [0.266, 0.276] | 0.340 [0.329, 0.351] | 0.069 [0.060, 0.077] | 0.000 | 0.000 | Wilcox_128 77%; Welch_128 23% | | |
| ANCOM-BC2 | 0.261 [0.253, 0.270] | 0.450 [0.444, 0.457] | 0.189 [0.182, 0.196] | 0.000 | 0.128 | BH 75%; Holm_NoFilter 20%; BH_NoFilter 3% | | |
| DESeq2 | 0.502 [0.494, 0.511] | 0.521 [0.511, 0.531] | 0.019 [0.017, 0.021] | 0.131 | 0.121 | LRT_poscounts 78%; Wald_poscounts 22% | | |
| LDM | 0.551 [0.545, 0.556] | 0.609 [0.600, 0.617] | 0.058 [0.052, 0.064] | 0.127 | 0.046 | CLR 45%; Omni 18%; Omni3 17% | | |
| LinDA | 0.591 [0.574, 0.608] | 0.641 [0.627, 0.654] | 0.050 [0.042, 0.057] | 0.065 | 0.033 | Pseudo_0.1 42%; Pseudo_0.5 19%; Pseudo_1 16% | | |
| MaAsLin 2 | 0.605 [0.589, 0.620] | 0.629 [0.615, 0.642] | 0.024 [0.021, 0.027] | 0.064 | 0.045 | TSS_LOG 40%; CLR_NONE 36%; TMM_LOG 24% | | |
| MaAsLin 3 | 0.533 [0.519, 0.547] | 0.633 [0.617, 0.649] | 0.100 [0.098, 0.102] | 0.035 | 0.051 | TSS_PLOG 60%; TSS_PLOG_Median 22%; CLR_NONE_Median 14% | | |
| Wilcoxon | | 0.593 [0.585, 0.601] | | | 0.051 | TSS 94%; CLR 6% | | |
| ZicoSeq | 0.600 [0.583, 0.618] | 0.650 [0.630, 0.669] | 0.049 [0.040, 0.058] | 0.058 | 0.046 | NoPosterior 83%; MultiLink 8%; Default 8% | | |
| edgeR | 0.482 [0.471, 0.493] | 0.520 [0.501, 0.539] | 0.038 [0.029, 0.048] | 0.465 | 0.389 | TMM_LRT 55%; TMM_QL 45% | | |
| limma-voom | | 0.498 [0.455, 0.540] | | | 0.244 | TMM_voom 62%; TMMwsp_voom 38% | | |
| ***Simulated, genus level*** | | | | | | | | |
| ALDEx2 | 0.419 [0.379, 0.458] | 0.432 [0.388, 0.475] | 0.013 [0.002, 0.024] | 0.031 | 0.026 | Welch_128 46%; Wilcox_128 29%; Welch_128_IQLR 18% | | |
| ANCOM-BC2 | 0.339 [0.310, 0.368] | 0.383 [0.338, 0.429] | 0.045 [0.012, 0.078] | 0.000 | 0.220 | BH 50%; Holm 22%; Holm_NoFilter 16% | | |
| DESeq2 | 0.453 [0.423, 0.484] | 0.460 [0.430, 0.490] | 0.007 [−0.002, 0.015] | 0.254 | 0.246 | Wald_poscounts 66%; LRT_poscounts 34% | | |
| LDM | 0.577 [0.546, 0.609] | 0.612 [0.569, 0.655] | 0.035 [0.014, 0.055] | 0.122 | 0.110 | CLR 40%; PA 26%; Omni 11% | | |
| LinDA | 0.620 [0.589, 0.652] | 0.615 [0.578, 0.652] | −0.005 [−0.014, 0.003] | 0.098 | 0.106 | Adaptive 31%; Imputation 27%; Pseudo_0.1 22% | | |
| MaAsLin 2 | 0.624 [0.600, 0.648] | 0.638 [0.619, 0.658] | 0.014 [−0.024, 0.052] | 0.098 | 0.074 | TSS_LOG 54%; TMM_LOG 32%; CLR_NONE 9% | | |
| MaAsLin 3 | 0.598 [0.557, 0.639] | 0.619 [0.578, 0.661] | 0.021 [0.016, 0.026] | 0.030 | 0.070 | TSS_PLOG 59%; TSS_LOG_Median 21%; TSS_PLOG_Median 11% | | |
| Wilcoxon | | 0.614 [0.566, 0.663] | | | 0.120 | TSS 72%; CLR 19%; Rarefied 8% | | |
| ZicoSeq | 0.555 [0.522, 0.588] | 0.613 [0.585, 0.642] | 0.058 [0.021, 0.096] | 0.068 | 0.071 | NoPosterior 44%; MultiLink 43%; Default 13% | | |
| edgeR | 0.419 [0.385, 0.453] | 0.495 [0.468, 0.523] | 0.076 [0.060, 0.092] | 0.593 | 0.391 | TMM_LRT 77%; TMM_QL 23% | | |
| limma-voom | | 0.542 [0.494, 0.590] | | | 0.163 | TMM_voom 56%; TMMwsp_voom 44% | | |
| ***IBD community, semi-synthetic, genus level*** | | | | | | | | |
| ALDEx2 | 0.322 [0.270, 0.373] | 0.372 [0.323, 0.422] | 0.051 [0.021, 0.080] | 0.000 | 0.000 | Wilcox_128 62%; Welch_128 38% | | |
| ANCOM-BC2 | 0.231 [0.156, 0.306] | 0.304 [0.183, 0.424] | 0.072 [−0.055, 0.200] | 0.000 | 0.334 | BH 50%; BH_NoFilter 25%; Holm_NoFilter 19% | | |
| DESeq2 | 0.456 [0.368, 0.544] | 0.447 [0.377, 0.517] | −0.009 [−0.043, 0.024] | 0.340 | 0.322 | Wald_poscounts 62%; LRT_poscounts 38% | | |
| LDM | 0.460 [0.315, 0.604] | 0.489 [0.314, 0.663] | 0.029 [−0.005, 0.063] | 0.018 | 0.025 | CLR 44%; Omni 25%; PA 12% | | |
| LinDA | 0.508 [0.384, 0.633] | 0.526 [0.425, 0.627] | 0.018 [−0.011, 0.047] | 0.033 | 0.041 | Adaptive 38%; Pseudo_1 31%; Pseudo_0.1 31% | | |
| MaAsLin 2 | 0.505 [0.378, 0.631] | 0.530 [0.419, 0.642] | 0.025 [−0.007, 0.058] | 0.023 | 0.028 | TSS_LOG 50%; CLR_NONE 44%; TMM_LOG 6% | | |

| Tool | MCC default | MCC chosen | Gain over default | FDR default | FDR chosen | Most often chosen | Gain over default, 20 replicates | FDR change, 20 replicates |
|---|---|---|---|---|---|---|---|---|
| MaAsLin 3 | 0.418 [0.271, 0.566] | 0.481 [0.365, 0.597] | 0.062 [0.004, 0.121] | 0.022 | 0.014 | TSS_PLOG 50%; TSS_PLOG_Median 31%; TSS_LOG_Median 19% | | |
| Wilcoxon | | 0.473 [0.357, 0.589] | | | 0.026 | CLR 50%; TSS 50% | | |
| ZicoSeq | 0.410 [0.304, 0.516] | 0.463 [0.363, 0.563] | 0.053 [−0.054, 0.161] | 0.059 | 0.074 | MultiLink 62%; Default 25%; NoPosterior 12% | | |
| edgeR | 0.310 [0.229, 0.392] | 0.376 [0.300, 0.452] | 0.066 [0.031, 0.100] | 0.713 | 0.519 | TMM_LRT 81%; TMM_QL 19% | | |
| limma-voom | | 0.446 [0.359, 0.532] | | | 0.205 | TMM_voom 81%; TMMwsp_voom 19% | | |

**Table S12. Secondary designs.** Group balance, fraction of features with a signal, a single feature increased with an effect size of 10, and balanced or unidirectional signals, at 100 and 200 samples and effect sizes of 1.5 to 2.5. Means with 95% confidence intervals across replicates.

| Level | Tool | Configuration | MCC | Recall | FDR |
|---|---|---|---|---|---|
| ***Group balance (proportion of cases), genus level*** | | | | | |
| 0.3 | LinDA | Pseudo_0.1 | 0.866 [0.776, 0.956] | 0.828 | 0.053 [0.000, 0.109] |
| 0.5 | LinDA | Pseudo_0.1 | 0.919 [0.904, 0.934] | 0.906 | 0.049 [0.000, 0.108] |
| 0.3 | MaAsLin 2 | TMM_LOG | 0.847 [0.785, 0.909] | 0.802 | 0.063 [0.000, 0.138] |
| 0.5 | MaAsLin 2 | TMM_LOG | 0.934 [0.907, 0.960] | 0.922 | 0.035 [0.000, 0.086] |
| 0.3 | LinDA | Imputation | 0.826 [0.740, 0.912] | 0.812 | 0.110 [0.000, 0.226] |
| 0.5 | LinDA | Imputation | 0.828 [0.721, 0.935] | 0.911 | 0.190 [0.073, 0.307] |
| 0.3 | MaAsLin 3 | TSS_PLOG_Median | 0.877 [0.782, 0.972] | 0.812 | 0.023 [0.000, 0.051] |
| 0.5 | MaAsLin 3 | TSS_PLOG_Median | 0.865 [0.754, 0.977] | 0.865 | 0.091 [0.000, 0.198] |
| 0.3 | LinDA | Pseudo_0.5 | 0.872 [0.797, 0.948] | 0.823 | 0.039 [0.000, 0.090] |
| 0.5 | LinDA | Pseudo_0.5 | 0.918 [0.881, 0.956] | 0.901 | 0.043 [0.000, 0.096] |
| 0.3 | MaAsLin 3 | TSS_PLOG | 0.840 [0.744, 0.935] | 0.823 | 0.099 [0.020, 0.178] |
| 0.5 | MaAsLin 3 | TSS_PLOG | 0.850 [0.721, 0.980] | 0.922 | 0.160 [0.010, 0.310] |
| 0.3 | MaAsLin 2 | TSS_LOG * | 0.833 [0.736, 0.930] | 0.818 | 0.103 [0.008, 0.199] |
| 0.5 | MaAsLin 2 | TSS_LOG * | 0.851 [0.733, 0.969] | 0.927 | 0.160 [0.016, 0.303] |
| 0.3 | Wilcoxon | TSS | 0.817 [0.729, 0.904] | 0.812 | 0.123 [0.032, 0.215] |
| 0.5 | Wilcoxon | TSS | 0.838 [0.713, 0.964] | 0.911 | 0.169 [0.018, 0.319] |
| 0.3 | LinDA | Adaptive * | 0.832 [0.740, 0.924] | 0.812 | 0.098 [0.000, 0.213] |
| 0.5 | LinDA | Adaptive * | 0.838 [0.729, 0.946] | 0.911 | 0.173 [0.060, 0.286] |
| 0.3 | LDM | CLR | 0.867 [0.791, 0.943] | 0.818 | 0.044 [0.000, 0.095] |
| 0.5 | LDM | CLR | 0.924 [0.898, 0.951] | 0.922 | 0.054 [0.000, 0.121] |
| 0.3 | LDM | Omni3 | 0.808 [0.734, 0.882] | 0.786 | 0.114 [0.026, 0.202] |
| 0.5 | LDM | Omni3 | 0.815 [0.655, 0.974] | 0.948 | 0.220 [0.042, 0.398] |
| 0.3 | LinDA | Adaptive_NoWinsor | 0.840 [0.734, 0.947] | 0.812 | 0.084 [0.005, 0.164] |
| 0.5 | LinDA | Adaptive_NoWinsor | 0.858 [0.764, 0.953] | 0.901 | 0.135 [0.046, 0.225] |
| 0.3 | MaAsLin 2 | CLR_NONE | 0.855 [0.771, 0.940] | 0.839 | 0.077 [0.000, 0.174] |
| 0.5 | MaAsLin 2 | CLR_NONE | 0.899 [0.814, 0.983] | 0.943 | 0.112 [0.000, 0.237] |
| 0.3 | limma-voom | TMM_voom | 0.798 [0.676, 0.920] | 0.865 | 0.161 [0.046, 0.276] |
| 0.5 | limma-voom | TMM_voom | 0.811 [0.647, 0.975] | 0.906 | 0.192 [0.019, 0.364] |
| 0.3 | LDM | PA | 0.745 [0.583, 0.908] | 0.661 | 0.074 [0.000, 0.157] |
| 0.5 | LDM | PA | 0.830 [0.685, 0.976] | 0.911 | 0.177 [0.036, 0.318] |
| 0.3 | LDM | Arcsin | 0.780 [0.691, 0.868] | 0.781 | 0.160 [0.103, 0.216] |
| 0.5 | LDM | Arcsin | 0.804 [0.642, 0.965] | 0.932 | 0.226 [0.038, 0.414] |
| 0.3 | ZicoSeq | MultiLink | 0.826 [0.687, 0.966] | 0.786 | 0.084 [0.000, 0.193] |
| 0.5 | ZicoSeq | MultiLink | 0.862 [0.743, 0.980] | 0.854 | 0.084 [0.000, 0.271] |
| 0.3 | LinDA | Pseudo_1 | 0.866 [0.798, 0.933] | 0.807 | 0.035 [0.000, 0.083] |
| 0.5 | LinDA | Pseudo_1 | 0.909 [0.875, 0.943] | 0.885 | 0.045 [0.000, 0.100] |
| 0.3 | ZicoSeq | NoPosterior | 0.814 [0.767, 0.860] | 0.750 | 0.072 [0.027, 0.116] |
| 0.5 | ZicoSeq | NoPosterior | 0.900 [0.826, 0.974] | 0.865 | 0.036 [0.000, 0.077] |
| 0.3 | MaAsLin 3 | TSS_LOG_Median * | 0.823 [0.704, 0.941] | 0.719 | 0.011 [0.000, 0.047] |
| 0.5 | MaAsLin 3 | TSS_LOG_Median * | 0.859 [0.803, 0.914] | 0.792 | 0.033 [0.003, 0.062] |
| 0.3 | LDM | Omni * | 0.780 [0.700, 0.859] | 0.781 | 0.162 [0.095, 0.228] |
| 0.5 | LDM | Omni * | 0.803 [0.620, 0.986] | 0.943 | 0.235 [0.018, 0.451] |
| 0.3 | Wilcoxon | CLR | 0.805 [0.735, 0.876] | 0.802 | 0.104 [0.000, 0.220] |
| 0.5 | Wilcoxon | CLR | 0.871 [0.781, 0.961] | 0.901 | 0.117 [0.000, 0.275] |
| 0.3 | limma-voom | TMMwsp_voom | 0.811 [0.691, 0.932] | 0.797 | 0.092 [0.000, 0.218] |
| 0.5 | limma-voom | TMMwsp_voom | 0.869 [0.774, 0.963] | 0.917 | 0.124 [0.000, 0.255] |
| 0.3 | MaAsLin 3 | CLR_NONE | 0.811 [0.666, 0.956] | 0.708 | 0.019 [0.000, 0.058] |

| Level | Tool | Configuration | MCC | Recall | FDR |
|---|---|---|---|---|---|
| 0.5 | MaAsLin 3 | CLR_NONE | 0.844 [0.780, 0.908] | 0.818 | 0.084 [0.047, 0.121] |
| 0.3 | MaAsLin 3 | CLR_NONE_Median | 0.808 [0.685, 0.931] | 0.703 | 0.013 [0.000, 0.037] |
| 0.5 | MaAsLin 3 | CLR_NONE_Median | 0.867 [0.832, 0.902] | 0.812 | 0.039 [0.026, 0.051] |
| 0.3 | MaAsLin 3 | TSS_LOG | 0.775 [0.645, 0.905] | 0.755 | 0.131 [0.000, 0.270] |
| 0.5 | MaAsLin 3 | TSS_LOG | 0.746 [0.637, 0.855] | 0.839 | 0.224 [0.099, 0.349] |
| 0.3 | DESeq2 | LRT_poscounts | 0.776 [0.646, 0.907] | 0.802 | 0.191 [0.078, 0.303] |
| 0.5 | DESeq2 | LRT_poscounts | 0.819 [0.727, 0.911] | 0.870 | 0.186 [0.076, 0.297] |
| 0.3 | DESeq2 | Wald_poscounts * | 0.770 [0.628, 0.911] | 0.807 | 0.211 [0.095, 0.326] |
| 0.5 | DESeq2 | Wald_poscounts * | 0.806 [0.697, 0.914] | 0.854 | 0.195 [0.074, 0.315] |
| 0.3 | ZicoSeq | Default * | 0.782 [0.653, 0.912] | 0.755 | 0.127 [0.000, 0.269] |
| 0.5 | ZicoSeq | Default * | 0.874 [0.763, 0.984] | 0.844 | 0.054 [0.000, 0.204] |
| 0.3 | Wilcoxon | Rarefied | 0.789 [0.688, 0.890] | 0.750 | 0.113 [0.073, 0.153] |
| 0.5 | Wilcoxon | Rarefied | 0.781 [0.673, 0.889] | 0.839 | 0.195 [0.044, 0.346] |
| 0.3 | edgeR | TMM_LRT | 0.743 [0.675, 0.811] | 0.901 | 0.332 [0.218, 0.445] |
| 0.5 | edgeR | TMM_LRT | 0.774 [0.706, 0.841] | 0.958 | 0.328 [0.265, 0.391] |
| 0.3 | LDM | Freq | 0.720 [0.641, 0.799] | 0.646 | 0.129 [0.039, 0.219] |
| 0.5 | LDM | Freq | 0.807 [0.653, 0.961] | 0.911 | 0.209 [0.002, 0.415] |
| 0.3 | ALDEx2 | Wilcox_128 | 0.772 [0.629, 0.915] | 0.651 | 0.000 [0.000, 0.000] |
| 0.5 | ALDEx2 | Wilcox_128 | 0.827 [0.805, 0.850] | 0.724 | 0.006 [0.000, 0.025] |
| 0.3 | edgeR | TMM_QL * | 0.618 [0.575, 0.660] | 0.943 | 0.529 [0.473, 0.584] |
| 0.5 | edgeR | TMM_QL * | 0.654 [0.572, 0.736] | 0.974 | 0.500 [0.414, 0.586] |
| 0.3 | ANCOM-BC2 | BH | 0.824 [0.709, 0.939] | 0.724 | 0.006 [0.000, 0.025] |
| 0.5 | ANCOM-BC2 | BH | 0.868 [0.829, 0.908] | 0.781 | 0.000 [0.000, 0.000] |
| 0.3 | ALDEx2 | Welch_128 * | 0.738 [0.630, 0.847] | 0.609 | 0.008 [0.000, 0.035] |
| 0.5 | ALDEx2 | Welch_128 * | 0.807 [0.783, 0.830] | 0.698 | 0.007 [0.000, 0.029] |
| 0.3 | ALDEx2 | Welch_32 | 0.742 [0.615, 0.869] | 0.615 | 0.008 [0.000, 0.035] |
| 0.5 | ALDEx2 | Welch_32 | 0.804 [0.777, 0.832] | 0.693 | 0.006 [0.000, 0.025] |
| 0.3 | ALDEx2 | Welch_128_IQLR | 0.732 [0.625, 0.839] | 0.599 | 0.008 [0.000, 0.035] |
| 0.5 | ALDEx2 | Welch_128_IQLR | 0.787 [0.756, 0.818] | 0.667 | 0.007 [0.000, 0.029] |
| 0.3 | ANCOM-BC2 | BH_StrucZero | 0.802 [0.696, 0.909] | 0.688 | 0.006 [0.000, 0.025] |
| 0.5 | ANCOM-BC2 | BH_StrucZero | 0.862 [0.831, 0.893] | 0.771 | 0.000 [0.000, 0.000] |
| 0.3 | MaAsLin 2 | TMM_NONE | 0.700 [0.608, 0.792] | 0.562 | 0.047 [0.000, 0.100] |
| 0.5 | MaAsLin 2 | TMM_NONE | 0.776 [0.670, 0.882] | 0.667 | 0.021 [0.000, 0.066] |
| 0.3 | ANCOM-BC2 | Holm * | 0.773 [0.681, 0.864] | 0.646 | 0.008 [0.000, 0.035] |
| 0.5 | ANCOM-BC2 | Holm * | 0.828 [0.796, 0.859] | 0.724 | 0.000 [0.000, 0.000] |
| 0.3 | ANCOM-BC2 | Holm_NoFilter | 0.635 [0.592, 0.677] | 0.792 | 0.416 [0.349, 0.483] |
| 0.5 | ANCOM-BC2 | Holm_NoFilter | 0.642 [0.548, 0.736] | 0.885 | 0.469 [0.390, 0.548] |
| 0.3 | ANCOM-BC2 | BH_NoFilter | 0.510 [0.469, 0.550] | 0.839 | 0.606 [0.588, 0.624] |
| 0.5 | ANCOM-BC2 | BH_NoFilter | 0.502 [0.408, 0.597] | 0.906 | 0.642 [0.581, 0.702] |
| ***Group balance (proportion of cases), OTU level*** | | | | | |
| 0.3 | LinDA | Pseudo_0.1 | 0.799 [0.777, 0.822] | 0.698 | 0.044 [0.029, 0.060] |
| 0.5 | LinDA | Pseudo_0.1 | 0.864 [0.841, 0.886] | 0.820 | 0.059 [0.019, 0.100] |
| 0.3 | MaAsLin 2 | TMM_LOG | 0.797 [0.789, 0.806] | 0.685 | 0.030 [0.000, 0.064] |
| 0.5 | MaAsLin 2 | TMM_LOG | 0.858 [0.837, 0.879] | 0.813 | 0.062 [0.021, 0.102] |
| 0.3 | LinDA | Imputation | 0.742 [0.718, 0.766] | 0.640 | 0.089 [0.053, 0.126] |
| 0.5 | LinDA | Imputation | 0.769 [0.739, 0.800] | 0.743 | 0.149 [0.088, 0.211] |
| 0.3 | MaAsLin 3 | TSS_PLOG_Median | 0.748 [0.725, 0.772] | 0.628 | 0.060 [0.023, 0.097] |
| 0.5 | MaAsLin 3 | TSS_PLOG_Median | 0.799 [0.776, 0.823] | 0.724 | 0.075 [0.026, 0.124] |
| 0.3 | LinDA | Pseudo_0.5 | 0.799 [0.777, 0.820] | 0.695 | 0.042 [0.025, 0.059] |
| 0.5 | LinDA | Pseudo_0.5 | 0.865 [0.838, 0.892] | 0.820 | 0.058 [0.021, 0.094] |
| 0.3 | MaAsLin 3 | TSS_PLOG | 0.794 [0.781, 0.807] | 0.702 | 0.058 [0.023, 0.093] |

| Level | Tool | Configuration | MCC | Recall | FDR |
|---|---|---|---|---|---|
| 0.5 | MaAsLin 3 | TSS_PLOG | 0.834 [0.787, 0.882] | 0.827 | 0.116 [0.044, 0.187] |
| 0.3 | MaAsLin 2 | TSS_LOG * | 0.780 [0.769, 0.792] | 0.701 | 0.082 [0.051, 0.113] |
| 0.5 | MaAsLin 2 | TSS_LOG * | 0.811 [0.762, 0.860] | 0.837 | 0.163 [0.094, 0.231] |
| 0.3 | Wilcoxon | TSS | 0.774 [0.766, 0.783] | 0.685 | 0.075 [0.046, 0.104] |
| 0.5 | Wilcoxon | TSS | 0.816 [0.782, 0.851] | 0.793 | 0.112 [0.054, 0.170] |
| 0.3 | LinDA | Adaptive * | 0.743 [0.720, 0.766] | 0.642 | 0.089 [0.053, 0.125] |
| 0.5 | LinDA | Adaptive * | 0.771 [0.739, 0.804] | 0.746 | 0.149 [0.089, 0.209] |
| 0.3 | LDM | CLR | 0.782 [0.755, 0.810] | 0.667 | 0.040 [0.016, 0.064] |
| 0.5 | LDM | CLR | 0.870 [0.843, 0.898] | 0.825 | 0.054 [0.017, 0.090] |
| 0.3 | LDM | Omni3 | 0.721 [0.705, 0.737] | 0.692 | 0.162 [0.134, 0.191] |
| 0.5 | LDM | Omni3 | 0.658 [0.624, 0.692] | 0.883 | 0.384 [0.338, 0.429] |
| 0.3 | LinDA | Adaptive_NoWinsor | 0.747 [0.722, 0.772] | 0.635 | 0.072 [0.033, 0.111] |
| 0.5 | LinDA | Adaptive_NoWinsor | 0.790 [0.761, 0.819] | 0.736 | 0.106 [0.047, 0.164] |
| 0.3 | MaAsLin 2 | CLR_NONE | 0.823 [0.808, 0.838] | 0.842 | 0.143 [0.102, 0.183] |
| 0.5 | MaAsLin 2 | CLR_NONE | 0.796 [0.751, 0.841] | 0.935 | 0.261 [0.169, 0.352] |
| 0.3 | limma-voom | TMM_voom | 0.478 [0.325, 0.632] | 0.921 | 0.588 [0.378, 0.799] |
| 0.5 | limma-voom | TMM_voom | 0.645 [0.559, 0.731] | 0.923 | 0.409 [0.355, 0.464] |
| 0.3 | LDM | PA | 0.685 [0.665, 0.704] | 0.687 | 0.212 [0.185, 0.240] |
| 0.5 | LDM | PA | 0.654 [0.623, 0.686] | 0.888 | 0.388 [0.339, 0.436] |
| 0.3 | LDM | Arcsin | 0.748 [0.727, 0.769] | 0.702 | 0.134 [0.088, 0.181] |
| 0.5 | LDM | Arcsin | 0.750 [0.705, 0.795] | 0.876 | 0.276 [0.220, 0.333] |
| 0.3 | ZicoSeq | MultiLink | 0.726 [0.701, 0.752] | 0.643 | 0.124 [0.072, 0.176] |
| 0.5 | ZicoSeq | MultiLink | 0.778 [0.726, 0.830] | 0.728 | 0.116 [0.029, 0.204] |
| 0.3 | LinDA | Pseudo_1 | 0.790 [0.770, 0.811] | 0.682 | 0.043 [0.028, 0.058] |
| 0.5 | LinDA | Pseudo_1 | 0.859 [0.830, 0.888] | 0.809 | 0.056 [0.020, 0.093] |
| 0.3 | ZicoSeq | NoPosterior | 0.787 [0.782, 0.792] | 0.681 | 0.047 [0.037, 0.057] |
| 0.5 | ZicoSeq | NoPosterior | 0.852 [0.816, 0.888] | 0.813 | 0.074 [0.021, 0.128] |
| 0.3 | MaAsLin 3 | TSS_LOG_Median * | 0.718 [0.712, 0.723] | 0.557 | 0.024 [0.017, 0.030] |
| 0.5 | MaAsLin 3 | TSS_LOG_Median * | 0.749 [0.737, 0.761] | 0.613 | 0.036 [0.017, 0.054] |
| 0.3 | LDM | Omni * | 0.742 [0.722, 0.762] | 0.686 | 0.124 [0.085, 0.163] |
| 0.5 | LDM | Omni * | 0.671 [0.639, 0.703] | 0.882 | 0.369 [0.327, 0.411] |
| 0.3 | Wilcoxon | CLR | 0.456 [0.427, 0.485] | 0.809 | 0.537 [0.490, 0.584] |
| 0.5 | Wilcoxon | CLR | 0.428 [0.272, 0.584] | 0.867 | 0.597 [0.446, 0.749] |
| 0.3 | limma-voom | TMMwsp_voom | 0.303 [0.236, 0.370] | 0.940 | 0.715 [0.635, 0.794] |
| 0.5 | limma-voom | TMMwsp_voom | 0.255 [0.189, 0.322] | 0.976 | 0.776 [0.694, 0.857] |
| 0.3 | MaAsLin 3 | CLR_NONE | 0.682 [0.671, 0.694] | 0.582 | 0.127 [0.096, 0.158] |
| 0.5 | MaAsLin 3 | CLR_NONE | 0.696 [0.682, 0.710] | 0.669 | 0.194 [0.182, 0.206] |
| 0.3 | MaAsLin 3 | CLR_NONE_Median | 0.730 [0.727, 0.734] | 0.581 | 0.032 [0.021, 0.042] |
| 0.5 | MaAsLin 3 | CLR_NONE_Median | 0.774 [0.762, 0.787] | 0.652 | 0.035 [0.018, 0.052] |
| 0.3 | MaAsLin 3 | TSS_LOG | 0.468 [0.396, 0.541] | 0.623 | 0.456 [0.352, 0.561] |
| 0.5 | MaAsLin 3 | TSS_LOG | 0.464 [0.397, 0.530] | 0.722 | 0.505 [0.412, 0.599] |
| 0.3 | DESeq2 | LRT_poscounts | 0.740 [0.701, 0.778] | 0.681 | 0.143 [0.057, 0.228] |
| 0.5 | DESeq2 | LRT_poscounts | 0.742 [0.703, 0.782] | 0.687 | 0.141 [0.071, 0.212] |
| 0.3 | DESeq2 | Wald_poscounts * | 0.694 [0.651, 0.738] | 0.655 | 0.200 [0.117, 0.284] |
| 0.5 | DESeq2 | Wald_poscounts * | 0.712 [0.662, 0.763] | 0.645 | 0.152 [0.062, 0.241] |
| 0.3 | ZicoSeq | Default * | 0.744 [0.725, 0.764] | 0.649 | 0.096 [0.057, 0.134] |
| 0.5 | ZicoSeq | Default * | 0.788 [0.739, 0.836] | 0.736 | 0.109 [0.034, 0.184] |
| 0.3 | Wilcoxon | Rarefied | 0.680 [0.668, 0.693] | 0.550 | 0.092 [0.076, 0.108] |
| 0.5 | Wilcoxon | Rarefied | 0.678 [0.648, 0.708] | 0.632 | 0.187 [0.145, 0.230] |
| 0.3 | edgeR | TMM_LRT | 0.798 [0.760, 0.836] | 0.876 | 0.229 [0.149, 0.309] |
| 0.5 | edgeR | TMM_LRT | 0.810 [0.766, 0.854] | 0.910 | 0.235 [0.173, 0.298] |

| Level | Tool | Configuration | MCC | Recall | FDR |
|---|---|---|---|---|---|
| 0.3 | LDM | Freq | 0.700 [0.691, 0.710] | 0.557 | 0.060 [0.044, 0.075] |
| 0.5 | LDM | Freq | 0.702 [0.670, 0.735] | 0.862 | 0.328 [0.287, 0.368] |
| 0.3 | ALDEx2 | Wilcox_128 | 0.633 [0.620, 0.646] | 0.437 | 0.000 [0.000, 0.000] |
| 0.5 | ALDEx2 | Wilcox_128 | 0.667 [0.653, 0.681] | 0.479 | 0.000 [0.000, 0.001] |
| 0.3 | edgeR | TMM_QL * | 0.705 [0.662, 0.749] | 0.894 | 0.383 [0.312, 0.454] |
| 0.5 | edgeR | TMM_QL * | 0.722 [0.675, 0.770] | 0.915 | 0.369 [0.309, 0.429] |
| 0.3 | ANCOM-BC2 | BH | 0.776 [0.760, 0.792] | 0.632 | 0.003 [0.000, 0.006] |
| 0.5 | ANCOM-BC2 | BH | 0.787 [0.786, 0.788] | 0.650 | 0.006 [0.000, 0.014] |
| 0.3 | ALDEx2 | Welch_128 * | 0.549 [0.540, 0.558] | 0.355 | 0.000 [0.000, 0.000] |
| 0.5 | ALDEx2 | Welch_128 * | 0.612 [0.604, 0.620] | 0.416 | 0.000 [0.000, 0.000] |
| 0.3 | ALDEx2 | Welch_32 | 0.550 [0.536, 0.563] | 0.355 | 0.000 [0.000, 0.000] |
| 0.5 | ALDEx2 | Welch_32 | 0.614 [0.602, 0.627] | 0.419 | 0.000 [0.000, 0.000] |
| 0.3 | ALDEx2 | Welch_128_IQLR | 0.561 [0.553, 0.568] | 0.365 | 0.000 [0.000, 0.000] |
| 0.5 | ALDEx2 | Welch_128_IQLR | 0.622 [0.609, 0.634] | 0.425 | 0.000 [0.000, 0.000] |
| 0.3 | ANCOM-BC2 | BH_StrucZero | 0.719 [0.708, 0.731] | 0.547 | 0.002 [0.000, 0.006] |
| 0.5 | ANCOM-BC2 | BH_StrucZero | 0.749 [0.738, 0.760] | 0.592 | 0.005 [0.000, 0.013] |
| 0.3 | MaAsLin 2 | TMM_NONE | 0.703 [0.696, 0.710] | 0.528 | 0.009 [0.001, 0.018] |
| 0.5 | MaAsLin 2 | TMM_NONE | 0.754 [0.728, 0.779] | 0.618 | 0.024 [0.000, 0.048] |
| 0.3 | ANCOM-BC2 | Holm * | 0.625 [0.614, 0.636] | 0.441 | 0.000 [0.000, 0.000] |
| 0.5 | ANCOM-BC2 | Holm * | 0.675 [0.661, 0.689] | 0.495 | 0.000 [0.000, 0.000] |
| 0.3 | ANCOM-BC2 | Holm_NoFilter | 0.641 [0.631, 0.651] | 0.680 | 0.323 [0.305, 0.340] |
| 0.5 | ANCOM-BC2 | Holm_NoFilter | 0.538 [0.528, 0.547] | 0.714 | 0.504 [0.494, 0.513] |
| 0.3 | ANCOM-BC2 | BH_NoFilter | 0.349 [0.340, 0.358] | 0.798 | 0.744 [0.737, 0.751] |
| 0.5 | ANCOM-BC2 | BH_NoFilter | 0.239 [0.233, 0.245] | 0.783 | 0.816 [0.815, 0.818] |
| ***Fraction with a signal, genus level*** | | | | | |
| 0.05 | LinDA | Pseudo_0.1 | 0.932 [0.866, 0.997] | 0.896 | 0.017 [0.000, 0.047] |
| 0.1 | LinDA | Pseudo_0.1 | 0.887 [0.840, 0.934] | 0.854 | 0.053 [0.000, 0.146] |
| 0.2 | LinDA | Pseudo_0.1 | 0.904 [0.879, 0.929] | 0.914 | 0.066 [0.020, 0.111] |
| 0.05 | MaAsLin 2 | TMM_LOG | 0.907 [0.799, 1.015] | 0.896 | 0.062 [0.000, 0.132] |
| 0.1 | MaAsLin 2 | TMM_LOG | 0.872 [0.783, 0.961] | 0.870 | 0.093 [0.000, 0.197] |
| 0.2 | MaAsLin 2 | TMM_LOG | 0.874 [0.802, 0.945] | 0.924 | 0.116 [0.000, 0.266] |
| 0.05 | LinDA | Imputation | 0.864 [0.779, 0.949] | 0.906 | 0.135 [0.050, 0.220] |
| 0.1 | LinDA | Imputation | 0.787 [0.757, 0.818] | 0.849 | 0.207 [0.165, 0.249] |
| 0.2 | LinDA | Imputation | 0.795 [0.713, 0.877] | 0.902 | 0.208 [0.107, 0.308] |
| 0.05 | MaAsLin 3 | TSS_PLOG_Median | 0.888 [0.794, 0.983] | 0.885 | 0.074 [0.000, 0.182] |
| 0.1 | MaAsLin 3 | TSS_PLOG_Median | 0.836 [0.810, 0.862] | 0.786 | 0.064 [0.002, 0.126] |
| 0.2 | MaAsLin 3 | TSS_PLOG_Median | 0.866 [0.826, 0.907] | 0.836 | 0.046 [0.002, 0.090] |
| 0.05 | LinDA | Pseudo_0.5 | 0.936 [0.874, 0.999] | 0.896 | 0.008 [0.000, 0.035] |
| 0.1 | LinDA | Pseudo_0.5 | 0.864 [0.801, 0.927] | 0.818 | 0.057 [0.000, 0.119] |
| 0.2 | LinDA | Pseudo_0.5 | 0.894 [0.875, 0.914] | 0.890 | 0.057 [0.012, 0.102] |
| 0.05 | MaAsLin 3 | TSS_PLOG | 0.893 [0.777, 1.008] | 0.917 | 0.090 [0.000, 0.227] |
| 0.1 | MaAsLin 3 | TSS_PLOG | 0.811 [0.786, 0.835] | 0.865 | 0.181 [0.139, 0.224] |
| 0.2 | MaAsLin 3 | TSS_PLOG | 0.836 [0.773, 0.898] | 0.934 | 0.177 [0.084, 0.269] |
| 0.05 | MaAsLin 2 | TSS_LOG * | 0.882 [0.775, 0.990] | 0.917 | 0.103 [0.000, 0.231] |
| 0.1 | MaAsLin 2 | TSS_LOG * | 0.811 [0.793, 0.829] | 0.870 | 0.183 [0.135, 0.231] |
| 0.2 | MaAsLin 2 | TSS_LOG * | 0.815 [0.742, 0.888] | 0.926 | 0.198 [0.092, 0.304] |
| 0.05 | Wilcoxon | TSS | 0.880 [0.794, 0.966] | 0.917 | 0.111 [0.002, 0.220] |
| 0.1 | Wilcoxon | TSS | 0.791 [0.740, 0.842] | 0.849 | 0.190 [0.107, 0.273] |
| 0.2 | Wilcoxon | TSS | 0.798 [0.725, 0.871] | 0.926 | 0.222 [0.127, 0.317] |
| 0.05 | LinDA | Adaptive * | 0.874 [0.801, 0.946] | 0.896 | 0.102 [0.042, 0.162] |
| 0.1 | LinDA | Adaptive * | 0.779 [0.761, 0.796] | 0.818 | 0.187 [0.117, 0.256] |

| Level | Tool | Configuration | MCC | Recall | FDR |
| --- | --- | --- | --- | --- | --- |
| 0.2 | LinDA | Adaptive * | 0.803 [0.724, 0.881] | 0.897 | 0.194 [0.105, 0.283] |
| 0.05 | LDM | CLR | 0.923 [0.821, 1.024] | 0.906 | 0.040 [0.000, 0.135] |
| 0.1 | LDM | CLR | 0.903 [0.871, 0.936] | 0.875 | 0.045 [0.000, 0.098] |
| 0.2 | LDM | CLR | 0.893 [0.888, 0.899] | 0.902 | 0.070 [0.034, 0.106] |
| 0.05 | LDM | Omni3 | 0.888 [0.773, 1.004] | 0.948 | 0.109 [0.000, 0.240] |
| 0.1 | LDM | Omni3 | 0.783 [0.709, 0.857] | 0.885 | 0.225 [0.128, 0.321] |
| 0.2 | LDM | Omni3 | 0.716 [0.563, 0.869] | 0.944 | 0.319 [0.164, 0.474] |
| 0.05 | LinDA | Adaptive_NoWinsor | 0.881 [0.816, 0.947] | 0.896 | 0.087 [0.013, 0.162] |
| 0.1 | LinDA | Adaptive_NoWinsor | 0.805 [0.777, 0.833] | 0.823 | 0.153 [0.080, 0.227] |
| 0.2 | LinDA | Adaptive_NoWinsor | 0.814 [0.726, 0.903] | 0.887 | 0.171 [0.063, 0.280] |
| 0.05 | MaAsLin 2 | CLR_NONE | 0.924 [0.841, 1.006] | 0.896 | 0.031 [0.000, 0.097] |
| 0.1 | MaAsLin 2 | CLR_NONE | 0.879 [0.840, 0.917] | 0.885 | 0.097 [0.050, 0.144] |
| 0.2 | MaAsLin 2 | CLR_NONE | 0.817 [0.762, 0.872] | 0.951 | 0.205 [0.151, 0.259] |
| 0.05 | limma-voom | TMM_voom | 0.826 [0.583, 1.069] | 0.896 | 0.161 [0.000, 0.380] |
| 0.1 | limma-voom | TMM_voom | 0.734 [0.479, 0.989] | 0.859 | 0.250 [0.000, 0.622] |
| 0.2 | limma-voom | TMM_voom | 0.640 [0.381, 0.899] | 0.917 | 0.354 [0.084, 0.625] |
| 0.05 | LDM | PA | 0.858 [0.721, 0.994] | 0.875 | 0.096 [0.000, 0.245] |
| 0.1 | LDM | PA | 0.771 [0.670, 0.872] | 0.818 | 0.191 [0.133, 0.249] |
| 0.2 | LDM | PA | 0.752 [0.634, 0.870] | 0.885 | 0.240 [0.119, 0.361] |
| 0.05 | LDM | Arcsin | 0.881 [0.762, 0.999] | 0.948 | 0.126 [0.000, 0.262] |
| 0.1 | LDM | Arcsin | 0.798 [0.733, 0.863] | 0.891 | 0.205 [0.094, 0.316] |
| 0.2 | LDM | Arcsin | 0.733 [0.597, 0.869] | 0.946 | 0.305 [0.158, 0.453] |
| 0.05 | ZicoSeq | MultiLink | 0.876 [0.772, 0.979] | 0.875 | 0.090 [0.061, 0.120] |
| 0.1 | ZicoSeq | MultiLink | 0.810 [0.664, 0.956] | 0.797 | 0.125 [0.000, 0.321] |
| 0.2 | ZicoSeq | MultiLink | 0.871 [0.846, 0.895] | 0.853 | 0.058 [0.000, 0.124] |
| 0.05 | LinDA | Pseudo_1 | 0.927 [0.865, 0.989] | 0.896 | 0.025 [0.000, 0.052] |
| 0.1 | LinDA | Pseudo_1 | 0.847 [0.775, 0.918] | 0.797 | 0.065 [0.017, 0.113] |
| 0.2 | LinDA | Pseudo_1 | 0.884 [0.874, 0.894] | 0.868 | 0.051 [0.012, 0.090] |
| 0.05 | ZicoSeq | NoPosterior | 0.890 [0.852, 0.928] | 0.844 | 0.039 [0.000, 0.098] |
| 0.1 | ZicoSeq | NoPosterior | 0.868 [0.815, 0.922] | 0.818 | 0.041 [0.000, 0.117] |
| 0.2 | ZicoSeq | NoPosterior | 0.883 [0.846, 0.919] | 0.900 | 0.084 [0.023, 0.145] |
| 0.05 | MaAsLin 3 | TSS_LOG_Median * | 0.862 [0.813, 0.910] | 0.792 | 0.037 [0.000, 0.087] |
| 0.1 | MaAsLin 3 | TSS_LOG_Median * | 0.828 [0.779, 0.876] | 0.750 | 0.050 [0.000, 0.118] |
| 0.2 | MaAsLin 3 | TSS_LOG_Median * | 0.875 [0.824, 0.926] | 0.824 | 0.022 [0.000, 0.047] |
| 0.05 | LDM | Omni * | 0.884 [0.760, 1.007] | 0.948 | 0.119 [0.000, 0.269] |
| 0.1 | LDM | Omni * | 0.791 [0.705, 0.877] | 0.885 | 0.207 [0.080, 0.334] |
| 0.2 | LDM | Omni * | 0.709 [0.551, 0.868] | 0.941 | 0.325 [0.154, 0.495] |
| 0.05 | Wilcoxon | CLR | 0.884 [0.802, 0.965] | 0.844 | 0.045 [0.000, 0.116] |
| 0.1 | Wilcoxon | CLR | 0.826 [0.779, 0.873] | 0.807 | 0.111 [0.040, 0.182] |
| 0.2 | Wilcoxon | CLR | 0.726 [0.653, 0.799] | 0.897 | 0.266 [0.165, 0.366] |
| 0.05 | limma-voom | TMMwsp_voom | 0.854 [0.733, 0.975] | 0.875 | 0.107 [0.000, 0.259] |
| 0.1 | limma-voom | TMMwsp_voom | 0.811 [0.760, 0.862] | 0.865 | 0.156 [0.004, 0.307] |
| 0.2 | limma-voom | TMMwsp_voom | 0.685 [0.528, 0.843] | 0.897 | 0.279 [0.141, 0.417] |
| 0.05 | MaAsLin 3 | CLR_NONE | 0.884 [0.838, 0.930] | 0.823 | 0.031 [0.000, 0.095] |
| 0.1 | MaAsLin 3 | CLR_NONE | 0.805 [0.737, 0.872] | 0.745 | 0.082 [0.022, 0.143] |
| 0.2 | MaAsLin 3 | CLR_NONE | 0.780 [0.703, 0.856] | 0.801 | 0.145 [0.055, 0.235] |
| 0.05 | MaAsLin 3 | CLR_NONE_Median | 0.883 [0.828, 0.937] | 0.802 | 0.010 [0.000, 0.044] |
| 0.1 | MaAsLin 3 | CLR_NONE_Median | 0.829 [0.748, 0.910] | 0.740 | 0.032 [0.000, 0.110] |
| 0.2 | MaAsLin 3 | CLR_NONE_Median | 0.861 [0.805, 0.917] | 0.799 | 0.017 [0.000, 0.046] |
| 0.05 | MaAsLin 3 | TSS_LOG | 0.795 [0.678, 0.911] | 0.823 | 0.154 [0.000, 0.308] |
| 0.1 | MaAsLin 3 | TSS_LOG | 0.724 [0.631, 0.818] | 0.792 | 0.223 [0.087, 0.360] |

| Level | Tool | Configuration | MCC | Recall | FDR |
|---|---|---|---|---|---|
| 0.2 | MaAsLin 3 | TSS_LOG | 0.652 [0.536, 0.767] | 0.846 | 0.332 [0.203, 0.461] |
| 0.05 | DESeq2 | LRT_poscounts | 0.852 [0.803, 0.902] | 0.885 | 0.153 [0.125, 0.180] |
| 0.1 | DESeq2 | LRT_poscounts | 0.799 [0.731, 0.867] | 0.823 | 0.176 [0.085, 0.267] |
| 0.2 | DESeq2 | LRT_poscounts | 0.753 [0.685, 0.821] | 0.848 | 0.228 [0.141, 0.315] |
| 0.05 | DESeq2 | Wald_poscounts * | 0.824 [0.783, 0.864] | 0.844 | 0.163 [0.119, 0.207] |
| 0.1 | DESeq2 | Wald_poscounts * | 0.766 [0.722, 0.809] | 0.786 | 0.196 [0.136, 0.255] |
| 0.2 | DESeq2 | Wald_poscounts * | 0.741 [0.657, 0.824] | 0.833 | 0.235 [0.138, 0.332] |
| 0.05 | ZicoSeq | Default * | 0.812 [0.643, 0.981] | 0.823 | 0.121 [0.107, 0.135] |
| 0.1 | ZicoSeq | Default * | 0.796 [0.709, 0.884] | 0.776 | 0.113 [0.000, 0.237] |
| 0.2 | ZicoSeq | Default * | 0.857 [0.781, 0.933] | 0.850 | 0.075 [0.000, 0.183] |
| 0.05 | Wilcoxon | Rarefied | 0.851 [0.736, 0.967] | 0.865 | 0.103 [0.000, 0.231] |
| 0.1 | Wilcoxon | Rarefied | 0.776 [0.727, 0.825] | 0.781 | 0.151 [0.076, 0.226] |
| 0.2 | Wilcoxon | Rarefied | 0.736 [0.650, 0.823] | 0.826 | 0.222 [0.097, 0.347] |
| 0.05 | edgeR | TMM_LRT | 0.759 [0.690, 0.827] | 0.958 | 0.355 [0.237, 0.474] |
| 0.1 | edgeR | TMM_LRT | 0.747 [0.696, 0.797] | 0.932 | 0.345 [0.276, 0.414] |
| 0.2 | edgeR | TMM_LRT | 0.733 [0.632, 0.835] | 0.963 | 0.337 [0.214, 0.460] |
| 0.05 | LDM | Freq | 0.871 [0.748, 0.995] | 0.917 | 0.122 [0.000, 0.256] |
| 0.1 | LDM | Freq | 0.810 [0.738, 0.883] | 0.859 | 0.165 [0.047, 0.283] |
| 0.2 | LDM | Freq | 0.741 [0.632, 0.850] | 0.909 | 0.277 [0.158, 0.395] |
| 0.05 | ALDEx2 | Wilcox_128 | 0.812 [0.660, 0.965] | 0.708 | 0.000 [0.000, 0.000] |
| 0.1 | ALDEx2 | Wilcox_128 | 0.768 [0.730, 0.805] | 0.630 | 0.013 [0.000, 0.037] |
| 0.2 | ALDEx2 | Wilcox_128 | 0.798 [0.766, 0.831] | 0.718 | 0.032 [0.000, 0.072] |
| 0.05 | edgeR | TMM_QL * | 0.584 [0.556, 0.612] | 0.990 | 0.615 [0.572, 0.657] |
| 0.1 | edgeR | TMM_QL * | 0.627 [0.568, 0.685] | 0.964 | 0.524 [0.442, 0.606] |
| 0.2 | edgeR | TMM_QL * | 0.641 [0.517, 0.765] | 0.978 | 0.448 [0.326, 0.569] |
| 0.05 | ANCOM-BC2 | BH | 0.865 [0.777, 0.952] | 0.771 | 0.000 [0.000, 0.000] |
| 0.1 | ANCOM-BC2 | BH | 0.818 [0.755, 0.881] | 0.703 | 0.006 [0.000, 0.025] |
| 0.2 | ANCOM-BC2 | BH | 0.860 [0.820, 0.900] | 0.782 | 0.000 [0.000, 0.000] |
| 0.05 | ALDEx2 | Welch_128 * | 0.788 [0.632, 0.944] | 0.677 | 0.000 [0.000, 0.000] |
| 0.1 | ALDEx2 | Welch_128 * | 0.741 [0.688, 0.795] | 0.583 | 0.000 [0.000, 0.000] |
| 0.2 | ALDEx2 | Welch_128 * | 0.790 [0.736, 0.845] | 0.696 | 0.021 [0.000, 0.051] |
| 0.05 | ALDEx2 | Welch_32 | 0.776 [0.603, 0.948] | 0.656 | 0.000 [0.000, 0.000] |
| 0.1 | ALDEx2 | Welch_32 | 0.753 [0.723, 0.782] | 0.599 | 0.000 [0.000, 0.000] |
| 0.2 | ALDEx2 | Welch_32 | 0.791 [0.741, 0.840] | 0.699 | 0.022 [0.000, 0.058] |
| 0.05 | ALDEx2 | Welch_128_IQLR | 0.788 [0.632, 0.944] | 0.677 | 0.000 [0.000, 0.000] |
| 0.1 | ALDEx2 | Welch_128_IQLR | 0.737 [0.677, 0.798] | 0.578 | 0.000 [0.000, 0.000] |
| 0.2 | ALDEx2 | Welch_128_IQLR | 0.800 [0.747, 0.854] | 0.696 | 0.003 [0.000, 0.012] |
| 0.05 | ANCOM-BC2 | BH_StrucZero | 0.858 [0.779, 0.937] | 0.760 | 0.000 [0.000, 0.000] |
| 0.1 | ANCOM-BC2 | BH_StrucZero | 0.815 [0.746, 0.884] | 0.698 | 0.006 [0.000, 0.025] |
| 0.2 | ANCOM-BC2 | BH_StrucZero | 0.842 [0.808, 0.877] | 0.755 | 0.000 [0.000, 0.000] |
| 0.05 | MaAsLin 2 | TMM_NONE | 0.727 [0.581, 0.872] | 0.625 | 0.017 [0.000, 0.047] |
| 0.1 | MaAsLin 2 | TMM_NONE | 0.716 [0.693, 0.739] | 0.557 | 0.013 [0.000, 0.054] |
| 0.2 | MaAsLin 2 | TMM_NONE | 0.711 [0.652, 0.770] | 0.674 | 0.126 [0.036, 0.216] |
| 0.05 | ANCOM-BC2 | Holm * | 0.827 [0.676, 0.978] | 0.729 | 0.000 [0.000, 0.000] |
| 0.1 | ANCOM-BC2 | Holm * | 0.783 [0.732, 0.835] | 0.646 | 0.000 [0.000, 0.000] |
| 0.2 | ANCOM-BC2 | Holm * | 0.810 [0.779, 0.841] | 0.708 | 0.000 [0.000, 0.000] |
| 0.05 | ANCOM-BC2 | Holm_NoFilter | 0.535 [0.406, 0.665] | 0.906 | 0.636 [0.482, 0.791] |
| 0.1 | ANCOM-BC2 | Holm_NoFilter | 0.550 [0.441, 0.658] | 0.781 | 0.527 [0.428, 0.626] |
| 0.2 | ANCOM-BC2 | Holm_NoFilter | 0.668 [0.600, 0.736] | 0.826 | 0.329 [0.255, 0.403] |
| 0.05 | ANCOM-BC2 | BH_NoFilter | 0.398 [0.303, 0.492] | 0.906 | 0.775 [0.693, 0.857] |
| 0.1 | ANCOM-BC2 | BH_NoFilter | 0.446 [0.338, 0.555] | 0.812 | 0.660 [0.563, 0.757] |

| Level | Tool | Configuration | MCC | Recall | FDR |
|---|---|---|---|---|---|
| 0.2 | ANCOM-BC2 | BH_NoFilter | 0.566 [0.525, 0.607] | 0.865 | 0.476 [0.441, 0.510] |
| ***Fraction with a signal, OTU level*** | | | | | |
| 0.05 | LinDA | Pseudo_0.1 | 0.849 [0.831, 0.867] | 0.779 | 0.055 [0.042, 0.067] |
| 0.1 | LinDA | Pseudo_0.1 | 0.861 [0.827, 0.895] | 0.809 | 0.054 [0.022, 0.085] |
| 0.2 | LinDA | Pseudo_0.1 | 0.886 [0.878, 0.894] | 0.862 | 0.043 [0.038, 0.049] |
| 0.05 | MaAsLin 2 | TMM_LOG | 0.845 [0.830, 0.860] | 0.764 | 0.045 [0.032, 0.057] |
| 0.1 | MaAsLin 2 | TMM_LOG | 0.852 [0.834, 0.870] | 0.804 | 0.062 [0.010, 0.114] |
| 0.2 | MaAsLin 2 | TMM_LOG | 0.873 [0.859, 0.887] | 0.860 | 0.060 [0.036, 0.084] |
| 0.05 | LinDA | Imputation | 0.791 [0.771, 0.811] | 0.714 | 0.099 [0.083, 0.115] |
| 0.1 | LinDA | Imputation | 0.773 [0.743, 0.803] | 0.741 | 0.142 [0.090, 0.195] |
| 0.2 | LinDA | Imputation | 0.726 [0.692, 0.759] | 0.751 | 0.186 [0.155, 0.218] |
| 0.05 | MaAsLin 3 | TSS_PLOG_Median | 0.803 [0.783, 0.822] | 0.702 | 0.059 [0.052, 0.067] |
| 0.1 | MaAsLin 3 | TSS_PLOG_Median | 0.795 [0.770, 0.820] | 0.717 | 0.076 [0.031, 0.122] |
| 0.2 | MaAsLin 3 | TSS_PLOG_Median | 0.776 [0.760, 0.792] | 0.724 | 0.082 [0.070, 0.095] |
| 0.05 | LinDA | Pseudo_0.5 | 0.851 [0.832, 0.869] | 0.777 | 0.049 [0.038, 0.060] |
| 0.1 | LinDA | Pseudo_0.5 | 0.863 [0.834, 0.893] | 0.812 | 0.053 [0.024, 0.082] |
| 0.2 | LinDA | Pseudo_0.5 | 0.890 [0.885, 0.895] | 0.866 | 0.041 [0.036, 0.045] |
| 0.05 | MaAsLin 3 | TSS_PLOG | 0.832 [0.815, 0.849] | 0.771 | 0.078 [0.051, 0.106] |
| 0.1 | MaAsLin 3 | TSS_PLOG | 0.836 [0.806, 0.865] | 0.819 | 0.106 [0.032, 0.180] |
| 0.2 | MaAsLin 3 | TSS_PLOG | 0.825 [0.799, 0.852] | 0.879 | 0.147 [0.108, 0.186] |
| 0.05 | MaAsLin 2 | TSS_LOG * | 0.822 [0.805, 0.838] | 0.770 | 0.096 [0.062, 0.129] |
| 0.1 | MaAsLin 2 | TSS_LOG * | 0.814 [0.768, 0.860] | 0.827 | 0.149 [0.058, 0.240] |
| 0.2 | MaAsLin 2 | TSS_LOG * | 0.780 [0.764, 0.795] | 0.890 | 0.216 [0.193, 0.238] |
| 0.05 | Wilcoxon | TSS | 0.820 [0.807, 0.832] | 0.748 | 0.075 [0.051, 0.099] |
| 0.1 | Wilcoxon | TSS | 0.814 [0.787, 0.841] | 0.789 | 0.114 [0.047, 0.181] |
| 0.2 | Wilcoxon | TSS | 0.819 [0.809, 0.828] | 0.841 | 0.124 [0.103, 0.145] |
| 0.05 | LinDA | Adaptive * | 0.795 [0.774, 0.815] | 0.719 | 0.096 [0.074, 0.118] |
| 0.1 | LinDA | Adaptive * | 0.776 [0.748, 0.804] | 0.742 | 0.139 [0.089, 0.190] |
| 0.2 | LinDA | Adaptive * | 0.726 [0.692, 0.759] | 0.751 | 0.186 [0.155, 0.218] |
| 0.05 | LDM | CLR | 0.852 [0.824, 0.881] | 0.784 | 0.054 [0.041, 0.068] |
| 0.1 | LDM | CLR | 0.869 [0.836, 0.903] | 0.822 | 0.052 [0.020, 0.085] |
| 0.2 | LDM | CLR | 0.894 [0.881, 0.907] | 0.868 | 0.037 [0.032, 0.042] |
| 0.05 | LDM | Omni3 | 0.774 [0.757, 0.792] | 0.807 | 0.206 [0.166, 0.246] |
| 0.1 | LDM | Omni3 | 0.663 [0.627, 0.699] | 0.874 | 0.378 [0.333, 0.423] |
| 0.2 | LDM | Omni3 | 0.506 [0.475, 0.536] | 0.942 | 0.510 [0.486, 0.534] |
| 0.05 | LinDA | Adaptive_NoWinsor | 0.805 [0.790, 0.820] | 0.721 | 0.078 [0.052, 0.103] |
| 0.1 | LinDA | Adaptive_NoWinsor | 0.791 [0.763, 0.819] | 0.730 | 0.099 [0.048, 0.149] |
| 0.2 | LinDA | Adaptive_NoWinsor | 0.756 [0.736, 0.775] | 0.736 | 0.126 [0.109, 0.142] |
| 0.05 | MaAsLin 2 | CLR_NONE | 0.851 [0.842, 0.861] | 0.832 | 0.103 [0.053, 0.154] |
| 0.1 | MaAsLin 2 | CLR_NONE | 0.792 [0.758, 0.826] | 0.927 | 0.257 [0.237, 0.276] |
| 0.2 | MaAsLin 2 | CLR_NONE | 0.615 [0.575, 0.656] | 0.992 | 0.461 [0.411, 0.511] |
| 0.05 | limma-voom | TMM_voom | 0.665 [0.561, 0.770] | 0.808 | 0.334 [0.198, 0.469] |
| 0.1 | limma-voom | TMM_voom | 0.639 [0.604, 0.674] | 0.951 | 0.446 [0.375, 0.517] |
| 0.2 | limma-voom | TMM_voom | 0.410 [0.207, 0.612] | 0.963 | 0.588 [0.392, 0.784] |
| 0.05 | LDM | PA | 0.772 [0.754, 0.789] | 0.805 | 0.208 [0.167, 0.248] |
| 0.1 | LDM | PA | 0.658 [0.616, 0.700] | 0.878 | 0.385 [0.334, 0.436] |
| 0.2 | LDM | PA | 0.498 [0.471, 0.526] | 0.945 | 0.517 [0.496, 0.537] |
| 0.05 | LDM | Arcsin | 0.810 [0.803, 0.816] | 0.809 | 0.155 [0.129, 0.180] |
| 0.1 | LDM | Arcsin | 0.756 [0.695, 0.817] | 0.865 | 0.265 [0.171, 0.358] |
| 0.2 | LDM | Arcsin | 0.647 [0.624, 0.670] | 0.930 | 0.384 [0.364, 0.404] |
| 0.05 | ZicoSeq | MultiLink | 0.792 [0.776, 0.808] | 0.692 | 0.068 [0.043, 0.093] |

| Level | Tool | Configuration | MCC | Recall | FDR |
|---|---|---|---|---|---|
| 0.1 | ZicoSeq | MultiLink | 0.785 [0.763, 0.807] | 0.717 | 0.096 [0.053, 0.140] |
| 0.2 | ZicoSeq | MultiLink | 0.500 [0.490, 0.509] | 0.862 | 0.429 [0.398, 0.459] |
| 0.05 | LinDA | Pseudo_1 | 0.846 [0.828, 0.864] | 0.766 | 0.047 [0.029, 0.064] |
| 0.1 | LinDA | Pseudo_1 | 0.858 [0.827, 0.889] | 0.800 | 0.050 [0.015, 0.084] |
| 0.2 | LinDA | Pseudo_1 | 0.886 [0.879, 0.893] | 0.860 | 0.041 [0.036, 0.046] |
| 0.05 | ZicoSeq | NoPosterior | 0.840 [0.833, 0.847] | 0.764 | 0.056 [0.051, 0.061] |
| 0.1 | ZicoSeq | NoPosterior | 0.849 [0.843, 0.855] | 0.799 | 0.066 [0.037, 0.095] |
| 0.2 | ZicoSeq | NoPosterior | 0.718 [0.679, 0.757] | 0.863 | 0.243 [0.180, 0.307] |
| 0.05 | MaAsLin 3 | TSS_LOG_Median * | 0.759 [0.741, 0.776] | 0.614 | 0.036 [0.028, 0.045] |
| 0.1 | MaAsLin 3 | TSS_LOG_Median * | 0.746 [0.733, 0.759] | 0.608 | 0.035 [0.018, 0.053] |
| 0.2 | MaAsLin 3 | TSS_LOG_Median * | 0.721 [0.712, 0.730] | 0.597 | 0.029 [0.024, 0.034] |
| 0.05 | LDM | Omni * | 0.783 [0.764, 0.802] | 0.807 | 0.195 [0.152, 0.237] |
| 0.1 | LDM | Omni * | 0.678 [0.638, 0.718] | 0.872 | 0.362 [0.310, 0.415] |
| 0.2 | LDM | Omni * | 0.524 [0.494, 0.554] | 0.941 | 0.496 [0.470, 0.521] |
| 0.05 | Wilcoxon | CLR | 0.628 [0.524, 0.731] | 0.719 | 0.292 [0.135, 0.448] |
| 0.1 | Wilcoxon | CLR | 0.412 [0.300, 0.524] | 0.870 | 0.617 [0.457, 0.777] |
| 0.2 | Wilcoxon | CLR | 0.142 [0.068, 0.215] | 0.908 | 0.759 [0.726, 0.793] |
| 0.05 | limma-voom | TMMwsp_voom | 0.481 [0.359, 0.603] | 0.881 | 0.561 [0.393, 0.729] |
| 0.1 | limma-voom | TMMwsp_voom | 0.255 [0.157, 0.352] | 0.960 | 0.764 [0.647, 0.881] |
| 0.2 | limma-voom | TMMwsp_voom | 0.040 [−0.029, 0.109] | 0.965 | 0.789 [0.770, 0.809] |
| 0.05 | MaAsLin 3 | CLR_NONE | 0.748 [0.730, 0.767] | 0.625 | 0.075 [0.051, 0.098] |
| 0.1 | MaAsLin 3 | CLR_NONE | 0.702 [0.684, 0.720] | 0.662 | 0.180 [0.155, 0.204] |
| 0.2 | MaAsLin 3 | CLR_NONE | 0.551 [0.540, 0.562] | 0.742 | 0.383 [0.365, 0.400] |
| 0.05 | MaAsLin 3 | CLR_NONE_Median | 0.769 [0.752, 0.785] | 0.627 | 0.033 [0.022, 0.045] |
| 0.1 | MaAsLin 3 | CLR_NONE_Median | 0.775 [0.767, 0.784] | 0.651 | 0.032 [0.013, 0.051] |
| 0.2 | MaAsLin 3 | CLR_NONE_Median | 0.783 [0.775, 0.792] | 0.692 | 0.030 [0.023, 0.038] |
| 0.05 | MaAsLin 3 | TSS_LOG | 0.612 [0.537, 0.686] | 0.651 | 0.316 [0.189, 0.442] |
| 0.1 | MaAsLin 3 | TSS_LOG | 0.474 [0.420, 0.529] | 0.723 | 0.491 [0.427, 0.555] |
| 0.2 | MaAsLin 3 | TSS_LOG | 0.300 [0.263, 0.338] | 0.810 | 0.623 [0.569, 0.677] |
| 0.05 | DESeq2 | LRT_poscounts | 0.740 [0.724, 0.755] | 0.655 | 0.135 [0.124, 0.146] |
| 0.1 | DESeq2 | LRT_poscounts | 0.746 [0.720, 0.771] | 0.684 | 0.134 [0.068, 0.199] |
| 0.2 | DESeq2 | LRT_poscounts | 0.694 [0.669, 0.719] | 0.743 | 0.225 [0.183, 0.267] |
| 0.05 | DESeq2 | Wald_poscounts * | 0.713 [0.703, 0.722] | 0.620 | 0.148 [0.132, 0.163] |
| 0.1 | DESeq2 | Wald_poscounts * | 0.716 [0.696, 0.737] | 0.644 | 0.143 [0.083, 0.203] |
| 0.2 | DESeq2 | Wald_poscounts * | 0.663 [0.639, 0.688] | 0.699 | 0.232 [0.191, 0.273] |
| 0.05 | ZicoSeq | Default * | 0.801 [0.786, 0.817] | 0.699 | 0.059 [0.036, 0.082] |
| 0.1 | ZicoSeq | Default * | 0.792 [0.776, 0.808] | 0.725 | 0.091 [0.057, 0.125] |
| 0.2 | ZicoSeq | Default * | 0.513 [0.498, 0.527] | 0.863 | 0.417 [0.379, 0.456] |
| 0.05 | Wilcoxon | Rarefied | 0.723 [0.712, 0.733] | 0.598 | 0.092 [0.068, 0.117] |
| 0.1 | Wilcoxon | Rarefied | 0.687 [0.658, 0.715] | 0.628 | 0.171 [0.124, 0.218] |
| 0.2 | Wilcoxon | Rarefied | 0.617 [0.611, 0.623] | 0.678 | 0.264 [0.257, 0.272] |
| 0.05 | edgeR | TMM_LRT | 0.755 [0.726, 0.784] | 0.881 | 0.323 [0.265, 0.382] |
| 0.1 | edgeR | TMM_LRT | 0.809 [0.789, 0.829] | 0.912 | 0.240 [0.201, 0.280] |
| 0.2 | edgeR | TMM_LRT | 0.818 [0.784, 0.853] | 0.940 | 0.212 [0.156, 0.268] |
| 0.05 | LDM | Freq | 0.784 [0.769, 0.799] | 0.792 | 0.181 [0.137, 0.225] |
| 0.1 | LDM | Freq | 0.710 [0.676, 0.743] | 0.853 | 0.316 [0.266, 0.367] |
| 0.2 | LDM | Freq | 0.600 [0.583, 0.618] | 0.903 | 0.421 [0.402, 0.440] |
| 0.05 | ALDEx2 | Wilcox_128 | 0.664 [0.659, 0.669] | 0.463 | 0.000 [0.000, 0.000] |
| 0.1 | ALDEx2 | Wilcox_128 | 0.668 [0.657, 0.679] | 0.479 | 0.001 [0.000, 0.003] |
| 0.2 | ALDEx2 | Wilcox_128 | 0.673 [0.670, 0.676] | 0.515 | 0.001 [0.000, 0.003] |
| 0.05 | edgeR | TMM_QL * | 0.631 [0.580, 0.682] | 0.890 | 0.514 [0.436, 0.592] |

| Level | Tool | Configuration | MCC | Recall | FDR |
|---|---|---|---|---|---|
| 0.1 | edgeR | TMM_QL * | 0.717 [0.695, 0.739] | 0.916 | 0.380 [0.345, 0.416] |
| 0.2 | edgeR | TMM_QL * | 0.752 [0.714, 0.791] | 0.942 | 0.301 [0.248, 0.354] |
| 0.05 | ANCOM-BC2 | BH | 0.779 [0.762, 0.795] | 0.623 | 0.004 [0.001, 0.006] |
| 0.1 | ANCOM-BC2 | BH | 0.785 [0.783, 0.787] | 0.645 | 0.004 [0.000, 0.016] |
| 0.2 | ANCOM-BC2 | BH | 0.791 [0.785, 0.797] | 0.679 | 0.002 [0.000, 0.004] |
| 0.05 | ALDEx2 | Welch_128 * | 0.604 [0.591, 0.617] | 0.399 | 0.000 [0.000, 0.000] |
| 0.1 | ALDEx2 | Welch_128 * | 0.611 [0.602, 0.621] | 0.416 | 0.000 [0.000, 0.000] |
| 0.2 | ALDEx2 | Welch_128 * | 0.620 [0.613, 0.627] | 0.450 | 0.000 [0.000, 0.001] |
| 0.05 | ALDEx2 | Welch_32 | 0.608 [0.592, 0.625] | 0.404 | 0.000 [0.000, 0.000] |
| 0.1 | ALDEx2 | Welch_32 | 0.614 [0.605, 0.623] | 0.418 | 0.000 [0.000, 0.000] |
| 0.2 | ALDEx2 | Welch_32 | 0.619 [0.614, 0.624] | 0.449 | 0.000 [0.000, 0.001] |
| 0.05 | ALDEx2 | Welch_128_IQLR | 0.608 [0.593, 0.624] | 0.403 | 0.000 [0.000, 0.000] |
| 0.1 | ALDEx2 | Welch_128_IQLR | 0.620 [0.611, 0.629] | 0.423 | 0.000 [0.000, 0.000] |
| 0.2 | ALDEx2 | Welch_128_IQLR | 0.626 [0.622, 0.631] | 0.454 | 0.000 [0.000, 0.000] |
| 0.05 | ANCOM-BC2 | BH_StrucZero | 0.750 [0.743, 0.757] | 0.578 | 0.004 [0.002, 0.005] |
| 0.1 | ANCOM-BC2 | BH_StrucZero | 0.749 [0.739, 0.758] | 0.591 | 0.005 [0.000, 0.017] |
| 0.2 | ANCOM-BC2 | BH_StrucZero | 0.745 [0.737, 0.753] | 0.611 | 0.002 [0.000, 0.004] |
| 0.05 | MaAsLin 2 | TMM_NONE | 0.719 [0.707, 0.731] | 0.546 | 0.011 [0.000, 0.023] |
| 0.1 | MaAsLin 2 | TMM_NONE | 0.748 [0.724, 0.772] | 0.611 | 0.027 [0.007, 0.046] |
| 0.2 | MaAsLin 2 | TMM_NONE | 0.775 [0.758, 0.793] | 0.700 | 0.049 [0.025, 0.073] |
| 0.05 | ANCOM-BC2 | Holm * | 0.689 [0.681, 0.696] | 0.500 | 0.000 [0.000, 0.000] |
| 0.1 | ANCOM-BC2 | Holm * | 0.678 [0.669, 0.686] | 0.497 | 0.000 [0.000, 0.000] |
| 0.2 | ANCOM-BC2 | Holm * | 0.662 [0.659, 0.665] | 0.503 | 0.000 [0.000, 0.000] |
| 0.05 | ANCOM-BC2 | Holm_NoFilter | 0.433 [0.426, 0.441] | 0.699 | 0.675 [0.664, 0.687] |
| 0.1 | ANCOM-BC2 | Holm_NoFilter | 0.537 [0.509, 0.566] | 0.700 | 0.494 [0.462, 0.526] |
| 0.2 | ANCOM-BC2 | Holm_NoFilter | 0.615 [0.601, 0.628] | 0.713 | 0.320 [0.301, 0.340] |
| 0.05 | ANCOM-BC2 | BH_NoFilter | 0.176 [0.166, 0.187] | 0.770 | 0.902 [0.899, 0.905] |
| 0.1 | ANCOM-BC2 | BH_NoFilter | 0.244 [0.226, 0.261] | 0.774 | 0.812 [0.803, 0.822] |
| 0.2 | ANCOM-BC2 | BH_NoFilter | 0.311 [0.297, 0.326] | 0.783 | 0.668 [0.659, 0.677] |
| ***Single extreme change, genus level*** | | | | | |
| single feature, log fold-change 10 | LinDA | Pseudo_0.1 | 0.872 [0.727, 1.017] | 1.000 | 0.208 [0.000, 0.438] |
| single feature, log fold-change 10 | MaAsLin 2 | TMM_LOG | 1.000 [1.000, 1.000] | 1.000 | 0.000 [0.000, 0.000] |
| single feature, log fold-change 10 | LinDA | Imputation | 0.124 [0.104, 0.144] | 1.000 | 0.972 [0.966, 0.979] |
| single feature, log fold-change 10 | MaAsLin 3 | TSS_PLOG_Median | 0.189 [0.165, 0.214] | 1.000 | 0.951 [0.941, 0.961] |
| single feature, log fold-change 10 | LinDA | Pseudo_0.5 | 0.909 [0.737, 1.082] | 1.000 | 0.146 [0.000, 0.419] |
| single feature, log fold-change 10 | MaAsLin 3 | TSS_PLOG | 0.148 [0.118, 0.178] | 1.000 | 0.965 [0.955, 0.975] |
| single feature, log fold-change 10 | MaAsLin 2 | TSS_LOG * | 0.115 [0.094, 0.136] | 1.000 | 0.974 [0.968, 0.980] |
| single feature, log fold-change 10 | Wilcoxon | TSS | 0.160 [0.155, 0.165] | 1.000 | 0.963 [0.961, 0.964] |
| single feature, log fold-change 10 | LinDA | Adaptive * | 0.124 [0.104, 0.144] | 1.000 | 0.972 [0.966, 0.979] |
| single feature, log fold-change 10 | LDM | CLR | 1.000 [1.000, 1.000] | 1.000 | 0.000 [0.000, 0.000] |
| single feature, log fold-change 10 | LDM | Omni3 | 0.030 [0.020, 0.041] | 1.000 | 0.986 [0.985, 0.988] |
| single feature, log fold-change 10 | LinDA | Adaptive_NoWinsor | 0.152 [0.138, 0.166] | 1.000 | 0.965 [0.960, 0.969] |
| single feature, log fold-change 10 | MaAsLin 2 | CLR_NONE | 0.666 [0.190, 1.142] | 1.000 | 0.428 [0.000, 1.021] |
| single feature, log fold-change 10 | limma-voom | TMM_voom | 1.000 [1.000, 1.000] | 1.000 | 0.000 [0.000, 0.000] |
| single feature, log fold-change 10 | LDM | PA | 0.037 [0.024, 0.050] | 1.000 | 0.986 [0.984, 0.988] |
| single feature, log fold-change 10 | LDM | Arcsin | 0.038 [0.021, 0.055] | 1.000 | 0.985 [0.982, 0.988] |
| single feature, log fold-change 10 | ZicoSeq | MultiLink | 0.445 [0.111, 0.778] | 1.000 | 0.674 [0.278, 1.070] |
| single feature, log fold-change 10 | LinDA | Pseudo_1 | 0.909 [0.737, 1.082] | 1.000 | 0.146 [0.000, 0.419] |
| single feature, log fold-change 10 | ZicoSeq | NoPosterior | 0.802 [0.517, 1.086] | 1.000 | 0.308 [0.000, 0.735] |
| single feature, log fold-change 10 | MaAsLin 3 | TSS_LOG_Median * | 1.000 [1.000, 1.000] | 1.000 | 0.000 [0.000, 0.000] |
| single feature, log fold-change 10 | LDM | Omni * | 0.029 [0.020, 0.039] | 1.000 | 0.987 [0.985, 0.988] |

| Level | Tool | Configuration | MCC | Recall | FDR |
|---|---|---|---|---|---|
| single feature, log fold-change 10 | Wilcoxon | CLR | 0.663 [0.291, 1.034] | 1.000 | 0.417 [0.000, 0.895] |
| single feature, log fold-change 10 | limma-voom | TMMwsp_voom | 0.892 [0.677, 1.108] | 1.000 | 0.163 [0.000, 0.477] |
| single feature, log fold-change 10 | MaAsLin 3 | CLR_NONE | 0.808 [0.542, 1.074] | 1.000 | 0.302 [0.000, 0.712] |
| single feature, log fold-change 10 | MaAsLin 3 | CLR_NONE_Median | 1.000 [1.000, 1.000] | 1.000 | 0.000 [0.000, 0.000] |
| single feature, log fold-change 10 | MaAsLin 3 | TSS_LOG | 0.027 [0.021, 0.034] | 1.000 | 0.987 [0.986, 0.988] |
| single feature, log fold-change 10 | DESeq2 | LRT_poscounts | 0.872 [0.727, 1.017] | 1.000 | 0.208 [0.000, 0.438] |
| single feature, log fold-change 10 | DESeq2 | Wald_poscounts * | 0.835 [0.597, 1.073] | 1.000 | 0.271 [0.000, 0.652] |
| single feature, log fold-change 10 | ZicoSeq | Default * | 0.547 [0.259, 0.836] | 1.000 | 0.566 [0.205, 0.927] |
| single feature, log fold-change 10 | Wilcoxon | Rarefied | 0.097 [0.088, 0.106] | 1.000 | 0.978 [0.976, 0.981] |
| single feature, log fold-change 10 | edgeR | TMM_LRT | 0.583 [0.385, 0.781] | 1.000 | 0.613 [0.345, 0.882] |
| single feature, log fold-change 10 | LDM | Freq | 0.063 [0.050, 0.075] | 1.000 | 0.983 [0.981, 0.986] |
| single feature, log fold-change 10 | ALDEx2 | Wilcox_128 | 1.000 [1.000, 1.000] | 1.000 | 0.000 [0.000, 0.000] |
| single feature, log fold-change 10 | edgeR | TMM_QL * | 0.377 [0.209, 0.545] | 1.000 | 0.832 [0.668, 0.995] |
| single feature, log fold-change 10 | ANCOM-BC2 | BH | 1.000 [1.000, 1.000] | 1.000 | 0.000 [0.000, 0.000] |
| single feature, log fold-change 10 | ALDEx2 | Welch_128 * | 1.000 [1.000, 1.000] | 1.000 | 0.000 [0.000, 0.000] |
| single feature, log fold-change 10 | ALDEx2 | Welch_32 | 1.000 [1.000, 1.000] | 1.000 | 0.000 [0.000, 0.000] |
| single feature, log fold-change 10 | ALDEx2 | Welch_128_IQLR | 1.000 [1.000, 1.000] | 1.000 | 0.000 [0.000, 0.000] |
| single feature, log fold-change 10 | ANCOM-BC2 | BH_StrucZero | 1.000 [1.000, 1.000] | 1.000 | 0.000 [0.000, 0.000] |
| single feature, log fold-change 10 | MaAsLin 2 | TMM_NONE | 0.750 [0.291, 1.209] | 0.750 | 0.000 [0.000, 0.000] |
| single feature, log fold-change 10 | ANCOM-BC2 | Holm * | 1.000 [1.000, 1.000] | 1.000 | 0.000 [0.000, 0.000] |
| single feature, log fold-change 10 | ANCOM-BC2 | Holm_NoFilter | 0.288 [0.243, 0.334] | 1.000 | 0.904 [0.877, 0.930] |
| single feature, log fold-change 10 | ANCOM-BC2 | BH_NoFilter | 0.201 [0.156, 0.246] | 1.000 | 0.947 [0.927, 0.966] |
| ***Single extreme change, OTU level*** | | | | | |
| single feature, log fold-change 10 | LinDA | Pseudo_0.1 | 0.811 [0.550, 1.073] | 1.000 | 0.302 [0.000, 0.712] |
| single feature, log fold-change 10 | MaAsLin 2 | TMM_LOG | 1.000 [1.000, 1.000] | 1.000 | 0.000 [0.000, 0.000] |
| single feature, log fold-change 10 | LinDA | Imputation | 0.053 [0.044, 0.061] | 1.000 | 0.997 [0.996, 0.998] |
| single feature, log fold-change 10 | MaAsLin 3 | TSS_PLOG_Median | 0.157 [0.047, 0.267] | 1.000 | 0.967 [0.917, 1.018] |
| single feature, log fold-change 10 | LinDA | Pseudo_0.5 | 0.874 [0.732, 1.016] | 1.000 | 0.208 [0.000, 0.438] |
| single feature, log fold-change 10 | MaAsLin 3 | TSS_PLOG | 0.064 [0.016, 0.112] | 1.000 | 0.994 [0.985, 1.003] |
| single feature, log fold-change 10 | MaAsLin 2 | TSS_LOG * | 0.038 [0.018, 0.057] | 1.000 | 0.998 [0.996, 1.000] |
| single feature, log fold-change 10 | Wilcoxon | TSS | 0.080 [0.062, 0.097] | 1.000 | 0.993 [0.989, 0.996] |
| single feature, log fold-change 10 | LinDA | Adaptive * | 0.053 [0.044, 0.061] | 1.000 | 0.997 [0.996, 0.998] |
| single feature, log fold-change 10 | LDM | CLR | 0.000 [0.000, 0.000] | 0.000 | 0.000 [0.000, 0.000] |
| single feature, log fold-change 10 | LDM | Omni3 | 0.002 [0.001, 0.003] | 1.000 | 1.000 [1.000, 1.000] |
| single feature, log fold-change 10 | LinDA | Adaptive_NoWinsor | 0.092 [0.046, 0.137] | 1.000 | 0.990 [0.977, 1.002] |
| single feature, log fold-change 10 | MaAsLin 2 | CLR_NONE | 0.963 [0.847, 1.080] | 1.000 | 0.062 [0.000, 0.261] |
| single feature, log fold-change 10 | limma-voom | TMM_voom | 0.642 [−0.021, 1.304] | 1.000 | 0.455 [0.000, 1.292] |
| single feature, log fold-change 10 | LDM | PA | 0.002 [0.001, 0.003] | 1.000 | 1.000 [1.000, 1.000] |
| single feature, log fold-change 10 | LDM | Arcsin | 0.007 [0.003, 0.010] | 1.000 | 1.000 [1.000, 1.000] |
| single feature, log fold-change 10 | ZicoSeq | MultiLink | 0.104 [0.074, 0.134] | 1.000 | 0.988 [0.982, 0.995] |
| single feature, log fold-change 10 | LinDA | Pseudo_1 | 0.947 [0.779, 1.115] | 1.000 | 0.083 [0.000, 0.349] |
| single feature, log fold-change 10 | ZicoSeq | NoPosterior | 0.567 [0.466, 0.668] | 1.000 | 0.637 [0.436, 0.838] |
| single feature, log fold-change 10 | MaAsLin 3 | TSS_LOG_Median * | 1.000 [1.000, 1.000] | 1.000 | 0.000 [0.000, 0.000] |
| single feature, log fold-change 10 | LDM | Omni * | 0.003 [0.002, 0.004] | 1.000 | 1.000 [1.000, 1.000] |
| single feature, log fold-change 10 | Wilcoxon | CLR | 0.910 [0.622, 1.197] | 1.000 | 0.115 [0.000, 0.483] |
| single feature, log fold-change 10 | limma-voom | TMMwsp_voom | 1.000 [1.000, 1.000] | 1.000 | 0.000 [0.000, 0.000] |
| single feature, log fold-change 10 | MaAsLin 3 | CLR_NONE | 0.963 [0.847, 1.080] | 1.000 | 0.062 [0.000, 0.261] |
| single feature, log fold-change 10 | MaAsLin 3 | CLR_NONE_Median | 0.963 [0.847, 1.080] | 1.000 | 0.062 [0.000, 0.261] |
| single feature, log fold-change 10 | MaAsLin 3 | TSS_LOG | 0.003 [0.001, 0.004] | 1.000 | 1.000 [1.000, 1.000] |
| single feature, log fold-change 10 | DESeq2 | LRT_poscounts | 0.894 [0.700, 1.089] | 1.000 | 0.167 [0.000, 0.473] |
| single feature, log fold-change 10 | DESeq2 | Wald_poscounts * | 0.805 [0.570, 1.039] | 1.000 | 0.312 [0.000, 0.693] |

| Level | Tool | Configuration | MCC | Recall | FDR |
|---|---|---|---|---|---|
| single feature, log fold-change 10 | ZicoSeq | Default * | 0.103 [0.080, 0.126] | 1.000 | 0.989 [0.984, 0.994] |
| single feature, log fold-change 10 | Wilcoxon | Rarefied | 0.028 [0.022, 0.034] | 1.000 | 0.999 [0.998, 0.999] |
| single feature, log fold-change 10 | edgeR | TMM_LRT | 0.288 [0.218, 0.359] | 1.000 | 0.902 [0.857, 0.947] |
| single feature, log fold-change 10 | LDM | Freq | 0.009 [0.007, 0.011] | 1.000 | 1.000 [0.999, 1.000] |
| single feature, log fold-change 10 | ALDEx2 | Wilcox_128 | 1.000 [1.000, 1.000] | 1.000 | 0.000 [0.000, 0.000] |
| single feature, log fold-change 10 | edgeR | TMM_QL * | 0.119 [0.073, 0.165] | 1.000 | 0.984 [0.970, 0.998] |
| single feature, log fold-change 10 | ANCOM-BC2 | BH | 1.000 [1.000, 1.000] | 1.000 | 0.000 [0.000, 0.000] |
| single feature, log fold-change 10 | ALDEx2 | Welch_128 * | 1.000 [1.000, 1.000] | 1.000 | 0.000 [0.000, 0.000] |
| single feature, log fold-change 10 | ALDEx2 | Welch_32 | 1.000 [1.000, 1.000] | 1.000 | 0.000 [0.000, 0.000] |
| single feature, log fold-change 10 | ALDEx2 | Welch_128_IQLR | 1.000 [1.000, 1.000] | 1.000 | 0.000 [0.000, 0.000] |
| single feature, log fold-change 10 | ANCOM-BC2 | BH_StrucZero | 1.000 [1.000, 1.000] | 1.000 | 0.000 [0.000, 0.000] |
| single feature, log fold-change 10 | MaAsLin 2 | TMM_NONE | 0.875 [0.477, 1.273] | 0.875 | 0.000 [0.000, 0.000] |
| single feature, log fold-change 10 | ANCOM-BC2 | Holm * | 1.000 [1.000, 1.000] | 1.000 | 0.000 [0.000, 0.000] |
| single feature, log fold-change 10 | ANCOM-BC2 | Holm_NoFilter | 0.067 [0.065, 0.070] | 1.000 | 0.995 [0.995, 0.995] |
| single feature, log fold-change 10 | ANCOM-BC2 | BH_NoFilter | 0.025 [0.024, 0.026] | 1.000 | 0.999 [0.999, 0.999] |
| ***Signal direction, genus level*** | | | | | |
| bidirectional | LinDA | Pseudo_0.1 | 0.870 [0.831, 0.908] | 0.865 | 0.094 [0.021, 0.167] |
| unidirectional | LinDA | Pseudo_0.1 | 0.951 [0.888, 1.013] | 1.000 | 0.082 [0.000, 0.184] |
| bidirectional | MaAsLin 2 | TMM_LOG | 0.924 [0.892, 0.956] | 0.891 | 0.024 [0.000, 0.082] |
| unidirectional | MaAsLin 2 | TMM_LOG | 0.910 [0.815, 1.005] | 1.000 | 0.147 [0.001, 0.293] |
| bidirectional | LinDA | Imputation | 0.876 [0.789, 0.963] | 0.880 | 0.096 [0.000, 0.241] |
| unidirectional | LinDA | Imputation | 0.757 [0.652, 0.862] | 0.984 | 0.345 [0.219, 0.471] |
| bidirectional | MaAsLin 3 | TSS_PLOG_Median | 0.896 [0.845, 0.947] | 0.854 | 0.038 [0.000, 0.078] |
| unidirectional | MaAsLin 3 | TSS_PLOG_Median | 0.902 [0.857, 0.947] | 0.990 | 0.148 [0.080, 0.216] |
| bidirectional | LinDA | Pseudo_0.5 | 0.859 [0.832, 0.886] | 0.849 | 0.094 [0.025, 0.164] |
| unidirectional | LinDA | Pseudo_0.5 | 0.952 [0.887, 1.017] | 1.000 | 0.078 [0.000, 0.183] |
| bidirectional | MaAsLin 3 | TSS_PLOG | 0.899 [0.837, 0.962] | 0.906 | 0.083 [0.015, 0.152] |
| unidirectional | MaAsLin 3 | TSS_PLOG | 0.774 [0.665, 0.882] | 0.974 | 0.319 [0.195, 0.442] |
| bidirectional | MaAsLin 2 | TSS_LOG * | 0.890 [0.832, 0.947] | 0.906 | 0.099 [0.002, 0.195] |
| unidirectional | MaAsLin 2 | TSS_LOG * | 0.753 [0.636, 0.869] | 0.974 | 0.339 [0.213, 0.465] |
| bidirectional | Wilcoxon | TSS | 0.883 [0.818, 0.948] | 0.891 | 0.097 [0.027, 0.167] |
| unidirectional | Wilcoxon | TSS | 0.730 [0.630, 0.829] | 0.979 | 0.377 [0.268, 0.486] |
| bidirectional | LinDA | Adaptive * | 0.853 [0.787, 0.919] | 0.865 | 0.121 [0.000, 0.249] |
| unidirectional | LinDA | Adaptive * | 0.759 [0.666, 0.853] | 0.984 | 0.343 [0.227, 0.459] |
| bidirectional | LDM | CLR | 0.908 [0.845, 0.971] | 0.896 | 0.059 [0.014, 0.103] |
| unidirectional | LDM | CLR | 0.963 [0.910, 1.016] | 1.000 | 0.063 [0.000, 0.151] |
| bidirectional | LDM | Omni3 | 0.866 [0.775, 0.956] | 0.917 | 0.146 [0.015, 0.277] |
| unidirectional | LDM | Omni3 | 0.625 [0.517, 0.733] | 0.995 | 0.493 [0.364, 0.621] |
| bidirectional | LinDA | Adaptive_NoWinsor | 0.877 [0.828, 0.926] | 0.870 | 0.086 [0.000, 0.187] |
| unidirectional | LinDA | Adaptive_NoWinsor | 0.785 [0.702, 0.867] | 0.995 | 0.313 [0.194, 0.433] |
| bidirectional | MaAsLin 2 | CLR_NONE | 0.859 [0.819, 0.899] | 0.880 | 0.112 [0.000, 0.237] |
| unidirectional | MaAsLin 2 | CLR_NONE | 0.563 [0.504, 0.621] | 1.000 | 0.568 [0.507, 0.628] |
| bidirectional | limma-voom | TMM_voom | 0.825 [0.691, 0.959] | 0.865 | 0.152 [0.000, 0.377] |
| unidirectional | limma-voom | TMM_voom | 0.544 [0.430, 0.658] | 1.000 | 0.557 [0.440, 0.675] |
| bidirectional | LDM | PA | 0.893 [0.824, 0.963] | 0.880 | 0.069 [0.000, 0.150] |
| unidirectional | LDM | PA | 0.698 [0.590, 0.805] | 0.891 | 0.348 [0.237, 0.459] |
| bidirectional | LDM | Arcsin | 0.855 [0.757, 0.953] | 0.911 | 0.160 [0.030, 0.290] |
| unidirectional | LDM | Arcsin | 0.644 [0.564, 0.724] | 0.984 | 0.474 [0.344, 0.604] |
| bidirectional | ZicoSeq | MultiLink | 0.849 [0.822, 0.876] | 0.839 | 0.100 [0.064, 0.137] |
| unidirectional | ZicoSeq | MultiLink | 0.961 [0.920, 1.002] | 0.995 | 0.062 [0.004, 0.120] |
| bidirectional | LinDA | Pseudo_1 | 0.849 [0.824, 0.874] | 0.833 | 0.097 [0.021, 0.172] |

| Level | Tool | Configuration | MCC | Recall | FDR |
|---|---|---|---|---|---|
| unidirectional | LinDA | Pseudo_1 | 0.957 [0.898, 1.016] | 1.000 | 0.072 [0.000, 0.169] |
| bidirectional | ZicoSeq | NoPosterior | 0.877 [0.840, 0.913] | 0.854 | 0.070 [0.038, 0.101] |
| unidirectional | ZicoSeq | NoPosterior | 0.943 [0.879, 1.006] | 0.953 | 0.046 [0.004, 0.089] |
| bidirectional | MaAsLin 3 | TSS_LOG_Median * | 0.867 [0.853, 0.881] | 0.797 | 0.027 [0.000, 0.069] |
| unidirectional | MaAsLin 3 | TSS_LOG_Median * | 0.974 [0.928, 1.020] | 0.984 | 0.029 [0.000, 0.103] |
| bidirectional | LDM | Omni * | 0.855 [0.750, 0.959] | 0.911 | 0.157 [0.020, 0.295] |
| unidirectional | LDM | Omni * | 0.617 [0.566, 0.669] | 0.984 | 0.503 [0.415, 0.591] |
| bidirectional | Wilcoxon | CLR | 0.808 [0.743, 0.872] | 0.854 | 0.155 [0.004, 0.307] |
| unidirectional | Wilcoxon | CLR | 0.479 [0.394, 0.564] | 0.958 | 0.609 [0.537, 0.682] |
| bidirectional | limma-voom | TMMwsp_voom | 0.794 [0.672, 0.915] | 0.865 | 0.183 [0.000, 0.424] |
| unidirectional | limma-voom | TMMwsp_voom | 0.430 [0.373, 0.487] | 0.974 | 0.654 [0.568, 0.740] |
| bidirectional | MaAsLin 3 | CLR_NONE | 0.855 [0.827, 0.882] | 0.786 | 0.039 [0.000, 0.091] |
| unidirectional | MaAsLin 3 | CLR_NONE | 0.846 [0.812, 0.879] | 0.953 | 0.194 [0.080, 0.308] |
| bidirectional | MaAsLin 3 | CLR_NONE_Median | 0.861 [0.850, 0.871] | 0.786 | 0.027 [0.000, 0.071] |
| unidirectional | MaAsLin 3 | CLR_NONE_Median | 0.955 [0.914, 0.997] | 0.953 | 0.029 [0.000, 0.103] |
| bidirectional | MaAsLin 3 | TSS_LOG | 0.797 [0.700, 0.894] | 0.802 | 0.144 [0.003, 0.286] |
| unidirectional | MaAsLin 3 | TSS_LOG | 0.639 [0.594, 0.683] | 0.964 | 0.453 [0.368, 0.537] |
| bidirectional | DESeq2 | LRT_poscounts | 0.788 [0.716, 0.859] | 0.823 | 0.191 [0.115, 0.266] |
| unidirectional | DESeq2 | LRT_poscounts | 0.679 [0.608, 0.750] | 0.995 | 0.470 [0.375, 0.564] |
| bidirectional | DESeq2 | Wald_poscounts * | 0.771 [0.685, 0.856] | 0.802 | 0.200 [0.111, 0.290] |
| unidirectional | DESeq2 | Wald_poscounts * | 0.673 [0.602, 0.744] | 0.995 | 0.478 [0.382, 0.573] |
| bidirectional | ZicoSeq | Default * | 0.824 [0.805, 0.843] | 0.802 | 0.107 [0.062, 0.151] |
| unidirectional | ZicoSeq | Default * | 0.955 [0.892, 1.018] | 0.979 | 0.059 [0.000, 0.132] |
| bidirectional | Wilcoxon | Rarefied | 0.824 [0.784, 0.864] | 0.807 | 0.113 [0.015, 0.211] |
| unidirectional | Wilcoxon | Rarefied | 0.734 [0.678, 0.790] | 0.979 | 0.367 [0.295, 0.439] |
| bidirectional | edgeR | TMM_LRT | 0.789 [0.754, 0.824] | 0.927 | 0.280 [0.229, 0.331] |
| unidirectional | edgeR | TMM_LRT | 0.691 [0.645, 0.737] | 0.990 | 0.458 [0.383, 0.533] |
| bidirectional | LDM | Freq | 0.850 [0.733, 0.968] | 0.880 | 0.130 [0.035, 0.226] |
| unidirectional | LDM | Freq | 0.640 [0.570, 0.710] | 0.927 | 0.449 [0.334, 0.565] |
| bidirectional | ALDEx2 | Wilcox_128 | 0.794 [0.744, 0.843] | 0.661 | 0.000 [0.000, 0.000] |
| unidirectional | ALDEx2 | Wilcox_128 | 0.904 [0.823, 0.986] | 0.938 | 0.099 [0.004, 0.195] |
| bidirectional | edgeR | TMM_QL * | 0.638 [0.613, 0.664] | 0.943 | 0.499 [0.458, 0.540] |
| unidirectional | edgeR | TMM_QL * | 0.571 [0.541, 0.602] | 0.995 | 0.603 [0.563, 0.642] |
| bidirectional | ANCOM-BC2 | BH | 0.862 [0.842, 0.881] | 0.766 | 0.000 [0.000, 0.000] |
| unidirectional | ANCOM-BC2 | BH | 0.973 [0.908, 1.038] | 0.964 | 0.005 [0.000, 0.022] |
| bidirectional | ALDEx2 | Welch_128 * | 0.761 [0.725, 0.797] | 0.615 | 0.000 [0.000, 0.000] |
| unidirectional | ALDEx2 | Welch_128 * | 0.897 [0.816, 0.979] | 0.885 | 0.064 [0.000, 0.154] |
| bidirectional | ALDEx2 | Welch_32 | 0.759 [0.731, 0.786] | 0.615 | 0.000 [0.000, 0.000] |
| unidirectional | ALDEx2 | Welch_32 | 0.906 [0.839, 0.974] | 0.896 | 0.053 [0.000, 0.143] |
| bidirectional | ALDEx2 | Welch_128_IQLR | 0.754 [0.715, 0.792] | 0.604 | 0.000 [0.000, 0.000] |
| unidirectional | ALDEx2 | Welch_128_IQLR | 0.934 [0.886, 0.981] | 0.911 | 0.022 [0.000, 0.076] |
| bidirectional | ANCOM-BC2 | BH_StrucZero | 0.839 [0.817, 0.862] | 0.729 | 0.000 [0.000, 0.000] |
| unidirectional | ANCOM-BC2 | BH_StrucZero | 0.967 [0.884, 1.050] | 0.953 | 0.005 [0.000, 0.022] |
| bidirectional | MaAsLin 2 | TMM_NONE | 0.702 [0.637, 0.767] | 0.573 | 0.026 [0.000, 0.110] |
| unidirectional | MaAsLin 2 | TMM_NONE | 0.813 [0.756, 0.870] | 0.786 | 0.109 [0.000, 0.231] |
| bidirectional | ANCOM-BC2 | Holm * | 0.822 [0.752, 0.893] | 0.708 | 0.000 [0.000, 0.000] |
| unidirectional | ANCOM-BC2 | Holm * | 0.957 [0.910, 1.005] | 0.932 | 0.000 [0.000, 0.000] |
| bidirectional | ANCOM-BC2 | Holm_NoFilter | 0.600 [0.495, 0.705] | 0.844 | 0.499 [0.398, 0.600] |
| unidirectional | ANCOM-BC2 | Holm_NoFilter | 0.677 [0.630, 0.724] | 0.990 | 0.476 [0.416, 0.535] |
| bidirectional | ANCOM-BC2 | BH_NoFilter | 0.474 [0.390, 0.559] | 0.870 | 0.653 [0.566, 0.740] |
| unidirectional | ANCOM-BC2 | BH_NoFilter | 0.525 [0.493, 0.556] | 0.990 | 0.648 [0.615, 0.680] |

| Level | Tool | Configuration | MCC | Recall | FDR |
|---|---|---|---|---|---|
| ***Signal direction, OTU level*** | | | | | |
| bidirectional | LinDA | Pseudo_0.1 | 0.860 [0.837, 0.884] | 0.810 | 0.056 [0.050, 0.063] |
| unidirectional | LinDA | Pseudo_0.1 | 0.969 [0.965, 0.974] | 0.996 | 0.050 [0.045, 0.055] |
| bidirectional | MaAsLin 2 | TMM_LOG | 0.863 [0.853, 0.872] | 0.810 | 0.052 [0.034, 0.069] |
| unidirectional | MaAsLin 2 | TMM_LOG | 0.953 [0.927, 0.978] | 0.999 | 0.080 [0.042, 0.118] |
| bidirectional | LinDA | Imputation | 0.775 [0.738, 0.812] | 0.742 | 0.141 [0.113, 0.170] |
| unidirectional | LinDA | Imputation | 0.840 [0.834, 0.845] | 0.996 | 0.255 [0.244, 0.266] |
| bidirectional | MaAsLin 3 | TSS_PLOG_Median | 0.798 [0.769, 0.826] | 0.720 | 0.076 [0.056, 0.096] |
| unidirectional | MaAsLin 3 | TSS_PLOG_Median | 0.944 [0.934, 0.954] | 0.996 | 0.092 [0.077, 0.107] |
| bidirectional | LinDA | Pseudo_0.5 | 0.866 [0.844, 0.887] | 0.815 | 0.052 [0.042, 0.062] |
| unidirectional | LinDA | Pseudo_0.5 | 0.971 [0.967, 0.975] | 0.998 | 0.050 [0.041, 0.058] |
| bidirectional | MaAsLin 3 | TSS_PLOG | 0.839 [0.820, 0.859] | 0.825 | 0.107 [0.075, 0.140] |
| unidirectional | MaAsLin 3 | TSS_PLOG | 0.867 [0.851, 0.884] | 0.993 | 0.206 [0.183, 0.229] |
| bidirectional | MaAsLin 2 | TSS_LOG * | 0.817 [0.803, 0.830] | 0.829 | 0.148 [0.127, 0.169] |
| unidirectional | MaAsLin 2 | TSS_LOG * | 0.789 [0.776, 0.803] | 0.995 | 0.316 [0.294, 0.338] |
| bidirectional | Wilcoxon | TSS | 0.828 [0.817, 0.840] | 0.800 | 0.101 [0.078, 0.124] |
| unidirectional | Wilcoxon | TSS | 0.872 [0.859, 0.884] | 0.988 | 0.200 [0.185, 0.216] |
| bidirectional | LinDA | Adaptive * | 0.776 [0.741, 0.811] | 0.744 | 0.141 [0.114, 0.167] |
| unidirectional | LinDA | Adaptive * | 0.840 [0.834, 0.845] | 0.996 | 0.255 [0.244, 0.266] |
| bidirectional | LDM | CLR | 0.874 [0.852, 0.896] | 0.832 | 0.055 [0.045, 0.065] |
| unidirectional | LDM | CLR | 0.973 [0.967, 0.979] | 0.998 | 0.046 [0.040, 0.052] |
| bidirectional | LDM | Omni3 | 0.660 [0.632, 0.687] | 0.878 | 0.386 [0.351, 0.421] |
| unidirectional | LDM | Omni3 | 0.442 [0.421, 0.463] | 0.996 | 0.654 [0.624, 0.683] |
| bidirectional | LinDA | Adaptive_NoWinsor | 0.794 [0.768, 0.819] | 0.733 | 0.097 [0.081, 0.113] |
| unidirectional | LinDA | Adaptive_NoWinsor | 0.900 [0.892, 0.907] | 0.997 | 0.166 [0.157, 0.174] |
| bidirectional | MaAsLin 2 | CLR_NONE | 0.802 [0.761, 0.842] | 0.940 | 0.255 [0.198, 0.312] |
| unidirectional | MaAsLin 2 | CLR_NONE | 0.427 [0.380, 0.474] | 0.995 | 0.701 [0.650, 0.752] |
| bidirectional | limma-voom | TMM_voom | 0.521 [0.368, 0.674] | 0.936 | 0.548 [0.365, 0.730] |
| unidirectional | limma-voom | TMM_voom | 0.266 [0.212, 0.319] | 0.986 | 0.798 [0.762, 0.835] |
| bidirectional | LDM | PA | 0.652 [0.626, 0.677] | 0.884 | 0.398 [0.370, 0.426] |
| unidirectional | LDM | PA | 0.430 [0.406, 0.455] | 0.996 | 0.661 [0.628, 0.694] |
| bidirectional | LDM | Arcsin | 0.756 [0.739, 0.773] | 0.868 | 0.268 [0.245, 0.290] |
| unidirectional | LDM | Arcsin | 0.598 [0.582, 0.614] | 0.994 | 0.520 [0.491, 0.549] |
| bidirectional | ZicoSeq | MultiLink | 0.795 [0.780, 0.809] | 0.729 | 0.090 [0.074, 0.106] |
| unidirectional | ZicoSeq | MultiLink | 0.565 [0.524, 0.606] | 0.999 | 0.479 [0.417, 0.542] |
| bidirectional | LinDA | Pseudo_1 | 0.865 [0.844, 0.886] | 0.812 | 0.049 [0.038, 0.060] |
| unidirectional | LinDA | Pseudo_1 | 0.973 [0.969, 0.977] | 0.998 | 0.046 [0.038, 0.054] |
| bidirectional | ZicoSeq | NoPosterior | 0.852 [0.840, 0.863] | 0.802 | 0.064 [0.043, 0.085] |
| unidirectional | ZicoSeq | NoPosterior | 0.830 [0.769, 0.891] | 0.998 | 0.213 [0.139, 0.286] |
| bidirectional | MaAsLin 3 | TSS_LOG_Median * | 0.755 [0.742, 0.767] | 0.617 | 0.030 [0.022, 0.038] |
| unidirectional | MaAsLin 3 | TSS_LOG_Median * | 0.976 [0.970, 0.982] | 0.991 | 0.034 [0.029, 0.039] |
| bidirectional | LDM | Omni * | 0.675 [0.648, 0.703] | 0.877 | 0.369 [0.332, 0.407] |
| unidirectional | LDM | Omni * | 0.463 [0.445, 0.480] | 0.994 | 0.637 [0.613, 0.662] |
| bidirectional | Wilcoxon | CLR | 0.394 [0.379, 0.408] | 0.878 | 0.634 [0.566, 0.702] |
| unidirectional | Wilcoxon | CLR | −0.049 [−0.084, −0.013] | 0.785 | 0.908 [0.904, 0.913] |
| bidirectional | limma-voom | TMMwsp_voom | 0.246 [0.106, 0.386] | 0.967 | 0.771 [0.602, 0.940] |
| unidirectional | limma-voom | TMMwsp_voom | −0.001 [−0.037, 0.034] | 0.939 | 0.900 [0.895, 0.906] |
| bidirectional | MaAsLin 3 | CLR_NONE | 0.708 [0.690, 0.726] | 0.678 | 0.185 [0.161, 0.209] |
| unidirectional | MaAsLin 3 | CLR_NONE | 0.668 [0.651, 0.686] | 0.987 | 0.451 [0.424, 0.478] |
| bidirectional | MaAsLin 3 | CLR_NONE_Median | 0.781 [0.770, 0.792] | 0.663 | 0.036 [0.024, 0.047] |
| unidirectional | MaAsLin 3 | CLR_NONE_Median | 0.979 [0.973, 0.984] | 0.992 | 0.030 [0.024, 0.036] |

| Level | Tool | Configuration | MCC | Recall | FDR |
|---|---|---|---|---|---|
| bidirectional | MaAsLin 3 | TSS_LOG | 0.468 [0.418, 0.517] | 0.735 | 0.516 [0.458, 0.573] |
| unidirectional | MaAsLin 3 | TSS_LOG | 0.325 [0.251, 0.399] | 0.978 | 0.750 [0.672, 0.827] |
| bidirectional | DESeq2 | LRT_poscounts | 0.753 [0.739, 0.767] | 0.694 | 0.132 [0.114, 0.151] |
| unidirectional | DESeq2 | LRT_poscounts | 0.793 [0.762, 0.825] | 0.992 | 0.318 [0.275, 0.362] |
| bidirectional | DESeq2 | Wald_poscounts * | 0.727 [0.712, 0.741] | 0.658 | 0.142 [0.123, 0.161] |
| unidirectional | DESeq2 | Wald_poscounts * | 0.793 [0.765, 0.821] | 0.993 | 0.320 [0.280, 0.360] |
| bidirectional | ZicoSeq | Default * | 0.800 [0.787, 0.813] | 0.737 | 0.089 [0.071, 0.107] |
| unidirectional | ZicoSeq | Default * | 0.568 [0.500, 0.635] | 0.997 | 0.480 [0.395, 0.565] |
| bidirectional | Wilcoxon | Rarefied | 0.685 [0.646, 0.724] | 0.635 | 0.178 [0.150, 0.205] |
| unidirectional | Wilcoxon | Rarefied | 0.718 [0.698, 0.739] | 0.938 | 0.363 [0.341, 0.386] |
| bidirectional | edgeR | TMM_LRT | 0.808 [0.773, 0.843] | 0.908 | 0.241 [0.195, 0.288] |
| unidirectional | edgeR | TMM_LRT | 0.797 [0.765, 0.830] | 0.994 | 0.311 [0.265, 0.358] |
| bidirectional | LDM | Freq | 0.708 [0.677, 0.738] | 0.857 | 0.323 [0.281, 0.365] |
| unidirectional | LDM | Freq | 0.536 [0.515, 0.556] | 0.969 | 0.580 [0.555, 0.604] |
| bidirectional | ALDEx2 | Wilcox_128 | 0.674 [0.660, 0.687] | 0.488 | 0.000 [0.000, 0.000] |
| unidirectional | ALDEx2 | Wilcox_128 | 0.926 [0.922, 0.929] | 0.877 | 0.000 [0.000, 0.000] |
| bidirectional | edgeR | TMM_QL * | 0.716 [0.685, 0.748] | 0.916 | 0.384 [0.349, 0.419] |
| unidirectional | edgeR | TMM_QL * | 0.708 [0.684, 0.732] | 0.996 | 0.435 [0.402, 0.467] |
| bidirectional | ANCOM-BC2 | BH | 0.794 [0.769, 0.820] | 0.660 | 0.004 [0.002, 0.006] |
| unidirectional | ANCOM-BC2 | BH | 0.992 [0.987, 0.996] | 0.988 | 0.003 [0.000, 0.006] |
| bidirectional | ALDEx2 | Welch_128 * | 0.620 [0.616, 0.625] | 0.426 | 0.000 [0.000, 0.000] |
| unidirectional | ALDEx2 | Welch_128 * | 0.865 [0.854, 0.875] | 0.789 | 0.000 [0.000, 0.000] |
| bidirectional | ALDEx2 | Welch_32 | 0.624 [0.618, 0.631] | 0.431 | 0.000 [0.000, 0.000] |
| unidirectional | ALDEx2 | Welch_32 | 0.868 [0.857, 0.880] | 0.794 | 0.000 [0.000, 0.000] |
| bidirectional | ALDEx2 | Welch_128_IQLR | 0.628 [0.623, 0.634] | 0.433 | 0.000 [0.000, 0.000] |
| unidirectional | ALDEx2 | Welch_128_IQLR | 0.908 [0.901, 0.915] | 0.852 | 0.000 [0.000, 0.000] |
| bidirectional | ANCOM-BC2 | BH_StrucZero | 0.755 [0.736, 0.775] | 0.600 | 0.004 [0.002, 0.006] |
| unidirectional | ANCOM-BC2 | BH_StrucZero | 0.985 [0.982, 0.989] | 0.977 | 0.003 [0.000, 0.006] |
| bidirectional | MaAsLin 2 | TMM_NONE | 0.756 [0.736, 0.776] | 0.620 | 0.021 [0.007, 0.034] |
| unidirectional | MaAsLin 2 | TMM_NONE | 0.921 [0.908, 0.934] | 0.927 | 0.065 [0.045, 0.085] |
| bidirectional | ANCOM-BC2 | Holm * | 0.685 [0.670, 0.700] | 0.508 | 0.000 [0.000, 0.000] |
| unidirectional | ANCOM-BC2 | Holm * | 0.917 [0.914, 0.921] | 0.870 | 0.000 [0.000, 0.000] |
| bidirectional | ANCOM-BC2 | Holm_NoFilter | 0.540 [0.526, 0.554] | 0.711 | 0.498 [0.488, 0.509] |
| unidirectional | ANCOM-BC2 | Holm_NoFilter | 0.718 [0.697, 0.739] | 0.992 | 0.427 [0.401, 0.454] |
| bidirectional | ANCOM-BC2 | BH_NoFilter | 0.243 [0.232, 0.254] | 0.781 | 0.814 [0.811, 0.817] |
| unidirectional | ANCOM-BC2 | BH_NoFilter | 0.365 [0.349, 0.382] | 0.999 | 0.779 [0.768, 0.789] |

**Table S13. Shifts of most taxa in one direction.** Stool genera at fixed read depth. In each dataset, 10%, 30%, 50% or 70% of the genera were decreased or increased together with an effect size of 1.5, at 100 and 200 samples, four replicates (eight datasets per cell). Mean MCC / mean empirical FDR against the implanted genera.

| Tool | Configuration | Decrease 10% | Decrease 30% | Decrease 50% | Decrease 70% | Increase 10% | Increase 30% | Increase 50% | Increase 70% |
|---|---|---|---|---|---|---|---|---|---|
| LinDA | Pseudo_0.1 | 0.70 / 0.03 | 0.71 / 0.15 | 0.64 / 0.10 | −0.01 / 0.31 | 0.87 / 0.14 | 0.86 / 0.16 | 0.49 / 0.27 | −0.43 / 0.53 |
| MaAsLin 2 | TMM_LOG | 0.69 / 0.08 | 0.70 / 0.17 | 0.55 / 0.17 | 0.26 / 0.15 | 0.94 / 0.10 | 0.77 / 0.27 | 0.51 / 0.27 | 0.47 / 0.13 |
| LinDA | Imputation | 0.68 / 0.08 | 0.65 / 0.21 | 0.58 / 0.15 | −0.02 / 0.34 | 0.89 / 0.13 | 0.78 / 0.23 | 0.43 / 0.30 | −0.33 / 0.49 |
| MaAsLin 3 | TSS_PLOG_Median | 0.70 / 0.07 | 0.64 / 0.14 | 0.32 / 0.26 | −0.42 / 0.67 | 0.92 / 0.11 | 0.88 / 0.12 | 0.21 / 0.38 | −0.71 / 0.69 |
| LinDA | Pseudo_0.5 | 0.68 / 0.04 | 0.66 / 0.17 | 0.59 / 0.12 | 0.00 / 0.33 | 0.86 / 0.15 | 0.82 / 0.20 | 0.42 / 0.30 | −0.56 / 0.60 |
| MaAsLin 3 | TSS_PLOG | 0.67 / 0.07 | 0.68 / 0.15 | 0.63 / 0.12 | 0.28 / 0.12 | 0.85 / 0.17 | 0.77 / 0.23 | 0.44 / 0.29 | 0.51 / 0.13 |
| MaAsLin 2 | TSS_LOG * | 0.66 / 0.08 | 0.65 / 0.19 | 0.60 / 0.14 | 0.18 / 0.17 | 0.86 / 0.17 | 0.76 / 0.25 | 0.42 / 0.30 | 0.48 / 0.15 |
| Wilcoxon | TSS | 0.68 / 0.06 | 0.69 / 0.17 | 0.62 / 0.13 | 0.28 / 0.12 | 0.84 / 0.19 | 0.75 / 0.26 | 0.43 / 0.29 | 0.48 / 0.14 |
| LinDA | Adaptive * | 0.67 / 0.07 | 0.66 / 0.17 | 0.59 / 0.12 | 0.02 / 0.31 | 0.86 / 0.15 | 0.81 / 0.21 | 0.44 / 0.30 | −0.56 / 0.60 |
| LDM | CLR | 0.69 / 0.09 | 0.69 / 0.17 | 0.29 / 0.32 | −0.46 / 0.64 | 0.90 / 0.13 | 0.79 / 0.22 | 0.25 / 0.38 | −0.70 / 0.69 |
| LDM | Omni3 | 0.69 / 0.08 | 0.62 / 0.26 | 0.51 / 0.24 | 0.18 / 0.23 | 0.85 / 0.18 | 0.65 / 0.36 | 0.31 / 0.39 | 0.38 / 0.20 |
| LinDA | Adaptive_NoWinsor | 0.69 / 0.07 | 0.68 / 0.18 | 0.62 / 0.13 | −0.08 / 0.37 | 0.88 / 0.13 | 0.84 / 0.18 | 0.49 / 0.27 | −0.47 / 0.57 |
| MaAsLin 2 | CLR_NONE | 0.62 / 0.14 | 0.31 / 0.48 | −0.12 / 0.55 | −0.49 / 0.67 | 0.81 / 0.25 | 0.26 / 0.59 | −0.14 / 0.53 | −0.60 / 0.64 |
| limma-voom | TMM_voom | 0.54 / 0.26 | 0.47 / 0.35 | 0.09 / 0.41 | 0.52 / 0.11 | 0.87 / 0.18 | 0.67 / 0.35 | 0.38 / 0.32 | 0.06 / 0.27 |
| LDM | PA | 0.66 / 0.07 | 0.65 / 0.14 | 0.62 / 0.12 | 0.28 / 0.16 | 0.82 / 0.14 | 0.74 / 0.24 | 0.38 / 0.34 | 0.46 / 0.16 |
| LDM | Arcsin | 0.69 / 0.05 | 0.63 / 0.26 | 0.52 / 0.23 | 0.16 / 0.25 | 0.85 / 0.18 | 0.62 / 0.39 | 0.25 / 0.42 | 0.35 / 0.21 |
| ZicoSeq | MultiLink | 0.58 / 0.12 | 0.60 / 0.17 | −0.18 / 0.66 | −0.13 / 0.48 | 0.98 / 0.03 | 0.54 / 0.36 | 0.11 / 0.46 | 0.08 / 0.30 |
| LinDA | Pseudo_1 | 0.68 / 0.05 | 0.63 / 0.19 | 0.56 / 0.14 | −0.00 / 0.33 | 0.86 / 0.15 | 0.80 / 0.22 | 0.36 / 0.33 | −0.37 / 0.51 |
| ZicoSeq | NoPosterior | 0.67 / 0.02 | 0.66 / 0.11 | 0.16 / 0.38 | −0.17 / 0.59 | 0.94 / 0.05 | 0.69 / 0.26 | 0.12 / 0.45 | 0.18 / 0.23 |
| MaAsLin 3 | TSS_LOG_Median * | 0.58 / 0.03 | 0.68 / 0.04 | 0.48 / 0.18 | 0.21 / 0.20 | 0.87 / 0.07 | 0.83 / 0.13 | 0.39 / 0.28 | 0.31 / 0.18 |
| LDM | Omni * | 0.70 / 0.05 | 0.60 / 0.27 | 0.49 / 0.25 | 0.10 / 0.28 | 0.85 / 0.19 | 0.62 / 0.39 | 0.25 / 0.42 | 0.31 / 0.22 |
| Wilcoxon | CLR | 0.56 / 0.12 | 0.22 / 0.51 | −0.24 / 0.65 | −0.46 / 0.70 | 0.65 / 0.33 | −0.01 / 0.65 | −0.24 / 0.59 | −0.64 / 0.68 |
| limma-voom | TMMwsp_voom | 0.52 / 0.22 | 0.36 / 0.43 | 0.08 / 0.44 | −0.22 / 0.51 | 0.78 / 0.24 | 0.24 / 0.55 | 0.03 / 0.48 | −0.39 / 0.57 |
| MaAsLin 3 | CLR_NONE | 0.58 / 0.06 | 0.59 / 0.12 | 0.53 / 0.14 | 0.34 / 0.10 | 0.81 / 0.18 | 0.67 / 0.29 | 0.48 / 0.22 | 0.42 / 0.12 |
| MaAsLin 3 | CLR_NONE_Median | 0.57 / 0.02 | 0.64 / 0.04 | 0.34 / 0.27 | 0.19 / 0.18 | 0.84 / 0.09 | 0.80 / 0.12 | 0.29 / 0.33 | 0.28 / 0.17 |
| MaAsLin 3 | TSS_LOG | 0.60 / 0.04 | 0.53 / 0.24 | 0.39 / 0.26 | 0.10 / 0.25 | 0.75 / 0.19 | 0.69 / 0.29 | 0.39 / 0.30 | 0.40 / 0.17 |
| DESeq2 | LRT_poscounts | 0.62 / 0.15 | 0.50 / 0.32 | 0.30 / 0.31 | 0.05 / 0.23 | 0.82 / 0.25 | 0.62 / 0.39 | 0.29 / 0.35 | 0.13 / 0.25 |
| DESeq2 | Wald_poscounts * | 0.59 / 0.14 | 0.47 / 0.34 | 0.29 / 0.31 | 0.05 / 0.23 | 0.82 / 0.25 | 0.62 / 0.38 | 0.28 / 0.35 | 0.12 / 0.25 |
| ZicoSeq | Default * | 0.50 / 0.16 | 0.37 / 0.31 | −0.36 / 0.81 | −0.22 / 0.54 | 0.97 / 0.06 | 0.42 / 0.40 | −0.02 / 0.54 | 0.04 / 0.34 |
| Wilcoxon | Rarefied | 0.58 / 0.07 | 0.58 / 0.20 | 0.48 / 0.16 | 0.06 / 0.25 | 0.84 / 0.20 | 0.72 / 0.26 | 0.32 / 0.33 | 0.34 / 0.19 |
| edgeR | TMM_LRT | 0.62 / 0.30 | 0.53 / 0.36 | 0.27 / 0.34 | 0.00 / 0.31 | 0.79 / 0.27 | 0.58 / 0.41 | 0.20 / 0.41 | 0.06 / 0.29 |
| LDM | Freq | 0.68 / 0.05 | 0.57 / 0.26 | 0.39 / 0.24 | −0.14 / 0.47 | 0.82 / 0.18 | 0.59 / 0.35 | 0.14 / 0.43 | 0.16 / 0.29 |
| ALDEx2 | Wilcox_128 | 0.52 / 0.00 | 0.30 / 0.36 | 0.07 / 0.47 | −0.22 / 0.70 | 0.79 / 0.10 | 0.42 / 0.40 | −0.10 / 0.65 | −0.24 / 0.67 |
| edgeR | TMM_QL * | 0.55 / 0.53 | 0.51 / 0.44 | 0.24 / 0.40 | −0.03 / 0.32 | 0.63 / 0.54 | 0.47 / 0.51 | 0.14 / 0.45 | 0.06 / 0.28 |
| ANCOM-BC2 | BH | 0.50 / 0.00 | 0.59 / 0.01 | −0.09 / 0.59 | −0.29 / 0.77 | 0.81 / 0.12 | 0.81 / 0.09 | −0.45 / 0.97 | −0.37 / 0.91 |
| ALDEx2 | Welch_128 * | 0.45 / 0.00 | 0.32 / 0.31 | 0.12 / 0.40 | −0.20 / 0.69 | 0.68 / 0.11 | 0.40 / 0.38 | −0.10 / 0.68 | −0.25 / 0.75 |
| ALDEx2 | Welch_32 | 0.43 / 0.00 | 0.33 / 0.30 | 0.12 / 0.40 | −0.21 / 0.71 | 0.69 / 0.12 | 0.41 / 0.38 | −0.10 / 0.68 | −0.25 / 0.76 |
| ALDEx2 | Welch_128_IQLR | 0.44 / 0.00 | 0.42 / 0.16 | 0.14 / 0.35 | −0.12 / 0.55 | 0.76 / 0.06 | 0.53 / 0.28 | −0.07 / 0.67 | −0.26 / 0.74 |
| ANCOM-BC2 | BH_StrucZero | 0.47 / 0.00 | 0.57 / 0.01 | −0.08 / 0.58 | −0.28 / 0.77 | 0.81 / 0.12 | 0.80 / 0.09 | −0.45 / 0.97 | −0.37 / 0.91 |
| MaAsLin 2 | TMM_NONE | 0.37 / 0.00 | 0.42 / 0.21 | 0.10 / 0.37 | −0.06 / 0.44 | 0.75 / 0.06 | 0.56 / 0.27 | 0.21 / 0.36 | 0.07 / 0.30 |
| ANCOM-BC2 | Holm * | 0.45 / 0.00 | 0.47 / 0.00 | −0.04 / 0.51 | −0.23 / 0.75 | 0.74 / 0.00 | 0.58 / 0.01 | −0.34 / 1.00 | −0.34 / 1.00 |
| ANCOM-BC2 | Holm_NoFilter | 0.41 / 0.53 | 0.53 / 0.28 | −0.02 / 0.52 | −0.18 / 0.49 | 0.72 / 0.41 | 0.61 / 0.24 | −0.35 / 0.80 | −0.21 / 0.54 |
| ANCOM-BC2 | BH_NoFilter | 0.29 / 0.74 | 0.44 / 0.42 | 0.06 / 0.47 | −0.08 / 0.36 | 0.58 / 0.60 | 0.58 / 0.41 | −0.26 / 0.69 | −0.10 / 0.40 |

**Table S14. Read depth that differs between groups, null data.** Stool genera without signal. The reads of every case sample were thinned binomially to the stated percentage of reads, which lowers the read depth of the cases without changing their expected composition, and every level was fitted with the same seeds as the unthinned dataset (10 datasets at each of 100 and 200 samples). The mean depth of the controls was 1.01, 1.12, 1.35, 2.01, 4.04 and 10.04 times that of the cases. In the last two columns, 60 or 100 real stool samples were split into their shallower and deeper halves, which differed 2.4-fold in depth, or at random (20 draws each). With 200 samples, nearly the whole cohort, the halves differed 2.5-fold and most configurations made calls in every depth split. These splits are in I_depthsplit.csv in Zenodo. Cells give the datasets with at least one call and, in brackets, the mean number of calls.

| Tool | Configuration | 100% of reads | 90% of reads | 75% of reads | 50% of reads | 25% of reads | 10% of reads | Split by depth | Random split |
|---|---|---|---|---|---|---|---|---|---|
| LinDA | Pseudo_0.1 | 0 of 20 (0.0) | 0 of 20 (0.0) | 4 of 20 (0.3) | 20 of 20 (8.8) | 20 of 20 (27.6) | 20 of 20 (46.3) | 39 of 40 (13.9) | 4 of 40 (0.1) |
| MaAsLin 2 | TMM_LOG | 0 of 20 (0.0) | 0 of 20 (0.0) | 1 of 20 (0.1) | 5 of 20 (0.3) | 15 of 20 (2.6) | 20 of 20 (24.9) | 23 of 40 (1.9) | 0 of 40 (0.0) |
| LinDA | Imputation | 0 of 20 (0.0) | 0 of 20 (0.0) | 0 of 20 (0.0) | 1 of 20 (0.1) | 3 of 20 (0.2) | 15 of 20 (4.2) | 14 of 40 (0.7) | 3 of 40 (0.1) |
| MaAsLin 3 | TSS_PLOG_Median | 0 of 20 (0.0) | 0 of 20 (0.0) | 0 of 20 (0.0) | 2 of 20 (0.1) | 6 of 20 (0.6) | 18 of 20 (5.8) | 15 of 40 (0.9) | 0 of 40 (0.0) |
| LinDA | Pseudo_0.5 | 0 of 20 (0.0) | 1 of 20 (0.1) | 5 of 20 (0.5) | 20 of 20 (10.8) | 20 of 20 (28.5) | 20 of 20 (47.5) | 40 of 40 (17.4) | 4 of 40 (0.1) |
| MaAsLin 3 | TSS_PLOG | 0 of 20 (0.0) | 0 of 20 (0.0) | 0 of 20 (0.0) | 4 of 20 (0.2) | 13 of 20 (1.2) | 20 of 20 (17.3) | 19 of 40 (1.7) | 0 of 40 (0.0) |
| MaAsLin 2 | TSS_LOG * | 0 of 20 (0.0) | 0 of 20 (0.0) | 0 of 20 (0.0) | 2 of 20 (0.1) | 8 of 20 (0.7) | 20 of 20 (9.6) | 16 of 40 (1.2) | 1 of 40 (0.0) |
| Wilcoxon | TSS | 1 of 20 (0.1) | 1 of 20 (0.1) | 1 of 20 (0.1) | 6 of 20 (0.3) | 13 of 20 (2.5) | 20 of 20 (23.2) | 24 of 40 (3.0) | 2 of 40 (0.1) |
| LinDA | Adaptive * | 0 of 20 (0.0) | 1 of 20 (0.1) | 0 of 20 (0.0) | 1 of 20 (0.1) | 3 of 20 (0.2) | 15 of 20 (4.2) | 14 of 40 (0.7) | 4 of 40 (0.1) |
| LDM | CLR | 0 of 20 (0.0) | 0 of 20 (0.0) | 1 of 20 (0.1) | 17 of 20 (11.7) | 20 of 20 (40.5) | 20 of 20 (55.5) | 40 of 40 (23.7) | 1 of 40 (0.0) |
| LDM | Omni3 | 0 of 20 (0.0) | 0 of 20 (0.0) | 0 of 20 (0.0) | 2 of 20 (0.1) | 4 of 20 (0.3) | 15 of 20 (3.2) | 14 of 40 (0.9) | 0 of 40 (0.0) |
| LinDA | Adaptive_NoWinsor | 0 of 20 (0.0) | 1 of 20 (0.1) | 0 of 20 (0.0) | 2 of 20 (0.1) | 3 of 20 (0.1) | 12 of 20 (2.5) | 15 of 40 (0.6) | 3 of 40 (0.1) |
| MaAsLin 2 | CLR_NONE | 0 of 20 (0.0) | 0 of 20 (0.0) | 3 of 20 (0.2) | 19 of 20 (24.9) | 20 of 20 (56.1) | 20 of 20 (65.9) | 40 of 40 (32.4) | 2 of 40 (0.1) |
| limma-voom | TMM_voom | 0 of 20 (0.0) | 2 of 20 (0.1) | 6 of 20 (3.5) | 19 of 20 (40.6) | 20 of 20 (61.2) | 20 of 20 (70.7) | 40 of 40 (38.4) | 4 of 40 (0.1) |
| LDM | PA | 1 of 20 (0.1) | 1 of 20 (0.1) | 1 of 20 (0.1) | 0 of 20 (0.0) | 1 of 20 (0.1) | 12 of 20 (0.9) | 12 of 40 (0.6) | 0 of 40 (0.0) |
| LDM | Arcsin | 0 of 20 (0.0) | 0 of 20 (0.0) | 0 of 20 (0.0) | 2 of 20 (0.1) | 4 of 20 (0.3) | 17 of 20 (4.7) | 14 of 40 (1.1) | 0 of 40 (0.0) |
| ZicoSeq | MultiLink | 1 of 20 (0.1) | 4 of 20 (0.3) | 3 of 20 (0.8) | 5 of 20 (2.8) | 12 of 20 (14.3) | 20 of 20 (39.5) | 22 of 40 (5.4) | 1 of 40 (0.0) |
| LinDA | Pseudo_1 | 1 of 20 (0.1) | 2 of 20 (0.1) | 5 of 20 (0.7) | 20 of 20 (11.6) | 20 of 20 (28.8) | 20 of 20 (48.0) | 40 of 40 (17.8) | 3 of 40 (0.1) |
| ZicoSeq | NoPosterior | 0 of 20 (0.0) | 0 of 20 (0.0) | 0 of 20 (0.0) | 2 of 20 (0.1) | 9 of 20 (1.1) | 20 of 20 (24.1) | 20 of 40 (1.4) | 1 of 40 (0.0) |
| MaAsLin 3 | TSS_LOG_Median * | 1 of 20 (0.1) | 2 of 20 (0.1) | 3 of 20 (0.1) | 10 of 20 (1.0) | 20 of 20 (15.5) | 20 of 20 (37.9) | 30 of 40 (7.1) | 0 of 40 (0.0) |
| LDM | Omni * | 0 of 20 (0.0) | 0 of 20 (0.0) | 0 of 20 (0.0) | 2 of 20 (0.1) | 4 of 20 (0.3) | 15 of 20 (3.8) | 14 of 40 (0.9) | 0 of 40 (0.0) |
| Wilcoxon | CLR | 2 of 20 (0.1) | 2 of 20 (0.1) | 5 of 20 (1.1) | 19 of 20 (37.4) | 20 of 20 (67.5) | 20 of 20 (74.5) | 40 of 40 (46.5) | 3 of 40 (0.1) |
| limma-voom | TMMwsp_voom | 1 of 20 (0.1) | 1 of 20 (0.3) | 9 of 20 (7.8) | 20 of 20 (44.1) | 20 of 20 (62.9) | 20 of 20 (70.8) | 40 of 40 (39.4) | 3 of 40 (0.1) |
| MaAsLin 3 | CLR_NONE | 1 of 20 (0.1) | 1 of 20 (0.1) | 2 of 20 (0.1) | 11 of 20 (1.1) | 20 of 20 (18.6) | 20 of 20 (35.2) | 40 of 40 (11.7) | 2 of 40 (0.1) |
| MaAsLin 3 | CLR_NONE_Median | 0 of 20 (0.0) | 0 of 20 (0.0) | 1 of 20 (0.1) | 5 of 20 (0.3) | 20 of 20 (9.3) | 20 of 20 (30.1) | 29 of 40 (4.1) | 2 of 40 (0.1) |
| MaAsLin 3 | TSS_LOG | 0 of 20 (0.0) | 1 of 20 (0.1) | 1 of 20 (0.1) | 10 of 20 (0.9) | 19 of 20 (15.6) | 20 of 20 (37.9) | 31 of 40 (6.0) | 1 of 40 (0.0) |
| DESeq2 | LRT_poscounts | 5 of 20 (0.5) | 4 of 20 (0.2) | 7 of 20 (0.4) | 10 of 20 (1.2) | 20 of 20 (9.0) | 20 of 20 (19.9) | 36 of 40 (4.4) | 8 of 40 (0.3) |
| DESeq2 | Wald_poscounts * | 6 of 20 (0.5) | 5 of 20 (0.3) | 8 of 20 (0.5) | 10 of 20 (1.2) | 20 of 20 (9.0) | 20 of 20 (19.0) | 37 of 40 (4.4) | 12 of 40 (0.5) |
| ZicoSeq | Default * | 2 of 20 (0.1) | 2 of 20 (0.1) | 5 of 20 (1.1) | 6 of 20 (2.8) | 15 of 20 (18.4) | 19 of 20 (37.3) | 19 of 40 (5.6) | 3 of 40 (0.1) |
| Wilcoxon | Rarefied | 1 of 20 (0.1) | 2 of 20 (0.1) | 0 of 20 (0.0) | 0 of 20 (0.0) | 1 of 20 (0.1) | 0 of 20 (0.0) | 2 of 40 (0.1) | 0 of 40 (0.0) |
| edgeR | TMM_LRT | 17 of 20 (2.5) | 17 of 20 (2.5) | 17 of 20 (2.1) | 16 of 20 (2.4) | 18 of 20 (2.9) | 19 of 20 (8.5) | 34 of 40 (3.4) | 20 of 40 (0.8) |
| LDM | Freq | 0 of 20 (0.0) | 0 of 20 (0.0) | 0 of 20 (0.0) | 0 of 20 (0.0) | 1 of 20 (0.1) | 12 of 20 (1.0) | 8 of 40 (0.3) | 0 of 40 (0.0) |
| ALDEx2 | Wilcox_128 | 0 of 20 (0.0) | 1 of 20 (0.1) | 2 of 20 (0.1) | 10 of 20 (1.1) | 20 of 20 (12.1) | 20 of 20 (18.8) | 23 of 40 (1.8) | 0 of 40 (0.0) |
| edgeR | TMM_QL * | 20 of 20 (7.2) | 20 of 20 (7.1) | 19 of 20 (7.2) | 20 of 20 (6.7) | 20 of 20 (7.5) | 20 of 20 (11.9) | 39 of 40 (7.0) | 40 of 40 (4.8) |
| ANCOM-BC2 | BH | 0 of 20 (0.0) | 0 of 20 (0.0) | 0 of 20 (0.0) | 2 of 20 (0.1) | 18 of 20 (5.3) | 20 of 20 (11.6) | 5 of 40 (0.2) | 0 of 40 (0.0) |
| ALDEx2 | Welch_128 * | 0 of 20 (0.0) | 0 of 20 (0.0) | 0 of 20 (0.0) | 7 of 20 (0.6) | 20 of 20 (8.9) | 20 of 20 (15.1) | 17 of 40 (0.7) | 0 of 40 (0.0) |
| ALDEx2 | Welch_32 | 0 of 20 (0.0) | 0 of 20 (0.0) | 0 of 20 (0.0) | 7 of 20 (0.7) | 20 of 20 (9.1) | 20 of 20 (15.4) | 16 of 40 (0.7) | 0 of 40 (0.0) |
| ALDEx2 | Welch_128_IQLR | 0 of 20 (0.0) | 0 of 20 (0.0) | 0 of 20 (0.0) | 3 of 20 (0.2) | 20 of 20 (8.2) | 20 of 20 (15.3) | 5 of 40 (0.2) | 0 of 40 (0.0) |
| ANCOM-BC2 | BH_StrucZero | 0 of 20 (0.0) | 0 of 20 (0.0) | 0 of 20 (0.0) | 2 of 20 (0.1) | 17 of 20 (4.8) | 20 of 20 (11.5) | 5 of 40 (0.1) | 0 of 40 (0.0) |
| MaAsLin 2 | TMM_NONE | 0 of 20 (0.0) | 0 of 20 (0.0) | 0 of 20 (0.0) | 0 of 20 (0.0) | 4 of 20 (0.2) | 12 of 20 (2.1) | 1 of 40 (0.0) | 0 of 40 (0.0) |
| ANCOM-BC2 | Holm * | 0 of 20 (0.0) | 0 of 20 (0.0) | 0 of 20 (0.0) | 1 of 20 (0.1) | 8 of 20 (1.3) | 18 of 20 (6.4) | 1 of 40 (0.0) | 0 of 40 (0.0) |
| ANCOM-BC2 | Holm_NoFilter | 20 of 20 (8.0) | 20 of 20 (7.9) | 20 of 20 (7.4) | 20 of 20 (8.9) | 20 of 20 (12.6) | 20 of 20 (20.1) | 40 of 40 (9.7) | 40 of 40 (6.2) |

| Tool | Configuration | 100% of reads | 90% of reads | 75% of reads | 50% of reads | 25% of reads | 10% of reads | Split by depth | Random split |
|---|---|---|---|---|---|---|---|---|---|
| ANCOM-BC2 | BH_NoFilter | 20 of 20 (14.7) | 20 of 20 (15.5) | 20 of 20 (13.2) | 20 of 20 (17.6) | 20 of 20 (26.1) | 20 of 20 (32.4) | 40 of 40 (21.3) | 40 of 40 (13.7) |

**Table S15. Confounding.** Empirical FDR and, in brackets, the mean number of confounded features called per dataset, by the Pearson correlation r between the covariate and the group, without and with the covariate in the model, at 200 samples. Empty cells mark configurations without a covariate-adjusted form.

| Tool | Configuration | r 0, unadjusted | r 0, adjusted | r 0.3, unadjusted | r 0.3, adjusted | r 0.6, unadjusted | r 0.6, adjusted |
|---|---|---|---|---|---|---|---|
| ***Genus level*** | | | | | | | |
| LinDA | Pseudo_0.1 | 0.04 (0.1) | 0.05 (0.1) | 0.17 (1.4) | 0.01 (0.0) | 0.36 (3.3) | 0.05 (0.2) |
| MaAsLin 2 | TMM_LOG | 0.05 (0.1) | 0.04 (0.2) | 0.22 (1.5) | 0.05 (0.1) | 0.36 (3.5) | 0.04 (0.0) |
| LinDA | Imputation | 0.06 (0.2) | 0.05 (0.1) | 0.28 (1.4) | 0.13 (0.2) | 0.44 (3.2) | 0.17 (0.4) |
| MaAsLin 3 | TSS_PLOG_Median | 0.01 (0.0) | 0.02 (0.0) | 0.12 (0.9) | 0.02 (0.1) | 0.33 (2.7) | 0.06 (0.1) |
| LinDA | Pseudo_0.5 | 0.01 (0.0) | 0.04 (0.2) | 0.18 (1.5) | 0.01 (0.0) | 0.39 (3.8) | 0.08 (0.2) |
| MaAsLin 3 | TSS_PLOG | 0.07 (0.1) | 0.07 (0.1) | 0.19 (1.0) | 0.07 (0.1) | 0.40 (2.8) | 0.11 (0.2) |
| MaAsLin 2 | TSS_LOG * | 0.07 (0.1) | 0.07 (0.1) | 0.23 (1.2) | 0.10 (0.2) | 0.44 (3.3) | 0.15 (0.3) |
| Wilcoxon | TSS | 0.11 (0.1) | | 0.24 (1.3) | | 0.39 (2.3) | |
| LinDA | Adaptive * | 0.03 (0.1) | 0.07 (0.2) | 0.29 (1.5) | 0.11 (0.2) | 0.46 (3.6) | 0.17 (0.4) |
| LDM | CLR | 0.02 (0.0) | 0.04 (0.0) | 0.17 (1.5) | 0.00 (0.0) | 0.37 (4.0) | 0.03 (0.0) |
| LDM | Omni3 | 0.15 (0.2) | 0.13 (0.2) | 0.30 (1.8) | 0.11 (0.0) | 0.55 (4.9) | 0.25 (0.3) |
| LinDA | Adaptive_NoWinsor | 0.05 (0.1) | 0.08 (0.1) | 0.24 (1.4) | 0.10 (0.2) | 0.45 (3.6) | 0.17 (0.3) |
| MaAsLin 2 | CLR_NONE | 0.13 (0.1) | 0.10 (0.0) | 0.23 (1.8) | 0.06 (0.1) | 0.47 (5.0) | 0.06 (0.0) |
| limma-voom | TMM_voom | 0.25 (0.5) | 0.25 (0.6) | 0.38 (2.5) | 0.22 (1.1) | 0.54 (4.3) | 0.22 (0.3) |
| LDM | PA | 0.05 (0.2) | 0.04 (0.2) | 0.20 (1.1) | 0.05 (0.0) | 0.47 (3.8) | 0.15 (0.4) |
| LDM | Arcsin | 0.13 (0.1) | 0.13 (0.1) | 0.30 (1.8) | 0.15 (0.2) | 0.54 (4.8) | 0.22 (0.4) |
| ZicoSeq | MultiLink | 0.08 (0.0) | 0.09 (0.0) | 0.24 (1.8) | 0.03 (0.1) | 0.41 (3.8) | 0.05 (0.0) |
| LinDA | Pseudo_1 | 0.02 (0.0) | 0.05 (0.2) | 0.18 (1.5) | 0.00 (0.0) | 0.42 (4.4) | 0.08 (0.2) |
| ZicoSeq | NoPosterior | 0.03 (0.0) | 0.04 (0.0) | 0.14 (0.9) | 0.07 (0.1) | 0.37 (3.5) | 0.08 (0.0) |
| MaAsLin 3 | TSS_LOG_Median * | 0.02 (0.1) | 0.02 (0.2) | 0.16 (1.4) | 0.00 (0.0) | 0.49 (6.2) | 0.01 (0.0) |
| LDM | Omni * | 0.15 (0.1) | 0.14 (0.1) | 0.30 (1.8) | 0.13 (0.1) | 0.56 (5.1) | 0.24 (0.2) |
| Wilcoxon | CLR | 0.14 (0.1) | | 0.22 (1.6) | | 0.43 (3.1) | |
| limma-voom | TMMwsp_voom | 0.15 (0.4) | 0.13 (0.2) | 0.38 (2.4) | 0.22 (0.4) | 0.52 (4.6) | 0.19 (0.2) |
| MaAsLin 3 | CLR_NONE | 0.03 (0.2) | 0.03 (0.2) | 0.22 (2.0) | 0.06 (0.2) | 0.53 (6.2) | 0.03 (0.0) |
| MaAsLin 3 | CLR_NONE_Median | 0.01 (0.1) | 0.01 (0.1) | 0.18 (1.8) | 0.00 (0.0) | 0.49 (6.2) | 0.02 (0.0) |
| MaAsLin 3 | TSS_LOG | 0.11 (0.2) | 0.11 (0.3) | 0.36 (2.4) | 0.20 (0.6) | 0.59 (6.4) | 0.22 (0.7) |
| DESeq2 | LRT_poscounts | 0.23 (0.8) | 0.28 (0.2) | 0.44 (3.9) | 0.27 (0.1) | 0.56 (5.3) | 0.30 (0.2) |
| DESeq2 | Wald_poscounts * | 0.22 (0.7) | 0.26 (0.3) | 0.44 (4.0) | 0.29 (0.1) | 0.56 (5.4) | 0.31 (0.3) |
| ZicoSeq | Default * | 0.14 (0.1) | 0.06 (0.0) | 0.19 (1.2) | 0.02 (0.1) | 0.42 (3.8) | 0.04 (0.0) |
| Wilcoxon | Rarefied | 0.09 (0.1) | | 0.26 (1.2) | | 0.48 (3.8) | |
| edgeR | TMM_LRT | 0.40 (1.9) | 0.27 (0.3) | 0.53 (4.2) | 0.35 (0.3) | 0.61 (6.4) | 0.31 (0.4) |
| LDM | Freq | 0.13 (0.1) | 0.12 (0.0) | 0.26 (1.5) | 0.12 (0.1) | 0.54 (4.8) | 0.19 (0.1) |
| ALDEx2 | Wilcox_128 | 0.02 (0.0) | | 0.14 (1.1) | | 0.26 (2.2) | |
| edgeR | TMM_QL * | 0.58 (3.2) | 0.49 (0.8) | 0.65 (5.4) | 0.50 (0.8) | 0.70 (7.2) | 0.50 (1.1) |
| ANCOM-BC2 | BH | 0.00 (0.0) | 0.00 (0.0) | 0.08 (0.7) | 0.00 (0.0) | 0.29 (2.8) | 0.00 (0.0) |
| ALDEx2 | Welch_128 * | 0.00 (0.0) | 0.00 (0.0) | 0.10 (0.8) | 0.00 (0.0) | 0.27 (2.2) | 0.00 (0.0) |
| ALDEx2 | Welch_32 | 0.01 (0.0) | 0.00 (0.0) | 0.10 (0.8) | 0.00 (0.0) | 0.26 (2.1) | 0.00 (0.0) |
| ALDEx2 | Welch_128_IQLR | 0.00 (0.0) | | 0.07 (0.6) | | 0.26 (2.2) | |
| ANCOM-BC2 | BH_StrucZero | 0.00 (0.0) | 0.00 (0.0) | 0.07 (0.6) | 0.00 (0.0) | 0.29 (2.8) | 0.00 (0.0) |
| MaAsLin 2 | TMM_NONE | 0.01 (0.0) | 0.01 (0.0) | 0.06 (0.2) | 0.03 (0.0) | 0.20 (1.3) | 0.01 (0.0) |
| ANCOM-BC2 | Holm * | 0.00 (0.0) | 0.00 (0.0) | 0.03 (0.2) | 0.00 (0.0) | 0.23 (1.9) | 0.00 (0.0) |
| ANCOM-BC2 | Holm_NoFilter | 0.42 (1.3) | 0.44 (1.1) | 0.63 (5.2) | 0.58 (1.1) | 0.68 (7.9) | 0.59 (1.5) |
| ANCOM-BC2 | BH_NoFilter | 0.61 (2.1) | 0.63 (1.9) | 0.72 (6.5) | 0.69 (1.6) | 0.75 (7.9) | 0.70 (2.2) |
| ***OTU level*** | | | | | | | |
| LinDA | Pseudo_0.1 | 0.04 (1.7) | 0.04 (1.6) | 0.07 (6.3) | 0.05 (0.8) | 0.16 (29.9) | 0.06 (1.3) |
| MaAsLin 2 | TMM_LOG | 0.04 (1.3) | 0.03 (0.8) | 0.08 (8.4) | 0.04 (0.7) | 0.22 (48.7) | 0.04 (0.8) |
| LinDA | Imputation | 0.09 (2.8) | 0.09 (2.1) | 0.13 (8.7) | 0.11 (2.1) | 0.23 (35.0) | 0.11 (2.6) |

| Tool | Configuration | r 0, unadjusted | r 0, adjusted | r 0.3, unadjusted | r 0.3, adjusted | r 0.6, unadjusted | r 0.6, adjusted |
|---|---|---|---|---|---|---|---|
| MaAsLin 3 | TSS_PLOG_Median | 0.05 (2.2) | 0.02 (1.1) | 0.07 (4.8) | 0.04 (0.8) | 0.14 (19.4) | 0.04 (0.6) |
| LinDA | Pseudo_0.5 | 0.04 (1.4) | 0.04 (1.4) | 0.08 (9.2) | 0.05 (0.9) | 0.23 (56.7) | 0.06 (0.8) |
| MaAsLin 3 | TSS_PLOG | 0.05 (2.1) | 0.03 (1.2) | 0.10 (6.6) | 0.05 (1.0) | 0.16 (22.1) | 0.06 (1.4) |
| MaAsLin 2 | TSS_LOG * | 0.08 (2.2) | 0.05 (1.3) | 0.12 (8.2) | 0.07 (1.7) | 0.22 (33.9) | 0.08 (1.9) |
| Wilcoxon | TSS | 0.06 (1.9) | | 0.10 (4.5) | | 0.14 (14.6) | |
| LinDA | Adaptive * | 0.09 (2.8) | 0.09 (2.1) | 0.13 (8.7) | 0.10 (1.8) | 0.23 (35.0) | 0.11 (2.6) |
| LDM | CLR | 0.03 (1.7) | 0.03 (1.5) | 0.07 (9.8) | 0.05 (1.0) | 0.24 (62.6) | 0.06 (0.2) |
| LDM | Omni3 | 0.37 (12.3) | 0.35 (10.7) | 0.40 (37.9) | 0.35 (12.6) | 0.54 (124.8) | 0.33 (10.2) |
| LinDA | Adaptive_NoWinsor | 0.05 (2.1) | 0.05 (1.9) | 0.09 (7.1) | 0.07 (1.2) | 0.20 (33.6) | 0.08 (1.6) |
| MaAsLin 2 | CLR_NONE | 0.32 (6.3) | 0.27 (3.7) | 0.31 (23.8) | 0.24 (2.8) | 0.46 (97.8) | 0.16 (0.9) |
| limma-voom | TMM_voom | 0.56 (65.2) | 0.56 (78.7) | 0.61 (85.2) | 0.60 (80.0) | 0.74 (130.7) | 0.56 (73.8) |
| LDM | PA | 0.38 (13.3) | 0.36 (12.8) | 0.40 (37.8) | 0.36 (11.9) | 0.53 (110.5) | 0.35 (14.3) |
| LDM | Arcsin | 0.23 (6.6) | 0.22 (6.1) | 0.29 (26.0) | 0.24 (6.2) | 0.44 (94.8) | 0.24 (3.2) |
| ZicoSeq | MultiLink | 0.06 (3.2) | 0.05 (2.7) | 0.15 (22.0) | 0.09 (2.8) | 0.39 (108.1) | 0.12 (2.1) |
| LinDA | Pseudo_1 | 0.03 (1.1) | 0.03 (1.0) | 0.08 (10.4) | 0.05 (0.6) | 0.26 (70.5) | 0.05 (0.5) |
| ZicoSeq | NoPosterior | 0.04 (1.2) | 0.04 (1.5) | 0.09 (10.8) | 0.07 (1.3) | 0.27 (66.9) | 0.08 (0.7) |
| MaAsLin 3 | TSS_LOG_Median * | 0.03 (0.8) | 0.03 (0.9) | 0.13 (25.2) | 0.04 (0.8) | 0.48 (170.9) | 0.05 (1.8) |
| LDM | Omni * | 0.35 (9.7) | 0.32 (7.8) | 0.38 (34.9) | 0.32 (9.3) | 0.53 (123.8) | 0.31 (4.3) |
| Wilcoxon | CLR | 0.72 (140.3) | | 0.63 (113.3) | | 0.73 (150.3) | |
| limma-voom | TMMwsp_voom | 0.80 (177.3) | 0.80 (191.6) | 0.78 (156.1) | 0.76 (164.7) | 0.87 (169.2) | 0.80 (150.0) |
| MaAsLin 3 | CLR_NONE | 0.23 (2.2) | 0.23 (7.6) | 0.30 (35.4) | 0.21 (7.1) | 0.56 (179.8) | 0.16 (5.4) |
| MaAsLin 3 | CLR_NONE_Median | 0.02 (0.7) | 0.03 (1.1) | 0.12 (24.1) | 0.04 (1.1) | 0.48 (180.2) | 0.06 (2.0) |
| MaAsLin 3 | TSS_LOG | 0.45 (12.6) | 0.45 (39.4) | 0.46 (54.0) | 0.39 (42.1) | 0.66 (168.9) | 0.37 (35.2) |
| DESeq2 | LRT_poscounts | 0.20 (18.7) | 0.17 (2.4) | 0.33 (57.8) | 0.17 (2.4) | 0.52 (153.8) | 0.18 (2.6) |
| DESeq2 | Wald_poscounts * | 0.21 (18.4) | 0.20 (5.1) | 0.35 (60.5) | 0.19 (3.5) | 0.53 (148.5) | 0.19 (4.1) |
| ZicoSeq | Default * | 0.06 (2.9) | 0.06 (2.7) | 0.15 (20.6) | 0.09 (2.2) | 0.36 (94.8) | 0.11 (1.3) |
| Wilcoxon | Rarefied | 0.14 (2.5) | | 0.19 (14.8) | | 0.34 (56.6) | |
| edgeR | TMM_LRT | 0.33 (59.6) | 0.20 (7.1) | 0.42 (107.6) | 0.21 (5.9) | 0.56 (213.6) | 0.22 (5.1) |
| LDM | Freq | 0.30 (5.5) | 0.28 (4.7) | 0.34 (26.1) | 0.28 (5.9) | 0.48 (105.4) | 0.27 (1.8) |
| ALDEx2 | Wilcox_128 | 0.00 (0.0) | | 0.00 (0.8) | | 0.05 (7.2) | |
| edgeR | TMM_QL * | 0.51 (109.5) | 0.37 (20.4) | 0.56 (158.8) | 0.38 (17.8) | 0.64 (239.6) | 0.38 (15.6) |
| ANCOM-BC2 | BH | 0.00 (0.3) | 0.00 (0.1) | 0.02 (2.9) | 0.00 (0.2) | 0.10 (20.1) | 0.01 (0.2) |
| ALDEx2 | Welch_128 * | 0.00 (0.0) | 0.00 (0.0) | 0.00 (0.5) | 0.00 (0.0) | 0.04 (5.4) | 0.00 (0.0) |
| ALDEx2 | Welch_32 | 0.00 (0.0) | 0.00 (0.0) | 0.00 (0.5) | 0.00 (0.0) | 0.04 (5.8) | 0.00 (0.0) |
| ALDEx2 | Welch_128_IQLR | 0.00 (0.0) | | 0.00 (0.4) | | 0.04 (5.3) | |
| ANCOM-BC2 | BH_StrucZero | 0.00 (0.2) | 0.00 (0.1) | 0.02 (2.4) | 0.00 (0.2) | 0.10 (19.5) | 0.01 (0.2) |
| MaAsLin 2 | TMM_NONE | 0.02 (0.1) | 0.02 (0.1) | 0.03 (0.8) | 0.03 (0.0) | 0.10 (14.7) | 0.03 (0.2) |
| ANCOM-BC2 | Holm * | 0.00 (0.0) | 0.00 (0.0) | 0.00 (0.1) | 0.00 (0.0) | 0.02 (3.0) | 0.00 (0.0) |
| ANCOM-BC2 | Holm_NoFilter | 0.60 (73.0) | 0.59 (37.8) | 0.65 (161.5) | 0.65 (51.0) | 0.71 (265.2) | 0.73 (66.2) |
| ANCOM-BC2 | BH_NoFilter | 0.83 (152.2) | 0.83 (122.6) | 0.84 (231.2) | 0.85 (137.3) | 0.84 (268.2) | 0.86 (147.5) |

**Table S16. Component tests of MaAsLin 3.** MCC, recall and FDR of the abundance, prevalence and joint tests of each MaAsLin 3 configuration, with 95% confidence intervals across replicates. The pseudo-log configurations have an abundance test only.

| Configuration [test] | MCC | Recall | FDR |
|---|---|---|---|
| ***Semi-synthetic, genus level, joint signal*** | | | |
| CLR_NONE [abundance] | 0.635 [0.541, 0.728] | 0.508 | 0.053 [0.027, 0.078] |
| CLR_NONE [joint] | 0.710 [0.684, 0.735] | 0.612 | 0.051 [0.024, 0.078] |
| CLR_NONE [prevalence] | 0.496 [0.448, 0.543] | 0.354 | 0.037 [0.000, 0.088] |
| CLR_NONE_Median [abundance] | 0.630 [0.562, 0.699] | 0.484 | 0.012 [0.000, 0.026] |
| CLR_NONE_Median [joint] | 0.707 [0.671, 0.743] | 0.599 | 0.017 [0.000, 0.040] |
| CLR_NONE_Median [prevalence] | 0.492 [0.449, 0.536] | 0.349 | 0.032 [0.000, 0.070] |
| TSS_LOG [abundance] | 0.570 [0.445, 0.695] | 0.510 | 0.129 [0.007, 0.251] |
| TSS_LOG [joint] | 0.647 [0.537, 0.757] | 0.622 | 0.124 [0.019, 0.228] |
| TSS_LOG [prevalence] | 0.499 [0.438, 0.561] | 0.362 | 0.018 [0.000, 0.060] |
| TSS_LOG_Median [abundance] | 0.629 [0.600, 0.658] | 0.487 | 0.025 [0.012, 0.039] |
| TSS_LOG_Median [joint] | 0.703 [0.695, 0.712] | 0.596 | 0.030 [0.012, 0.047] |
| TSS_LOG_Median [prevalence] | 0.493 [0.409, 0.576] | 0.352 | 0.007 [0.000, 0.021] |
| TSS_PLOG [abundance] | 0.710 [0.622, 0.799] | 0.680 | 0.095 [0.008, 0.182] |
| TSS_PLOG_Median [abundance] | 0.730 [0.644, 0.816] | 0.661 | 0.049 [0.000, 0.118] |
| ***Semi-synthetic, genus level, prevalence-only signal*** | | | |
| CLR_NONE [abundance] | 0.002 [−0.025, 0.030] | 0.003 | 0.125 [0.000, 0.296] |
| CLR_NONE [joint] | 0.446 [0.393, 0.498] | 0.328 | 0.024 [0.000, 0.061] |
| CLR_NONE [prevalence] | 0.455 [0.413, 0.498] | 0.328 | 0.003 [0.000, 0.012] |
| CLR_NONE_Median [abundance] | −0.003 [−0.007, 0.001] | 0.000 | 0.083 [0.000, 0.192] |
| CLR_NONE_Median [joint] | 0.448 [0.399, 0.498] | 0.326 | 0.017 [0.000, 0.035] |
| CLR_NONE_Median [prevalence] | 0.455 [0.413, 0.497] | 0.326 | 0.000 [0.000, 0.000] |
| TSS_LOG [abundance] | −0.006 [−0.007, −0.005] | 0.000 | 0.167 [0.167, 0.167] |
| TSS_LOG [joint] | 0.493 [0.452, 0.533] | 0.393 | 0.049 [0.000, 0.099] |
| TSS_LOG [prevalence] | 0.521 [0.471, 0.570] | 0.409 | 0.000 [0.000, 0.000] |
| TSS_LOG_Median [abundance] | −0.004 [−0.008, 0.001] | 0.000 | 0.104 [0.000, 0.231] |
| TSS_LOG_Median [joint] | 0.494 [0.455, 0.533] | 0.396 | 0.033 [0.002, 0.063] |
| TSS_LOG_Median [prevalence] | 0.523 [0.471, 0.575] | 0.411 | 0.000 [0.000, 0.000] |
| TSS_PLOG [abundance] | 0.584 [0.460, 0.707] | 0.495 | 0.028 [0.007, 0.049] |
| TSS_PLOG_Median [abundance] | 0.577 [0.462, 0.693] | 0.477 | 0.019 [0.000, 0.043] |
| ***Semi-synthetic, OTU level, joint signal*** | | | |
| CLR_NONE [abundance] | 0.529 [0.523, 0.535] | 0.417 | 0.167 [0.157, 0.176] |
| CLR_NONE [joint] | 0.604 [0.598, 0.610] | 0.500 | 0.121 [0.111, 0.130] |
| CLR_NONE [prevalence] | 0.542 [0.528, 0.556] | 0.410 | 0.019 [0.009, 0.029] |
| CLR_NONE_Median [abundance] | 0.602 [0.591, 0.613] | 0.424 | 0.039 [0.035, 0.043] |
| CLR_NONE_Median [joint] | 0.663 [0.651, 0.675] | 0.508 | 0.031 [0.023, 0.039] |
| CLR_NONE_Median [prevalence] | 0.542 [0.529, 0.556] | 0.405 | 0.025 [0.009, 0.040] |
| TSS_LOG [abundance] | 0.360 [0.340, 0.379] | 0.407 | 0.352 [0.321, 0.383] |
| TSS_LOG [joint] | 0.435 [0.416, 0.453] | 0.503 | 0.319 [0.287, 0.351] |
| TSS_LOG [prevalence] | 0.534 [0.512, 0.557] | 0.428 | 0.034 [0.012, 0.057] |
| TSS_LOG_Median [abundance] | 0.586 [0.577, 0.596] | 0.409 | 0.030 [0.022, 0.038] |
| TSS_LOG_Median [joint] | 0.630 [0.618, 0.643] | 0.470 | 0.025 [0.017, 0.034] |
| TSS_LOG_Median [prevalence] | 0.531 [0.518, 0.543] | 0.396 | 0.021 [0.001, 0.041] |
| TSS_PLOG [abundance] | 0.679 [0.659, 0.698] | 0.604 | 0.078 [0.061, 0.095] |
| TSS_PLOG_Median [abundance] | 0.658 [0.647, 0.668] | 0.543 | 0.058 [0.048, 0.068] |
| ***Semi-synthetic, OTU level, prevalence-only signal*** | | | |
| CLR_NONE [abundance] | 0.001 [−0.010, 0.011] | 0.001 | 0.547 [0.377, 0.718] |
| CLR_NONE [joint] | 0.445 [0.430, 0.461] | 0.340 | 0.024 [0.018, 0.030] |

| Configuration [test] | MCC | Recall | FDR |
|---|---|---|---|
| CLR_NONE [prevalence] | 0.451 [0.441, 0.461] | 0.339 | 0.005 [0.003, 0.008] |
| CLR_NONE_Median [abundance] | 0.002 [−0.010, 0.014] | 0.001 | 0.534 [0.322, 0.747] |
| CLR_NONE_Median [joint] | 0.443 [0.427, 0.460] | 0.339 | 0.024 [0.018, 0.030] |
| CLR_NONE_Median [prevalence] | 0.451 [0.440, 0.461] | 0.339 | 0.005 [0.003, 0.008] |
| TSS_LOG [abundance] | 0.010 [−0.004, 0.025] | 0.003 | 0.497 [0.379, 0.614] |
| TSS_LOG [joint] | 0.436 [0.420, 0.452] | 0.334 | 0.041 [0.000, 0.103] |
| TSS_LOG [prevalence] | 0.443 [0.426, 0.459] | 0.335 | 0.005 [0.001, 0.008] |
| TSS_LOG_Median [abundance] | 0.005 [−0.008, 0.019] | 0.002 | 0.513 [0.365, 0.661] |
| TSS_LOG_Median [joint] | 0.436 [0.420, 0.451] | 0.334 | 0.045 [0.000, 0.124] |
| TSS_LOG_Median [prevalence] | 0.442 [0.426, 0.457] | 0.334 | 0.005 [0.001, 0.008] |
| TSS_PLOG [abundance] | 0.562 [0.535, 0.590] | 0.457 | 0.026 [0.015, 0.038] |
| TSS_PLOG_Median [abundance] | 0.562 [0.531, 0.593] | 0.452 | 0.023 [0.014, 0.031] |
| ***Simulated, genus level, joint signal*** | | | |
| CLR_NONE [abundance] | 0.544 [0.466, 0.622] | 0.444 | 0.159 [0.113, 0.205] |
| CLR_NONE [joint] | 0.643 [0.566, 0.721] | 0.580 | 0.134 [0.075, 0.193] |
| CLR_NONE [prevalence] | 0.556 [0.514, 0.598] | 0.447 | 0.053 [0.026, 0.079] |
| CLR_NONE_Median [abundance] | 0.583 [0.489, 0.676] | 0.430 | 0.065 [0.034, 0.096] |
| CLR_NONE_Median [joint] | 0.692 [0.623, 0.761] | 0.583 | 0.055 [0.028, 0.082] |
| CLR_NONE_Median [prevalence] | 0.558 [0.526, 0.591] | 0.444 | 0.045 [0.010, 0.079] |
| TSS_LOG [abundance] | 0.505 [0.429, 0.581] | 0.416 | 0.173 [0.073, 0.273] |
| TSS_LOG [joint] | 0.650 [0.575, 0.725] | 0.613 | 0.147 [0.046, 0.247] |
| TSS_LOG [prevalence] | 0.579 [0.569, 0.590] | 0.484 | 0.052 [0.022, 0.081] |
| TSS_LOG_Median [abundance] | 0.568 [0.482, 0.654] | 0.416 | 0.059 [0.019, 0.099] |
| TSS_LOG_Median [joint] | 0.713 [0.661, 0.765] | 0.614 | 0.047 [0.008, 0.085] |
| TSS_LOG_Median [prevalence] | 0.580 [0.557, 0.602] | 0.478 | 0.037 [0.001, 0.072] |
| TSS_PLOG [abundance] | 0.748 [0.691, 0.805] | 0.742 | 0.129 [0.066, 0.192] |
| TSS_PLOG_Median [abundance] | 0.742 [0.698, 0.787] | 0.672 | 0.066 [0.016, 0.117] |
| ***Simulated, genus level, prevalence-only signal*** | | | |
| CLR_NONE [abundance] | 0.001 [−0.012, 0.014] | 0.002 | 0.075 [0.000, 0.155] |
| CLR_NONE [joint] | 0.411 [0.379, 0.444] | 0.281 | 0.046 [0.017, 0.074] |
| CLR_NONE [prevalence] | 0.414 [0.382, 0.446] | 0.280 | 0.008 [0.000, 0.017] |
| CLR_NONE_Median [abundance] | −0.004 [−0.006, −0.001] | 0.000 | 0.100 [0.035, 0.165] |
| CLR_NONE_Median [joint] | 0.405 [0.368, 0.441] | 0.280 | 0.055 [0.000, 0.109] |
| CLR_NONE_Median [prevalence] | 0.414 [0.382, 0.446] | 0.280 | 0.008 [0.000, 0.017] |
| TSS_LOG [abundance] | 0.001 [−0.014, 0.016] | 0.002 | 0.087 [0.048, 0.127] |
| TSS_LOG [joint] | 0.483 [0.438, 0.529] | 0.375 | 0.023 [0.000, 0.047] |
| TSS_LOG [prevalence] | 0.489 [0.443, 0.536] | 0.373 | 0.003 [0.000, 0.013] |
| TSS_LOG_Median [abundance] | −0.001 [−0.003, 0.000] | 0.000 | 0.037 [0.000, 0.077] |
| TSS_LOG_Median [joint] | 0.483 [0.432, 0.534] | 0.373 | 0.013 [0.000, 0.031] |
| TSS_LOG_Median [prevalence] | 0.489 [0.443, 0.536] | 0.373 | 0.003 [0.000, 0.013] |
| TSS_PLOG [abundance] | 0.533 [0.490, 0.576] | 0.428 | 0.064 [0.014, 0.114] |
| TSS_PLOG_Median [abundance] | 0.514 [0.477, 0.551] | 0.397 | 0.049 [0.000, 0.100] |

**Table S17. Effect-size estimation.** Estimated against implanted log2 fold change for all tested implanted features and for those called significant, which are selected for large estimates. Abundance-only signals change the non-zero counts only. Only configurations with a coefficient on an interpretable scale are included.

| Tool | Configuration | Median ratio of estimate to truth | Median relative error | Correct sign | Number of features |
|---|---|---|---|---|---|
| ***Called implanted features, genus level, abundance-only signal*** | | | | | |
| LinDA | Pseudo_0.1 | 0.891 | 0.180 | 1.000 | 59 |
| MaAsLin 2 | TMM_LOG | 0.867 | 0.155 | 1.000 | 66 |
| LinDA | Imputation | 0.831 | 0.195 | 1.000 | 71 |
| MaAsLin 3 | TSS_PLOG_Median | 0.923 | 0.132 | 1.000 | 55 |
| LinDA | Pseudo_0.5 | 0.760 | 0.284 | 1.000 | 80 |
| MaAsLin 3 | TSS_PLOG | 0.923 | 0.122 | 1.000 | 55 |
| MaAsLin 2 | TSS_LOG * | 0.900 | 0.146 | 1.000 | 64 |
| LinDA | Adaptive * | 0.803 | 0.203 | 1.000 | 74 |
| LinDA | Adaptive_NoWinsor | 0.824 | 0.202 | 1.000 | 77 |
| MaAsLin 2 | CLR_NONE | 0.694 | 0.313 | 1.000 | 89 |
| limma-voom | TMM_voom | 0.753 | 0.255 | 1.000 | 82 |
| LinDA | Pseudo_1 | 0.669 | 0.332 | 1.000 | 85 |
| MaAsLin 3 | TSS_LOG_Median * | 0.932 | 0.115 | 1.000 | 141 |
| limma-voom | TMMwsp_voom | 0.793 | 0.231 | 1.000 | 81 |
| MaAsLin 3 | CLR_NONE | 0.937 | 0.108 | 1.000 | 157 |
| MaAsLin 3 | CLR_NONE_Median | 0.937 | 0.108 | 1.000 | 152 |
| MaAsLin 3 | TSS_LOG | 0.925 | 0.143 | 1.000 | 155 |
| DESeq2 | LRT_poscounts | 1.004 | 0.148 | 1.000 | 132 |
| DESeq2 | Wald_poscounts * | 1.014 | 0.149 | 1.000 | 132 |
| edgeR | TMM_LRT | 0.967 | 0.180 | 1.000 | 152 |
| ALDEx2 | Wilcox_128 | 0.945 | 0.094 | 1.000 | 58 |
| edgeR | TMM_QL * | 0.956 | 0.186 | 1.000 | 185 |
| ANCOM-BC2 | BH | 0.970 | 0.078 | 1.000 | 47 |
| ALDEx2 | Welch_128 * | 0.959 | 0.087 | 1.000 | 48 |
| ALDEx2 | Welch_32 | 0.959 | 0.102 | 1.000 | 49 |
| ALDEx2 | Welch_128_IQLR | 0.980 | 0.105 | 1.000 | 45 |
| ANCOM-BC2 | BH_StrucZero | 0.970 | 0.078 | 1.000 | 47 |
| ANCOM-BC2 | Holm * | 0.982 | 0.082 | 1.000 | 41 |
| ANCOM-BC2 | Holm_NoFilter | 0.919 | 0.152 | 1.000 | 223 |
| ANCOM-BC2 | BH_NoFilter | 0.898 | 0.162 | 1.000 | 236 |
| ***Called implanted features, genus level, joint signal*** | | | | | |
| LinDA | Pseudo_0.1 | 1.226 | 0.376 | 1.000 | 212 |
| MaAsLin 2 | TMM_LOG | 1.138 | 0.308 | 1.000 | 211 |
| LinDA | Imputation | 1.111 | 0.290 | 1.000 | 212 |
| MaAsLin 3 | TSS_PLOG_Median | 1.335 | 0.417 | 1.000 | 192 |
| LinDA | Pseudo_0.5 | 1.019 | 0.234 | 1.000 | 207 |
| MaAsLin 3 | TSS_PLOG | 1.270 | 0.449 | 1.000 | 209 |
| MaAsLin 2 | TSS_LOG * | 1.088 | 0.292 | 1.000 | 215 |
| LinDA | Adaptive * | 1.024 | 0.259 | 1.000 | 209 |
| LinDA | Adaptive_NoWinsor | 1.030 | 0.269 | 1.000 | 209 |
| MaAsLin 2 | CLR_NONE | 0.860 | 0.256 | 1.000 | 215 |
| limma-voom | TMM_voom | 0.963 | 0.219 | 1.000 | 209 |
| LinDA | Pseudo_1 | 0.890 | 0.240 | 1.000 | 201 |
| MaAsLin 3 | TSS_LOG_Median * | 0.987 | 0.114 | 1.000 | 147 |
| limma-voom | TMMwsp_voom | 0.961 | 0.226 | 1.000 | 202 |
| MaAsLin 3 | CLR_NONE | 0.969 | 0.110 | 1.000 | 149 |
| MaAsLin 3 | CLR_NONE_Median | 0.971 | 0.110 | 1.000 | 146 |

| Tool | Configuration | Median ratio of estimate to truth | Median relative error | Correct sign | Number of features |
|---|---|---|---|---|---|
| MaAsLin 3 | TSS_LOG | 0.984 | 0.118 | 1.000 | 151 |
| DESeq2 | LRT_poscounts | 1.349 | 0.352 | 1.000 | 194 |
| DESeq2 | Wald_poscounts * | 1.350 | 0.353 | 1.000 | 191 |
| edgeR | TMM_LRT | 1.332 | 0.365 | 1.000 | 226 |
| ALDEx2 | Wilcox_128 | 1.281 | 0.295 | 1.000 | 154 |
| edgeR | TMM_QL * | 1.335 | 0.365 | 1.000 | 236 |
| ANCOM-BC2 | BH | 1.027 | 0.143 | 1.000 | 169 |
| ALDEx2 | Welch_128 * | 1.295 | 0.295 | 1.000 | 134 |
| ALDEx2 | Welch_32 | 1.315 | 0.318 | 1.000 | 136 |
| ALDEx2 | Welch_128_IQLR | 1.320 | 0.329 | 1.000 | 134 |
| ANCOM-BC2 | BH_StrucZero | 1.027 | 0.143 | 1.000 | 167 |
| ANCOM-BC2 | Holm * | 1.032 | 0.130 | 1.000 | 148 |
| ANCOM-BC2 | Holm_NoFilter | 1.017 | 0.161 | 1.000 | 198 |
| ANCOM-BC2 | BH_NoFilter | 1.001 | 0.178 | 0.995 | 217 |
| ***Called implanted features, OTU level, abundance-only signal*** | | | | | |
| LinDA | Pseudo_0.1 | 0.645 | 0.355 | 1.000 | 131 |
| MaAsLin 2 | TMM_LOG | 0.515 | 0.485 | 1.000 | 369 |
| LinDA | Imputation | 0.544 | 0.456 | 1.000 | 284 |
| MaAsLin 3 | TSS_PLOG_Median | 0.750 | 0.250 | 1.000 | 70 |
| LinDA | Pseudo_0.5 | 0.418 | 0.582 | 1.000 | 460 |
| MaAsLin 3 | TSS_PLOG | 0.722 | 0.278 | 1.000 | 81 |
| MaAsLin 2 | TSS_LOG * | 0.551 | 0.449 | 1.000 | 246 |
| LinDA | Adaptive * | 0.517 | 0.483 | 1.000 | 293 |
| LinDA | Adaptive_NoWinsor | 0.524 | 0.476 | 1.000 | 329 |
| MaAsLin 2 | CLR_NONE | 0.327 | 0.673 | 1.000 | 722 |
| limma-voom | TMM_voom | 0.276 | 0.724 | 0.999 | 949 |
| LinDA | Pseudo_1 | 0.324 | 0.676 | 1.000 | 692 |
| MaAsLin 3 | TSS_LOG_Median * | 0.872 | 0.172 | 1.000 | 3538 |
| limma-voom | TMMwsp_voom | 0.305 | 0.695 | 0.995 | 787 |
| MaAsLin 3 | CLR_NONE | 0.915 | 0.155 | 1.000 | 4623 |
| MaAsLin 3 | CLR_NONE_Median | 0.923 | 0.145 | 1.000 | 4467 |
| MaAsLin 3 | TSS_LOG | 0.849 | 0.195 | 1.000 | 3818 |
| DESeq2 | LRT_poscounts | 0.942 | 0.171 | 1.000 | 2427 |
| DESeq2 | Wald_poscounts * | 0.952 | 0.168 | 1.000 | 2434 |
| edgeR | TMM_LRT | 0.920 | 0.190 | 1.000 | 3190 |
| ALDEx2 | Wilcox_128 | 0.851 | 0.167 | 1.000 | 102 |
| edgeR | TMM_QL * | 0.905 | 0.205 | 1.000 | 4072 |
| ANCOM-BC2 | BH | 0.934 | 0.094 | 1.000 | 78 |
| ALDEx2 | Welch_128 * | 0.954 | 0.078 | 1.000 | 23 |
| ALDEx2 | Welch_32 | 0.937 | 0.091 | 1.000 | 24 |
| ALDEx2 | Welch_128_IQLR | 0.963 | 0.084 | 1.000 | 22 |
| ANCOM-BC2 | BH_StrucZero | 0.934 | 0.094 | 1.000 | 78 |
| ANCOM-BC2 | Holm * | 0.968 | 0.071 | 1.000 | 42 |
| ANCOM-BC2 | Holm_NoFilter | 0.854 | 0.210 | 1.000 | 6763 |
| ANCOM-BC2 | BH_NoFilter | 0.766 | 0.288 | 0.999 | 8143 |
| ***Called implanted features, OTU level, joint signal*** | | | | | |
| LinDA | Pseudo_0.1 | 1.035 | 0.371 | 1.000 | 6271 |
| MaAsLin 2 | TMM_LOG | 0.812 | 0.286 | 1.000 | 6295 |
| LinDA | Imputation | 0.965 | 0.224 | 0.998 | 5691 |
| MaAsLin 3 | TSS_PLOG_Median | 1.208 | 0.351 | 1.000 | 5476 |
| LinDA | Pseudo_0.5 | 0.707 | 0.313 | 1.000 | 6319 |

| Tool | Configuration | Median ratio of estimate to truth | Median relative error | Correct sign | Number of features |
|---|---|---|---|---|---|
| MaAsLin 3 | TSS_PLOG | 1.131 | 0.393 | 1.000 | 6212 |
| MaAsLin 2 | TSS_LOG * | 0.815 | 0.259 | 1.000 | 6352 |
| LinDA | Adaptive * | 0.918 | 0.240 | 0.998 | 5730 |
| LinDA | Adaptive_NoWinsor | 0.943 | 0.247 | 1.000 | 5715 |
| MaAsLin 2 | CLR_NONE | 0.499 | 0.501 | 1.000 | 7075 |
| limma-voom | TMM_voom | 0.548 | 0.454 | 1.000 | 7216 |
| LinDA | Pseudo_1 | 0.567 | 0.434 | 1.000 | 6286 |
| MaAsLin 3 | TSS_LOG_Median * | 0.829 | 0.214 | 1.000 | 4084 |
| limma-voom | TMMwsp_voom | 0.529 | 0.473 | 1.000 | 7257 |
| MaAsLin 3 | CLR_NONE | 0.892 | 0.136 | 1.000 | 4302 |
| MaAsLin 3 | CLR_NONE_Median | 0.899 | 0.131 | 1.000 | 4195 |
| MaAsLin 3 | TSS_LOG | 0.828 | 0.211 | 1.000 | 4484 |
| DESeq2 | LRT_poscounts | 1.388 | 0.398 | 1.000 | 5446 |
| DESeq2 | Wald_poscounts * | 1.404 | 0.406 | 1.000 | 5157 |
| edgeR | TMM_LRT | 1.306 | 0.385 | 1.000 | 7176 |
| ALDEx2 | Wilcox_128 | 1.247 | 0.263 | 1.000 | 3373 |
| edgeR | TMM_QL * | 1.304 | 0.381 | 1.000 | 7457 |
| ANCOM-BC2 | BH | 0.943 | 0.121 | 1.000 | 4778 |
| ALDEx2 | Welch_128 * | 1.272 | 0.279 | 1.000 | 2809 |
| ALDEx2 | Welch_32 | 1.270 | 0.279 | 1.000 | 2840 |
| ALDEx2 | Welch_128_IQLR | 1.315 | 0.320 | 1.000 | 2866 |
| ANCOM-BC2 | BH_StrucZero | 0.955 | 0.111 | 1.000 | 4456 |
| ANCOM-BC2 | Holm * | 0.975 | 0.092 | 1.000 | 3375 |
| ANCOM-BC2 | Holm_NoFilter | 0.937 | 0.139 | 1.000 | 5972 |
| ANCOM-BC2 | BH_NoFilter | 0.899 | 0.173 | 0.998 | 6813 |
| ***All tested implanted features, genus level, abundance-only signal*** | | | | | |
| LinDA | Pseudo_0.1 | 0.314 | 0.686 | 0.902 | 256 |
| MaAsLin 2 | TMM_LOG | 0.317 | 0.683 | 0.926 | 256 |
| LinDA | Imputation | 0.291 | 0.709 | 0.906 | 256 |
| MaAsLin 3 | TSS_PLOG_Median | 0.304 | 0.696 | 0.902 | 256 |
| LinDA | Pseudo_0.5 | 0.273 | 0.727 | 0.934 | 256 |
| MaAsLin 3 | TSS_PLOG | 0.304 | 0.696 | 0.902 | 256 |
| MaAsLin 2 | TSS_LOG * | 0.304 | 0.696 | 0.926 | 256 |
| LinDA | Adaptive * | 0.278 | 0.722 | 0.918 | 256 |
| LinDA | Adaptive_NoWinsor | 0.304 | 0.696 | 0.918 | 256 |
| MaAsLin 2 | CLR_NONE | 0.257 | 0.743 | 0.938 | 256 |
| limma-voom | TMM_voom | 0.276 | 0.724 | 0.922 | 256 |
| LinDA | Pseudo_1 | 0.249 | 0.751 | 0.941 | 256 |
| MaAsLin 3 | TSS_LOG_Median * | 0.841 | 0.196 | 0.996 | 256 |
| limma-voom | TMMwsp_voom | 0.269 | 0.731 | 0.922 | 256 |
| MaAsLin 3 | CLR_NONE | 0.908 | 0.144 | 0.995 | 221 |
| MaAsLin 3 | CLR_NONE_Median | 0.908 | 0.144 | 0.995 | 221 |
| MaAsLin 3 | TSS_LOG | 0.841 | 0.196 | 0.996 | 256 |
| DESeq2 | LRT_poscounts | 0.772 | 0.320 | 0.957 | 256 |
| DESeq2 | Wald_poscounts * | 0.772 | 0.320 | 0.957 | 256 |
| edgeR | TMM_LRT | 0.859 | 0.248 | 0.953 | 256 |
| ALDEx2 | Wilcox_128 | 0.312 | 0.688 | 0.891 | 256 |
| edgeR | TMM_QL * | 0.855 | 0.249 | 0.953 | 256 |
| ANCOM-BC2 | BH | 0.868 | 0.180 | 0.992 | 256 |
| ALDEx2 | Welch_128 * | 0.312 | 0.688 | 0.891 | 256 |
| ALDEx2 | Welch_32 | 0.312 | 0.692 | 0.887 | 256 |

| Tool | Configuration | Median ratio of estimate to truth | Median relative error | Correct sign | Number of features |
|---|---|---|---|---|---|
| ALDEx2 | Welch_128_IQLR | 0.292 | 0.710 | 0.902 | 256 |
| ANCOM-BC2 | BH_StrucZero | 0.871 | 0.177 | 0.992 | 253 |
| ANCOM-BC2 | Holm * | 0.868 | 0.180 | 0.992 | 256 |
| ANCOM-BC2 | Holm_NoFilter | 0.868 | 0.180 | 0.992 | 256 |
| ANCOM-BC2 | BH_NoFilter | 0.868 | 0.180 | 0.992 | 256 |
| ***All tested implanted features, genus level, joint signal*** | | | | | |
| LinDA | Pseudo_0.1 | 1.105 | 0.437 | 1.000 | 256 |
| MaAsLin 2 | TMM_LOG | 1.043 | 0.364 | 0.996 | 256 |
| LinDA | Imputation | 1.032 | 0.387 | 0.996 | 256 |
| MaAsLin 3 | TSS_PLOG_Median | 1.153 | 0.509 | 0.996 | 256 |
| LinDA | Pseudo_0.5 | 0.904 | 0.335 | 1.000 | 256 |
| MaAsLin 3 | TSS_PLOG | 1.153 | 0.509 | 0.996 | 256 |
| MaAsLin 2 | TSS_LOG * | 1.001 | 0.364 | 0.996 | 256 |
| LinDA | Adaptive * | 0.931 | 0.374 | 0.996 | 256 |
| LinDA | Adaptive_NoWinsor | 0.944 | 0.359 | 0.996 | 256 |
| MaAsLin 2 | CLR_NONE | 0.784 | 0.362 | 0.992 | 256 |
| limma-voom | TMM_voom | 0.853 | 0.328 | 0.977 | 256 |
| LinDA | Pseudo_1 | 0.779 | 0.381 | 1.000 | 256 |
| MaAsLin 3 | TSS_LOG_Median * | 0.900 | 0.192 | 0.969 | 254 |
| limma-voom | TMMwsp_voom | 0.859 | 0.312 | 0.980 | 256 |
| MaAsLin 3 | CLR_NONE | 0.931 | 0.149 | 0.991 | 211 |
| MaAsLin 3 | CLR_NONE_Median | 0.931 | 0.149 | 0.991 | 211 |
| MaAsLin 3 | TSS_LOG | 0.900 | 0.192 | 0.969 | 254 |
| DESeq2 | LRT_poscounts | 1.243 | 0.401 | 0.996 | 256 |
| DESeq2 | Wald_poscounts * | 1.243 | 0.401 | 0.996 | 256 |
| edgeR | TMM_LRT | 1.274 | 0.383 | 0.992 | 256 |
| ALDEx2 | Wilcox_128 | 1.069 | 0.377 | 0.996 | 256 |
| edgeR | TMM_QL * | 1.303 | 0.382 | 0.992 | 256 |
| ANCOM-BC2 | BH | 0.966 | 0.192 | 0.983 | 234 |
| ALDEx2 | Welch_128 * | 1.069 | 0.377 | 0.996 | 256 |
| ALDEx2 | Welch_32 | 1.070 | 0.377 | 0.996 | 256 |
| ALDEx2 | Welch_128_IQLR | 1.088 | 0.399 | 0.992 | 256 |
| ANCOM-BC2 | BH_StrucZero | 0.999 | 0.185 | 0.986 | 216 |
| ANCOM-BC2 | Holm * | 0.966 | 0.192 | 0.983 | 234 |
| ANCOM-BC2 | Holm_NoFilter | 0.966 | 0.192 | 0.983 | 234 |
| ANCOM-BC2 | BH_NoFilter | 0.966 | 0.192 | 0.983 | 234 |
| ***All tested implanted features, OTU level, abundance-only signal*** | | | | | |
| LinDA | Pseudo_0.1 | 0.116 | 0.884 | 0.778 | 8640 |
| MaAsLin 2 | TMM_LOG | 0.136 | 0.864 | 0.882 | 8640 |
| LinDA | Imputation | 0.122 | 0.878 | 0.809 | 8640 |
| MaAsLin 3 | TSS_PLOG_Median | 0.144 | 0.856 | 0.808 | 8640 |
| LinDA | Pseudo_0.5 | 0.096 | 0.904 | 0.835 | 8640 |
| MaAsLin 3 | TSS_PLOG | 0.144 | 0.856 | 0.808 | 8640 |
| MaAsLin 2 | TSS_LOG * | 0.140 | 0.860 | 0.866 | 8640 |
| LinDA | Adaptive * | 0.108 | 0.892 | 0.822 | 8640 |
| LinDA | Adaptive_NoWinsor | 0.128 | 0.872 | 0.858 | 8640 |
| MaAsLin 2 | CLR_NONE | 0.099 | 0.901 | 0.908 | 8640 |
| limma-voom | TMM_voom | 0.114 | 0.886 | 0.871 | 8640 |
| LinDA | Pseudo_1 | 0.082 | 0.918 | 0.859 | 8640 |
| MaAsLin 3 | TSS_LOG_Median * | 0.700 | 0.334 | 0.978 | 8640 |
| limma-voom | TMMwsp_voom | 0.115 | 0.885 | 0.852 | 8640 |

| Tool | Configuration | Median ratio of estimate to truth | Median relative error | Correct sign | Number of features |
|---|---|---|---|---|---|
| MaAsLin 3 | CLR_NONE | 0.753 | 0.300 | 0.994 | 8297 |
| MaAsLin 3 | CLR_NONE_Median | 0.753 | 0.300 | 0.994 | 8297 |
| MaAsLin 3 | TSS_LOG | 0.700 | 0.334 | 0.978 | 8640 |
| DESeq2 | LRT_poscounts | 0.599 | 0.434 | 0.949 | 8640 |
| DESeq2 | Wald_poscounts * | 0.599 | 0.434 | 0.949 | 8640 |
| edgeR | TMM_LRT | 0.660 | 0.391 | 0.953 | 8640 |
| ALDEx2 | Wilcox_128 | 0.119 | 0.881 | 0.815 | 8640 |
| edgeR | TMM_QL * | 0.664 | 0.388 | 0.953 | 8640 |
| ANCOM-BC2 | BH | 0.723 | 0.323 | 0.988 | 8640 |
| ALDEx2 | Welch_128 * | 0.119 | 0.881 | 0.815 | 8640 |
| ALDEx2 | Welch_32 | 0.120 | 0.880 | 0.810 | 8640 |
| ALDEx2 | Welch_128_IQLR | 0.117 | 0.883 | 0.807 | 8640 |
| ANCOM-BC2 | BH_StrucZero | 0.724 | 0.320 | 0.988 | 8475 |
| ANCOM-BC2 | Holm * | 0.723 | 0.323 | 0.988 | 8640 |
| ANCOM-BC2 | Holm_NoFilter | 0.723 | 0.323 | 0.988 | 8640 |
| ANCOM-BC2 | BH_NoFilter | 0.723 | 0.323 | 0.988 | 8640 |
| ***All tested implanted features, OTU level, joint signal*** | | | | | |
| LinDA | Pseudo_0.1 | 0.769 | 0.466 | 0.994 | 8640 |
| MaAsLin 2 | TMM_LOG | 0.594 | 0.432 | 0.999 | 8640 |
| LinDA | Imputation | 0.726 | 0.383 | 0.984 | 8640 |
| MaAsLin 3 | TSS_PLOG_Median | 0.902 | 0.464 | 0.999 | 8640 |
| LinDA | Pseudo_0.5 | 0.492 | 0.513 | 0.994 | 8640 |
| MaAsLin 3 | TSS_PLOG | 0.902 | 0.464 | 0.999 | 8640 |
| MaAsLin 2 | TSS_LOG * | 0.661 | 0.388 | 0.999 | 8640 |
| LinDA | Adaptive * | 0.618 | 0.437 | 0.985 | 8640 |
| LinDA | Adaptive_NoWinsor | 0.633 | 0.437 | 0.994 | 8640 |
| MaAsLin 2 | CLR_NONE | 0.394 | 0.606 | 0.999 | 8640 |
| limma-voom | TMM_voom | 0.477 | 0.523 | 0.998 | 8640 |
| LinDA | Pseudo_1 | 0.376 | 0.624 | 0.994 | 8640 |
| MaAsLin 3 | TSS_LOG_Median * | 0.744 | 0.287 | 0.973 | 8559 |
| limma-voom | TMMwsp_voom | 0.483 | 0.518 | 0.999 | 8640 |
| MaAsLin 3 | CLR_NONE | 0.773 | 0.252 | 0.991 | 8208 |
| MaAsLin 3 | CLR_NONE_Median | 0.773 | 0.252 | 0.991 | 8208 |
| MaAsLin 3 | TSS_LOG | 0.744 | 0.287 | 0.973 | 8559 |
| DESeq2 | LRT_poscounts | 1.184 | 0.442 | 1.000 | 8640 |
| DESeq2 | Wald_poscounts * | 1.184 | 0.442 | 1.000 | 8640 |
| edgeR | TMM_LRT | 1.224 | 0.386 | 0.999 | 8640 |
| ALDEx2 | Wilcox_128 | 0.745 | 0.436 | 0.998 | 8640 |
| edgeR | TMM_QL * | 1.225 | 0.385 | 0.999 | 8640 |
| ANCOM-BC2 | BH | 0.884 | 0.189 | 0.988 | 7109 |
| ALDEx2 | Welch_128 * | 0.745 | 0.436 | 0.998 | 8640 |
| ALDEx2 | Welch_32 | 0.746 | 0.435 | 0.997 | 8640 |
| ALDEx2 | Welch_128_IQLR | 0.752 | 0.471 | 0.997 | 8640 |
| ANCOM-BC2 | BH_StrucZero | 0.932 | 0.143 | 0.995 | 5979 |
| ANCOM-BC2 | Holm * | 0.884 | 0.189 | 0.988 | 7109 |
| ANCOM-BC2 | Holm_NoFilter | 0.884 | 0.189 | 0.988 | 7109 |
| ANCOM-BC2 | BH_NoFilter | 0.884 | 0.189 | 0.988 | 7109 |

**Table S18. Transfer of the configuration choice.** The configuration of each tool with the highest mean MCC on semi-synthetic stool genera, or at 200 samples, applied to other data or to 50 samples. Chosen in target is the configuration chosen on three replicates of the target data and scored on the fourth. limma-voom and the Wilcoxon test have no single default.

| Tool | Configuration transferred | MCC | FDR | MCC chosen in target | MCC default | Difference from chosen | Difference from default |
|---|---|---|---|---|---|---|---|
| ***Stool genus, semi-synthetic to semi-synthetic, OTU*** | | | | | | | |
| MaAsLin 2 | TMM_LOG | 0.613 | 0.030 | 0.629 | 0.605 | −0.016 | 0.008 |
| MaAsLin 3 | TSS_PLOG_Median | 0.610 | 0.041 | 0.633 | 0.533 | −0.023 | 0.077 |
| LinDA | Pseudo_0.1 | 0.630 | 0.038 | 0.641 | 0.591 | −0.010 | 0.039 |
| ZicoSeq | MultiLink | 0.604 | 0.061 | 0.650 | 0.600 | −0.045 | 0.004 |
| LDM | CLR | 0.609 | 0.035 | 0.609 | 0.551 | 0.001 | 0.059 |
| ALDEx2 | Wilcox_128 | 0.341 | 0.000 | 0.340 | 0.271 | 0.001 | 0.070 |
| ANCOM-BC2 | BH | 0.413 | 0.006 | 0.450 | 0.261 | −0.037 | 0.152 |
| DESeq2 | LRT_poscounts | 0.518 | 0.100 | 0.521 | 0.502 | −0.003 | 0.015 |
| edgeR | TMM_LRT | 0.503 | 0.388 | 0.520 | 0.482 | −0.017 | 0.021 |
| limma-voom | TMM_voom | 0.497 | 0.274 | 0.498 | | −0.000 | |
| Wilcoxon | TSS | 0.595 | 0.051 | 0.593 | | 0.002 | |
| ***Stool genus, semi-synthetic to simulated, genus*** | | | | | | | |
| MaAsLin 2 | TMM_LOG | 0.637 | 0.079 | 0.638 | 0.624 | −0.002 | 0.012 |
| MaAsLin 3 | TSS_PLOG_Median | 0.628 | 0.058 | 0.619 | 0.598 | 0.009 | 0.030 |
| LinDA | Pseudo_0.1 | 0.637 | 0.089 | 0.615 | 0.620 | 0.022 | 0.017 |
| ZicoSeq | MultiLink | 0.604 | 0.075 | 0.613 | 0.555 | −0.009 | 0.049 |
| LDM | CLR | 0.615 | 0.102 | 0.612 | 0.577 | 0.003 | 0.038 |
| ALDEx2 | Wilcox_128 | 0.435 | 0.045 | 0.432 | 0.419 | 0.004 | 0.017 |
| ANCOM-BC2 | BH | 0.393 | 0.006 | 0.383 | 0.339 | 0.010 | 0.055 |
| DESeq2 | LRT_poscounts | 0.458 | 0.231 | 0.460 | 0.453 | −0.002 | 0.005 |
| edgeR | TMM_LRT | 0.492 | 0.323 | 0.495 | 0.419 | −0.004 | 0.072 |
| limma-voom | TMM_voom | 0.550 | 0.149 | 0.542 | | 0.007 | |
| Wilcoxon | TSS | 0.625 | 0.104 | 0.614 | | 0.011 | |
| ***Stool genus, semi-synthetic to IBD semi-synthetic, genus*** | | | | | | | |
| MaAsLin 2 | TMM_LOG | 0.492 | 0.034 | 0.530 | 0.505 | −0.038 | −0.013 |
| MaAsLin 3 | TSS_PLOG_Median | 0.498 | 0.019 | 0.481 | 0.418 | 0.017 | 0.079 |
| LinDA | Pseudo_0.1 | 0.519 | 0.041 | 0.526 | 0.508 | −0.008 | 0.010 |
| ZicoSeq | MultiLink | 0.480 | 0.127 | 0.463 | 0.410 | 0.016 | 0.070 |
| LDM | CLR | 0.494 | 0.025 | 0.489 | 0.460 | 0.005 | 0.034 |
| ALDEx2 | Wilcox_128 | 0.384 | 0.000 | 0.372 | 0.322 | 0.011 | 0.062 |
| ANCOM-BC2 | BH | 0.284 | 0.005 | 0.304 | 0.231 | −0.020 | 0.053 |
| DESeq2 | LRT_poscounts | 0.429 | 0.168 | 0.447 | 0.456 | −0.018 | −0.027 |
| edgeR | TMM_LRT | 0.377 | 0.510 | 0.376 | 0.310 | 0.001 | 0.067 |
| limma-voom | TMM_voom | 0.469 | 0.215 | 0.446 | | 0.023 | |
| Wilcoxon | TSS | 0.473 | 0.022 | 0.473 | | −0.000 | |
| ***200 samples, genus to 50 samples, genus*** | | | | | | | |
| MaAsLin 2 | TMM_LOG | 0.493 | 0.000 | | 0.457 | | 0.036 |
| MaAsLin 3 | TSS_PLOG_Median | 0.449 | 0.000 | | 0.376 | | 0.073 |
| LinDA | Pseudo_0.1 | 0.477 | 0.041 | | 0.475 | | 0.002 |
| ZicoSeq | NoPosterior | 0.424 | 0.023 | | 0.386 | | 0.038 |
| LDM | CLR | 0.408 | 0.034 | | 0.425 | | −0.016 |
| ALDEx2 | Wilcox_128 | 0.307 | 0.000 | | 0.244 | | 0.063 |
| ANCOM-BC2 | BH | 0.283 | 0.000 | | 0.230 | | 0.053 |
| DESeq2 | LRT_poscounts | 0.417 | 0.072 | | 0.413 | | 0.004 |
| edgeR | TMM_LRT | 0.370 | 0.274 | | 0.350 | | 0.020 |

| Tool | Configuration transferred | MCC | FDR | MCC chosen in target | MCC default | Difference from chosen | Difference from default |
|---|---|---|---|---|---|---|---|
| limma-voom | TMM_voom | 0.439 | 0.072 | | | | |
| Wilcoxon | TSS | 0.453 | 0.005 | | | | |
| ***200 samples, OTU to 50 samples, OTU*** | | | | | | | |
| MaAsLin 2 | TMM_LOG | 0.390 | 0.013 | | 0.393 | | −0.003 |
| MaAsLin 3 | TSS_PLOG | 0.402 | 0.025 | | 0.306 | | 0.095 |
| LinDA | Pseudo_0.1 | 0.419 | 0.028 | | 0.387 | | 0.033 |
| ZicoSeq | NoPosterior | 0.444 | 0.049 | | 0.420 | | 0.024 |
| LDM | CLR | 0.389 | 0.025 | | 0.367 | | 0.022 |
| ALDEx2 | Wilcox_128 | 0.180 | 0.000 | | 0.120 | | 0.060 |
| ANCOM-BC2 | BH | 0.254 | 0.006 | | 0.141 | | 0.114 |
| DESeq2 | LRT_poscounts | 0.369 | 0.073 | | 0.367 | | 0.001 |
| edgeR | TMM_LRT | 0.288 | 0.569 | | 0.323 | | −0.035 |
| limma-voom | TMM_voom | 0.301 | 0.252 | | | | |
| Wilcoxon | TSS | 0.358 | 0.024 | | | | |

**Table S19. Semi-synthetic against simulated data.** The 24 genus-level scenarios common to both approaches. Mean MCC and FDR with 95% confidence intervals across replicates, and the difference (simulated minus semi-synthetic) with a Welch 95% interval. Null calls give the number of the 12 null datasets at 50, 100 and 200 samples with at least one call.

| Tool | Configuration | MCC semi-synthetic | MCC simulated | MCC difference | FDR semi-synthetic | FDR simulated | FDR difference | Null calls, semi-synthetic | Null calls, simulated |
|---|---|---|---|---|---|---|---|---|---|
| LinDA | Pseudo_0.1 | 0.675 [0.637, 0.714] | 0.540 [0.447, 0.633] | −0.135 [−0.223, −0.047] | 0.055 [0.034, 0.076] | 0.083 [0.006, 0.160] | 0.028 [−0.046, 0.102] | 1 | 2 |
| MaAsLin 2 | TMM_LOG | 0.675 [0.628, 0.723] | 0.528 [0.500, 0.555] | −0.148 [−0.193, −0.103] | 0.041 [0.029, 0.054] | 0.075 [0.049, 0.102] | 0.034 [0.009, 0.059] | 1 | 1 |
| LinDA | Imputation | 0.662 [0.624, 0.699] | 0.522 [0.426, 0.617] | −0.140 [−0.231, −0.050] | 0.059 [0.004, 0.114] | 0.106 [0.026, 0.186] | 0.047 [−0.030, 0.124] | 1 | 1 |
| MaAsLin 3 | TSS_PLOG_Median | 0.654 [0.596, 0.712] | 0.519 [0.452, 0.586] | −0.135 [−0.203, −0.067] | 0.034 [0.000, 0.080] | 0.062 [0.013, 0.111] | 0.029 [−0.023, 0.081] | 1 | 1 |
| LinDA | Pseudo_0.5 | 0.650 [0.625, 0.675] | 0.517 [0.441, 0.592] | −0.133 [−0.205, −0.061] | 0.048 [0.015, 0.082] | 0.100 [0.040, 0.160] | 0.052 [−0.005, 0.108] | 1 | 2 |
| MaAsLin 3 | TSS_PLOG | 0.647 [0.601, 0.693] | 0.531 [0.471, 0.591] | −0.116 [−0.176, −0.057] | 0.061 [0.009, 0.114] | 0.089 [0.041, 0.138] | 0.028 [−0.027, 0.083] | 1 | 1 |
| MaAsLin 2 | TSS_LOG * | 0.641 [0.604, 0.677] | 0.511 [0.444, 0.578] | −0.130 [−0.193, −0.067] | 0.064 [0.010, 0.119] | 0.090 [0.003, 0.177] | 0.026 [−0.057, 0.108] | 2 | 1 |
| Wilcoxon | TSS | 0.639 [0.598, 0.680] | 0.513 [0.439, 0.587] | −0.126 [−0.196, −0.056] | 0.068 [0.003, 0.133] | 0.093 [0.040, 0.146] | 0.025 [−0.040, 0.090] | 2 | 1 |
| LinDA | Adaptive * | 0.634 [0.611, 0.657] | 0.518 [0.443, 0.592] | −0.117 [−0.188, −0.046] | 0.059 [0.020, 0.098] | 0.098 [0.036, 0.161] | 0.040 [−0.020, 0.099] | 1 | 2 |
| LDM | CLR | 0.631 [0.623, 0.638] | 0.503 [0.432, 0.573] | −0.128 [−0.198, −0.058] | 0.035 [0.000, 0.075] | 0.105 [0.041, 0.170] | 0.070 [0.009, 0.131] | 0 | 1 |
| LDM | Omni3 | 0.629 [0.579, 0.679] | 0.491 [0.409, 0.573] | −0.138 [−0.216, −0.060] | 0.074 [0.029, 0.120] | 0.104 [0.032, 0.176] | 0.030 [−0.039, 0.098] | 2 | 0 |
| LinDA | Adaptive_NoWinsor | 0.626 [0.591, 0.661] | 0.516 [0.432, 0.601] | −0.110 [−0.190, −0.030] | 0.060 [0.015, 0.104] | 0.097 [0.032, 0.162] | 0.037 [−0.025, 0.100] | 1 | 2 |
| MaAsLin 2 | CLR_NONE | 0.613 [0.605, 0.620] | 0.479 [0.415, 0.542] | −0.134 [−0.197, −0.071] | 0.048 [0.014, 0.082] | 0.126 [0.068, 0.184] | 0.078 [0.023, 0.133] | 1 | 3 |
| limma-voom | TMM_voom | 0.612 [0.569, 0.655] | 0.453 [0.382, 0.524] | −0.159 [−0.226, −0.091] | 0.094 [0.049, 0.139] | 0.154 [0.094, 0.213] | 0.060 [0.001, 0.118] | 1 | 1 |
| LDM | PA | 0.606 [0.552, 0.659] | 0.467 [0.401, 0.533] | −0.138 [−0.204, −0.072] | 0.056 [0.004, 0.108] | 0.082 [0.010, 0.154] | 0.026 [−0.044, 0.096] | 1 | 0 |
| LDM | Arcsin | 0.604 [0.554, 0.655] | 0.465 [0.389, 0.541] | −0.139 [−0.212, −0.067] | 0.081 [0.045, 0.116] | 0.115 [0.031, 0.199] | 0.034 [−0.045, 0.114] | 2 | 0 |
| ZicoSeq | MultiLink | 0.604 [0.571, 0.636] | 0.512 [0.443, 0.580] | −0.092 [−0.156, −0.028] | 0.072 [0.050, 0.094] | 0.094 [0.067, 0.120] | 0.022 [−0.005, 0.048] | 0 | 0 |
| LinDA | Pseudo_1 | 0.603 [0.586, 0.620] | 0.500 [0.416, 0.584] | −0.103 [−0.185, −0.021] | 0.045 [0.011, 0.079] | 0.102 [0.023, 0.181] | 0.057 [−0.017, 0.132] | 1 | 2 |
| ZicoSeq | NoPosterior | 0.602 [0.567, 0.637] | 0.486 [0.467, 0.505] | −0.116 [−0.150, −0.083] | 0.042 [0.032, 0.052] | 0.082 [0.017, 0.146] | 0.040 [−0.024, 0.104] | 0 | 0 |
| MaAsLin 3 | TSS_LOG_Median * | 0.598 [0.580, 0.617] | 0.471 [0.401, 0.542] | −0.127 [−0.196, −0.059] | 0.031 [0.009, 0.053] | 0.034 [0.000, 0.080] | 0.003 [−0.040, 0.046] | 1 | 1 |
| LDM | Omni * | 0.595 [0.529, 0.660] | 0.464 [0.389, 0.539] | −0.130 [−0.208, −0.053] | 0.090 [0.063, 0.118] | 0.114 [0.026, 0.202] | 0.024 [−0.060, 0.108] | 2 | 0 |
| Wilcoxon | CLR | 0.590 [0.545, 0.635] | 0.444 [0.391, 0.496] | −0.147 [−0.200, −0.093] | 0.059 [0.037, 0.080] | 0.119 [0.087, 0.151] | 0.060 [0.030, 0.091] | 1 | 1 |
| limma-voom | TMMwsp_voom | 0.578 [0.564, 0.593] | 0.444 [0.365, 0.522] | −0.135 [−0.212, −0.058] | 0.085 [0.000, 0.189] | 0.199 [0.107, 0.292] | 0.114 [0.007, 0.221] | 0 | 1 |
| MaAsLin 3 | CLR_NONE | 0.578 [0.548, 0.607] | 0.429 [0.377, 0.481] | −0.149 [−0.198, −0.100] | 0.038 [0.017, 0.059] | 0.078 [0.025, 0.132] | 0.041 [−0.009, 0.091] | 0 | 2 |
| MaAsLin 3 | CLR_NONE_Median | 0.577 [0.541, 0.614] | 0.444 [0.381, 0.507] | −0.133 [−0.193, −0.074] | 0.017 [0.000, 0.034] | 0.059 [0.000, 0.131] | 0.042 [−0.028, 0.112] | 0 | 1 |
| MaAsLin 3 | TSS_LOG | 0.570 [0.508, 0.632] | 0.438 [0.368, 0.507] | −0.132 [−0.204, −0.060] | 0.086 [0.040, 0.132] | 0.089 [0.004, 0.174] | 0.003 [−0.077, 0.083] | 1 | 1 |
| DESeq2 | LRT_poscounts | 0.540 [0.509, 0.571] | 0.397 [0.373, 0.422] | −0.143 [−0.173, −0.112] | 0.136 [0.071, 0.201] | 0.211 [0.072, 0.349] | 0.075 [−0.056, 0.205] | 5 | 1 |
| DESeq2 | Wald_poscounts * | 0.536 [0.499, 0.573] | 0.400 [0.355, 0.445] | −0.135 [−0.181, −0.090] | 0.163 [0.084, 0.242] | 0.248 [0.095, 0.400] | 0.084 [−0.059, 0.227] | 5 | 4 |
| ZicoSeq | Default * | 0.518 [0.458, 0.579] | 0.455 [0.394, 0.516] | −0.064 [−0.130, 0.002] | 0.116 [0.075, 0.158] | 0.073 [0.000, 0.156] | −0.043 [−0.121, 0.035] | 0 | 2 |
| Wilcoxon | Rarefied | 0.505 [0.483, 0.526] | 0.420 [0.340, 0.499] | −0.085 [−0.162, −0.008] | 0.080 [0.056, 0.104] | 0.088 [0.000, 0.189] | 0.009 [−0.090, 0.107] | 1 | 0 |
| edgeR | TMM_LRT | 0.496 [0.468, 0.524] | 0.423 [0.375, 0.471] | −0.073 [−0.118, −0.028] | 0.341 [0.308, 0.374] | 0.285 [0.211, 0.359] | −0.056 [−0.126, 0.014] | 9 | 8 |
| LDM | Freq | 0.494 [0.470, 0.518] | 0.416 [0.362, 0.469] | −0.079 [−0.129, −0.028] | 0.081 [0.028, 0.134] | 0.108 [0.000, 0.217] | 0.027 [−0.075, 0.129] | 2 | 0 |
| ALDEx2 | Wilcox_128 | 0.461 [0.411, 0.511] | 0.339 [0.279, 0.398] | −0.122 [−0.183, −0.062] | 0.005 [0.000, 0.016] | 0.038 [0.000, 0.087] | 0.033 [−0.015, 0.081] | 1 | 0 |
| edgeR | TMM_QL * | 0.434 [0.394, 0.474] | 0.373 [0.323, 0.423] | −0.061 [−0.111, −0.011] | 0.571 [0.527, 0.615] | 0.585 [0.474, 0.696] | 0.014 [−0.091, 0.119] | 12 | 12 |
| ANCOM-BC2 | BH | 0.420 [0.355, 0.485] | 0.321 [0.291, 0.350] | −0.099 [−0.160, −0.038] | 0.005 [0.000, 0.016] | 0.008 [0.000, 0.023] | 0.003 [−0.011, 0.017] | 1 | 0 |
| ALDEx2 | Welch_128 * | 0.411 [0.372, 0.450] | 0.313 [0.269, 0.358] | −0.098 [−0.143, −0.052] | 0.000 [0.000, 0.000] | 0.025 [0.000, 0.059] | 0.025 [−0.009, 0.059] | 0 | 0 |
| ALDEx2 | Welch_32 | 0.407 [0.377, 0.437] | 0.318 [0.270, 0.366] | −0.089 [−0.134, −0.043] | 0.000 [0.000, 0.000] | 0.026 [0.000, 0.061] | 0.026 [−0.009, 0.061] | 0 | 0 |
| ALDEx2 | Welch_128_IQLR | 0.406 [0.368, 0.444] | 0.312 [0.270, 0.354] | −0.094 [−0.138, −0.051] | 0.000 [0.000, 0.000] | 0.029 [0.000, 0.086] | 0.029 [−0.027, 0.086] | 0 | 0 |
| ANCOM-BC2 | BH_StrucZero | 0.404 [0.355, 0.453] | 0.276 [0.241, 0.310] | −0.128 [−0.176, −0.081] | 0.005 [0.000, 0.016] | 0.008 [0.000, 0.023] | 0.003 [−0.011, 0.017] | 1 | 0 |
| MaAsLin 2 | TMM_NONE | 0.366 [0.360, 0.372] | 0.272 [0.225, 0.320] | −0.094 [−0.141, −0.047] | 0.025 [0.000, 0.055] | 0.030 [0.006, 0.055] | 0.006 [−0.024, 0.036] | 1 | 0 |
| ANCOM-BC2 | Holm * | 0.353 [0.328, 0.379] | 0.262 [0.235, 0.290] | −0.091 [−0.120, −0.062] | 0.000 [0.000, 0.000] | 0.000 [0.000, 0.000] | 0.000 [0.000, 0.000] | 1 | 0 |
| ANCOM-BC2 | Holm_NoFilter | 0.309 [0.261, 0.357] | 0.280 [0.236, 0.324] | −0.028 [−0.079, 0.022] | 0.647 [0.573, 0.722] | 0.647 [0.573, 0.721] | −0.000 [−0.081, 0.081] | 12 | 12 |
| ANCOM-BC2 | BH_NoFilter | 0.273 [0.221, 0.326] | 0.235 [0.205, 0.265] | −0.038 [−0.088, 0.011] | 0.739 [0.694, 0.785] | 0.742 [0.707, 0.778] | 0.003 [−0.042, 0.048] | 12 | 12 |

**Table S20. Semi-synthetic against simulated data by design factor.** Mean over the 43 configurations with 95% confidence intervals across replicates. Sample size uses every scenario of each approach, 50 to 200 samples in semi-synthetic and 50 to 500 in simulated data. Effect size and signal type use the common scenarios.

| Level | MCC semi-synthetic | FDR semi-synthetic | MCC simulated | FDR simulated |
|---|---|---|---|---|
| ***Total sample size*** | | | | |
| 50 | 0.381 [0.355, 0.407] | 0.075 [0.060, 0.089] | 0.276 [0.217, 0.335] | 0.109 [0.053, 0.165] |
| 100 | 0.569 [0.532, 0.606] | 0.109 [0.096, 0.122] | 0.465 [0.425, 0.504] | 0.110 [0.074, 0.145] |
| 200 | 0.685 [0.645, 0.724] | 0.125 [0.081, 0.169] | 0.551 [0.460, 0.642] | 0.179 [0.080, 0.279] |
| 300 | | | 0.648 [0.578, 0.717] | 0.130 [0.046, 0.214] |
| 500 | | | 0.690 [0.643, 0.737] | 0.146 [0.126, 0.166] |
| ***Effect size*** | | | | |
| 1.0 | 0.293 [0.251, 0.335] | 0.091 [0.065, 0.118] | 0.244 [0.170, 0.317] | 0.122 [0.092, 0.152] |
| 1.5 | 0.514 [0.496, 0.532] | 0.103 [0.067, 0.139] | 0.412 [0.353, 0.471] | 0.136 [0.067, 0.206] |
| 2.0 | 0.656 [0.607, 0.704] | 0.111 [0.068, 0.153] | 0.519 [0.485, 0.554] | 0.098 [0.079, 0.118] |
| 2.5 | 0.717 [0.692, 0.741] | 0.106 [0.073, 0.139] | 0.547 [0.465, 0.630] | 0.174 [0.062, 0.285] |
| ***Signal type*** | | | | |
| joint | 0.697 [0.660, 0.735] | 0.109 [0.082, 0.136] | 0.601 [0.559, 0.644] | 0.151 [0.072, 0.231] |
| prevalence only | 0.392 [0.359, 0.425] | 0.096 [0.086, 0.106] | 0.260 [0.212, 0.308] | 0.114 [0.088, 0.139] |

**Table S21. Real and simulated stool tables.** The real stool genus table (203 samples, 85 genera) against 20 tables simulated without signal from the fitted model for the same genera and number of samples. Pair correlations are Spearman correlations between the relative abundances of genera present in at least 10% of the samples.

| Property | Real stool | SparseDOSSA 2 |
|---|---|---|
| Zero counts (%) | 60.7 | 64.9 |
| Coefficient of variation of read depth | 0.89 | 0.69 |
| Median genera observed per sample | 34.0 | 30.0 |
| Median Shannon index | 1.90 | 1.82 |
| Slope of log variance on log mean | 1.67 | 1.69 |
| Median absolute pair correlation | 0.10 | 0.02 |
| Median Bray–Curtis dissimilarity | 0.47 | 0.52 |
| Mean spread of log10 abundance in carriers | 0.54 | 0.49 |

**Table S22. Profile of every configuration across conditions.** MCC / empirical FDR in each condition. Genus, OTU and simulated refer to the main detection design. Prevalence only, unidirectional, single extreme and confounder are at genus level, and the IBD community at genus level with 50 samples. Real cohorts give, for 95 women against 108 men of HMP (sex) and for 48 patients with IBD against 19 non-IBD controls of HMP2 (IBD), the number of genera called with the true labels and, in brackets, the percentage of 200 label permutations with at least one call. A calibrated configuration makes a call in about 5% of permutations.

| Tool | Configuration | Genus | OTU | Simulated | Prevalence only | Unidirectional | Single extreme | Confounder unadjusted | Confounder adjusted | IBD | Real cohorts, sex / IBD, calls (% of permutations with a call) |
|---|---|---|---|---|---|---|---|---|---|---|---|
| LinDA | Pseudo_0.1 | 0.68 / 0.06 | 0.63 / 0.04 | 0.64 / 0.09 | 0.59 / 0.04 | 0.95 / 0.08 | 0.87 / 0.21 | 0.76 / 0.36 | 0.92 / 0.05 | 0.52 / 0.04 | 1 (7) / 1 (8) |
| MaAsLin 2 | TMM_LOG | 0.68 / 0.04 | 0.61 / 0.03 | 0.64 / 0.08 | 0.54 / 0.03 | 0.91 / 0.15 | 1.00 / 0.00 | 0.77 / 0.36 | 0.91 / 0.04 | 0.49 / 0.03 | 1 (2.5) / 1 (5) |
| LinDA | Imputation | 0.66 / 0.06 | 0.60 / 0.06 | 0.62 / 0.11 | 0.56 / 0.02 | 0.76 / 0.34 | 0.12 / 0.97 | 0.68 / 0.44 | 0.82 / 0.17 | 0.50 / 0.07 | 1 (3.5) / 1 (7.5) |
| MaAsLin 3 | TSS_PLOG_Median | 0.65 / 0.03 | 0.61 / 0.04 | 0.63 / 0.06 | 0.58 / 0.02 | 0.90 / 0.15 | 0.19 / 0.95 | 0.76 / 0.33 | 0.87 / 0.06 | 0.50 / 0.02 | 0 (5) / 1 (9) |
| LinDA | Pseudo_0.5 | 0.65 / 0.05 | 0.62 / 0.03 | 0.62 / 0.10 | 0.52 / 0.03 | 0.95 / 0.08 | 0.91 / 0.15 | 0.72 / 0.39 | 0.88 / 0.08 | 0.51 / 0.04 | 1 (8) / 1 (8) |
| MaAsLin 3 | TSS_PLOG | 0.65 / 0.06 | 0.62 / 0.05 | 0.64 / 0.10 | 0.58 / 0.03 | 0.77 / 0.32 | 0.15 / 0.96 | 0.73 / 0.40 | 0.86 / 0.11 | 0.48 / 0.03 | 1 (3.5) / 1 (5) |
| MaAsLin 2 | TSS_LOG * | 0.64 / 0.06 | 0.60 / 0.06 | 0.62 / 0.10 | 0.54 / 0.03 | 0.75 / 0.34 | 0.12 / 0.97 | 0.68 / 0.45 | 0.84 / 0.15 | 0.51 / 0.02 | 1 (2) / 1 (4.5) |
| Wilcoxon | TSS | 0.64 / 0.07 | 0.59 / 0.05 | 0.62 / 0.10 | 0.57 / 0.04 | 0.73 / 0.38 | 0.16 / 0.96 | 0.73 / 0.39 | | 0.47 / 0.02 | 0 (3) / 1 (5) |
| LinDA | Adaptive * | 0.63 / 0.06 | 0.59 / 0.07 | 0.62 / 0.10 | 0.52 / 0.02 | 0.76 / 0.34 | 0.12 / 0.97 | 0.66 / 0.46 | 0.82 / 0.17 | 0.51 / 0.03 | 1 (8) / 1 (8) |
| LDM | CLR | 0.63 / 0.04 | 0.61 / 0.04 | 0.61 / 0.10 | 0.48 / 0.02 | 0.96 / 0.06 | 1.00 / 0.00 | 0.76 / 0.37 | 0.94 / 0.03 | 0.49 / 0.03 | 1 (2.5) / 1 (3) |
| LDM | Omni3 | 0.63 / 0.07 | 0.56 / 0.13 | 0.60 / 0.12 | 0.53 / 0.03 | 0.62 / 0.49 | 0.03 / 0.99 | 0.60 / 0.55 | 0.76 / 0.25 | 0.46 / 0.02 | 1 (6.5) / 1 (5) |
| LinDA | Adaptive_NoWinsor | 0.63 / 0.06 | 0.59 / 0.05 | 0.62 / 0.09 | 0.51 / 0.03 | 0.79 / 0.31 | 0.15 / 0.96 | 0.67 / 0.45 | 0.82 / 0.17 | 0.50 / 0.04 | 1 (7.5) / 1 (9) |
| MaAsLin 2 | CLR_NONE | 0.61 / 0.05 | 0.59 / 0.09 | 0.58 / 0.13 | 0.45 / 0.01 | 0.56 / 0.57 | 0.67 / 0.43 | 0.68 / 0.47 | 0.94 / 0.06 | 0.51 / 0.05 | 1 (4) / 1 (7.5) |
| limma-voom | TMM_voom | 0.61 / 0.09 | 0.50 / 0.27 | 0.55 / 0.15 | 0.48 / 0.06 | 0.54 / 0.56 | 1.00 / 0.00 | 0.57 / 0.54 | 0.80 / 0.22 | 0.47 / 0.21 | 1 (5) / 2 (23.5) |
| LDM | PA | 0.61 / 0.06 | 0.54 / 0.14 | 0.60 / 0.08 | 0.55 / 0.02 | 0.70 / 0.35 | 0.04 / 0.99 | 0.65 / 0.47 | 0.77 / 0.15 | 0.45 / 0.03 | 0 (6) / 1 (2.5) |
| LDM | Arcsin | 0.60 / 0.08 | 0.57 / 0.10 | 0.58 / 0.12 | 0.48 / 0.03 | 0.64 / 0.47 | 0.04 / 0.98 | 0.61 / 0.54 | 0.78 / 0.22 | 0.48 / 0.02 | 1 (3.5) / 0 (3) |
| ZicoSeq | MultiLink | 0.60 / 0.07 | 0.60 / 0.06 | 0.60 / 0.07 | 0.43 / 0.08 | 0.96 / 0.06 | 0.45 / 0.67 | 0.67 / 0.41 | 0.86 / 0.05 | 0.48 / 0.13 | 1 (9) / 1 (9) |
| LinDA | Pseudo_1 | 0.60 / 0.04 | 0.61 / 0.03 | 0.60 / 0.10 | 0.43 / 0.02 | 0.96 / 0.07 | 0.91 / 0.15 | 0.70 / 0.42 | 0.88 / 0.08 | 0.50 / 0.04 | 1 (8) / 1 (6) |
| ZicoSeq | NoPosterior | 0.60 / 0.04 | 0.64 / 0.05 | 0.59 / 0.07 | 0.47 / 0.04 | 0.94 / 0.05 | 0.80 / 0.31 | 0.73 / 0.37 | 0.80 / 0.09 | 0.43 / 0.05 | 1 (4) / 1 (6.5) |
| MaAsLin 3 | TSS_LOG_Median * | 0.60 / 0.03 | 0.53 / 0.04 | 0.60 / 0.03 | 0.49 / 0.03 | 0.97 / 0.03 | 1.00 / 0.00 | 0.60 / 0.49 | 0.86 / 0.01 | 0.42 / 0.02 | 0 (6.5) / 0 (0.5) |
| LDM | Omni * | 0.59 / 0.09 | 0.55 / 0.13 | 0.58 / 0.12 | 0.47 / 0.04 | 0.62 / 0.50 | 0.03 / 0.99 | 0.60 / 0.56 | 0.77 / 0.24 | 0.46 / 0.02 | 1 (5.5) / 0 (5) |
| Wilcoxon | CLR | 0.59 / 0.06 | 0.43 / 0.19 | 0.53 / 0.12 | 0.45 / 0.03 | 0.48 / 0.61 | 0.66 / 0.42 | 0.69 / 0.43 | | 0.46 / 0.01 | 0 (2.5) / 1 (5) |
| limma-voom | TMMwsp_voom | 0.58 / 0.09 | 0.39 / 0.35 | 0.54 / 0.18 | 0.45 / 0.03 | 0.43 / 0.65 | 0.89 / 0.16 | 0.61 / 0.52 | 0.80 / 0.19 | 0.42 / 0.29 | 0 (5) / 2 (29.5) |
| MaAsLin 3 | CLR_NONE | 0.58 / 0.04 | 0.53 / 0.07 | 0.53 / 0.09 | 0.45 / 0.02 | 0.85 / 0.19 | 0.81 / 0.30 | 0.58 / 0.53 | 0.87 / 0.03 | 0.42 / 0.03 | 0 (7) / 0 (1.5) |
| MaAsLin 3 | CLR_NONE_Median | 0.58 / 0.02 | 0.55 / 0.03 | 0.55 / 0.06 | 0.45 / 0.02 | 0.95 / 0.03 | 1.00 / 0.00 | 0.61 / 0.49 | 0.87 / 0.02 | 0.40 / 0.02 | 0 (5) / 0 (0.5) |
| MaAsLin 3 | TSS_LOG | 0.57 / 0.09 | 0.43 / 0.18 | 0.57 / 0.09 | 0.49 / 0.05 | 0.64 / 0.45 | 0.03 / 0.99 | 0.54 / 0.59 | 0.74 / 0.22 | 0.41 / 0.03 | 0 (6.5) / 0 (2) |
| DESeq2 | LRT_poscounts | 0.54 / 0.14 | 0.52 / 0.10 | 0.46 / 0.23 | 0.35 / 0.12 | 0.68 / 0.47 | 0.87 / 0.21 | 0.57 / 0.56 | 0.73 / 0.30 | 0.43 / 0.17 | 2 (28.5) / 4 (24.5) |
| DESeq2 | Wald_poscounts * | 0.54 / 0.16 | 0.50 / 0.13 | 0.45 / 0.25 | 0.36 / 0.17 | 0.67 / 0.48 | 0.83 / 0.27 | 0.56 / 0.56 | 0.73 / 0.31 | 0.46 / 0.34 | 2 (33) / 5 (60.5) |
| ZicoSeq | Default * | 0.52 / 0.12 | 0.60 / 0.06 | 0.56 / 0.07 | 0.32 / 0.16 | 0.95 / 0.06 | 0.55 / 0.57 | 0.67 / 0.42 | 0.78 / 0.04 | 0.41 / 0.06 | 1 (12.5) / 2 (7) |
| Wilcoxon | Rarefied | 0.51 / 0.08 | 0.38 / 0.05 | 0.52 / 0.10 | 0.35 / 0.04 | 0.73 / 0.37 | 0.10 / 0.98 | 0.62 / 0.48 | | 0.35 / 0.01 | 0 (3) / 1 (2.5) |
| edgeR | TMM_LRT | 0.50 / 0.34 | 0.50 / 0.39 | 0.49 / 0.32 | 0.32 / 0.37 | 0.69 / 0.46 | 0.58 / 0.61 | 0.57 / 0.61 | 0.78 / 0.31 | 0.38 / 0.51 | 3 (90.5) / 4 (80) |
| LDM | Freq | 0.49 / 0.08 | 0.50 / 0.11 | 0.52 / 0.10 | 0.30 / 0.04 | 0.64 / 0.45 | 0.06 / 0.98 | 0.61 / 0.54 | 0.72 / 0.19 | 0.36 / 0.02 | 1 (5.5) / 0 (5) |
| ALDEx2 | Wilcox_128 | 0.46 / 0.01 | 0.34 / 0.00 | 0.43 / 0.04 | 0.25 / 0.01 | 0.90 / 0.10 | 1.00 / 0.00 | 0.75 / 0.26 | | 0.38 / 0.00 | 0 (0) / 0 (0.5) |
| edgeR | TMM_QL * | 0.43 / 0.57 | 0.48 / 0.47 | 0.42 / 0.59 | 0.27 / 0.65 | 0.57 / 0.60 | 0.38 / 0.83 | 0.47 / 0.70 | 0.65 / 0.50 | 0.31 / 0.71 | 12 (99.5) / 19 (100) |
| ANCOM-BC2 | BH | 0.42 / 0.01 | 0.41 / 0.01 | 0.39 / 0.01 | 0.16 / 0.01 | 0.97 / 0.01 | 1.00 / 0.00 | 0.75 / 0.29 | 0.86 / 0.00 | 0.28 / 0.01 | 0 (0.5) / 0 (0.5) |
| ALDEx2 | Welch_128 * | 0.41 / 0.00 | 0.27 / 0.00 | 0.42 / 0.03 | 0.20 / 0.00 | 0.90 / 0.06 | 1.00 / 0.00 | 0.72 / 0.27 | 0.83 / 0.00 | 0.32 / 0.00 | 0 (0) / 0 (0.5) |
| ALDEx2 | Welch_32 | 0.41 / 0.00 | 0.28 / 0.00 | 0.42 / 0.03 | 0.20 / 0.00 | 0.91 / 0.05 | 1.00 / 0.00 | 0.72 / 0.26 | 0.81 / 0.00 | 0.31 / 0.00 | 0 (0.5) / 0 (0.5) |
| ALDEx2 | Welch_128_IQLR | 0.41 / 0.00 | 0.28 / 0.00 | 0.42 / 0.02 | 0.20 / 0.00 | 0.93 / 0.02 | 1.00 / 0.00 | 0.72 / 0.26 | | 0.32 / 0.00 | 0 (0) / 0 (0.5) |
| ANCOM-BC2 | BH_StrucZero | 0.40 / 0.01 | 0.38 / 0.01 | 0.36 / 0.01 | 0.15 / 0.01 | 0.97 / 0.01 | 1.00 / 0.00 | 0.74 / 0.29 | 0.86 / 0.00 | 0.24 / 0.01 | 0 (0.5) / 0 (0.5) |
| MaAsLin 2 | TMM_NONE | 0.37 / 0.03 | 0.41 / 0.01 | 0.40 / 0.03 | 0.17 / 0.03 | 0.81 / 0.11 | 0.75 / 0.00 | 0.78 / 0.20 | 0.73 / 0.01 | 0.15 / 0.00 | 0 (1) / 0 (3) |
| ANCOM-BC2 | Holm * | 0.35 / 0.00 | 0.26 / 0.00 | 0.34 / 0.00 | 0.08 / 0.00 | 0.96 / 0.00 | 1.00 / 0.00 | 0.77 / 0.23 | 0.82 / 0.00 | 0.23 / 0.00 | 0 (0) / 0 (0) |
| ANCOM-BC2 | Holm_NoFilter | 0.31 / 0.65 | 0.28 / 0.62 | 0.28 / 0.66 | 0.06 / 0.85 | 0.68 / 0.48 | 0.29 / 0.90 | 0.46 / 0.68 | 0.53 / 0.59 | 0.28 / 0.62 | 9 (100) / 11 (100) |
| ANCOM-BC2 | BH_NoFilter | 0.27 / 0.74 | 0.12 / 0.85 | 0.23 / 0.75 | 0.07 / 0.86 | 0.53 / 0.65 | 0.20 / 0.95 | 0.38 / 0.75 | 0.44 / 0.70 | 0.22 / 0.77 | 11 (100) / 28 (100) |

**Table S23. Read depth that differs between groups, with a signal.** Stool genera with joint signals implanted into 10% of the genera in both directions, effect sizes 1.5 and 2.0, at 100 and 200 samples, four replicates (16 datasets per level), with the case samples thinned as in Table S14. Mean MCC / mean empirical FDR.

| Tool | Configuration | 100% of reads | 75% of reads | 50% of reads | 25% of reads |
|---|---|---|---|---|---|
| LinDA | Pseudo_0.1 | 0.85 / 0.08 | 0.80 / 0.17 | 0.61 / 0.43 | 0.29 / 0.77 |
| MaAsLin 2 | TMM_LOG | 0.88 / 0.06 | 0.87 / 0.08 | 0.80 / 0.20 | 0.66 / 0.43 |
| LinDA | Imputation | 0.81 / 0.15 | 0.80 / 0.15 | 0.79 / 0.16 | 0.76 / 0.24 |
| MaAsLin 3 | TSS_PLOG_Median | 0.86 / 0.07 | 0.86 / 0.07 | 0.83 / 0.08 | 0.77 / 0.19 |
| LinDA | Pseudo_0.5 | 0.84 / 0.11 | 0.78 / 0.19 | 0.55 / 0.51 | 0.28 / 0.78 |
| MaAsLin 3 | TSS_PLOG | 0.85 / 0.14 | 0.85 / 0.13 | 0.82 / 0.19 | 0.68 / 0.39 |
| MaAsLin 2 | TSS_LOG * | 0.85 / 0.13 | 0.84 / 0.13 | 0.83 / 0.17 | 0.74 / 0.31 |
| Wilcoxon | TSS | 0.82 / 0.16 | 0.82 / 0.16 | 0.78 / 0.23 | 0.65 / 0.44 |
| LinDA | Adaptive * | 0.80 / 0.16 | 0.81 / 0.14 | 0.79 / 0.16 | 0.76 / 0.24 |
| LDM | CLR | 0.88 / 0.07 | 0.79 / 0.17 | 0.50 / 0.55 | 0.19 / 0.83 |
| LDM | Omni3 | 0.83 / 0.17 | 0.83 / 0.15 | 0.82 / 0.17 | 0.75 / 0.31 |
| LinDA | Adaptive_NoWinsor | 0.81 / 0.16 | 0.81 / 0.13 | 0.78 / 0.18 | 0.78 / 0.18 |
| MaAsLin 2 | CLR_NONE | 0.88 / 0.08 | 0.79 / 0.16 | 0.44 / 0.56 | 0.11 / 0.88 |
| limma-voom | TMM_voom | 0.84 / 0.11 | 0.73 / 0.21 | 0.36 / 0.65 | 0.13 / 0.88 |
| LDM | PA | 0.84 / 0.12 | 0.84 / 0.12 | 0.83 / 0.14 | 0.80 / 0.19 |
| LDM | Arcsin | 0.83 / 0.17 | 0.83 / 0.16 | 0.80 / 0.19 | 0.74 / 0.32 |
| ZicoSeq | MultiLink | 0.83 / 0.10 | 0.81 / 0.11 | 0.68 / 0.25 | 0.41 / 0.58 |
| LinDA | Pseudo_1 | 0.84 / 0.11 | 0.77 / 0.20 | 0.53 / 0.53 | 0.27 / 0.79 |
| ZicoSeq | NoPosterior | 0.87 / 0.05 | 0.88 / 0.05 | 0.86 / 0.05 | 0.73 / 0.28 |
| MaAsLin 3 | TSS_LOG_Median * | 0.82 / 0.02 | 0.83 / 0.05 | 0.73 / 0.18 | 0.39 / 0.62 |
| LDM | Omni * | 0.82 / 0.15 | 0.82 / 0.16 | 0.81 / 0.18 | 0.74 / 0.31 |
| Wilcoxon | CLR | 0.82 / 0.13 | 0.73 / 0.17 | 0.38 / 0.58 | 0.04 / 0.90 |
| limma-voom | TMMwsp_voom | 0.78 / 0.19 | 0.71 / 0.16 | 0.36 / 0.58 | 0.11 / 0.88 |
| MaAsLin 3 | CLR_NONE | 0.81 / 0.04 | 0.75 / 0.14 | 0.60 / 0.38 | 0.35 / 0.69 |
| MaAsLin 3 | CLR_NONE_Median | 0.82 / 0.01 | 0.83 / 0.03 | 0.77 / 0.10 | 0.52 / 0.47 |
| MaAsLin 3 | TSS_LOG | 0.75 / 0.16 | 0.77 / 0.16 | 0.68 / 0.31 | 0.42 / 0.66 |
| DESeq2 | LRT_poscounts | 0.76 / 0.22 | 0.69 / 0.33 | 0.57 / 0.49 | 0.41 / 0.67 |
| DESeq2 | Wald_poscounts * | 0.75 / 0.23 | 0.70 / 0.31 | 0.55 / 0.50 | 0.39 / 0.67 |
| ZicoSeq | Default * | 0.82 / 0.08 | 0.78 / 0.14 | 0.61 / 0.34 | 0.36 / 0.60 |
| Wilcoxon | Rarefied | 0.77 / 0.15 | 0.77 / 0.14 | 0.76 / 0.11 | 0.67 / 0.15 |
| edgeR | TMM_LRT | 0.76 / 0.30 | 0.73 / 0.33 | 0.66 / 0.43 | 0.63 / 0.48 |
| LDM | Freq | 0.85 / 0.12 | 0.84 / 0.12 | 0.83 / 0.14 | 0.81 / 0.18 |
| ALDEx2 | Wilcox_128 | 0.74 / 0.00 | 0.71 / 0.04 | 0.57 / 0.22 | 0.30 / 0.72 |
| edgeR | TMM_QL * | 0.65 / 0.49 | 0.63 / 0.50 | 0.56 / 0.59 | 0.53 / 0.61 |
| ANCOM-BC2 | BH | 0.83 / 0.00 | 0.81 / 0.01 | 0.76 / 0.03 | 0.58 / 0.25 |
| ALDEx2 | Welch_128 * | 0.69 / 0.00 | 0.68 / 0.03 | 0.57 / 0.20 | 0.31 / 0.69 |
| ALDEx2 | Welch_32 | 0.69 / 0.00 | 0.65 / 0.03 | 0.55 / 0.20 | 0.32 / 0.68 |
| ALDEx2 | Welch_128_IQLR | 0.67 / 0.00 | 0.64 / 0.02 | 0.60 / 0.13 | 0.38 / 0.59 |
| ANCOM-BC2 | BH_StrucZero | 0.81 / 0.00 | 0.79 / 0.01 | 0.73 / 0.03 | 0.57 / 0.25 |
| MaAsLin 2 | TMM_NONE | 0.71 / 0.01 | 0.67 / 0.02 | 0.64 / 0.10 | 0.59 / 0.22 |
| ANCOM-BC2 | Holm * | 0.76 / 0.00 | 0.75 / 0.00 | 0.67 / 0.00 | 0.57 / 0.04 |
| ANCOM-BC2 | Holm_NoFilter | 0.63 / 0.46 | 0.61 / 0.46 | 0.55 / 0.48 | 0.42 / 0.58 |
| ANCOM-BC2 | BH_NoFilter | 0.48 / 0.64 | 0.48 / 0.64 | 0.46 / 0.63 | 0.30 / 0.76 |

**Table S24. Data sources and community tables.** Samples with fewer than 1,000 reads were removed, and one sample was kept per subject. Genera present in at least 5% of the samples and OTUs present in at least 10% were kept.

| Community | Resolution | Source | Samples in source | Subjects in source | Features before filter | Subjects kept | Features kept | Reads kept (%) | Zeros (%) | Median depth | Median prevalence (%) | Reads in top feature (%) |
|---|---|---|---|---|---|---|---|---|---|---|---|---|
| Stool | Genus | HMP V3-V5, stool (HMP16SData V35) | 319 | 210 | 151 | 203 | 85 | 99.9 | 60.7 | 4478 | 24.1 | 46.3 |
| Stool | OTU | HMP V3-V5, stool (HMP16SData V35) | 319 | 210 | 11145 | 203 | 2697 | 81.7 | 76.0 | 3755 | 18.7 | 2.4 |
| IBD | Genus | HMP2 inflammatory bowel disease cohort, mucosal biopsies of CD and UC patients (HMP2Data IBD16S) | 132 | 59 | 285 | 56 | 168 | 99.6 | 65.5 | 16678 | 25.0 | 30.5 |
| IBD | OTU | HMP2 inflammatory bowel disease cohort, mucosal biopsies of CD and UC patients (HMP2Data IBD16S) | 132 | 59 | 650 | 56 | 195 | 96.3 | 63.9 | 16448 | 26.8 | 15.1 |
