## Supplementary Text for "A configuration-resolved benchmark of differential abundance analysis methods for human gut 16S rRNA microbiome data"

### Text S1. Supplementary results and methods

A configuration-resolved benchmark of differential abundance analysis methods for human gut 16S rRNA microbiome data

Nader Abdelmalek, Carola Di Meo, Giovanni Fiorito, Sergio Uzzau and Alessandro Tanca

#### 1. Supplementary results

##### 1.1 Leading configurations and package defaults

Eight of the eleven leading configurations are linear models on log or log-ratio scales that handle zeros with a small pseudo-count or by imputation, namely LinDA, MaAsLin 2 with log transformation, MaAsLin 3 with pseudo-log transformation and LDM on CLR values. The other three are the rank test on proportions, which makes no distributional assumption, and two LDM tests that include presence and absence. The log scale stabilises the variance of abundances that span orders of magnitude, and a small pseudo-count keeps zeros from dominating the estimates.

In semi-synthetic data, the LinDA default ranked below both options between which it switches. It chose imputation where read depth was associated with group, where a pseudo-count gave a higher MCC, and a pseudo-count elsewhere, where imputation gave a higher MCC. Implantation raises the read depth of the cases, and the switch therefore chose imputation in most datasets with signals in one direction or a bloom, where imputation was liberal. The MaAsLin 3 default, with median comparison and log transformation, traded recall for error control.

ALDEx2 was insensitive to its settings. The number of Monte Carlo instances and the denominator hardly changed its MCC, and 32 instances instead of 128 cut run time four- to fivefold. The interquartile log-ratio denominator lowered the FDR under signals in one direction (Table S12), and the Wilcoxon test gave a higher MCC than the Welch test from the same fit. Configuration sensitivity is thus itself tool dependent. Run time differed by more than three orders of magnitude at OTU level and did not increase the MCC (Figure S14, Table S2). LDM, which assesses each test by permutation (1), and ANCOM-BC2, which repeats its fit over a range of pseudo-counts (2), were the slowest.

##### 1.2 Error control

With the 60 null stool datasets of the 20-replicate design and Benjamini–Hochberg adjustment at 0.05, the configurations other than DESeq2, edgeR and ANCOM-BC2 without its sensitivity analysis stayed within the binomial uncertainty of the nominal rate of datasets with a call (Tables S4 and S6). At OTU level, limma-voom with TMM also made calls in 5 of 12 null datasets, and in simulated data MaAsLin 2 CLR_NONE made calls in 4 of 20 (Table S6). The few calls of the other configurations occurred mostly in the same datasets, often in opposite directions for the configurations on proportions and for limma-voom, the pattern expected from a chance compositional difference between two random groups. DESeq2 made calls in more than a third of the null datasets and edgeR in most of them, and

ANCOM-BC2 without its sensitivity analysis called about a third of the OTU table on average. DESeq2 and edgeR shrink the dispersion of each feature towards a common trend, which sparse taxa with many zeros probably fit poorly. ANCOM-BC2 leaves zeros out of its primary fit, and its sensitivity analysis removes the taxa whose result depends on the pseudo-count used to replace them.

LinDA with a pseudo-count of 0.5 tracked the nominal level at every threshold (Figure S1, Table S7). It corrects the compositional bias of each coefficient with the mode of all coefficients and centres its test on the unaffected majority of taxa (3). The LinDA default and LinDA with imputation stayed above the nominal level at every threshold, as did MaAsLin 2 and MaAsLin 3 on proportions without median comparison, and at OTU level the intervals of both TSS_LOG configurations excluded it. Once small studies and small effects were excluded, the empirical FDR of MaAsLin 2 and MaAsLin 3 on proportions without median comparison was 0.12 to 0.14 at genus level at a nominal 0.05. ALDEx2 and ANCOM-BC2 stayed well below the nominal level, because their internal stringency rather than the threshold limits their calls. ALDEx2 averages p-values over Monte Carlo draws from a Dirichlet posterior, which is conservative for low-count taxa.

**1.3 Contribution of design, tool and configuration**

The design term of the MCC was led by effect size, signal type and sample size (Figure 3A). MaAsLin 2 TMM_NONE fits normalised counts without a log transformation, and the few large counts of each taxon dominate its fit. It made few false calls but also found few true ones, whereas the two ANCOM-BC2 configurations without sensitivity analysis made many (Figure S3). Without edgeR and these two ANCOM-BC2 configurations, the four most liberal configurations, the tool explained 5.0% and the configuration 2.2% of the variation in MCC at genus level, and 5.5% and 1.2% of the variation in empirical FDR, of which 69% lay between replicates. At OTU level the tool still explained 20% of the variation in empirical FDR and the configuration 5%. Replicates, which differ in the subjects and signals drawn, explained about 15% of the variation in MCC at genus level and 2% at OTU level. With about 30 times more features, the MCC of a single OTU dataset rests on many more calls and varies less around its scenario, and the replicate term is therefore smaller at OTU level. The tool and configuration terms were nearly unchanged with 20 replicates, whereas the replicate term increased (Table S8). These contributions reflect the settings tested here rather than every possible one.

At genus level, the gap between the best and the median tool exceeded the gain from configuring each tool in all but one of the 24 scenarios (Table S9). At OTU level, the gain exceeded the gap in 9 of the 24 scenarios, five of them at 200 samples. OTU tables are sparser, and the handling of zeros and the reference matter more there. In large OTU-level studies, configuration deserves as much attention as the tool.

The out-of-sample gain over the default could not be distinguished from zero for MaAsLin 2, LDM, DESeq2 and ALDEx2 at genus level, and at OTU level the gains were mostly larger, led by ANCOM-BC2 (Table S11). For edgeR, the likelihood-ratio test gave a higher MCC and was less liberal than the quasi-likelihood default. For ANCOM-BC2, choosing by MCC alone selected a configuration without sensitivity analysis in a quarter to a third of the stool datasets and raised the FDR. Keeping the sensitivity analysis and replacing Holm with Benjamini–Hochberg adjustment gave most of the gain at an FDR near zero. Holm adjustment controls the family-wise error rate (4), which is stricter than the FDR targeted by the other tools.

A configuration chosen at 200 samples also outperformed the default at 50 samples for most tools (Table S18).

A report of ANCOM-BC2 error control is uninterpretable unless it states whether the sensitivity analysis was active, because switching it off moves the FDR from the lowest in the benchmark to the highest without changing the model. Median comparison in MaAsLin 3 removes the shift that compositionality imposes on all taxa when some of them change (5). Posterior sampling in ZicoSeq is designed to stabilise the proportions of rare taxa (6), yet it lowered the MCC. In LinDA, a pseudo-count of 1 compresses the log-ratio differences of low-count taxa and lowered the MCC relative to 0.1 and 0.5 (Table S10). Comparing tools at their defaults can thus hide the settings that make a tool liberal or conservative.

**1.4 Sample size and effect size**

At genus level, the mean MCC of the leading configurations increased nearly twofold between 50 and 200 samples (Figure 3B, Figure S4A). The standard error of a difference decreases with the square root of the sample size, and a genus present in 20% of the samples is seen in only about five samples per group in a study of 50. ALDEx2 and ANCOM-BC2 started lower, and with 200 samples both remained below the leading configurations at 100. The MCC of the leading configurations increased more than twofold from an effect size of 1.0 to 2.5, that is, from a 2.7-fold to a 12-fold change in the non-zero counts (Figure 3C, Figure S5A).

The order of the configurations changed little with sample size or effect size. At genus level in semi-synthetic data, MaAsLin 2 and LinDA had the best mean ranks and ALDEx2 the worst, but the intervals of the leading tools overlapped (Figure S15A). The leading tools differed at OTU level and in simulated data (Figure S15B and C). Single datasets agreed less in small studies, and Kendall's W at 50 samples was about half its value at larger sizes in semi-synthetic data and a fifth in simulated data (Figure S15D and E).

Over all sample sizes, 24 configurations exceeded the nominal FDR in simulated data, against 11 on the scenarios common to both approaches and 10 in the semi-synthetic data (Tables S3 and S19). Semi-synthetic data showed the same increase of the FDR with sample size (Figure S6A and B). MaAsLin 3 with median comparison, LDM CLR and LinDA with a pseudo-count of 0.1 stayed near the nominal level at every sample size and effect size in semi-synthetic data. At 500 simulated samples, LinDA with a fixed pseudo-count and LDM CLR reached an empirical FDR of about 0.1. MaAsLin 3 with median comparison, ZicoSeq without posterior sampling and ANCOM-BC2 with sensitivity analysis stayed below 0.04 (Figure S6). The compositional changes behind this increase are real in relative but not in absolute abundance (7, 8).

The empirical FDR is the product of the proportion of datasets with at least one call and the FDR of the non-empty lists (9). For MaAsLin 2 TSS_LOG, the second term increased most with sample size (Figure S7B and D). ALDEx2 and ANCOM-BC2 made a call in fewer than half of the datasets at 50 samples (Figure S7A and C), which keeps their FDR at zero but leaves most small studies without a result. At OTU level, the FDR of the LDM omnibus test increased strongly with effect size under joint signals (Figure S8B). For methods on proportions, the FDR increases with study size and signal strength, which are the conditions under which results are trusted most.

Placed on one scale, the design factors differed in cost (Figure S16A and B). For the default of each tool at genus level, a small effect size cost the most MCC, followed by prevalence-only signals, an ignored confounder and a halved sample size (Figure S16A). Unbalanced groups, unidirectional signals and a larger fraction of differential features cost little. Only an ignored confounder and unidirectional signals raised the FDR markedly (Figure S16B). The MCC is therefore limited mainly by the study and the signal, and false discoveries arise mainly from the direction of the signal and from omitted covariates.

##### 1.5 Signal structures and confounding

Under signals in one direction, MaAsLin 3 TSS_PLOG_Median, LinDA with imputation and the LinDA default were not protected by their references, unlike MaAsLin 3 with median comparison on log or CLR values and LinDA with a fixed pseudo-count (Figure 4B, Table S12).

Under a single bloom at implantation depth, median comparison did not protect MaAsLin 3 on the pseudo-log scale, and LinDA with imputation failed as well. MaAsLin 2 TMM_LOG, limma-voom with TMM, LDM CLR, MaAsLin 3 with median comparison on log or CLR values, ANCOM-BC2 with sensitivity analysis and ALDEx2 kept the FDR near the nominal level (Figure 4). The bloom raised the read depth of the cases about 200-fold. Configurations on proportions called the bloom together with multiple unchanged taxa.

With 20% of the features changed, LinDA with imputation also exceeded the nominal FDR, whereas a fixed pseudo-count kept LinDA close to it (Figure S9A and B).

At a correlation of 0.6, the confounded taxa differ between groups slightly more than the true taxa, and an unadjusted analysis cannot distinguish them (Figure S11A and B). With adjustment, DESeq2, edgeR, limma-voom, MaAsLin 3 TSS_LOG and the LDM tests on proportions, on arcsine-root proportions and the omnibus test remained liberal, largely because their FDR was already above 0.1 without confounding. LinDA with imputation and its default, MaAsLin 2 TSS_LOG and the LDM presence-absence test also stayed at 0.15 to 0.17 at genus level, although their FDR without confounding was 0.07 or less (Figure S11C and D, Table S15). This agrees with confounder-aware benchmarks of observational microbiome studies (10).

##### 1.6 Twenty replicates, fixed read depth and unequal depth

With 20 replicates per scenario, the order of the configurations was unchanged and the same configurations led (Table S4). Every configuration that exceeded the nominal FDR with four replicates still did, except the Wilcoxon test on rarefied counts. Five more configurations now clearly controlled the FDR, namely MaAsLin 2 TMM_LOG and TMM_NONE, MaAsLin 3 TSS_LOG_Median and TSS_PLOG_Median and LDM CLR. Seven configurations newly exceeded it, namely LinDA with imputation, LDM Omni3 and arcsine, both limma-voom configurations, the Wilcoxon test on CLR values and MaAsLin 3 TSS_LOG. Their empirical FDR was 0.063 to 0.068, except limma-voom at 0.095 and 0.097, about twice the nominal level. Resampling every sample to its depth before implantation preserved the order of the configurations under balanced signals (Table S5).

Under signals in one direction at fixed depth, ANCOM-BC2 with sensitivity analysis and MaAsLin 3 with median comparison on log or CLR values led the ten configurations that could not be distinguished from the nominal FDR, followed by ZicoSeq (Table S5). This order rests on four replicates. Nearly all extra calls of LinDA, LDM CLR and the other liberal configurations were unchanged taxa called decreased. Against a truth defined by relative abundance, the configurations on proportions made almost no false calls, and whether their calls count as errors depends on whether the question concerns relative or absolute abundance.

Read depth can differ between groups for technical reasons alone. Thinning lowered the depth of the cases without changing their expected composition (Figure S10A, Table S14). At 2-fold, MaAsLin 3 with median comparison on log or CLR values called taxa in a quarter to half of the null datasets, possibly because detection, which its prevalence model tests, depends on read depth. LDM on proportions and on presence and LinDA with imputation stayed within the binomial uncertainty of the nominal rate up to 4-fold, and only the Wilcoxon test on rarefied counts stayed near it at 10-fold, in line with the robustness of rarefaction to uneven sequencing effort in diversity analyses (11).

With a signal and the reads of the cases halved, which gave a median depth ratio of about 1.8 because implantation had raised the depth of the cases, LinDA with a fixed pseudo-count lost MCC and its empirical FDR increased several-fold. LinDA with imputation kept its MCC and empirical FDR close to their values at equal depth, although that FDR was above the nominal level (Figure S10B, Table S23). In the depth split, the two halves differed 2.4-fold in depth. The configurations with a fixed pseudo-count on a log-ratio scale and limma-voom made calls in nearly all of these splits but rarely in random splits (Table S14). LinDA with imputation, MaAsLin 3 TSS_PLOG_Median, ANCOM-BC2 with sensitivity analysis and the Wilcoxon test on rarefied counts made calls in fewer of them, and ZicoSeq without posterior sampling in half. With 200 samples, nearly the whole cohort, the halves differed 2.5-fold, and most configurations made calls in every depth split. Only ANCOM-BC2 with sensitivity analysis, MaAsLin 2 TMM_NONE, the LDM tests on proportions, on arcsine-root proportions and the LDM omnibus test, and the Wilcoxon test on rarefied counts made calls in fewer than half of these splits. Depth alone does not explain all of these calls. LinDA with imputation and the LDM presence-absence test made calls in at most 3 of 20 null datasets thinned to a 4-fold difference, but in 19 or 20 of the 20 depth splits at 200 samples and in none of the random splits. Shallow and deep stool samples therefore appear to differ in composition, and part of these calls may be real.

In the shift design with 30% of the genera shifted at fixed depth, MaAsLin 3 with median comparison on log or CLR values and ANCOM-BC2 with sensitivity analysis kept the empirical FDR at 0.13 or less. The ZicoSeq default reached a negative MCC from 50% of the genera shifted (Figure S10C and D, Table S13). MaAsLin 3 TSS_PLOG and CLR_NONE, MaAsLin 2 TSS_LOG and TMM_LOG and the Wilcoxon test on proportions degraded more gently and kept a positive MCC.

**1.7 Prevalence signals and effect-size estimation**

In the HMP2 IBD multi-omics data, MaAsLin 3 corroborated previously reported associations and identified 77% of them through prevalence, not abundance (5). The pseudo-log transformation lets the abundance model of MaAsLin 3 detect prevalence-only signals, because it keeps the zeros on the log scale (Figure 5A and B, Table S16). ANCOM-BC2 has no prevalence component and is unsuitable when presence-absence shifts are expected, despite its low FDR on abundance signals.

Effect sizes matter for meta-analysis and biomarker prioritisation, which compare them across studies. edgeR and DESeq2 estimate the log ratio of mean counts, which equals the change among carriers when only abundance changes (Figure 5C, Table S17). DESeq2 should therefore match edgeR, and its lower value is a bias whose source we did not isolate. MaAsLin 3 with pseudo-log transformation and ALDEx2 also recovered less than a third of the implanted change, like LinDA, MaAsLin 2 and limma-voom. ANCOM-BC2 and MaAsLin 3 on log or CLR values leave zeros out of the abundance model and estimate the change among carriers.

With 100 samples, the called features of LinDA and MaAsLin 2 came close to the implanted change, against less than a third over all tested features under abundance-only signals (Figure 5D, Figure S17, Table S17). The sign of the estimate was correct for most implanted features (Figure 5E), and ANCOM-BC2 and MaAsLin 3 without pseudo-log transformation made the fewest sign errors. Among the true positives of the main design, the sign was correct for 99.9% under joint signals, and for every configuration for at least 98.7% (MaAsLin 3 TSS_LOG at genus level). Under prevalence-only signals, it was correct for 54% to 84% of the true positives of ANCOM-BC2 and 58% to 65% of those of MaAsLin 3 on log or CLR values, against more than 99.8% for the other configurations with a coefficient. These models fit abundance on non-zero values only, and a change in presence alone gives their coefficient no defined direction. Effect sizes of taxa called in small studies are inflated by this selection and should be reported with this caveat. The configurations best suited to estimation are not those that maximise MCC.

##### 1.8 Taxonomic levels and the IBD community

MaAsLin 2 TMM_LOG uses the same normalisation and scale as limma-voom without exceeding the nominal level at OTU level, and the excess of limma-voom therefore probably comes from how it estimates the variance of sparse taxa (Figure 6B). MaAsLin 3 with median comparison on log or CLR values, LinDA with a fixed pseudo-count, LDM CLR and ANCOM-BC2 with sensitivity analysis stayed near the nominal level at both taxonomic levels (Figure 6B and C). Run time increased about 20-fold from genus to OTU level (Figure S14, Table S2).

The dependence of differential abundance results on cohort type (12) affected the magnitude of performance in the IBD community rather than the choice of method (Figure 6D).

##### 1.9 Semi-synthetic and simulated data

SparseDOSSA 2 reproduced the prevalence, mean abundance and variance of each genus (Figure S13A, Figure S18, Table S21), but the correlations between genera were close to zero in simulated tables (Figure S13B). About a quarter of the simulated true genera were rarer than the 10% prevalence that implantation requires (Figure S13C). Nearly all of each implanted change reached the counts, whereas part of each simulated change was lost when the reads were drawn (Figure S13D and E). Under joint signals, the prevalence part of each simulated signal reached the counts in full, and the loss came from the abundance among carriers (Figure S18H). Both differences lower power.

On the 24 common scenarios, the interval of the difference in mean MCC excluded zero for 40 of the 43 configurations (Figure S19A, Table S19). The loss came almost entirely from recall and was largest for prevalence-only signals (Figure S13F, Table S20). The empirical FDR increased slightly (Figure S13G, Figure S19B), because the same compositional false positives form a larger fraction of shorter lists. The same seven configurations exceeded the nominal FDR in both approaches, and edgeR and ANCOM-BC2 without sensitivity analysis made calls in most null datasets of both (Figure S19C). The semi-synthetic approach, which keeps the real correlations and places signals on observed genera, is the more realistic of the two (10).

##### 1.10 Real cohorts

A configuration that controls the FDR at 0.05 should make a call in about 5% of permutations, and up to 15 of 200 are compatible with this level (Figure 7, Table S22). With the sex labels, Akkermansia was the only genus called by more than half of the configurations, and with the IBD labels the Lachnospiraceae ND3007 group (Table S22). Different tools are known to return different taxa on the same data (12), and the configuration within a tool does so as well. In both cohorts, both configurations of edgeR and DESeq2 and the two ANCOM-BC2 configurations without sensitivity analysis made a call in 24.5% to 100% of permutations (Figure 7C and E). Both limma-voom configurations did so in 23.5% and 29.5% of the IBD permutations but in 5% of the sex permutations, and the ZicoSeq default in 12.5% of the sex permutations. Several LinDA, ZicoSeq and MaAsLin 3 configurations made calls in 8% to 9% of permutations in one or both cohorts, slightly above the range compatible with the nominal level. The lists of the clearly liberal configurations on the true labels are therefore largely noise. Across configurations, the proportion of permutations with a call correlated with the benchmark FDR (Spearman 0.72 for sex and 0.74 for IBD). It also correlated with the proportion of null datasets without signal with a call (0.60 for stool and 0.57 for the IBD community). The IBD contrast itself was weak in these rectal biopsies. A Wilcoxon test on proportions gave 11 of 164 genera with an unadjusted p-value below 0.05, close to the 8 expected by chance, and one after Benjamini–Hochberg adjustment, the Lachnospiraceae ND3007 group. Thirty of the 48 patients had Crohn's disease.

With the first 20 permutations, the correlations with the benchmark FDR were 0.70 for sex and 0.65 for IBD, and the proportions from 20 and 200 permutations correlated at 0.88 and 0.70. The same six configurations were clearly liberal. With 20 permutations, the proportion is resolved only in steps of 5%, and up to three of 20 are compatible with the nominal level. With the sex labels, LDM Omni3, MaAsLin 3 CLR_NONE and the ZicoSeq default made calls in 4 of 20 permutations. With 200, LDM Omni3 and MaAsLin 3 CLR_NONE decreased to 6.5% and 7%. Conversely, limma-voom TMM_voom made calls in 2 of 20 IBD permutations and in 23.5% of 200. About 200 permutations are therefore needed to separate mildly liberal configurations from sampling noise.

#### 2. Supplementary methods

##### 2.1 Data sources and preparation

Stool profiles of the healthy adults of the Human Microbiome Project (HMP) V3–V5 16S rRNA data (13) were obtained with the HMP16SData package (14). Mucosal biopsy profiles of patients with Crohn's disease or ulcerative colitis from the HMP2 inflammatory bowel disease (IBD) cohort (15) were obtained with the HMP2Data package (16). Samples with fewer than 1,000 reads were removed. One sample was kept per subject, the earliest visit for HMP and the rectal biopsy when available for HMP2, making the samples of a dataset independent. Only public, de-identified data were used, and no ethics approval was required.

Two taxonomic levels were built from each table. For the genus table, the counts of the operational taxonomic units (OTUs) assigned to a genus were summed. OTUs without a genus assignment were grouped under their lowest classified rank, for example Lachnospiraceae_unclassified. Genera present in at least 5% of the samples were kept. For the OTU table, OTUs present in at least 10% of the samples were kept. The OTU table is the finest taxonomic level available in these data. Amplicon sequence variants from denoising pipelines are finer still, usually come from deeper sequencing and were not tested here.

The stool data comprise 203 subjects, 85 genera and 2,697 OTUs. Zero counts account for 60.7% of the genus table and 76.0% of the OTU table, and the median depth is 4,478 reads in the genus table and 3,755 in the OTU table. The IBD community (56 subjects, 168 genera) is a dysbiotic mucosal community with a median depth of 16,678 reads. Stool was analysed at both taxonomic levels and the IBD community at genus level (Figure 1A, Table S24). The stool genus table of the same 203 subjects, 95 women and 108 men, was also analysed with its sex labels, and HMP2 rectal biopsies with their IBD labels.

**2.2 Signal implantation**

For each dataset, the required number of subjects was drawn without replacement from the community table and assigned at random to cases and controls. A fraction of the features, 10% by default, was selected to carry a signal. Only features present in at least 10% of the drawn samples, and in at least three samples, were eligible. Half of the selected features were increased in the cases and half decreased, unless the design specified a unidirectional signal. The effect size $e$ has the same definition as in SparseDOSSA 2 (17), and the two approaches use the same nominal effects. How much of each effect reaches the counts differs between them.

An abundance effect multiplies the non-zero counts of a feature in the case samples by $\exp(\pm e)$, with rounding and a minimum of one. It never changes whether the feature is present in a sample. A prevalence effect shifts the log-odds of presence in the cases by $\pm e$. With $p_0$ the prevalence of the feature in the drawn samples, the target prevalence is $p_1 = \text{expit}(\text{logit}(p_0) \pm e)$. For an increase, a fraction $(p_1 - p_0)/(1 - p_0)$ of the case samples in which the feature is absent receive it. For a decrease, a fraction $(p_0 - p_1)/p_0$ of the case samples that carry the feature lose it. The number of changed samples is the rounded expected number, with a minimum of one. A new non-zero value is the relative abundance of the feature in a randomly chosen sample that carries it, multiplied by the depth of the receiving sample. For signals acting on prevalence alone, a feature had to be present in at least three cases and absent from at least three cases, and every true feature could therefore change. A joint signal applies both effects to the same features.

All other counts were left as observed. The mean-variance relationship, the zero pattern and the correlations of the unaffected features are those of the real subjects drawn. Apart from the change in read depth that the implanted counts cause, the only systematic difference between the two groups is the implanted effect. The implanted features are those that truly differ between groups, and they are the true positives against which every method is scored. The empirical FDR therefore measures recovery of the implanted features, not false biological associations on every abundance scale. Replicates of a scenario differ in the subjects drawn, the assignment to groups, the features that carry the signal and the direction of each effect. When the sample size approaches the number of subjects, replicates have most of their subjects in common, and their spread then reflects the assignment and the placement of the signal more than the sampling of subjects.

#### 2.3 Simulated data

Simulated communities were generated with SparseDOSSA 2 (17), fitted to the stool genus table (85 genera, 203 samples) with fit_SparseDOSSA2, using the default penalty on feature correlations ($\lambda = 1$) and at most 100 iterations of the expectation-maximisation algorithm. Datasets were simulated for the fitted genera themselves. SparseDOSSA 2 draws the read depth of each sample from a log-normal distribution fitted to the real table and rescales it to the median depth of that table (4,478 reads). The simulated depths therefore vary between samples, though less than the real ones (Table S21). Abundance and prevalence effects were set with the same definitions as for the semi-synthetic data. SparseDOSSA 2 applies them to the latent abundance of each genus, converts the abundances to proportions and then draws the reads of each sample from a multinomial distribution, whereas implantation changes the observed counts. Table S21 also compares other properties of tables simulated without signal with the real one. The simulated data also reach total sample sizes of 300 and 500, divided equally between cases and controls, which the 203 stool subjects cannot. They were used for the main detection and null designs, as a check that the ranking does not depend on how the truth is generated. The secondary designs were run on real profiles, which keep the real community around each implanted signal.

#### 2.4 Comparison of semi-synthetic and simulated data

The two approaches were compared at three levels. First, the real stool genus table was compared with 20 tables simulated without signal for the same genera and number of samples. Per genus, we compared prevalence, mean relative abundance and the standard deviation of the log relative abundance among the samples that carry the genus. Per sample, we compared read depth, observed genera and the Shannon index, and per pair of samples the Bray–Curtis dissimilarity. Per pair of genera present in at least 10% of the samples, we compared the Spearman correlation of their relative abundances.

Second, every dataset of the main detection design of both approaches was regenerated from its seed, and the change that each signal produced in the counts was measured at the sample sizes common to both approaches. For each true feature, the realised change is the difference between cases and controls in the log-odds of presence and in the mean log relative abundance among carriers, divided by the nominal effect. A ratio of one means that the full signal reached the counts. The Wilcoxon test on proportions was rerun on the regenerated datasets to relate recall to the prevalence of the true genera in the controls.

Third, the configurations were compared on the 24 genus-level scenarios common to both approaches. For each configuration, the difference between approaches in mean MCC, recall and FDR is reported with a Welch 95% interval across replicates. Agreement is the Spearman correlation across configurations, computed over all common scenarios and within each scenario.

#### 2.5 Designs

Table S1 gives the complete grid of designs. Every design is a two-group comparison that varies one factor around the main design. Sample size is the total number of samples in a dataset, divided equally between cases and controls unless the group-balance design specifies otherwise. Every tool was called at a nominal FDR level of 0.05 except in the threshold design. The signal type is either joint, which changes both how much of a taxon is present among its carriers and how often it is detected, or prevalence-only, which changes only how often it is detected. The main design was run on stool at genus and OTU level. Unless stated, the secondary designs were run on stool at both levels with 100 and 200 samples, joint signals and effect sizes of 1.5, 2.0 and 2.5. The designs on stool were the following.

**Main detection.** Effect sizes of 1.0, 1.5, 2.0 and 2.5 were crossed with 50, 100 and 200 samples and with both signal types, giving 24 scenarios per taxonomic level. Signals were implanted into 10% of the features.

**Null.** No signal was implanted, at 50, 100 and 200 samples, and every call is a false discovery.

**Group balance.** The case group comprised 50% or 30% of the samples (50/50 and 30/70). Under the unbalanced split, the smaller group is the case group.

**Fraction of features with a signal.** Signals were implanted into 5%, 10% (as in the main design) or 20% of the features.

**Signal direction.** Balanced signals, in which taxa increased and decreased in equal numbers, were compared with signals in one direction, in which all changed taxa increased.

**Single extreme change.** A single feature was increased with an effect size of 10, the limiting case of a signal in one direction.

**Effect-size estimation.** A signal on abundance alone or a joint signal was implanted at effect sizes of 1.0 to 2.5, and the coefficient of every implanted feature that a configuration tested was compared with the implanted effect. ZicoSeq, LDM and the Wilcoxon test return no coefficient on an interpretable scale and were not included.

**Confounding effects.** Confounding was introduced through a continuous covariate correlated with the exposure. The strength of the association was set with a Pearson correlation coefficient $r$ of 0, 0.3 and 0.6 at 200 samples, where $r = 0$ leaves the covariate independent of the exposure. A second, disjoint set of features, as large as the true set, was multiplied by $\exp(\pm e \cdot z)$ in every sample, with $z$ the standardised covariate, and these features have no group effect of their own. Each scenario was analysed twice, first with the covariate included as a fixed effect and then without it. These are referred to as the adjusted and unadjusted analyses. The covariate entered every tool that supports covariates as a fixed effect.

**Nominal FDR level.** Calls were made at nominal levels of 0.05, 0.10 and 0.25 with both signal types. Tools whose fit depends on the threshold were refitted at each level.

Two further data sources ran a subset of these designs. The IBD community was analysed at genus level with 50 samples, effect sizes of 1.5 and 2.5 and both signal types, the scenarios that it has in common with the stool main design, and with a null design at 50 samples. The simulated data ran the main detection and null designs at genus level with total sample sizes of 50, 100, 200, 300 and 500, giving 25 to 250 samples per group.

**2.6 Tools and configurations**

Eleven tools were evaluated across 43 tool-configuration combinations (Table S2). The configurations of a tool vary the settings that it exposes to the user, namely normalisation, transformation, reference, zero handling, test and filtering. The package default of each tool is marked in all tables and figures. edgeR has no single default test, and the quasi-likelihood test recommended in its user guide was treated as its default, whereas limma-voom and the Wilcoxon test have no single default. Prevalence and abundance filters inside the tools were switched off, and every configuration therefore tested the same features, with the exceptions stated below. A feature was called when its adjusted p-value was at or below the nominal level.

MaAsLin 2 was run with total sum scaling (TSS), centred log-ratio (CLR) or TMM normalisation followed by an optional log transformation, retaining the configurations TSS_LOG (the default), CLR_NONE, TMM_LOG and TMM_NONE (18, 19). MaAsLin 2 applies the log transformation to all values after replacing zeros by half the smallest non-zero value of each feature, and CLR normalisation adds a pseudo-count of one. Both are fixed in the package. The default 10% prevalence filter was switched off.

MaAsLin 3 was run with TSS or CLR normalisation. Under TSS, the plain-log and pseudo-log transformations were applied, whereas under CLR no further transformation was applied. Each of these was run with and without median comparison, yielding the six configurations TSS_LOG, TSS_LOG_Median (the default), TSS_PLOG, TSS_PLOG_Median, CLR_NONE and CLR_NONE_Median (5). MaAsLin 3 models prevalence and abundance separately and reports an abundance test, a prevalence test and a joint test that combines them. Under the plain-log transformation its abundance model is fitted on non-zero values only, with zeros excluded rather than replaced. The pseudo-log option restores a pseudo-count to zeros before the log transformation and applies to the abundance model alone, as the prevalence model is not compatible with pseudo-log. For the pseudo-log configurations we therefore scored the abundance test, and for the remaining configurations the joint test. TSS_LOG and TSS_PLOG therefore differ in the test as well as in the transformation, and Table S16 gives the abundance, prevalence and joint tests of every configuration. MaAsLin 3 computes CLR values with the natural logarithm and log values with base 2, and coefficients were converted to the log2 scale accordingly. Median comparison tests each abundance coefficient against the median coefficient instead of zero. The pseudo-log configurations were fitted with the pseudo-log transformation of the package and the abundance model only.

edgeR was run with TMM normalisation under the likelihood-ratio and quasi-likelihood tests (20, 21). At fixed read depth, two further fits followed the edgeR user guide more closely, the quasi-likelihood test after filtering with filterByExpr and with robust dispersion and quasi-likelihood estimation, and the quasi-likelihood fit of edgeR 3. DESeq2 was also run with the settings its vignette recommends for sparse counts, the likelihood-ratio test with a minimum mean of $10^{-6}$, without outlier replacement and with the glmGamPoi dispersion fit. These fits are reported in Table S5 and are not among the 43 configurations.

ALDEx2 was run in four configurations, the Welch t test with 128 (the default) and with 32 Monte Carlo instances, the Wilcoxon test with 128 Monte Carlo instances and the Welch t test with the interquartile log-ratio set as the denominator, giving Welch_128, Welch_32, Wilcox_128 and Welch_128_IQLR (22). The expected Benjamini–Hochberg adjusted p-value across instances was used. The run with 32 instances tests whether a run four times shorter loses MCC.

ANCOM-BC2 (2) adds no pseudo-count in its primary fit and leaves zeros out. Its sensitivity analysis repeats the test with zeros replaced by pseudo-counts from 0.01 to 0.5 and discards taxa whose result depends on the pseudo-count, and it thus acts as a filter on the results. Three configurations keep the sensitivity analysis active, namely the default with Holm adjustment (4), Benjamini–Hochberg adjustment and Benjamini–Hochberg adjustment with detection of structural zeros by their asymptotic lower bound. Two further configurations disable it, with either adjustment, and are labelled NoFilter. The 10% prevalence filter of the package stays active in all five, and taxa removed by it count as not called. When the model stopped because a taxon had zero variance within a group, that taxon was removed and the model refitted, as the package advises.

LinDA (3) fits linear models to CLR-transformed counts and corrects the compositional bias with the mode of the regression coefficients. It was run with its adaptive default, which uses imputation of zeros when log library size is associated with a term of the model ($P \leq 0.1$) and a pseudo-count of 0.5 otherwise, with a fixed pseudo-count of 0.1, 0.5 or 1, with imputation and with the adaptive default without winsorisation. The pseudo-count series isolates the effect of this choice on the same model. In MicrobiomeStat 1.4 the adaptive switch never takes effect, because the function sets the zero handling to “Imputation” and then tests for “imputation”. The default of the CRAN release therefore always uses a pseudo-count of 0.5 and gives the same result as LinDA with a fixed pseudo-count of 0.5. The development version of the package on GitHub (1.5.1) sets the value in lower case and applies the switch. We applied the switch as documented before each call in every analysis of this study, and both adaptive configurations follow the method as described (3).

ZicoSeq (6) is a permutation-based linear model with a reference-based correction for compositionality and posterior sampling of the underlying proportions. It was run with its default square-root link and posterior sampling (Default), with three links (powers of 0.25, 0.5 and 0.75) combined by their maximum (MultiLink), and without posterior sampling (NoPosterior). Each run used 99 permutations, the package default. ZicoSeq estimates the FDR from the permuted statistics of all features together, and its adjusted p-values are therefore not limited by the number of permutations. A pilot with 999 permutations returned the same calls at seven times the run time. Features that are constant after winsorisation cannot be tested by ZicoSeq and were counted as not called.

LDM (1) tests each feature by permutation within a linear decomposition model. One fit returns tests on relative abundances and on arcsine-root transformed relative abundances, their omnibus (the default), a test of presence and absence over rarefactions (23) and the omnibus of all three (24). Each was scored as a configuration. LDM on CLR-transformed counts with a pseudo-count of 0.5 (25) was run as a separate fit, which tests the effect of each taxon against the median effect of all taxa (the package default). LDM controls the FDR with a sequential permutation procedure tied to the nominal level and was refitted at each level of the threshold design. The number of permutations was limited to 5,000, because the default limit increases with the number of features and did not finish on OTU tables in a pilot.

DESeq2 (26) was run with the Wald test (the default) and the likelihood-ratio test. Size factors were estimated with the poscounts method, because the default estimator needs a feature without zeros, which 16S tables rarely contain. Coefficients are the maximum-likelihood estimates, without shrinkage. Independent filtering and the Cook's distance cutoff were kept at their defaults. Switching them off gave identical calls in a pilot.

limma-voom (27, 28) was run with TMM normalisation or its variant for tables with many zeros (TMMwsp) (20), voom precision weights and the moderated t test.

The Wilcoxon rank-sum test (29, 30) was applied to each feature after one of three preparations, namely relative abundances, CLR values with a pseudo-count of one or counts rarefied to the smallest library of the dataset.

Most tools applied Benjamini–Hochberg (BH) correction at the stated threshold (31). ANCOM-BC2 uses Holm adjustment in its default, ALDEx2 reports the expected BH value across Monte Carlo instances, and LDM and ZicoSeq adjust permutation p-values. In the confounding design the covariate was added as a fixed effect in every tool that supports it, and ALDEx2 was then fitted with its generalised linear model. The ALDEx2 Wilcoxon and interquartile log-ratio configurations and the Wilcoxon rank-sum test have no adjusted form and were not run in the adjusted analysis.

##### 2.7 Metrics

Performance was quantified by the Matthews correlation coefficient (MCC) as the primary metric, computed per dataset from the confusion matrix of implanted features against the features each method called significant (32). With TP, FP, TN and FN the numbers of true positives, false positives, true negatives and false negatives, $MCC = (TP \cdot TN - FP \cdot FN)/\sqrt{[(TP + FP)(TP + FN)(TN + FP)(TN + FN)]}$. It ranges from −1 to 1, where 0 is the value expected from a random call set and 1 is a perfect call set. The MCC was chosen as the primary metric because it uses all four cells of the confusion matrix and stays informative when classes are imbalanced (33). In differential abundance analysis the true features are a small fraction of those tested, and accuracy is then dominated by the true negatives, whereas the F1 score ignores them (33). Datasets in which a configuration ran but returned no significant features leave MCC undefined, because no feature is called positive. These were assigned an MCC of 0, the value of a non-informative classifier, and their proportion is reported. A recent simulation benchmark of differential abundance methods also used the MCC as its main metric (34).

Recall, F1 score and false discovery rate (FDR) were reported alongside it. The false discovery proportion of a dataset is FP/(TP + FP), set to 0 when nothing is called, and its mean across datasets is the empirical FDR. Recall is TP/(TP + FN) and the F1 score is 2TP/(2TP + FP + FN). For the estimation analysis, the coefficient of each implanted feature was converted to the log2 scale and compared against its implanted true effect, *e*/ln 2. The median ratio of estimate to truth is reported, because its distribution is heavy tailed, together with the proportion of implanted features whose estimate has the correct sign.

##### 2.8 Statistical analysis

Metrics were first averaged over the scenarios of a design or stratum within each replicate. They are reported as the mean of the four replicate values with a 95% confidence interval from the t distribution. The empirical FDR of a configuration was classed as controlled when the upper limit of its interval was at or below the nominal level, as exceeded when the lower limit was above it and as not distinguishable from the nominal level otherwise. These intervals were computed for each configuration separately and were not adjusted for the number of configurations. Proportions of null datasets with a call are given with exact binomial (Clopper–Pearson) intervals.

The variation of MCC, recall and FDR across scenarios, replicates and configurations was decomposed, over all scenarios, configurations and replicates of the main detection design, into sums of squares for scenario, tool, configuration within tool, the interactions of scenario with tool and with configuration, and the residual between replicates, which reflects the random draw of subjects, groups and signals. Each term is reported as a percentage of the total. Only scenario and replicate cells in which every configuration returned a result were used, which keeps the layout balanced and the terms additive. In the main detection design, where sample size, effect size and signal type are fully crossed, the scenario term was further divided into these factors and their interactions, and the interaction of tool with scenario into the corresponding terms. The decomposition was repeated without the three extreme configurations, MaAsLin 2 TMM_NONE and the two ANCOM-BC2 configurations without sensitivity analysis, and without the four configurations with the highest empirical FDR, the two edgeR configurations and the same two ANCOM-BC2 configurations.

To estimate the gain from changing the configuration of a tool without overfitting, the configuration was chosen on three replicates of a scenario and scored on the fourth, in rotation. The configuration with the highest mean MCC on the three replicates was selected, with ties resolved in favour of the default. The gain over the default is the mean difference on the held-out replicates, with a 95% interval across the held-out replicates. The interval is approximate, because the training sets of the rotations overlap. Within each scenario, the gain from choosing the configuration was compared with the gap between tools. The gain is the mean over tools of the difference between the chosen and the median configuration of each tool. The gap is the difference between the best and the median tool, each represented by its chosen configuration. To test whether a choice transfers, the configuration of each tool with the highest mean MCC on semi-synthetic stool genera was applied to OTU tables, to the simulated data and to the other communities. The configuration chosen at 200 samples was likewise applied at 50 samples.

Configurations were compared with a two-stage rank-based procedure. Within each scenario the configurations were ranked by MCC averaged over replicates, with rank 1 assigned to the best configuration and ties resolved by averaging. A Friedman test (35) was then applied to test the global hypothesis that all configurations have equal mean rank, with Kendall's W as the effect size. The Nemenyi post-hoc test (36) was used to identify which pairs of configurations differ, controlling the family-wise error rate across all pairwise comparisons (37). Because every configuration was evaluated on an identical set of scenarios, the Nemenyi test yields a single critical difference that applies to every pair. The paired difference across scenarios between each configuration and the best one was also tested with a t test and Holm adjustment (4). Tools were compared in the same way on every dataset of the main design, each represented by the configuration chosen on the other three replicates, overall and within each sample size and effect size.

The effects of sample size and effect size were summarised by averaging the main detection design over the other factors at each level, within each replicate. Estimation accuracy was summarised in the same way by effect size and by sample size, over all tested implanted features and over those that the configuration called. The second set shows the selection bias of estimates that pass a significance threshold.

Every dataset was also scored at ten nominal FDR levels from 0.001 to 0.5. For each configuration this gives the empirical FDR as a function of the nominal level, and the recall at the largest nominal level at which the empirical FDR stays at or below 0.05. For tools whose fit depends on the nominal level (ANCOM-BC2 with sensitivity analysis, LDM and DESeq2), the curve applies the lower thresholds to the fit made at 0.05 and is not used above 0.05. Their calibration at higher levels comes from the threshold design, in which they were refitted. The empirical FDR was also divided into the proportion of datasets with at least one call and the positive FDR, the mean false discovery proportion of the datasets with calls, whose product it is (9). The effect of each design factor was measured as the difference in mean MCC and FDR from the matched reference condition, with a Welch 95% interval across replicates. It was measured on the package default of each tool, or on limma-voom with TMM normalisation and the Wilcoxon test on proportions for the two tools without one.

Stool was compared with the IBD community on the four scenarios that they have in common. Agreement is the Spearman correlation across configurations of mean MCC, recall and FDR.

##### 2.9 Robustness analyses

Three further analyses tested whether the conclusions depend on the number of replicates, on read depth and on the breadth of a shift. All used genus-level data. First, the main detection design on stool and the null design on stool and IBD profiles were extended to 20 replicates per scenario, giving 480 main and 80 null datasets (60 stool and 20 IBD). The summaries, the FDR verdicts, the variance decomposition and the configuration choice were repeated on them, with the configuration chosen on 19 replicates and scored on the twentieth.

Second, implantation changes the read depth of the case samples, because an increase adds reads and a decrease removes them, whereas sequencing fixes the depth of each sample. The main detection, signal direction, single extreme change and null designs on stool genera were therefore analysed again after every sample, cases and controls alike, was resampled from a multinomial distribution to its depth before implantation. Each configuration was scored against two truths. The absolute truth is the implanted features, as everywhere else in this study. The relative truth adds every feature whose mean relative abundance in the cases changed by more than 0.2 on the natural-log scale through the implantation. We used the arithmetic mean of proportions, the fraction of the community that a taxon holds, because it needs no reference. Log-ratio methods estimate changes against a reference such as the geometric mean of all taxa, which neither truth matches exactly.

Third, three designs on stool genera separated read depth from the signal and extended signals in one direction to most of the community. In the depth design, the reads of every case sample were thinned binomially to 90%, 75%, 50%, 25% or 10%, which lowers the depth of the cases without changing their expected composition. Thinning was applied to null datasets (10 replicates at each of 100 and 200 samples) and, at 75%, 50% and 25%, to datasets with joint signals implanted into 10% of the genera at effect sizes of 1.5 and 2.0 (100 and 200 samples, four replicates). Each level was fitted with the same seeds as the unthinned dataset. In the depth split, 60, 100 or 200 real stool samples were divided into their shallower and deeper halves, and the same samples divided at random served as the control (20 draws each). In the shift design, 10%, 30%, 50% or 70% of the genera were decreased or increased together at an effect size of 1.5 (100 and 200 samples, four replicates), and every dataset was scored as implanted and at fixed read depth as above.

##### 2.10 Real cohorts

Every configuration was applied to two real comparisons at genus level, prepared like the benchmark tables. The first was the stool genus table of the 95 women and 108 men from HMP with their sex labels. Few genus-level differences are expected between the sexes (38), and this cohort therefore shows how liberal each configuration is on real data. The second was the HMP2 rectal biopsies of 48 patients with IBD (Crohn's disease or ulcerative colitis) and 19 non-IBD controls, one biopsy per subject, with 164 genera. These patients overlap with the IBD community used for implantation but not with the stool subjects on which the benchmark FDR was measured. A genus was called when its adjusted p-value was 0.05 or less, as in the benchmark. The labels of each cohort were permuted 200 times. On permuted labels every call is false, and the proportion of permutations with at least one call shows how often a configuration reports a difference where none exists. For a calibrated configuration this proportion is about 5%, and up to 15 of 200 permutations (7.5%) are compatible with this level by a one-sided binomial test. The proportion of permutations with a call and the number of calls on the true labels were compared with the mean empirical FDR of each configuration in the semi-synthetic main detection design at genus level. They were also compared with the proportion of datasets without signal in which the configuration made a call. This used the stool null design at 200 samples for the sex labels and the IBD community at 50 samples for the IBD labels. The estimated FDR reported for each configuration is the mean number of calls on permuted labels divided by the number of calls on the true labels, capped at one, and is not defined when nothing is called on the true labels. Agreement is the Spearman correlation across configurations. The first 20 permutations of each cohort are those of an earlier run with 20 permutations, which allows the two to be compared. Permuting the labels removes every difference between the groups, including the compositional shifts that a true signal causes, and the check therefore detects miscalibrated variance but not the compositional false discoveries studied in the benchmark.

##### 2.11 Implementation

The benchmark was implemented in R 4.5.2. Datasets were generated with HMP16SData 1.30.0 (14), HMP2Data 0.99.4 from GitHub (16) and SparseDOSSA2 0.99.2 (17). The tools were run as MaAsLin 2 1.24.1 (19), MaAsLin 3 1.2.0 (5), edgeR 4.8.2 (21), ALDEx2 1.42.0 (22), ANCOMBC 2.13.1 (2), a development version compatible with current CVXR releases, MicrobiomeStat 1.4 for LinDA (3), GUniFrac 1.9 for ZicoSeq (6), LDM 6.0.1 (1), DESeq2 1.50.2 (26) and limma 3.66.0 (28). Simulation seeds were fixed for every scenario and replicate. Each dataset and implanted feature set can therefore be regenerated exactly. The code and all result tables are deposited in Zenodo.

#### References


1. Hu YJ, Satten GA. Testing hypotheses about the microbiome using the linear decomposition model (LDM). Bioinformatics. 2020;36:4106-4115.

2. Lin H, Peddada SD. Multigroup analysis of compositions of microbiomes with covariate adjustments and repeated measures. Nat Methods. 2024;21:83-91.

3. Zhou H, He K, Chen J, Zhang X. LinDA: linear models for differential abundance analysis of microbiome compositional data. Genome Biol. 2022;23:95.

4. Holm S. A simple sequentially rejective multiple test procedure. Scand J Stat. 1979;6:65-70.

5. Nickols WA, Kuntz T, Shen J, et al. MaAsLin 3: refining and extending generalized multivariable linear models for meta-omic association discovery. Nat Methods. 2026;23:554-564. doi:10.1038/s41592-025-02923-9.

6. Yang L, Chen J. A comprehensive evaluation of microbial differential abundance analysis methods. Microbiome. 2022;10:130.

7. Gloor GB, Macklaim JM, Pawlowsky-Glahn V, Egozcue JJ. Microbiome datasets are compositional and this is not optional. Front Microbiol. 2017;8:2224.

8. Kumar MS, Slud EV, Okrah K, et al. Analysis and correction of compositional bias in sparse sequencing count data. BMC Genomics. 2018;19:799.

9. Storey JD. A direct approach to false discovery rates. J R Stat Soc B. 2002;64:479-498.

10. Wirbel J, Essex M, Forslund SK, Zeller G. A realistic benchmark for differential abundance testing and confounder adjustment in human microbiome studies. Genome Biol. 2024;25:247.

11. Schloss PD. Rarefaction is currently the best approach to control for uneven sequencing effort in amplicon sequence analyses. mSphere. 2024;9:e00354-23.

12. Nearing JT, Douglas GM, Hayes MG, et al. Microbiome differential abundance methods produce different results across 38 datasets. Nat Commun. 2022;13:342.

13. Human Microbiome Project Consortium. Structure, function and diversity of the healthy human microbiome. Nature. 2012;486:207-214.

14. Schiffer L, Azhar R, Shepherd L, et al. HMP16SData: efficient access to the Human Microbiome Project through Bioconductor. Am J Epidemiol. 2019;188:1023-1026.

15. Lloyd-Price J, Arze C, Ananthakrishnan AN, et al. Multi-omics of the gut microbial ecosystem in inflammatory bowel diseases. Nature. 2019;569:655-662.

16. Stansfield J, Smirnova E, Zhao N, et al. HMP2Data: 16s rRNA sequencing data from the Human Microbiome Project 2. R package version 0.99.4. https://github.com/dozmorovlab/HMP2Data. doi:10.18129/B9.bioc.HMP2Data.

17. Ma S, Ren B, Mallick H, et al. A statistical model for describing and simulating microbial community profiles. PLoS Comput Biol. 2021;17:e1008913.

18. Robinson MD, Oshlack A. A scaling normalization method for differential expression analysis of RNA-seq data. Genome Biol. 2010;11:R25.

19. Mallick H, Rahnavard A, McIver LJ, et al. Multivariable association discovery in population-scale meta-omics studies. PLoS Comput Biol. 2021;17:e1009442.

20. Robinson MD, McCarthy DJ, Smyth GK. edgeR: a Bioconductor package for differential expression analysis of digital gene expression data. Bioinformatics. 2010;26:139-140.

21. Chen Y, Chen L, Lun ATL, Baldoni PL, Smyth GK. edgeR v4: powerful differential analysis of sequencing data with expanded functionality and improved support for small counts and larger datasets. Nucleic Acids Res. 2025;53:gkaf018.

22. Fernandes AD, Reid JN, Macklaim JM, et al. Unifying the analysis of high-throughput sequencing datasets: characterizing RNA-seq, 16S rRNA gene sequencing and selective growth experiments by compositional data analysis. Microbiome. 2014;2:15.

23. Hu YJ, Lane A, Satten GA. A rarefaction-based extension of the LDM for testing presence-absence associations in the microbiome. Bioinformatics. 2021;37:1652-1657.

24. Zhu Z, Satten GA, Hu YJ. Integrative analysis of relative abundance data and presence-absence data of the microbiome using the LDM. Bioinformatics. 2022;38:2915-2917.

25. Hu Y, Satten GA. Compositional analysis of microbiome data using the linear decomposition model (LDM). bioRxiv. 2023. doi:10.1101/2023.05.26.542540 (preprint).

26. Love MI, Huber W, Anders S. Moderated estimation of fold change and dispersion for RNA-seq data with DESeq2. Genome Biol. 2014;15:550.

27. Law CW, Chen Y, Shi W, Smyth GK. voom: precision weights unlock linear model analysis tools for RNA-seq read counts. Genome Biol. 2014;15:R29.

28. Ritchie ME, Phipson B, Wu D, et al. limma powers differential expression analyses for RNA-sequencing and microarray studies. Nucleic Acids Res. 2015;43:e47.

29. Mann HB, Whitney DR. On a test of whether one of two random variables is stochastically larger than the other. Ann Math Stat. 1947;18:50-60.

30. Wilcoxon F. Individual comparisons by ranking methods. Biometrics Bull. 1945;1:80-83.

31. Benjamini Y, Hochberg Y. Controlling the false discovery rate: a practical and powerful approach to multiple testing. J R Stat Soc B. 1995;57:289-300.

32. Matthews BW. Comparison of the predicted and observed secondary structure of T4 phage lysozyme. Biochim Biophys Acta. 1975;405:442-451.

33. Chicco D, Jurman G. The advantages of the Matthews correlation coefficient (MCC) over F1 score and accuracy in binary classification evaluation. BMC Genomics. 2020;21:6.

34. Cappellato M, Baruzzo G, Di Camillo B. Investigating differential abundance methods in microbiome data: a benchmark study. PLoS Comput Biol. 2022;18:e1010467.

35. Friedman M. The use of ranks to avoid the assumption of normality implicit in the analysis of variance. J Am Stat Assoc. 1937;32:675-701.

36. Nemenyi PB. Distribution-free multiple comparisons. PhD thesis, Princeton University; 1963.

37. Demšar J. Statistical comparisons of classifiers over multiple data sets. J Mach Learn Res. 2006;7:1-30.

38. Falony G, Joossens M, Vieira-Silva S, et al. Population-level analysis of gut microbiome variation. Science. 2016;352:560-564.
